# *Thermoaminiphila catenidiffluenda* gen. nov., sp. nov.: A novel thermophilic, strictly anaerobic bacterium representing *Thermoaminiphilia* class nov., a newly described class thriving in hydrocarbon-rich habitats and biogas fermenters

**DOI:** 10.64898/2026.04.13.718153

**Authors:** Eva Maria Prem, Mathias Wunderer, Andja Mullaymeri, Julia Zoehrer, Zuzanna Dutkiewicz, Abhijeet Singh, Mahmoud M. Habashy, Anna Neubeck, Sepehr Shakeri Yekta, Christian Rinke, Andreas Otto Wagner

## Abstract

Axenic cultivation of novel bacterial lineages, referred to as “gold standard in microbiology”, remains challenging for fastidious or uncultured taxa due the challenges of replicating adequate growth conditions. We isolated strain PM69, a representative of the previously undescribed *Bacillota* class SHA-98, from a phenyl acid degrading, oligotrophic batch culture. By employing a broad spectrum of (anaerobic) culture techniques, biochemical, physiological, and genomic analyses, we characterised the strain as *Thermoaminiphila catenidiffluenda*, gen. nov., sp. nov., a thermophilic, strictly anaerobic, bacterium fermenting monosaccharides to acetate. Its motility, biofilm forming capacity, and ecological niche in biogas fermenters and hydrocarbon-associated habitats suggest adaptive strategies for harsh environments exhibiting e.g., high concentrations of aromatic compounds. This description of a new bacterial class not only expands the taxonomic diversity of phylum *Bacillota* but also provides insights into the metabolic versatility of yet uncultured microorganisms, with implications for carbon cycling and biotechnological applications.

**Graphical abstract:** 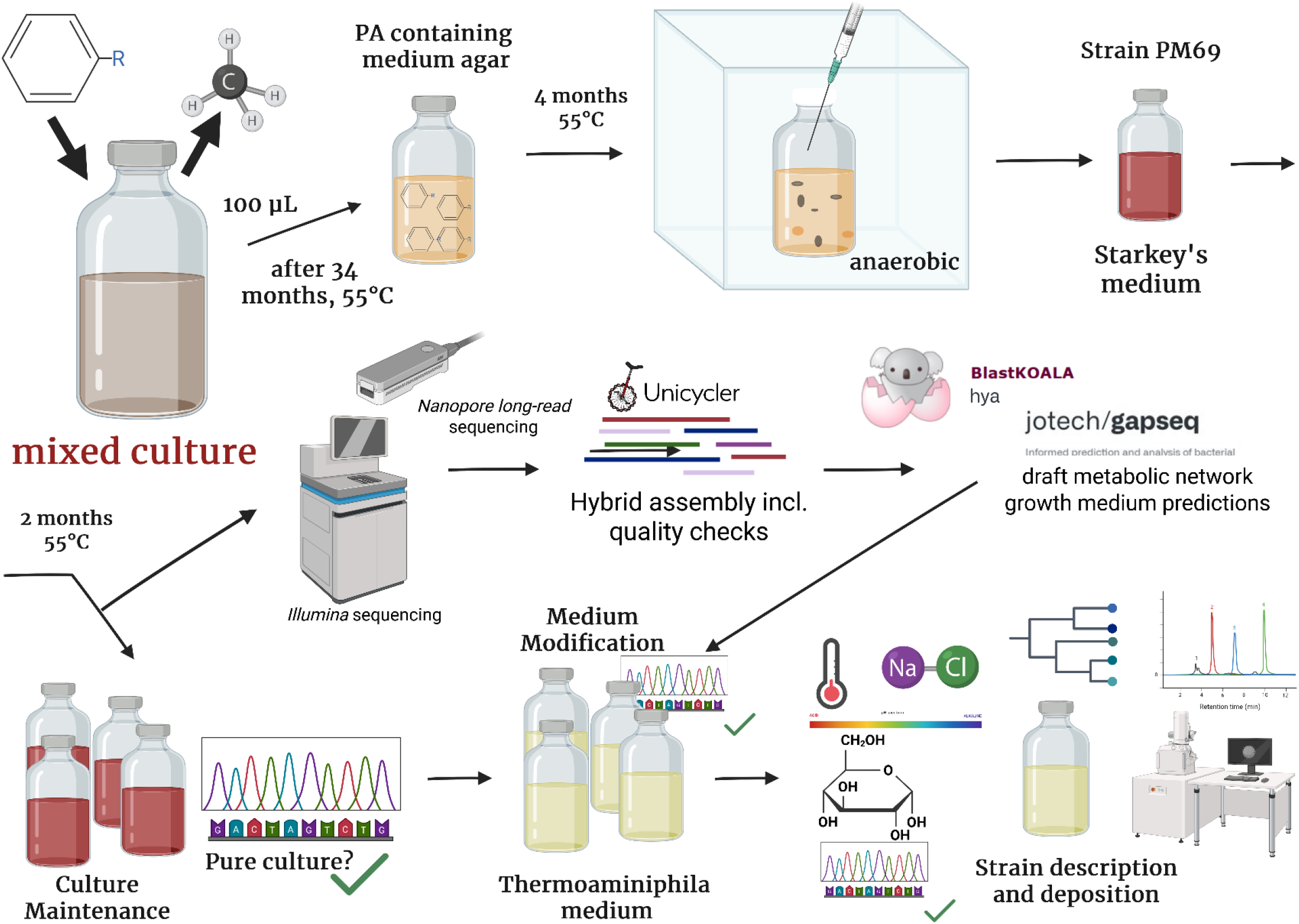

## 1. Introduction

The International Code of Nomenclature of Prokaryotes (*ICNP*) decided on using the suffix *-ota* for phylum names as proposed previously ^1,2^. In this regard, phylum *Firmicutes* was renamed *Bacillota* after its type genus *Bacillus* in 2021, and this change was implemented in the Genome Taxonomy Data Base (GTDB) Release 214 in 2023 ^3^. The GTDB classifies prokaryotic sequences obtained from the NCBI Assembly database by using relative evolutionary divergence (RED) for higher rank taxa and average nucleotide identity (ANI) for species-level clustering ^4^. The GTDB includes genomes of cultivated representatives as well as metagenome assembled genomes (MAGs). These genomes are either assigned placeholder names or named using *Seqcode* which provides a nomenclature for uncultivated procaryotes based on their genomic characteristics ^5^. The description of new cultured bacterial classes that are i) available as isolates, ii) deposited in two or more recognised culture repositories, and ii) validly published in agreement with the International Code of Nomenclature of Prokaryotes (ICNP) is indeed rare, with the List of Prokaryotic Names with Standing in Nomenclature (LPSN) documenting fewer than ten between 2020 and 2025 ^6^. This clearly reflects the ongoing challenges of replicating adequate growth conditions in axenic cultures which was recently referred to as “gold standard” in microbiological sciences ^7^.

The class SHA-98 (placeholder name, phylum *Bacillota*) was first mentioned in Release 202 of the GTDB in April 2021 and contains two orders (UBA4971 and Ch115), three families, six genera, and 19 species in the current release 226. As elaborated with the S*ingleM* software suite implemented in *Sandpiper* ^8^, class *SHA-98* can be found globally and primarily in anaerobic digesters but also in (crude) oil-, hydrocarbon-, and coal associated habitats, wetlands, hot springs, groundwater, manure, and landfills (https://sandpiper.qut.edu.au/taxonomy/c__SHA-98): For instance, UBA6256 sp012516125 showed a relative abundance (RA) of up to 30% in thermophilic anaerobic digestion (AD) reactors supplemented with a volatile fatty acids (VFA) mixture ^9^. UBA4971 sp900019985 thrived with a RA of 9% in mesophilic (41°C) reactors digesting maize silage ^10^. Class *SHA-98* was found at elevated ammonia concentrations caused by anaerobic digestion of protein-rich substrates such as slaughterhouse waste ^11^. Ch115 sp013178415, the only documented species of this genus, was sampled from habitats such as (thermophilic) anaerobic digesters, deep subsurface shale gas wells^12^, and dispersed coal-bearing rocks near an underground coal fire^13^ (https://sandpiper.qut.edu.au/taxonomy/s__Ch115%20sp013178415). The GTDB species representative was released in May 2020 and derived from a 2.8-km deep subsurface aquifer. However, neither this species nor other representatives of class SHA-98 have yet been isolated and thus described as pure cultures.

In this study, we present the isolation, physiological characterisation, and genomic analysis of strain PM69, closely related to MAG *Ch115 sp013178415* (ANI: 98.22), which represents the first isolated member of the class *SHA-98*. Based on its physiological and phenotypic characteristics, we propose the name *Thermoaminiphila catenidiffluenda gen. nov, sp. nov*. and have named the higher taxonomic ranks accordingly. We isolated strain PM69 from a thermophilic batch reactor with phenyl acids as carbon source but with otherwise oligotrophic conditions.

## 2. Methods

### 2.1. Experimental setup and separation of pure cultures

Strain PM69 derived from a thermophilic (55°C), 250 mL semi-batch lab reactor system anaerobically digesting a mixture of phenyl acids (PA) in a long-term experiment. The inoculum of the batch reactor derived from an industrial scale, organic fraction of municipal solid waste (OFMSW) digesting reactor (Roppen, Tyrol) operated at thermophilic temperatures. After an adaption phase with also other substrates, phenylacetic acid (PAA), 3-phenylpropionic acid (PPA), 3-phenylbutyric acid (3-PBA), and 4-phenylbutyric acid (4-PBA) were the sole carbon sources for the microbial community. Feeding intervals, substrate amounts, and liquid exchange rates varied to prevent over-feeding with potentially problematic compounds (PAs) but keep microbial community function alive and active.

After 34 months, we prepared an anaerobic, PA containing medium agar in 120 mL serum flasks sealed with butyl rubber stoppers and aluminium caps. Via syringe and cannula, we added the above-mentioned batch culture to still liquid but already viscous medium, thoroughly mixed and immediately cooled down the medium. The serum flask was incubated at 55°C for 4 months. With a cannula and syringe, we picked one isolated culture and transferred it to a new infusion flask containing Starkey’s medium under anaerobic conditions. The culture was then incubated at 55°C for additional 2 months. Following multiple generations and preliminary metabolic reconstructions, we switched to a medium with a more defined formulation comprising D-glucose as main carbon source and casamino acids (*Thermoaminiphila* medium). For the extended methodology, refer to Supplementary Methods (Section MM1).

### 2.2. Growth characteristics

Unless stated otherwise, after medium optimisation, PM69 was cultured in *Thermoaminiphilia* medium with D-glucose as main carbohydrate source at 55°C under anaerobic conditions. Growth was determined as optical density at 600 nm on a LAMBDA^®^ 365+ double-beam UV/Vis spectrometer (PerkinElmer, USA) with the respective *Thermoaminiphila* medium as reference after 0, 1, 2, 3, and 7 days (8% inoculum, 3 replicates). We used *Thermoaminiphila* medium at different temperatures (10°C, 20°C, 30°C, 37°C, 45°C, 50°C, 55°C, 60°C, or 65°C). By replacing D-glucose in the medium, we also tested galactose, maltose, arabinose, saccharose, starch, fructose, xylose, and lactose as carbon sources (5 g L^-1^). We incubated PM69 at an initial pH of 5.0, 6.0, 7.0, 7.5, 8.0, and 8.5 (adjustment with 1 M HCl or 1 M NaOH) and determined growth at 0, 0.3, 0.10, 0.50, 1.00, 2.00, 5.00, and 10.00% NaCl. By checking turbidity visually for 2 weeks, the growth in absence of yeast extract or casamino acids, yeast extract and casamino acids, and of D-glucose was also tested. To check susceptibility of PM69 to oxygen, we additionally incubated the strain at reduced aerophilic (16% oxygen) and aerophilic conditions.

### 2.3. Biochemical analyses

Gas production (gas pressure) and composition (H_2_, O_2_, CH_4_, CO_2_) (GC-TCD) were determined as described previously^14^. VFA (formate, acetate, propionate, i-butyrate, butyrate) were quantified as outlined earlier via high-performance liquid chromatography ^15^ with the modification that a Rezex ROA-Organic Acid H+ (8%, 300 x 7.8 mm) column was used at a flow rate of 0.6 mL/min. For analyses of cellular fatty acids and an extended screening of metabolites of cultures at two different growth stages via GC and HPLC, refer to the Supplementary Methods (Section MM2 and MM3, respectively).

### 2.4. Cell Morphology

Cells were visualised by phase-contrast and interference microscopy (Nikon Eclipse Ni with a Nikon DS-Ri2 camera). NucBlue™ Live ReadyProbes™ Reagent (Hoechst 33342, Thermo Fisher Scientific, USA) was used for cell staining as it is excited at 360 nm when bound to DNA (emission maximum at 430 nm). Gram staining (COLOR Gram 2 Kit, Biomérieux, France) was done according to the manufacturer’s protocol. Next, we used 3% (w/v) KOH for the assessment of further gram characteristics ^16^. For scanning electron microscopy (SEM), microbial cells were separated from the medium with a 0.8 µm nylon membrane filter through vacuum suction and fixated in a solution of 2.5% glutaraldehyde (25% solution, Sigma-Aldrich) and 0.1 M phosphate-buffered saline (PBS, Roche, Sigma-Aldrich, pH 7.3) for four hours ^17^. Thereafter, cells were washed with PBS (0.01 M, VWR) thrice for 10 minutes and dried via the ethanol gradient drying method as described previously ^18^. The filter was submerged in hexamethyldisilane (≥99 % HMDS, Sigma-Aldrich) for 30 seconds. The solution was removed with a pipette followed by drying overnight. A small part of the filter was dissected, mounted and coated with gold-palladium for 180 seconds for the field emission SEM analysis (Zeiss Supra 35 VP, Zeiss, Germany). A detailed description can be found in file Supplementary Methods (Section MM4).

### 2.5. Sanger sequencing of the 16S rRNA gene

Detailed protocols for DNA extraction, PCR amplification, quality checks, and constructing the 16S rRNA gene phylogenetic tree are provided in the Supplementary Method file (Section MM5). To assess the culture purity and evaluate the quality of the received 27-f/1492-r Sanger sequences, trace files were inspected for heterozygous positions using the CLC Main Workbench 23.0.2 (QIAGEN, Denmark) throughout the study (n=16). Sequences were classified with the SILVA Alignment, Classification and Tree Service (SILVA ACT: SINA Aligner v1.2.12, SINA Search and Classify, LCA method, https://www.arb-silva.de/aligner/) using default parameters and all available taxonomies ^*19*^. Additionally, consensus sequences ^20^ were queried individually against the NCBI nucleotide database using BLASTn; closely related reference sequences (> 95% coverage) were retrieved and dedublicated^21^. Maximum-likelihood phylogenetic inference was conducted using IQ-TREE (v2.0.7) ^22^ using the best-fitting nucleotide substitution model (TIM3+F+I+G4) for tree inference. The final 16S rRNA tree was visualized and annotated using package *ggtree* (v4.0.4) ^23^.

### 2.6. Whole genome sequencing and genome assembly

Genomic DNA was extracted using the Quick-DNA HMW MagBead Kit (ZYMO RESEARCH, USA) according to the manufacturer’s protocol. After quality checks and quantification, we sent an aliquot to Microsynth AG (Switzerland) for whole genome sequencing on a NovaSeq device (2×150bp, RRID:SCR_024569) according to internal protocols with demultiplexing and trimming of Illumina adapter residuals post sequencing. We did long-read sequencing in-house with the Native Barcoding Kit (SQK-NBD114.24, V14 chemistry, Oxford Nanopore, UK), companion products (NEB, USA), and the Oxford Nanopore MinION device (RRID:SCR_017985) using a FLO-MIN114 flow cell and the MinKnow software v24.06.5. After long-read sequencing, we base-called pod5 files, demultiplexed, trimmed, and corrected fastq files with *dorado* v0.7.3 according to Oxford Nanopore protocols.

*Illumina shot-gun* sequences were filtered with *Trimmomatic* PE v0.39 ^24^ (-phred64, ILLUMINACLIP:TruSeq3-PE.fa:2:30:10:2:True, LEADING:3, TRAILING:3 MINLEN:36) and *Nanopore long-read* sequences with *Filtlong* ^25^ (removed reads with a length below 1kb). We visualised *Illumina* sequences with *fastqc* v0.12.1 (https://www.bioinformatics.babraham.ac.uk/projects/fastqc) and *Nanopore* sequences with *nanoQC* v0.10.0 ^26^ prior and after filtering.

Hybrid assembly (short-read-first) was done with *Unicycler* v0.5.1 ^27^. *Via seqkit* ^*28*^, all contigs with a length below 1000 bases were excluded from further analyses. Quality assessment of the assemblies was done with QUAST 5.2.0 ^29^, *Bandage v0*.*8*.*1* ^30^, and *CheckM2* ^31^. Additionally, we did a chimera and quality check with *GUNC* 1.0.6 ^32^ using both, the *proGenomes* 2.1 ^33^ and *GTDB* databases with the optional flags *--sensitive* and –*contig_taxonomy_output*. Next, we searched for rRNA and tRNA sequences with *barrnap* v.0.9 ^34^ and *tRNAscan-SE* v2.0.12 ^35^, respectively, using default settings, and used ABRicate (https://github.com/tseemann/abricate) to screen for antimicrobial resistance and virulence genes with the databases NCBI Parminderpal’s ^36^, CARD ^37^, Resfinder ^38^, ARG-ANNOT ^39^, VFDB ^40^, ecoli_vf (https://github.com/phac-nml/ecoli_vf), PlasmidFinder ^41^, EcOH ^42^, and MEGARes 2.0 ^43^.

### 2.7. Phylogeny and Biogeography

Besides the recovered genome of PM69, we also included all binned GTDB species representatives of class SHA-98 obtained from NCBI RefSeq as well as MAGs deriving from the same or analogue batch reactors (chapter 2.1, Supplementary Methods, Table MM1). Phylogenomic inferences were based on the bacterial GTDB alignment (R10-RS226) comprising 120 marker proteins, whereby the alignment was created with GTDB-Tk (v2.3.2; GTDB-Tk reference data version r226) ^44^. To account for differences in substitution patterns between lineages, i.e. branch-wise compositional heterogeneity, we removed the top 30% most compositionally biased sites from the alignment, ranked according to chi-square scores by the alignment_pruner.pl script ^45^. Next, we inferred two phylogenetic trees 1) with 772 taxa and 2) with 1412 taxa (GTDB RS10-226 order representative). Initial trees were calculated with FastTree2 ^46^ and the output was used as a starting tree for IQ-TREE (v2.3.6) ^22,47^ to infer the phylogeny with the command “iqtree -s <input alignment> -nt 12 -m LG+C20+F+G -ft <starting tree> -pre <prefix of output files> --bb 1000”, with 1000 bootstrap support values. Resulting trees were decorated with PhyloRank (v0.1.12; https://github.com/donovan-h-parks/PhyloRank) and visualized with the software package ARB ^48^ and online display tool iTOL ^49^. For biogeographic information, we applied the sandpiper software suite ^8^ for class SHA-98 and genome Ch115 sp013178415.

### 2.8. Core metabolism of T. catenidiffluenda sp. nov

Protein prediction was done with *Prodigal* v2.6.3 ^50^ for strain PM69. Resulting protein FASTA files were used for KO annotation and *KEGG* mapping via *BlastKEGG Orthology Ank Links 164 Annotation* (*BlastKOALA*). Next, we used the *NCBI-datasets-cli* to download the GTDB representative genome of Ch115 sp013178415 (Accession GCA_013178415.1) and other representative sequences of class SHA-98 which were co-analysed with the assembly of strain PM69. For this purpose, we used *gapseq* v1.4.0 ^51^ (*Bacteria* database: 2024-11-06) which allowed pathway and transporter predictions. For further metabolic reconstructions for strain PM69, we reconstructed a draft metabolic network, performed growth medium predictions and gap filling. For the latter two steps, anaerobic conditions (cpd00007:0) were defined. After adapting the phenyotype medium considering first predictions, the gapseq medium was modified in a second run by removing lactose (cpd00208:0), xylose (cpd00154:0) and L-lactate (cpd00159:0), and by adding casamino acids (cpd30698:5). This allowed us to compare growth rates, predict metabolic products, and to perform gap filling for a second genome-scale metabolic model (phenotype model). Subsequently, we conducted parsimonious flux-balance analyses (PFBA) with minimization of total flux (MTF) using R version 4.4.3 ^52^ and the packages *cobrar* ^53^ and *data*.*table* ^54^. The *Additional_constraints*.*Rm* script (https://github.com/Waschina/cobrar/tree/main/vignettes) was applied to assess growth on specific carbon sources (Section MM6, Supplementary Methods). When growth was incorrectly predicted for carbon sources on which the phenotype could not grow, we consulted the reaction tracing file (https://gapseq.readthedocs.io/en/latest/tutorials/traceability.html).

Reactions identified as erroneously added during gapfilling - or missing from KEGG predictions - were subsequently removed.

The symbiont classifier *symclatron* v0.6.4 was also applied to predict the life style (free-living or symbiont) ^55^ for genome *Ch115 sp013178415*.

## 3. Results

### 3.1. Biochemical analyses

#### 3.1.1. Gas and volatile fatty acids production, and metabolite screening

Headspace gas of a two-, three-, and five-day old culture in *Thermoaminiphilia* medium comprised 11.1%, 14.6% and 18.9% H_2_, as well as 27.3%, 28.8%, and 29.6% CO_2_, and produced 0.75, 1.24, and

2.01 NmL H_2_, as well as 1.36, 2.19, and 3.09 NmL CO_2_, respectively. Long-term VFA production was assessed across multiple generations (20^th^ – 63^rd^) and ages of the cultures (3 – 66 days old, Supplementary Results, Table S1.): Maximum acetate concentration was 15.2 mM in a 7-days old culture (59^th^ generation). Average acetate concentration was 9.94 ± 1.91 mM with no significant differences among ages of the respective cultures. Further metabolite analyses showed that PM69 does not produce short-chain (C_3_ – C_7_) or medium-chain (C_8_ – C_12_) carboxylic acids other than acetate, regardless of the age of the culture. Additionally, ethanol accumulation reached up to 0.5 g L^-1^ (Supplementary Results 1, Table S2).

#### 3.1.2. Cellular fatty acids

Major, saturated fatty acid methyl esters of PM69 were iso-15:0 (29.1%), iso-17:0 (19.1%), anteiso-15:0 (17.2%), anteiso-17:0 (8.7%), iso-13:0 3-OH (7.7%), and 16:0 (6.0%). Unsaturated FAME were 18:1 w9c (1.2%), 16:1 (summed feature, 0.20%), and 18:1 (summed feature, 0.40%). For a full list of FAME, refer to the Supplementary Results 1 file (Table S3).

### 3.2. Cell morphology

PM69 was rod shaped with a usual length between 1.5 and 3 µm and width between 0.4 and 0.6 µm. Most cells sticked together and formed long chains with up to over 100 µm in length (Fig. 1). PM69 also appeared in short chains of at least two cells or as independent rods. SEM pictures indicated 2-3 flagella on one end of the rods. Cells were gram variable, whereby the KOH test showed gram positive characteristics. Spore formation was not observed, regardless of the age of the culture.

**Figure 1.**
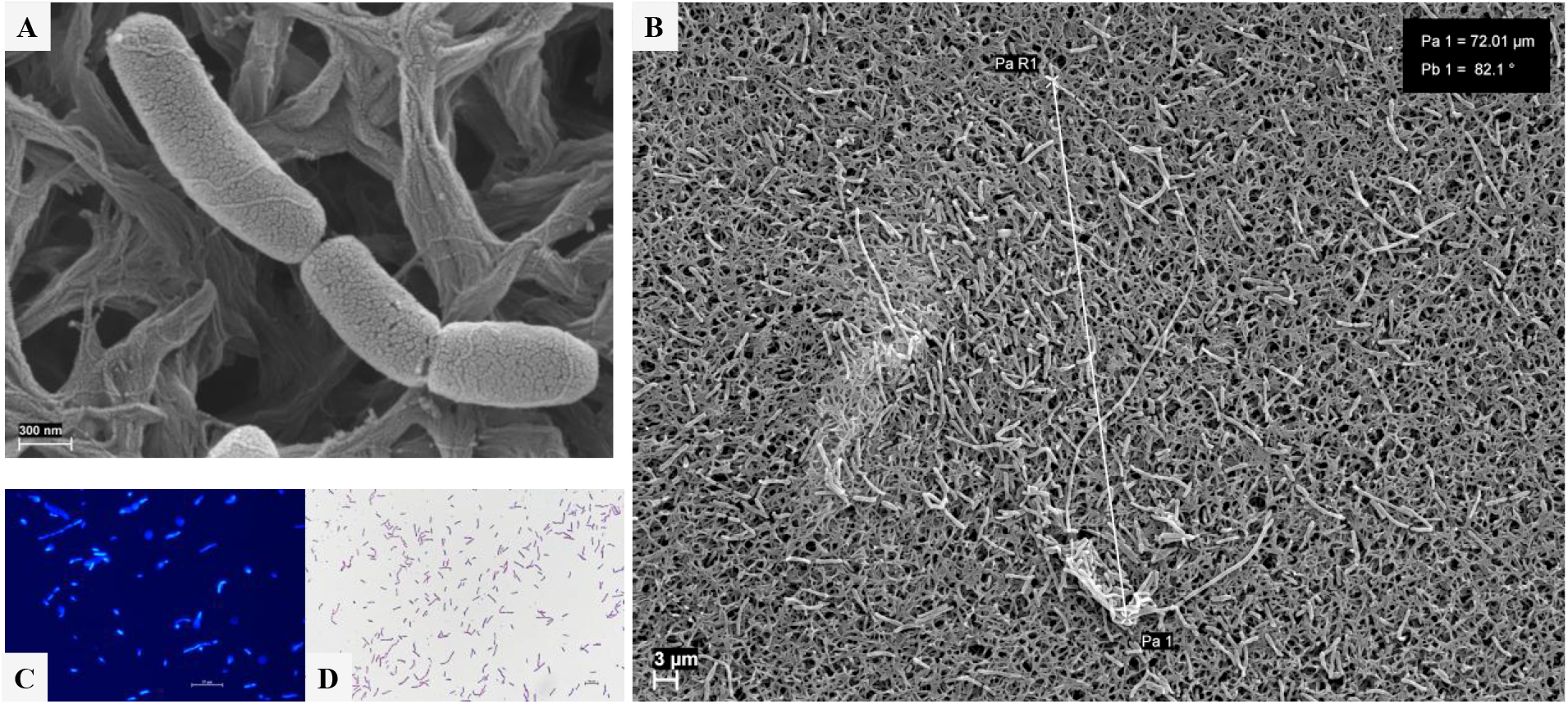
Visualisation of strain PM69 via SEM showing cells with 2-3 flagella at each end of the short chain (A) as well as cell-chains longer than 100 µm (B). Single rods and cell-chains were also visualized using a light microscope and Hoechst 33342 live-cell (C) and gram staining (D).

### 3.3. Growth characteristics

When inoculated in exponential phase into *Thermoaminiphila* medium, visible turbidity was observed after one day of incubation, high turbidity after two days. During the cultivation procedure, restricted or no growth was experienced when the inoculum was older than 4 days. PM69 demonstrated growth across a temperature range from 45°C to 60°C with optimal growth occurring at 55°C (Fig. 2). It exhibited reduced proliferation at pH 5.0 – 6.0 but thrived at pH 7.0 - 8.5 with a maximum at pH 8.5. Within seven days, PM69 could utilise D-glucose, fructose, and xylose. Moderate growth was observed with galactose as carbon source. The strain could not grow on L-lactate, lactose, sucrose, maltose, arabinose, and starch. The strain tolerated NaCl concentrations up to 1%, (w/v) with an optimal concentration at 0.1 – 0.5% (w/v). Without yeast extract addition, after three slowly growing generations (10% (v/v) inoculum), PM69 could not be recultivated. No growth was observed with casamino acids as sole carbon source or when both, yeast extract and casamino acids were absent from the medium (with glucose as main carbon source). PM69 was able to grow without casamino acids for 16 generations albeit at a slightly reduced rate. However, casamino acid addition was essential during the process of switching from a complex (Starkey’s medium) to a more defined medium (*Thermoaminiphila* medium) as PM69 exhibited restricted growth during this process and recovered after the casamino acid amendment. PM69 did not grow under reduced aerophilic or aerobic conditions and did not show genes for antimicrobial resistance, virulence and stress response according to genomic analyses.

**Figure 2.**
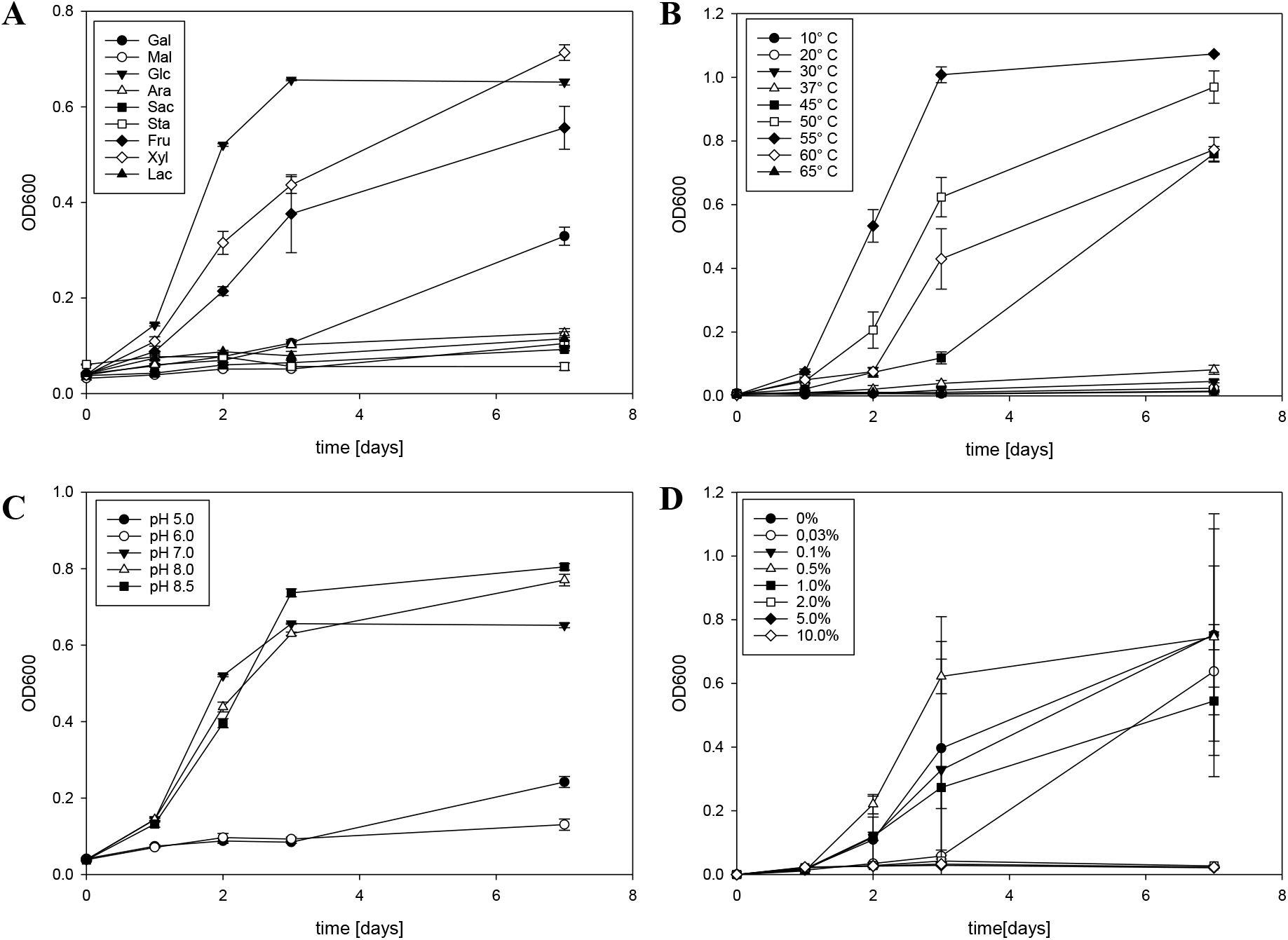
Depiction of growth characteristics of strain PM69 determined as optical density at 600 nm on various substrates (A), at different temperature (B), across a range of pH levels (C), and at different NaCl concentrations (D) over a 7-day incubation period.

### 3.4. Sanger Sequencing, and Whole Genome Sequencing and Genome Assembly

Sanger sequencing reads were trimmed using a strict quality threshold (Phred-Score ≥30, corresponding to an error probability of 0.001) resulting in a final sequence length of 1017 ± 25 base pairs. Electropherograms of cultures grown in Starkey’s and *Thermoaminiphila* medium showed single, clean sequence trace without overlapping or ambiguous peaks. SILVA ACT classified sequences as “unclassified” – regardless of the database.

For all results regarding quality control of whole genome sequencing and genome assembly, refer to Supplementary Results 1. After filtering, about 96% of *Illumina* reads (4 663 453 sequences) remained for R1 and R2 with a sequence length ranging from 36 – 125 bp (Table S4). *Checkm2* predicted a completeness of 100% for the assembly. After removal of all contigs <1kb, contamination decreased from 1.39% to 1.32%. The genome showed a total length of 3,303,196 bp (19 contigs, average length: 173,852.4 bp, maximum length: 791,104 bp, N50: 381,840 bp, N90: 183,514 bp, Table S5 and S6). The re-alignment process mapped 98.4% of sequences to the assembly, with 97.7% properly paired. GC content was 51.3%. The assembly with and without contigs <1kb contained 49 and 47 tRNA coding sequences, respectively (Table S5). Predictions were made for the 5S, 16S, and 23S rRNA regions (Table S7), and the 16S rRNA gene sequences, elaborated via PCR, are provided in Table S8. Both assemblies passed GUNC analysis, confirming high quality and the absence of chimeric sequence contamination (Table S9).

### 3.5. Phylogeny and Biogeography

For class SHA-98, 16 of 19 binned GTDB representative species met the MIMAG criteria for high-quality genomes with a completeness of >90% and contamination of <5% and were included for taxonomic assignment and phylogenomic interference. Additionally, we used 13 MAGs recovered from similar or analogue batch reactors (Supplementary Methods, Table MM1) and stated their quality in Fig. 3A. Taxonomic assignment confirmed their affiliation to one of two orders (Ch115 or UBA4971) within the class SHA-98. Within the order Ch115, strain PM69 belongs to the family and genus Ch115 and shows high similarity with genome s__Ch115 sp013178415 (ANI: 98.22, AF Query: 947, AF Reference:1094). The matrix of queries, references, and corresponding ANI and AF values used for the phylogenetic tree (Fig. 3A) is available in Supplementary Results 2.

**Figure 3.**
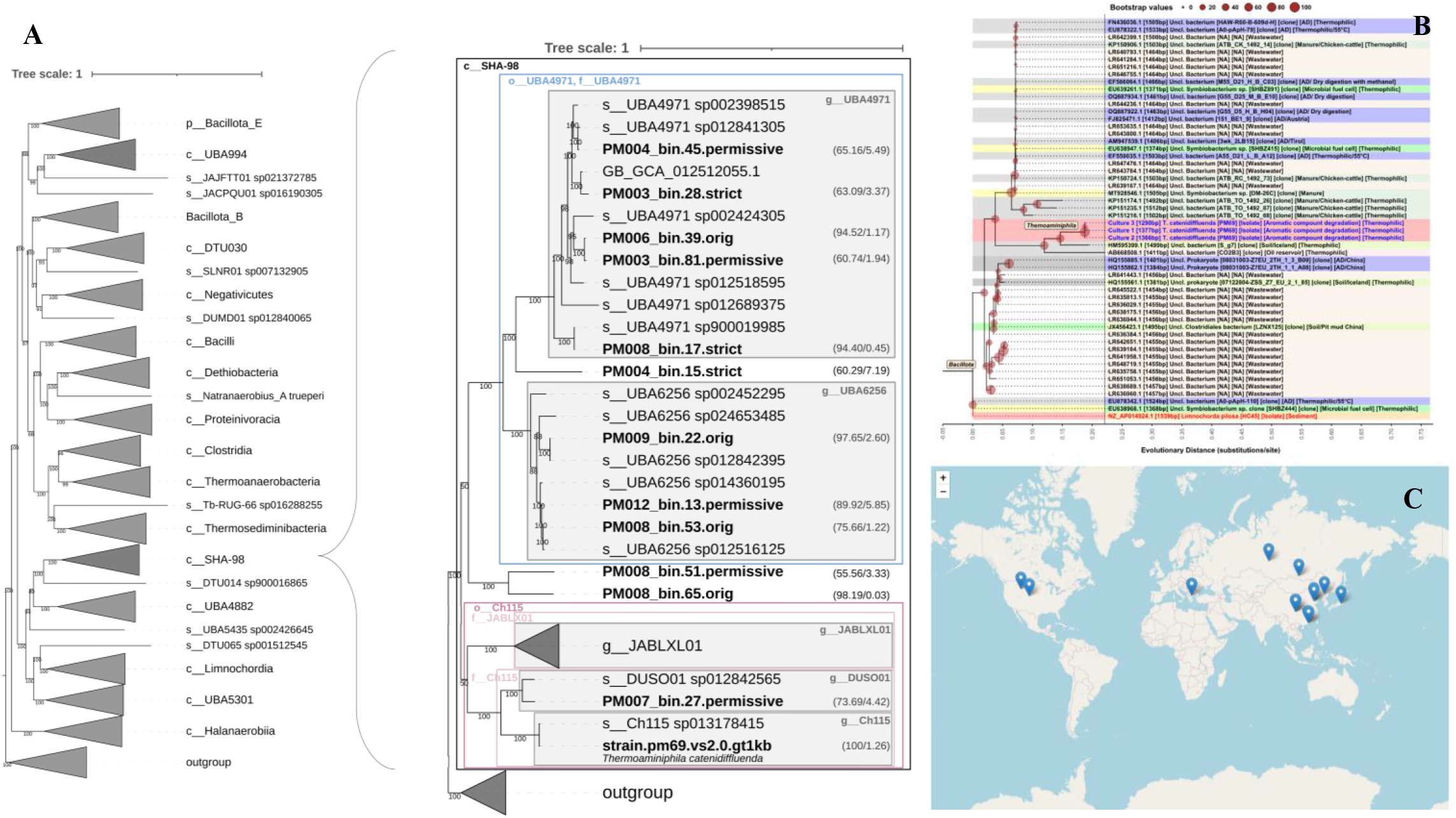
**A**: Phylogenetic tree based on the whole genome of strain PM69, GTDB representative MAGs of class SHA-98 (beginning with “s__“or “g__“) as well as MAGs of class SHA-98 found in the same or similar batch reactors (bold, beginning with “PM0”); the quality of those MAGs is provided in brackets. Detailed information on additional MAGs can be found in Supplementary Methods, Table MM1. **B**: Phylogenetic tree based on 16S rRNA gene sequences. The colours on the leaves represent the isolation source/environment of the sequence. Sequences of strain PM69 are highlighted in red. **C**: MAGs of *T. catenidiffluenda* sp. nov. (designated as Ch115 sp013178415) identified globally using sandpiper (April 2026).

Using sandpiper (Fig. 3C), we identified 33 matching samples for Ch115 sp013178415 which was predominantly detected in (thermophilic) biogas fermenters with RAs up to 1.36% ^8^. Some substrates for these reactors were i.e., acetate, CO_2_ and H_2_ ^56^, or activated sludge ^57^. Notably, some of these reactors were exposed to microplastics and antibiotics (https://sandpiper.qut.edu.au/run/SRR31359994). The species is also associated with terrestrial and subsurface environments rich in coal, gas, and oil, as well as paddy soils and hot springs. In these environments, its RA was consistently below 0.5% (https://sandpiper.qut.edu.au/taxonomy/s__Ch115%20sp013178415).

### 3.6. Core metabolism and cellular processes of T. catenidiffluenda sp. nov

For a complete list of pathways and complete KEGG modules as well as *gapseq* results, refer to Supplementary Results 1, Fig. S1 – S4 as well as file Supplementary Results 3, respectively. The phenotype of PM69 could grow with D-glucose, xylose, D-fructose and – to a lesser extent - also on D-galactose but not on arabinose as well as di- and polysaccharides (Fig. 2). Indeed, PM69 contains the complete glycolysis and pyruvate oxidation pathway (glucose to acetate) as well a complete module for galactose degradation (Leloir pathway). *Gapseq* predicted the glycolysis VI pathway from fructose as true with 81% completeness; however, this pathway has only been observed in mammals so far. By gap filling the draft model with the predicted medium, a maximal growth rate of 1.11 h^-1^ (gram-positive biomass) was predicted with D-glucose, xylose, lactose, L-serine, L-tyrosine, and L-lactate as substrates. Major products of this model were H_2_, CO_2_, acetate, and H^+^ with 58.1, 32.2, 27.5, and 19.6 mmol g^-1^ DW h^-1^, respectively. Entner-Doudoroff pathways as well as the Wood-Ljungdahl pathway were predicted as false. Flavin-based electron bifurcation (FBEB) including a lactate dehydrogenase (electron bifurcating LDH/Etf complex) was predicted as true (100%). With the phenotype medium, *gapseq fill* predicted a final growth rate of 0.30 h^-1^with D-glucose and L-Serine as substrates (maximum uptake rate: 5 and 0.1 mmol g^-1^ DW h^-1^, respectively). Major products of the phenotype model were H_2_ (12.5), H^+^ (7.65), CO_2_ (7.11), and acetate (5.78 mmol g^-1^ DW h^-1^). By providing only one specific carbon source to the GEM, we could confirm considerable growth on D-fructose (maximum growth rate 0.30 h^-1^), galactose (0.24 h^-1^), and xylose (0.20 h^-1^) and confirmed very slow growth on L-lactate (0.06 h^-1^) and arabinose (0.002 h^-1^). According to the FBA and MTF analyses, PM69 could also grow on lactose (0.57 h^-1^), maltose (0.62 h^-1^), and sucrose (0.62 h^-1^). Those reactions (EX_cpd00208_e0, EX_cpd00179_e0, and EX_cpd00076_e0) were added during gap filling (step7: exchange reactions); consequently, we removed those again from the phenotype model. PM69 further encodes the enzymatic repertoire (100% completeness) for degradation of L-alanine, L-aspartate, L-glutamate, L-cysteine, L-arginine, L-alanine, L-phenylalanine, L-threonine, L-glutamate, L-serine, and L-threonine. PM69 is potentially capable of sulfur respiration via the sulfur reduction III pathway. Pentose-phosphate pathways are incompletely encoded: A possible way to form 5-phosphoribosyl 1-pyrophosphate (PRPP) is via D-ribulose-5P. Besides, the sedoheptulose bisphosphate bypass was predicted as true (100% completeness). PM69 possesses complete modules for both, F-type ATPase (*Bacteria*) and V/A-type ATPase (*Bacteria, Archaea*) and harbours >30 genes for quorum sensing and 33 genes for flagella assembly whereby genes for H- and T-rings, needed in gram-negative bacteria, are missing. Notably, strain PM69 is the only GTDB representative to be potentially capable of cyanide degradation (P401-PWY) within the class SHA-98 (Fig. 4).

**Figure 4.**
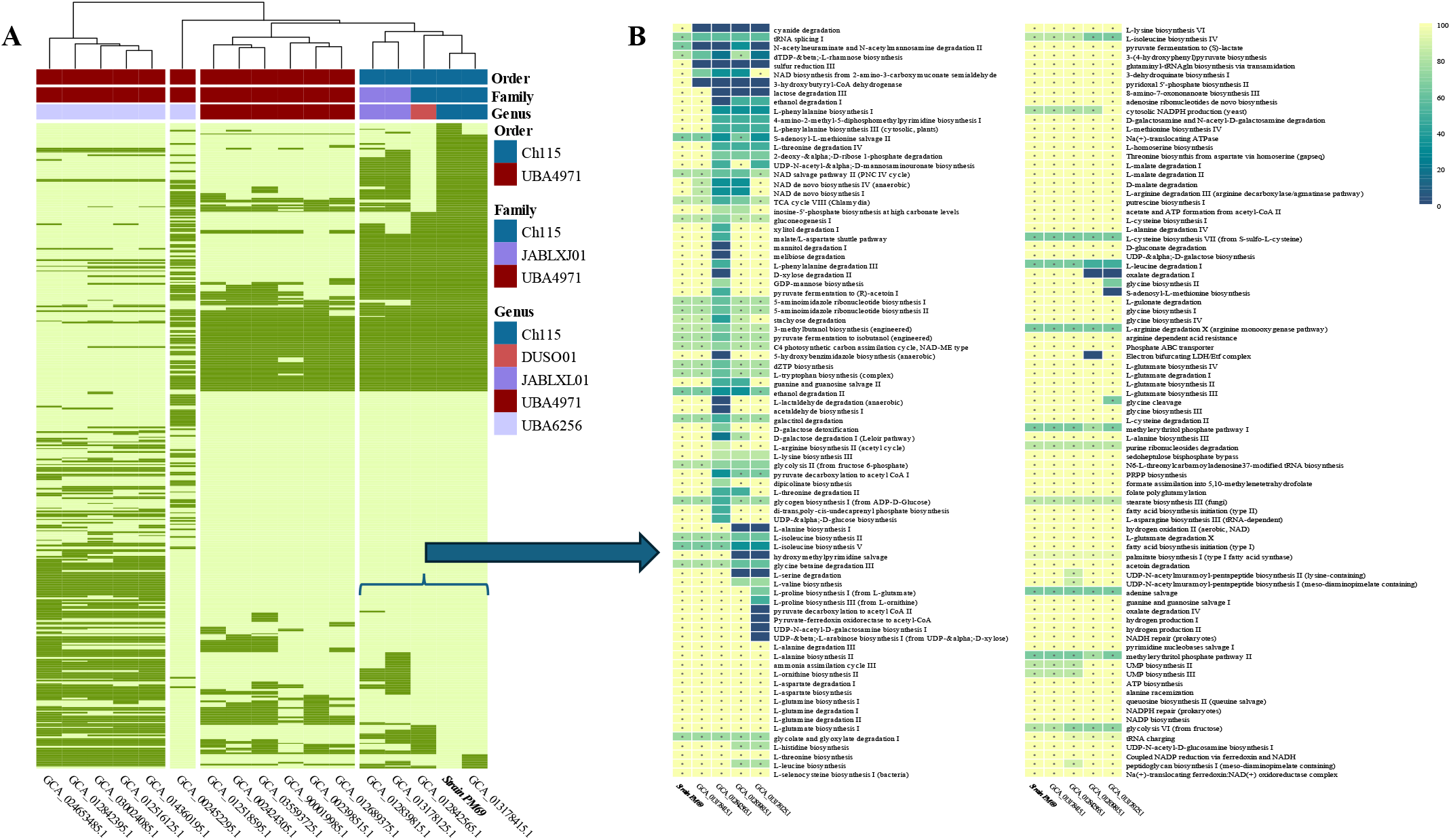
**A:** Overview of predicted metabolic profiles of GTDB representative MAGs of class SHA-98 in comparison to strain PM69: Dark green and light green indicate pathways predicted to be true and false, respectively. The rightmost accession number, GCA_013178415.1, corresponds to CH115 sp013178415 which is - alongside strain PM69 – designated as *T. catenidiffluenda* gen. nov., sp. nov. **B:** A detailed comparison of metabolic pathways predicted to be present (asterisks) in strain PM69 relative to other members of order CH115: Pathway completeness is indicated by colour, ranging from 100% (yellow) to 0% (blue).

*Symclatron* predicted a free -living lifestyle for *Ch115 sp*. genomes (>98% completeness, confidence >0.98). For comparison, strain PM69 – its autonomous growth already proven in lab experiments – showed 100% completeness (confidence > 0.99).

## 4. Discussion

*T. catenidiffluenda* gen. nov., sp. nov. is a monosaccharide-fermenting, strictly anaerobic, and thermophilic bacterial species: Strain PM69 utilises five- and six-carbon monosaccharides as carbon sources but does not metabolise disaccharides or sugar polymers. Genomic analysis further supported this observation, as pathways for sucrose degradation and extracellular cellulose and starch hydrolysis were predicted to be absent. Within the cascade of polymer degradation to biogas, *T. catenidiffluenda* sp. nov. is most likely involved in the acidogenesis stage with hydrogen, carbon dioxide, and acetate as major metabolic end products during anaerobic oxidation. These intermediates are critical for downstream processes during AD. Notably, while strain PM69 cannot grow on L-lactate as the sole carbon source, *in silico* metabolic modelling (via gapseq) predicted an increased growth rate in its presence, suggesting a role for lactate in energy conservation. Lactate plays a dual role in the metabolism of strictly anaerobic microorganisms. It can serve as an electron donor for NAD^+^ regeneration, a process often coupled with the simultaneous oxidation of reduced ferredoxin via electron confurcation ^58,59^.

Indeed, strain PM69 encodes multiple genes for L- and D-lactate dehydrogenases. Notably, L-lactate concentrations were observed to increase slightly in the growth medium (Supplementary Results 1, Table S2), suggesting that electron bifurcation to lactate—rather than confurcation—may prevail in strain PM69 growing with *Thermoaminiphila* medium. PM69 also exhibits other features typical of obligate anaerobes, including the absence of the oxidative branch of the pentose phosphate pathway and the Entner–Doudoroff pathway ^60,61^. Instead, it exclusively employs glycolysis (Embden-Meyerhof-Parnas pathway) for glucose degradation, and it most likely uses the sedoheptulose bisphosphate bypass for five-ring sugar metabolism and anabolic processes (Fig. 5).

**Figure 5.**
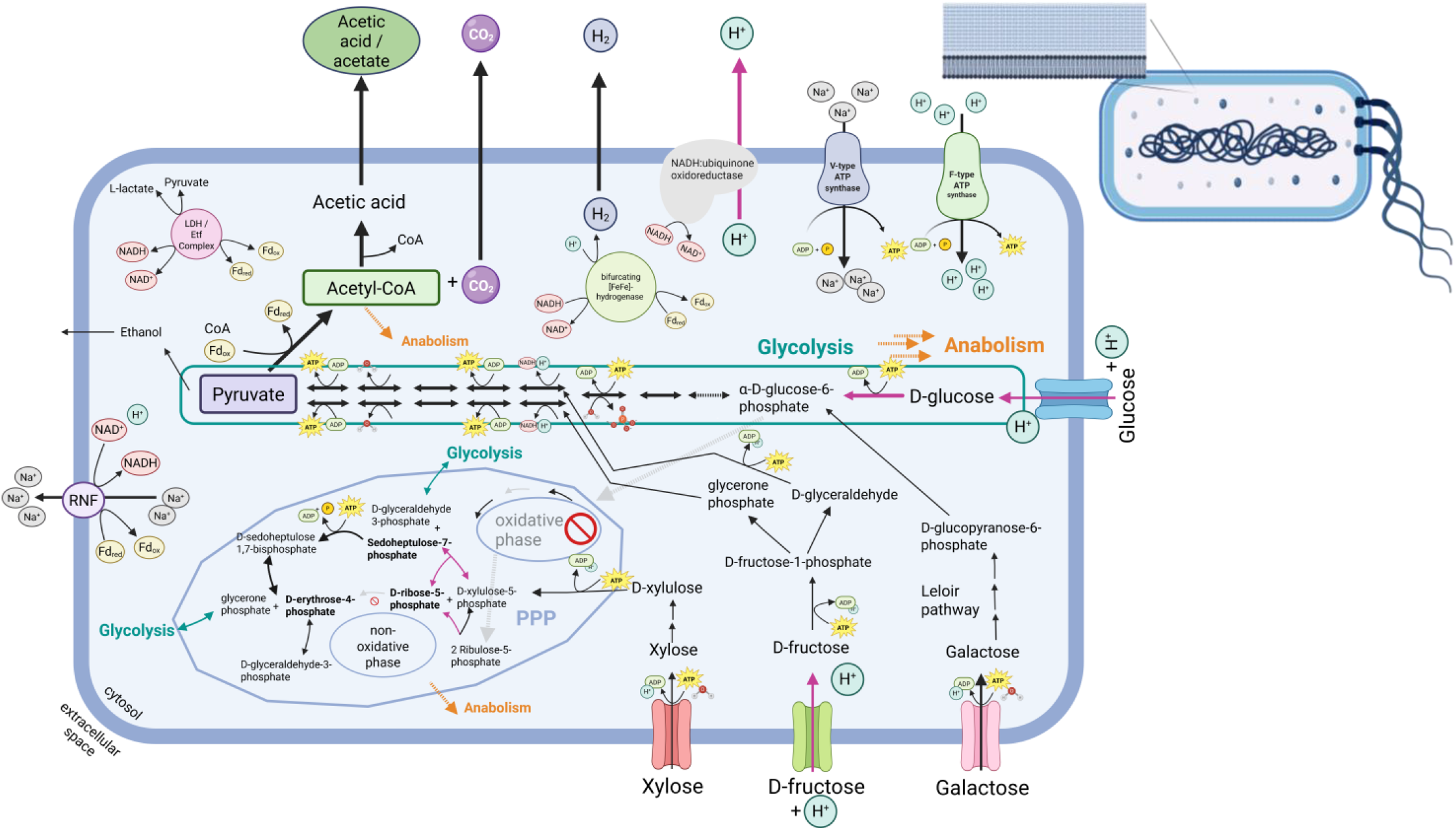
Cell architecture and proposed core metabolism of *T. catenidiffluenda* sp. nov. strain PM69 with acetic acid, CO_2_ and H_2_ as major products during monosaccharide fermentation. Arrows in purple indicate reactions inferred through metabolic reconstruction using *gapseq*. Strain PM69 encodes both, V-type as well as F-type ATPases and suggests potential for electron bi- and confurcation and utilisation of pyruvate / lactate to maintain redox balance. SEM pictures confirmed the presence of 1-3 flagella which is consistent with genomic predictions for flagellar and peptidoglycan biosynthesis. RNF: Rhodobacter nitrogen fixation, LDH: Lactate dehydrogenase, Etf: Electron transfer flavoprotein.

Strain PM69 exhibited rapid growth kinetics during monosaccharide fermentation and high biomass yields under lab conditions thus has an efficient glycolytic pathway. It remains noteworthy that this species but also other representatives of class *Thermoaminiphilia* are repeatedly associated with hydrocarbons (oil, gas, coal), aromatic compounds (e.g., phenyl acids, Table MM1), and oligo- or autotrophic habitats. Strain PM69 was isolated from a thermophilic, phenyl acid degrading batch reactor where its MAG exhibited a RA ranging from 0.87 to 2.66% (these results constitute a sub-analysis of an ongoing project, the full results of which are currently in preparation for publication). Metabolic reconstruction convincingly demonstrated that T. *catenidiffluenda* cannot cleave benzene rings which is a distinct and highly energy demanding process under anaerobic conditions. Nevertheless, the presence of a diverse set of genes associated with electron bifurcating LDH/Etf enzyme complex, cyanide detoxification (Fig. 4B), biofilm formation (e.g., complete modules of polyamine and glycan metabolism), quorum sensing, and flagellar assembly suggests strong capacity of environmental adaption in or avoidance of harsh and fluctuating habitats and colonising distinct ecological niches within the acetate producing stages of anaerobic digestion. The capability of *T. catenidiffluenda* to form long chains also underscores the possible relevance of biofilm formation ^62^, as well as cooperative nutrient uptake and stress response ^63^ for this bacterium in unfavourable environments. Previous attempts to isolate microorganisms of interest for biofilm formation have only been successful in rare instances ^64^. From both a technological and ecological perspective, the isolation of strain PM69 – the first isolated representative of class *Thermoaminiphilia* - may therefore be of relevance for researchers and justifying further investigations.

## 5. Description of cultivated strain PM69 in accordance with ICNP

### 5.1. Thermoaminiphila gen. nov

The description of genus *Thermoaminiphila* (ther.mo.a.mi’ni.phi.la Gr. adj. *thermos* warm; N.L. *amino* amino; Gr. masc. adj. *philos* loving; N.L. fem. adj. *Thermoaminiphila* heat-and-amino-acids -loving) is based on a single cultivated species, which is a thermophilic, monosaccharide-fermenting bacterium requiring obligate anaerobic conditions. Type species of this genus is *T. catenidiffluenda gen. nov*., *sp. nov*. strain PM69, obtained from a thermophilic, phenyl acid degrading batch reactor in Austria.

### 5.2. Thermoaminiphila catenidiffluenda sp. nov

*T. catenidiffluenda* (ca.te.ni.dif.flu’en.da M.L. fem. *catena* chain; M.L. fem. adj. *diffluenda* (gerundive of M.L. v. *diffluere*) to be disintegrated) is represented by the strain PM69 isolated from a thermophilic, phenyl acid degrading batch culture inoculated with digestate from an organic fraction of municipal solid waste digesting, thermophilic biogas reactor (Roppen, Austria). It is rod-shaped, gram variable, and non-spore-forming. PM69 appears as long (> 100 µm) but also short chains and rods. The strain requires anaerobic conditions and a temperature range of 45°C – 60°C with optimal growth at 55°C. It prefers moderately alkaline conditions, with maximum growth at pH 8.5. GC content is 51.3 mol%. Major fatty acid methyl esters are iso-15:0 (29.1%), iso-17:0 (19.1%), and anteiso-15:0 (17.2%). PM69 primarily produces hydrogen, carbon dioxide, and acetate during degradation of D-glucose and casamino acids. It can also grow with xylose and galactose as carbon sources. Genotypically, it can utilise lactate / pyruvate for redox balance. Type strain PM69 can be obtained from the German Collection of Microorganisms and Cell Cultures GmbH (DSM 121036) and Japan Collection of Microorganisms (JCM 39671).

### 5.3. Thermoaminiphilaceae fam. nov

*Thermoaminiphilaceae* (ther.mo.a.mi.ni.phi.la’ce.ae fem. n. *Thermoaminiphila*, type genus; *-ceae* ending to denote family) is based on one cultivated species (*T. catenidiffluenda gen. nov*., *sp. nov*. strain PM69). The description is the same as for type genus *Thermoaminiphila* gen. nov.

### 5.4. Thermoaminiphilales ord. nov

The description of *Thermoaminiphilales* (ther.mo.a.mi.ni.phi’la.les fem. n. *Thermoaminiphila*, type genus; *-ales* ending to denote order) is based on type genus *Thermoaminiphila* gen. nov.

### 5.5. Thermoaminiphilia class nov

The description of Thermoaminiphilia (ther.mo.a.mi.ni’phi.li.a fem. n. Thermoaminiphila, type species; *-ia* ending to denote class) is based on type genus *Thermoaminiphila* gen. nov.

## 6. Conclusions

We could successfully isolate strain PM69 from an oligotrophic, methanogenic consortium reliant on phenyl acids as sole carbon sources. Due to morphological and growth characteristics, we propose the name *Thermoaminiphila catenidiffluenda* gen. nov. sp. nov. of the yet undescribed, monophyletic class *Thermoaminiphilia* class nov. of phylum *Bacillota*. The interplay of broad spectrum of techniques to assess phenotypic and genomic characteristics led to the picture that *T. catenidiffluenda* is a thermophilic, monosaccharide-fermenting bacterium integrated in the acidogenesis stage of biogas production. Despite its relatively fast growth under lab conditions, its biogeography points to a niche existence in diverse environments under moderate conditions as well as in presence of elevated concentrations of hydrocarbons and aromatic compounds. Further investigation into its resistance to and/or behavioural avoidance of harsh or extreme habitats are of considerable scientific interest.

## Supporting information

Supplementary Results 1

Supplementary Results 2

Supplementary Results 3

## 7. List of abbreviations

AD: Anaerobic Digestion
ICNP: International Code of Nomenclature of Prokaryotes
GTDB: Genome Taxonomy Database
MAG: Metagenome assembled genome
LPSN: List of Prokaryotic Names with Standing in Nomenclature
RED: Relative evolutionary divergence
ANI: Average nucleotide identity
PA: Phenyl acids
OFMSW: Organic fraction of municipal solid waste
WWTP: Wastewater treatment plant
PAA: Phenylacetic acid
PPA: Phenylpropionic acid
PBA: Phenylbutyric acid
GC-TCD: Gas chromatograph equipped with a thermal conductivity detector
VFA: Volatile fatty acids
SEM: Scanning electron microscopy
PBS: Phosphate buffer solution
MM: Materials and Methods
LCA: Least common ancestor
NCBI: National Center for Biotechnology Information
PFBA: Parsimonious flux-balance analyses
MTF: Minimization of total flux
AF: Alignment fraction
PPP: Pentose phosphate pathway
RA: Relative abundance
LDH: Lactate dehydrogenase
Etf: Electron-transferring flavoprotein

## 9. Author information

### 9.1. Contributions

AOW and EMP isolated strain PM69 and organised the deposition of the strain. EMP, MW, AM, and AOW were responsible for the culture maintenance and assessment of growth characteristics. EMP, MW, JZ did light microscopic analyses and AS conducted SEM and ABRicate analyses. EMP and MMH carried out biochemical analyses. MW and JZ performed *Nanopore* sequencing incl. library preparations. EMP was responsible for *Illumina* sequencing, processed *Sanger* and whole genome sequences, performed genome assembly, and carried out metabolic modelling. Phylogenetic analyses were conducted by ZD (whole genome) and AS (16S rRNA gene). EMP and AOW secured funding. EMP drafted the manuscript. AN, SSY, CR and AOW supervised the findings of the study. All authors reviewed and approved the manuscript.

## 10. Acknowledgements

Computational results have been achieved using the LEO5 HPC infrastructure of the University of Innsbruck. We especially thank Hermann Schwärzler for his kind support.

This research was funded in part by the alpha+ Foundation and Austrian Science Fund (grant number ESP7170024). This work was also supported in part by the Austrian Science Fund grant P36711.

## 11. Supplementary information

Supplementary Methods

Supplementary Results 1

Supplementary Results 2

Supplementary Results 3

## 12. Availability of data and material

The raw sequence reads (Illumina, ONT) as well as Sanger sequences and the quality-filtered assembly (contigs > 1kb) have been submitted to the European Nucleotide Archive (ENA) under study accession number PRJEB101082. Biochemical data can be obtained from the open-access digital repository *Zenodo* (10.5281/zenodo.17396118). After peer review, type strain PM69 can be purchased from DSMZ – German Collection of Microorganisms and Cell Cultures GmbH (DSM 121036) as well as from the Japan Collection of Microorganisms (JCM 39671).

