## Supplementary Methods for "*Thermoaminiphila catenidiffluenda* gen. nov., sp. nov.: A novel thermophilic, strictly anaerobic bacterium representing *Thermoaminiphilia* class nov., a newly described class thriving in hydrocarbon-rich habitats and biogas fermenters"

Title

Author Information

Eva Maria Prem[1*], Mathias Wunderer[1], Andja Mullaymeri[1], Julia Zoehrer[1], Zuzanna Dutkiewicz[1], Abhijeet Singh[2,4,5], Mahmoud M. Habashy[3], Anna Neubeck[4], Sepehr Shakeri Yekta[3], Christian Rinke[1], and Andreas Otto Wagner[1]

[1] Department of Microbiology, Universität Innsbruck, Innsbruck, Austria
[2] Department of Molecular Sciences, Swedish University of Agricultural Sciences, Uppsala, Sweden
[3] Department of Thematic Studies, Environmental Change, Linköping University, Sweden
[4] Department of Earth Sciences, Uppsala University, Uppsala, Sweden
[5] Faculty of Agricultural Sciences and Technology, Ganpat University, India

**MM1.** Separation of strain PM69

Culture PM69 derived from a thermophilic, 250 mL semi-batch lab reactor system anaerobically digesting phenyl acids (PA) in a long-term experiment. The inoculum of the batch reactor derived from an organic fraction of municipal solid waste (OFMSW) digesting reactor (Roppen, Tyrol) operated at thermophilic temperatures. We incubated the sludge for several weeks to remove easily degradable residuals. Thereafter, we diluted it with deionised water (1:4) and stored at 4°C. All growth media were adjusted to pH 7.2 after PA addition and subsequently autoclaved. For stage 1, we prepared anaerobic 250 mL serum flasks with 60 mL start medium containing 8 mM phenylacetic acid (PAA), 8 mM 3-phenylpropionic acid (3-PPA) and 8 mM 2-phenylbutyric acid (2-PBA), 15 mM sodium formate, 2.4 mM sodium acetate, 3.7 mM ammonium chloride, 0.02% (w/v) of yeast extract, and trace elements and vitamins (DSM 119), as well as nitrogen as headspace gas. Reactors were sealed with butyl-rubber stoppers and aluminium caps. We added 60 mL of the respective inoculum to the reactors and incubated the communities at 55°C for one year. Reactors were biochemically monitored (pH, biogas production, methane concentration, concentrations of volatile fatty and phenyl acid), close meshed in the first three months. In stage 2, we changed from batch to semi-batch feeding (2 mM PA, 2.5 – 5% volume exchange), whereby substrate amount, medium replacement and feeding interval depended on the methane production and VFA concentrations of the reactors elaborated in stage 1. From stage 3 onwards, also in semi-batch mode, the cultures did not receive any more sodium formate and acetate and had to solely grow on phenyl acids as carbon source and minimal medium. Yeast extract addition was kept to a minimum (0.02% (w/v)). After 34 months, we prepared an anaerobic, PA containing medium agar (per litre: 12 mM PAA, 10 mM 2-PPA, 10 mM 3-PPA, 9 mM 3-PBA, 9 mM 4-PBA, 0.4 g $MgCl_2$ x 6 $H_2O$, 0.4 g NaCl, 0.4 g $NH_4Cl$, 0.2 g $MgSO_4$ x 7 $H_2O$, 0.1 g yeast extract, 0.05g $CaCl_2$ x 2 $H_2O$, 1 mL trace element solution, 1 mL vitamin solution (DSM119), 15 g agar, pH 7.5) in 120 mL serum flasks sealed with butyl rubber stoppers and aluminium caps. Via syringe and cannula, we added 100 µL of the above-mentioned batch culture to still liquid but already viscous medium, thoroughly mixed and immediately cooled down the medium to prevent excessive settling of cells. The serum flask was incubated at 55°C for 4 months. Thereafter, we prepared an anaerobic hood constantly flushed with sterile nitrogen and carefully opened agar flasks. With a 120 mm long cannula and syringe, we picked and transferred the culture to a new infusion flask containing Starkey's medium (per litre: 5 g peptone, 3 g meat extract, 0.2 g yeast extract, 5 g glucose, 1.5 g $MgSO_4$ x 7 $H_2O$, 0.1 g $NH_4Fe(SO_4)_2$, pH 7.4, headspace: 70% $N_2$, 30% $CO_2$). The culture was then incubated at 55°C for additional 2 months. After multiple generations, the medium was transitioned to a more defined formula (per litre: 5 g D-glucose, 4 g casamino acids, 1 g yeast extract, 0.5 g L-cysteine, 1.5 g $MgSO_4$ x 7 $H_2O$, 15 mL phosphate solution Japan Collection of Microorganisms (JCM) 770 [1], 12 mL mineral salt solution JCM 770 [1], 0.5 mL resazurin solution, 1 mL selenite-tungstate solution JCM 431 [2], 1mL trace element solution JCM 243 [3],

1 mL vitamin solution JCM 770 [1](added after autoclaving), pH 7.4 ± 0.2, headspace: 70% $N_2$, 30% $CO_2$, inoculation volume 5 – 10%). We prepared anaerobic media according to previous protocols [4].

**MM2.** Cellular fatty acids

We centrifuged 250 mL of culture at 4.000 x g for 10 minutes and resuspended 600 mg of wet biomass in 9 mL isopropanol (99.9% v/v). The conversion to fatty acid methyl esters (FAME) and GC measurements (GC-FID and -MS) were carried out by DSMZ Services, Leibnitz-Institut DSMZ – Deutsche Sammlung von Mikroorganismen und Zellkulturen GmbH, Braunschweig, Germany.

**MM3.** Metabolite screening of cultures at two different growth stages

For an extended metabolite screening via HPLC and GC, we centrifuged a 11- to 20-day old culture pool ("Culture 1"), a 7-day old culture pool ("Culture 2"), as well as uncultivated medium at 7,000 g for 10 mins and filtrated (0,20 µm) the supernatant into fresh falcon tubes. The pellet of culture 2 ("cells") was washed with 30 mL phosphate buffer twice (with a final decanting step). Samples were stored at -20°C and shipped frozen to Linköping, Sweden.

The short chain carboxylic acid concentration (including caproic "$C_6$" and heptanoic "$C_7$" acids) was analysed using a gas chromatograph with a flame-ionization detector (Agilent 8860 GC System, Agilent Technologies, Santa Clara, California, USA) according to previous protocols [5] with some modifications. The analysis was performed at a constant pressure of 10.8 psi and nitrogen as a carrier gas. The temperature program was set at 80°C then increased to 150°C (5°C/min) followed by a further increase to 200°C (10°C/min). Moreover, the presence of medium chain carboxylic acid ($C_8$ to $C_{12}$) was assessed using a high-performance liquid chromatograph (HPLC) with a Shim-pack GIST C18-AQ column and an SPD-40V UV-VIS detector (Nexera, Shimadzu, Kyoto, Japan). The column and detector were operated at 40°C while using 10% phosphate buffer (pH 2.1) and 90% methanol as a mobile phase at a flow rate of 0.7 mL/min. The ethanol and lactic acid concentrations were measured using the same HPLC and column, where SPD-40V UV-VIS detector was used for the lactic acid measurement and RID-20A refractive index detector was used for the ethanol measurement. The analyses of ethanol and lactic acid were done according to Westerholm et al. [6] with some modifications. The sample was centrifuged at 12000 for 10 minutes, followed by addition of 70 µL of 5 M $H_2SO_4$ to 700 µL of the supernatant. Subsequently, the sample was frozen. After another centrifugation step (12000 for 10 minutes), the supernatant was filtered into a glass vial using 0.2 µm syringe filter. The column and detectors were operated at 40°C while using 100% phosphate buffer (pH 2.1) as a mobile phase at a flow rate of 0.7 mL/min.

**MM4.** Scanning electron microscopy (SEM)

Media solution was removed, and microbial cells were separated on a 0.8 µm nylon membrane filter (Whatman®) under vacuum. The filter was then transferred and submerged into a solution of glutaraldehyde (4 ml, 25 % Glutaraldehyde, Sigma-Aldrich), phosphate buffer solution (0.4 ml of 10 M PBS, Roche) in deionized water (32 ml) for four hours. To remove glutaraldehyde the filter was submerged and washed with PBS (0.01 M), thrice for 10 minutes each. The ethanol gradient (50 %, 70 %, 80 %, 85 %, 90 %, 95 % and 100 %) drying method was used to desiccate the cells. The filter was submerged in respective ethanol solutions for 10 minutes each. The final drying step was repeated thrice with 100 % ethanol for 10 minutes. The filter was submerged for 30 seconds in hexamethyldisilane (≥99 % HMDS, Sigma-Aldrich) and solution was removed with a pipette followed by drying overnight. A small part of the filter was dissected, mounted and coated with gold-palladium for 180 seconds for the field emission SEM analysis (Zeiss Supra 35 VP, Zeiss, Germany). SEM images were processed and pseudo-coloured with GIMP (v3.0.4).

**MM5.** Sanger sequencing of the 16S rRNA gene

Genomic DNA of two 4-day old cultures (6 mL each) was extracted with the Monarch Spin gDNA Extraction Kit with following modifications: After enzymatic treatment on a thermo-block, samples underwent overnight digestion at 55°C. DNA concentration and purity were assessed using a NanoDrop 2000c spectrophotometer (Thermo Fisher Scientific, USA). The 16S rRNA gene (primer pair 27-f / 1492-r) was amplified using the Q5® High-Fidelity 2X Master Mix (NEB, USA) according to the manufacturer's protocol. After PCR clean-up (Monarch® Spin PCR & DNA Cleanup Kit (5 µg), NEB, USA), we checked amplificants on a QIAxcel Instrument using the QIAxcel DNA Screening Kit (Qiagen: RRID:SCR008539, Netherlands). For Sanger sequencing, we split each sample and added either the forward or the reverse primer to create premixed samples and sent those to Eurofins Genomics, Germany. As preliminary check, we assessed the presence of *Archaea* by using the primer pair 109-f / 1492-r.

For preparing the 16S tree, forward and reverse sequences (reverse complements) from each sample were assembled into consensus sequences using *AliView*[7]. Consensus sequences were queried individually against the NCBI nucleotide database using BLASTn, and closely related reference sequences with > 95 % query coverage were retrieved. Query and reference sequences were combined into a single dataset, and duplicate sequences were removed using *DupRemover*[8]. We performed multiple sequence alignment using MAFFT (v7.490). Maximum-likelihood phylogenetic inference was conducted using IQ-TREE (v2.0.7)[9] The best-fitting nucleotide substitution model (TIM3+F+I+G4) was selected using ModelFinder in automatic mode and used for tree inference. Tree search was performed under the selected model using the IQ-TREE stochastic algorithm including thorough nearest-neighbor interchange (NNI) moves and polytomy collapse. Branch support was assessed using 100 nonparametric bootstrap replicates and 100 jackknife replicates, and node support was additionally evaluated using Transfer Bootstrap Expectation (TBE) values calculated from bootstrap trees. Branch lengths were optimized during bootstrap analysis. The final phylogenetic tree was visualized and annotated using package *ggtree* (v4.0.4)[10].

**MM6.** Check for growth on specific carbon sources via *gapseq* and the *Additional Constraints* pipeline

```
#Additional constraints vignette
#checking growth exclusively on specific carbon sources, e.g., on fructose:

mod2_fru <- changeBounds(mod2, react = c("EX_cpd00027_e0", "EX_cpd00082_e0"), lb = c(0,-5))
mtf2_fru <- pfba(mod2_fru)
ex2_fru <- findExchReact(mod2_fru)
metname2_fru <- mod2_fru@met_name
dist2_fru <- mtf2_fru@fluxes
ex.dt2_fru <- data.table(ex=ex2_fru$react_id, met=metname2_fru[ex2_fru$met_pos],
flux=dist2_fru[ex2_fru$react_pos])
substrates2_fru <- ex.dt2_fru[flux<0][order(flux)] #substrates
products2_fru <- ex.dt2_fru[flux>0][order(flux)] #products


#removing reactions from the model
mod2_1 <- rmReact(mod2, "EX_cpd00076_e0") #sucrose
mod2_2 <- rmReact(mod2_1, "EX_cpd00208_e0", rm_met = TRUE) #lactose
mod2_3 <- rmReact(mod2_2, "EX_cpd00179_e0", rm_met = TRUE) #maltose
#all three reactions added in gapseq step 7


#list reactions
react_id2_3 <- table(mod2_3@react_id)
write.xlsx(react_id2_3, "react_id2_3.xlsx")



#parsimonious flux-balance analyses (PFBA) with minimization of total flux (MTF)
mtf2_3 <- pfba(mod2_3)
ex2_3 <- findExchReact(mod2_3)
metname2_3 <- mod2_3@met_name
dist2_3 <- mtf2_3@fluxes
ex.dt2_3 <- data.table(ex=ex2_3$react_id, met=metname2_3[ex2_3$met_pos],
flux=dist2_3[ex2_3$react_pos])
substrates2_3 <- ex.dt2_3[flux<0][order(flux)] #substrates
products2_3 <- ex.dt2_3[flux>0][order(flux)] #products


#another check with new model, e.g., with fructose:
mod2_3_fru <- changeBounds(mod2_3, react = c("EX_cpd00027_e0", "EX_cpd00082_e0"), lb = c(0,-5))
mtf2_3_fru <- pfba(mod2_3_fru)
ex2_3_fru <- findExchReact(mod2_3_fru)
metname2_3_fru <- mod2_3_fru@met_name
dist2_3_fru <- mtf2_3_fru@fluxes
ex.dt2_3_fru <- data.table(ex=ex2_3_fru$react_id, met=metname2_3_fru[ex2_3_fru$met_pos],
flux=dist2_3_fru[ex2_3_fru$react_pos])
substrates2_3_fru <- ex.dt2_3_fru[flux<0][order(flux)] #substrates
```

```
products2_3_fru <- ex.dt2_3_fru[flux>0][order(flux)] #products
```

**Table MM1.** Class SHA-98 MAGs recovered from the same or additional, oligotrophic batch reactors. Bin in bold was recovered from the same batch reactor as strain PM69 and was also classified as s__Ch115 sp013178415. Most bins derived from the OFMSW digester (black bold frame). Bins in bold red frame were recovered from the same batch reactor type.

| MAG | Preliminary bin name | Batch reactor temperature | Sludge origin | Temperature large-scale reactor |
|---|---|---|---|---|
| PM003_bin.28.strict | s__UBA4971 sp033809295 | 37°C | WWTP | 37°C |
| PM003_bin.81.permissive | g__UBA4971 unknown species | 37°C | WWTP | 37°C |
| PM004_bin.15.strict | f__UBA4971 unknown species | 37°C | OFMSW | 55°C |
| PM004_bin.45.permissive | g__UBA4971 unknown species | 37°C | OFMSW | 55°C |
| PM006_bin.39.orig | g__UBA4971 unknown species | 37°C | OFMSW | 55°C |
| **PM007_bin.19.strict** | **s__Ch115 sp013178415** | **55°C** | **OFMSW** | **55°C** |
| PM007_bin.27.permissive | g__DUSO01 unknown species | 55°C | OFMSW | 55°C |
| PM008_bin.17.strict | s__UBA4971 sp900019985 | 55°C | OFMSW | 55°C |
| PM008_bin.51.permissive | c__SHA-98 unknown species | 55°C | OFMSW | 55°C |
| PM008_bin.53.orig | s__UBA6256 sp012516125 | 55°C | OFMSW | 55°C |
| PM008_bin.65.orig | o__UBA4971 unknown species | 55°C | OFMSW | 55°C |
| PM009_bin.22.orig | s__UBA6256 sp012842395 | 55°C | OFMSW | 55°C |
| PM012_bin.13.permissive | g__UBA6256 unknown species | 55°C | WWTP | 37°C |
