## Supplementary Results 1 for "*Thermoaminiphila catenidiffluenda* gen. nov., sp. nov.: A novel thermophilic, strictly anaerobic bacterium representing *Thermoaminiphilia* class nov., a newly described class thriving in hydrocarbon-rich habitats and biogas fermenters"

Title

Author Information

Eva Maria Prem[1*], Mathias Wunderer[1], Andja Mullaymeri[1], Julia Zoehrer[1], Zuzanna Dutkiewicz[1], Abhijeet Singh[2,4,5], Mahmoud M. Habashy[3], Anna Neubeck[4], Sepehr Shakeri Yekta[3], Christian Rinke[1], and Andreas Otto Wagner[1]

[1] Department of Microbiology, Universität Innsbruck, Innsbruck, Austria
[2] Department of Molecular Sciences, Swedish University of Agricultural Sciences, Uppsala, Sweden
[3] Department of Thematic Studies, Environmental Change, Linköping University, Sweden
[4] Department of Earth Sciences, Uppsala University, Uppsala, Sweden
[5] Faculty of Agricultural Sciences and Technology, Ganpat University, India

**Table S1.** VFA production of difference generations of *T. catenidiffluena strain PM69*. Samples were taken at different ages. "Medium" means VFA concentrations in the non-inoculated medium.

| **Sample** | **Generation** | **Age of Culture** [d] | **Formate** [mM] | **Acetate** [mM] | **i-Butyrate** [mM] | **Butyrate** [mM] |
|---|---|---|---|---|---|---|
| Medium | -- | -- | 0.00 | 0.00 | 0.00 | 0.00 |
| Culture T. catenidiffluenda strain PM69 | 31 | 3 | 0.00 | 7.48 | 0.62 | 0.00 |
| | 53 | 4 | 0.01 | 10.54 | 0.00 | 0.11 |
| | 20 | 7 | 0.00 | 7.46 | 0.00 | 0.07 |
| | 21 | 7 | 0.04 | 9.83 | 0.00 | 0.11 |
| | 22 | 7 | 0.00 | 9.64 | 0.00 | 0.00 |
| | 23 | 7 | 0.00 | 10.06 | 0.00 | 0.00 |
| | 25 | 7 | 0.00 | 9.94 | 0.00 | 0.00 |
| | 26 | 7 | 0.00 | 9.00 | 0.00 | 0.00 |
| | 27 | 7 | 0.00 | 5.30 | 0.00 | 0.00 |
| | 28 | 7 | 0.41 | 8.43 | 0.00 | 0.00 |
| | 52 | 7 | 0.02 | 11.91 | 0.00 | 0.10 |
| | 54 | 7 | 0.00 | 11.20 | 0.00 | 0.11 |
| | 55 | 7 | 0.02 | 10.81 | 0.00 | 0.10 |
| | 56 | 7 | 0.02 | 11.30 | 0.00 | 0.11 |
| | 57 | 7 | 0.00 | 10.86 | 0.00 | 0.00 |
| | 58 | 7 | 0.00 | 3.83 | 0.00 | 0.00 |
| | 59 | 7 | 0.00 | 5.31 | 0.00 | 0.00 |
| | 59 | 7 | 0.00 | 15.20 | 0.00 | 0.00 |
| | 60 | 7 | 0.00 | 8.65 | 0.00 | 0.00 |
| | 61 | 7 | 0.00 | 10.11 | 0.00 | 0.00 |
| | 62 | 7 | 0.00 | 10.88 | 0.00 | 0.00 |
| | 63 | 7 | 0.63 | 9.73 | 0.00 | 0.00 |
| | 24 | 10 | 0.00 | 10.90 | 0.00 | 0.00 |
| | 51 | 11 | 0.00 | 10.81 | 0.00 | 0.07 |
| | 50 | 14 | 0.02 | 10.74 | 0.00 | 0.07 |
| | 49 | 17 | 0.03 | 10.84 | 0.00 | 0.07 |
| | 48 | 18 | 0.04 | 11.87 | 0.00 | 0.09 |
| | 47 | 21 | 0.02 | 10.54 | 0.00 | 0.07 |
| | 46 | 24 | 0.00 | 10.15 | 0.00 | 0.09 |
| | 45 | 26 | 0.00 | 10.34 | 0.00 | 0.10 |
| | 44 | 28 | 0.00 | 11.01 | 0.00 | 0.10 |
| | 43 | 32 | 0.00 | 10.04 | 0.00 | 0.09 |
| | 42 | 35 | 0.00 | 10.31 | 0.00 | 0.08 |
| | 40 | 42 | 0.06 | 10.20 | 0.00 | 0.08 |
| | 39 | 45 | 0.00 | 10.09 | 0.00 | 0.10 |
| | 38 | 47 | 0.00 | 10.49 | 0.00 | 0.09 |
| | 37 | 49 | 0.07 | 10.64 | 0.00 | 0.09 |
| | 36 | 52 | 0.03 | 9.67 | 0.00 | 0.09 |
| | 35 | 55 | 0.08 | 11.35 | 0.00 | 0.09 |
| | 30 | 63 | 0.04 | 8.76 | 0.00 | 0.05 |
| | 32 | 63 | 0.15 | 10.23 | 0.00 | 0.11 |
| | 31 | 66 | 0.12 | 10.99 | 0.00 | 0.10 |

**Table S2.** Extended metabolite screening for *Thermoaminiphila* medium (“Medium”), a 11-to-20-day old culture pool (“Culture 1”), and a 7-day old culture pool (“Culture 2”)

| | Concentration (mM) | | | | | | | | | | | | | | Concentration (gL) | |
|---|---|---|---|---|---|---|---|---|---|---|---|---|---|---|---|---|
| | **Acetic acid** | **Propionic acid** | **Iso-butyric acid** | **Butyric acid** | **Iso-valeric acid** | **n-valeric acid** | **Iso-caproic acid** | **n-caproic acid** | **Heptanoic acid** | **Caprylic acid** | **Nonanoic acid** | **Capric acid** | **Undecanoic acid** | **Lauric acid** | **Lactic acid** | **Ethanol** |
| Medium | 1.0 | 0.0 | 0.0 | 0.0 | 0.0 | 0.0 | 0.0 | 0.0 | 0.0 | 0.0 | 0.0 | 0.0 | 0.0 | 0.0 | 0.1 | 0.0 |
| Culture 1 | 9.6 | 0.0 | 0.0 | 0.0 | 0.1 | 0.0 | 0.0 | 0.0 | 0.0 | 0.0 | 0.0 | 0.0 | 0.0 | 0.0 | 0.3 | 0.5 |
| Culture 1 | 9.7 | 0.0 | 0.0 | 0.0 | 0.0 | 0.0 | 0.0 | 0.0 | 0.0 | 0.0 | 0.0 | 0.0 | 0.0 | 0.0 | 0.1 | 0.5 |
| Culture 2 | 9.1 | 0.0 | 0.0 | 0.0 | 0.0 | 0.0 | 0.0 | 0.0 | 0.0 | 0.0 | 0.0 | 0.0 | 0.0 | 0.0 | 0.2 | 0.5 |
| Culture 2 | 9.5 | 0.0 | 0.0 | 0.0 | 0.0 | 0.0 | 0.0 | 0.0 | 0.0 | 0.0 | 0.0 | 0.0 | 0.0 | 0.0 | 0.2 | 0.4 |

**Table S3.** Composition of cellular fatty acids via conversion to fatty acid methyl esters (FAME) for strain PM69

| Fatty acids | % |
|---|---|
| **saturated** | |
| 11:0 iso 3OH | 1.40 |
| 13:0 iso | 0.70 |
| 13:0 anteiso | 0.50 |
| 12:0 3OH | 0.50 |
| 14:0 | 1.00 |
| 13:0 iso 3OH | 7.70 |
| 13:0 2OH | 3.50 |
| 15:0 iso | 29.1 |
| 15:0 anteiso | 17.2 |
| 16:0 iso | 0.50 |
| 16:0 | 6.00 |
| 17:0 iso | 19.1 |
| 17:0 anteiso | 8.70 |
| 18:0 | 1.10 |
| 19:0 iso | 0.40 |
| 19:0 anteiso | 0.30 |
| 19:0 cyclo w8c | 0.40 |
| **unsaturated** | |
| Summed feature 3 (16:1 w7c/16:1 w6c) | 0.20 |
| Summed feature 8 (18:1 w7c or 18:1 w6c) | 0.40 |
| 18:1 w9c | 1.20 |

**Table S4**. Basic quality assessment of *Illumina* shot-gun sequences via **fastqc** prior and after filtering

| | Quality filtering | Total sequences | Total bases [Mbp] | Sequence length [bp] | Poor quality sequences | GC % |
|---|---|---|---|---|---|---|
| *Illumina* R1 fastq file | prior | 4663453 | 616.3 | 10 - 151 | 0 | 51 |
| | after (paired reads) | 4486933 | 507.9 | 36 - 125 | 0 | 51 |
| *Illumina* R2 fastq file | prior | 4663453 | 617.9 | 10 - 151 | 0 | 50 |
| | after (paired reads) | 4486933 | 507.5 | 36 - 125 | 0 | 50 |

**Table S5**. Quality assessment of the assembly with and without contigs <1kb via ***QUAST** and* ***tRNAscan-SE***

| Parameters | Assembly | Assembly (removal of <1kb contigs) |
|---|---|---|
| # contigs (>= 0 bp) | 77 | 19 |
| # contigs (>= 1000 bp) | 19 | 19 |
| # contigs (>= 5000 bp) | 14 | 14 |
| # contigs (>= 10000 bp) | 14 | 14 |
| # contigs (>= 25000 bp) | 13 | 13 |
| # contigs (>= 50000 bp) | 10 | 10 |
| Total length (>= 0 bp) | 3319540 | 3303196 |
| Total length (>= 1000 bp) | 3303196 | 3303196 |
| Total length (>= 5000 bp) | 3292864 | 3292864 |
| Total length (>= 10000 bp) | 3292864 | 3292864 |
| Total length (>= 25000 bp) | 3277704 | 3277704 |
| Total length (>= 50000 bp) | 3155302 | 3155302 |
| # contigs | 29 | 19 |
| Largest contig | 791104 | 791104 |
| Total length | 3309741 | 3303196 |
| GC (%) | 51.32 | 51.32 |
| N50 | 381840 | 381840 |
| N90 | 183514 | 183514 |
| auN | 416810.2 | 417634.8 |
| L50 | 3 | 3 |
| L90 | 9 | 9 |
| # total reads | 8973866 | 8973866 |
| # left | 4486933 | 4486933 |
| # right | 4486933 | 4486933 |
| Mapped (%) | 98.65 | 98.35 |
| Properly paired (%) | 97.9 | 97.65 |
| Avg. coverage depth | 301 | 301 |
| Coverage >= 1x (%) | 100 | 100 |
| # N's per 100 kbp | 0 | 0 |
| **tRNAscan-SE** | | |
| # tRNAs (standard 20 AA) | 47 | 45 |
| # Selenocysteine rRNAs | 1 | 1 |
| # predicted pseudogenes | 1 | 1 |

**Table S6**. Quality assessment of assemblies via ***Bandage***

| Parameters | Assembly | Assembly (removal of <1kb contigs) |
|---|---|---|
| **Graph size** | | |
| Node count | 127 | 19 |
| Edge count | 165 | 0 |
| Edge overlaps | 0 bp | n/a |
| Total length | 3 320 572 bp | 3 303 196 bp |
| **Node size** | | |
| N50 | 381 840 bp | 381 840 bp |
| Shortest node | 1 bp | 1 052 bp |
| Lower quartile node | 22bp | 9 322 bp |
| Median node | 140 bp | 57 405 bp |
| Upper quartile node | 371 bp | 245 107 bp |
| Longest node | 791 104 bp | 791 104 bp |
| **Graph connectivity** | | |
| Dead ends | 0 | 38 |
| Connected components | 1 | 19 |
| Largest component | 3 320 572 bp (100%) | 791 104 bp (23.95%) |
| Total length orphaned nodes | 0 bp (0%) | 3 303 196 bp (100%) |
| **Depth** | | |
| Median depth | 1.00 x | 1.00 x |
| Estimated sequence length | 3 332 294 bp | 3 303 196 bp |

**Table S7.** Predicted (partial) rRNA sequences of PM69 found using all contigs via *barrnap v0.9*.

| rRNA | Contig | Length [bp] | Note |
|---|---|---|---|
| **16S ribosomal RNA (partial)** | **19** | **1051** | **aligned 66 % of the 16S ribosomal RNA** |
| >16S_rRNA: 19:0-1051(+)<br>AAGCCTGACCGAGCGACGCCGCGTGAGGGAAGAAGGTCTTCGGATTGTAAACCTCTGTCTTGGGGGATGAGAAAGGACAGTACCCCAGGAGGAAGCCCCGGCTAACTACGTGCCAGCAGCCGCGGT<br>AATACGTAGGGGGCGAGCGTTGTCCGGAATTACTGGGCGTAAAGGGCGTGCAGGTGGTCTCTTAAGTTAGGTGGGAAATCCCATAGCTCAACTATGGGGGTGCGCCTAAAACTGGGGGGCTAGAGGG<br>CAAGAGAGGGAAGCGGAATTCCCGGTGTAGCGGTGAAATGCGTAGATATCGGGAGGAACACCAGTGGCGAAGGCGGCTTCCTGGATTGCACCTGACACTGAGGCGCGAAAGCCAGGGGAGCGAACG<br>GGATTAGATACCCCGGTAGTCCTGGCCGTAAACGATGGATGCTAGGTGTGGGAGGTATCGACCCCTTCCGTGCCGCAGCTAACGCATTAAGCATCCCGCCTGGGGAGTACGGCCGCAAGGTTGAAAC<br>TCAAAGGAATTGACGGGGGCCCGCACAAGCGGTGGAGCATGTGGTTTAATTCGACGCAACGCGAAGGACCTTACCAGGGTTTGACATGCTGGTGGTACTGAACCGAAAGGGGAAGGACCCAGGCAA<br>CTGGGAGCCAGCACAGGTGGTGCATGGCTGTCGTCAGCTCGTGTCGTGAGATGTTGGGTTAAGTCCCGCAACGAGCGCAACCCCTACATTCAGTTGCTAACGGGTAGAGCCGAGCACTCTGGATGGAC<br>TGCCGGGGATGACCCGGAGGAAGGTGGGGATGACGTCAAGTCATCATGCCCTTTATGCCCTGGGCCACACACGTGCTACAATGGCCTGTACAGAGGGAGGCAAACCCGCGAGGGGGAGCGGATCCC<br>AAAAAGCAGGTCTAAGTTCGGATCGCAGGCTGCAACTCGCCTGCGTGAAGCCGGAATCGCTAGTAATCGCGGGTCAGCATACCGCGGTGAATACGTTCCCGGGCCTTGTACACACCGCCCGTCACAC<br>CACGAAAGTTTGTCACACCCGAAGCCGGTGGCCTAACC | | | |
| **rRNA** | **Contig** | **Length [bp]** | **Note** |
| **23S ribosomal RNA (partial)** | **16** | **2362** | **aligned 73 % of the 23S ribosomal RNA** |
| >23S_rRNA: 16:2-2362(-)<br>GCCTGACTGCGTACTTTTTGTAGAACGGACCGGCGAGTTACAGTATGTGGCGAGGTTAAGCAGGAGATGCGGAGCCGCAGCGAAAGCGAGTCTTAAGAGGGCGCAAGTCGCATGCTGTAGACCCGAA<br>ACCGGGTGATCTATCCATGGCCAGGGTGAAGCGGAGTTAAAACTCCGTGGAGGCCCGAACCACGTTGTCGTTGAAAAGGCATGGGATGAGCTGTGGATAGGGGTGAAATGCCAATCGAACTCGGAG<br>ATAGCTGGTTCTCCCCGAAATAGCTTTAGGGCTAGCCTTGGGTAAGAGAGTTATGGAGGTAGAGCACTGATCGGGCTAGGGGCCCAAAAAGGTTACCGAACCCTTTCAAACTCCGAATTCCATAAGTT<br>GTTGCCCAGGAGTCAGACTGTGGGGGCTAAGCTTCATAGTCGAGAGGGAAACAGCCCAGATCGTCTGCTAAGGTCCCAAAGTATGAGCTAAGTGGGAAAGGATGTAGGATTGCACAGACAGCCAGG<br>ATGTTGGCTTAGAAGCAGCCATTCATTCAAAGAGTGCGTAATAGCTCACTGGTCGAGTGATTCTGCGCCGAAAATGTAACGGGGCTGAAGCTCATCACCGAAGCAGCGGAATACCGGAAGGTATTGG<br>TAGGGGAGCGTTGTGCCGGAGTAGAAGCAGGACGGGAACGACCTGTGGATCCGACACAAGTGAGAATGCCGGTATAAGTAGCGAAAAGAGGGGTGAGAATCCCCTTCGCCGAAAGCCTAAGGTTTC<br>CTGAGGAAGGGTCGTCCTCTCAGGGTTAGTCGGGGCCTAAGCCGAGGCTGGATAGCGTAGGCGATGGACAACCGGTTAATATTCCGGTACCACCGAGAGCCGGCCATGGTGAATATGTCTGGAATGG<br>AGGGGCGCAAGTTCCGAAAGGAAGGGCATATGAGCCAAGCAATGGGGTGACGCAGGAGGGTAGGCCATCAGGCTGATGGAATAGCCTGTCCAAGCGAGTAGGGGGTGCGGCAGGGAAATCCGCTG<br>CACAAGACCCTGAGGCGCGATGGGGAGCGAAATTATAGTAGCGGAACTGGTTGAACCCATACTGCCTAGAAAAGCCTCTAGCGAGGAAGCAGGTGCCCGTACCGCAAACCGACACAGGTAGGCGGG<br>GAGAGAATCCTAAGGCGCGCGAGAGAACCCCTGTTAAGGAACTCGGCAAAATGATCCCGTAACTTCGGGAGAAGGGATGCCTCGGTAGGGTGATAGTGAGAGCTAAAGCCCGAGGAGGTCGCAGAG<br>AAACGGCCCAAGCGACTGTTTATCAAAAACACAGGTCTCTGCTAAGTCGAAAGACGAAGTATAGGGGCTGACGCCTGCCCAGTGCCGGAAGGTTAAGGGGAGGGGTTAGGGGAACCGAAGCTCCGA<br>ACCAAAGCCCCGGTGAACGGCGGCCGTAACTATAACGGTCCTAAGGTAGCGAAATTCCTTGTCGGGTAAGTTCCGACCCGCACGAATGGCGTAACGACTTGGGCGCTGTCTCGATGGGGGGCTCGGC<br>GAAATTGTAAGTACCTGTGAAGATGCAGGTTACCCACGACTAGACGGAAAGACCCCGTGGAGCTTTACTGTAGCTTGAGATTGGATTTTGGCATGTTATGTACAGGATAGGTGGGAGGCGTAGAAGG<br>AGGGGCGCTAGCTCTTCTGGAGCCGGCCTTGGGATACCACCCTTAATGTGCTGGAGTTCTAACCCTGAGCAGTGATCCTGCCAGGGGACATTCTCAGGTGGGCAGTTTGACTGGGGCGGTCGCCTCCT<br>AAAGGGTAACGGAGGCGCCCAAAGGTTCCCTCAGCGCGGTTAGAAATCGCGCGTTGAGTGTAAAGGCAGAAGGGAGCTTGACTGCGAGAGACACAGCTCGAGCAGGGTGGAAACACGGCTTAGTG<br>ATCCGGTGGCGCTGAGTGGGCTGGCCATCGCTCAACGGATAAAAGCTACCCCGGGGATAACAGGCTGATCTCCCCCAAGAGTCCACATCGACGGGGAGGTTTGGCACCTCGATGTCGGCTCATCGCA<br>TCCTGGGGCTGAATTAGGTCCCAAGGGTTGGGCTGTTCGCCCATTAAAGCGGTACGTGAGCTGGGTTCAGAACGTCGTGAGACAGTTCGGTCCCTATCTGTCGTGGGCGCAGGAGATTTGAGGGGATC<br>TGTCCCTAGTACGAGAGGACCGGGATGGACATACCTCTGGTGTACCAGTTGTCCCGCCAGGGGCACGGCTGGGTAGCCAAGTATGGACGGGATAAACGCTGAAAGCATCTAAGCGTGAAGCCCTCCC<br>CAAGATTAGATCTCCCACAGGGTCGACCTGGTAAGACCCCAGCGAGAAGAGCTGGTAGATAGGCCGGGTGTGTAAGGGCA | | | |
| **rRNA** | **Contig** | **Length [bp]** | **Note** |
| **5S ribosomal RNA** | **47** | **110** | **aligned 73 % of the 23S ribosomal RNA** |
| **>5S_rRNA: 47:43-153(-)**<br>GGTGGCAATGACGGAGGGGAAACACCTGTTCCCATTCCGAACACAGAAGTTAAGCCCTCCAGTGCCGATGGTACTGCGCTGGCGACGGCGTGGGAGAGTAGGTCGCTGCC | | | |

**Table S8.** Elaborated 16S rRNA sequence of strain PM69 using primer pair 27f/1492r and Sanger sequencing.

| | |
|---|---|
| **PM69 culture 1** | >primer 27-forward (1151 bp)<br>AAGCGGGGGTAACAAAGTAGTTTACTATGGAGTTACCCTAGCGGCGGACGGGTGAGTAATGCGTGAACAATCTACCTCAAAGACTGGGATAACAGCTCGAAAGGGCTGCTAATACCGGATATGCTCA<br>AGGTTCCGCATGGGGCATTGAGGAAAGGGGGGACCCGCTTTGAGATGGGTTCACGTCCCATCAGCTAGTTGGTGAGGTAACGGCTCACCAAGGCGACGACGGGTAGCCGGCCTGAGAGGGTGGTCGG<br>CCACACTGGAACTGAGACACGGTCCAGACTCCTACGGGGGGCAGCAGTAGGGAATCTTCGGCAATGGGCGAAAGCCTGACCGAGCGACGCCGCGTGAGGGAAGAAGGTCTTCGGATTGTAAACCTCT<br>GTCTTGGGGGATGAGAAAGGACAGTACCCCAGGAGGAAGCCCCGGCTAACTACGTGCCAGCAGCCGCGGTAATACGTAGGGGGCGAGCGTTGTCCGGAATTACTGGGCGTAAAGGGCGTGCAGGTG<br>GTCTCTTAAGTTAGGTGGGAAATCCCATAGCTCAACTATGGGGGTGCGCCTAAAACTGGGGGGCTAGAGGGCAAGAGAGGGAAGCGGAATTCCCGGTGTAGCGGTGAAATGCGTAGATATCGGGAG<br>GAACACCAGTGGCGAAGGCGGCTTCCTGGATTGCACCTGACACTGAGGCGCGAAAGCCAGGGGAGCGAACGGGATTAGATACCCCGGTAGTCCTGGCCGTAAACGATGGATGCTAGGTGTGGGAGG<br>TATCGACCCCTTCCGTGCCGCAGCTAACGCATTAAGCATCCCGCCTGGGGAGTACGGCCGCAAGGTTGAAACTCAAAGGAATTGACGGGGGCCCGCACAAGCGGTGGAGCATGTGGTTTAATTCGAC<br>GCAACGCGAAGGACCTTACCAGGGTTTGACATGCTGGTGGTACTGAACCGAAAGGGGAAGGACCCAGGCAACTGGGAGCCAGCACAGGTGGTGCATGGCTGTCGTCAGCTCGTGTCGTGAGATGTTG<br>GGTTAAGTCCCGCAACGAGCGCAACCCCTACATTCAGTTGCTAACGGGTAGAGCCGAGCACTCTGGATGGACTGCCGGGGATGACCCGGAGAAAGTGGGGATGACGTCAGTCATCATGCCCTTATGC<br>CTGGGCCACCA |
| | >primer 1492-reverse (1115 bp)<br>GTTAAGGGCACCGGCTTCGGGTGTGACAAACTTTCGTGGTGTTGACGGGCGGTGTGTACAAGGCCCGGGAACGTATTCACCGCGGTATGCTGACCCGCGATTACTAGCGATTCCGGCTTCACGCAGGC<br>GAGTTGCAGCCTGCGATCCGAACTTAGACCTGCTTTTGGGGATCCGCTCCCCCTCGCGGGTTTGCCTCCCTCTGTACAGGCCATTGTAGCACGTGTGTGGCCCAGGGCATAAAGGGCATGATGACTTGA<br>CGTCATCCCCACCTTCCTCCGGGTCATCCCCGGCAGTCCATCCAGAGTGCTCGGCTCTACCCGTTAGCAACTGAATGTAGGGGTTGCGCTCGTTGCGGGACTTAACCCAACATCTCACGACACGAGCTG<br>ACGACAGCCATGCACCACCTGTGCTGGCTCCCAGTTGCCTGGGTCCTTCCCCTTTCGGTTCAGTACCACCAGCATGTCAAACCCTGGTAAGGTCCTTCGCGTTGCGTCGAATTAAACCACATGCTCCAC<br>CGCTTGTGCGGGCCCCCGTCAATTCCTTTGAGTTTCAACCTTGCGGCCGTACTCCCCAGGCGGGATGCTTAATGCGTTAGCTGCGGCACGGAAGGGGTCGATACCTCCCACACCTAGCATCCATCGTTT<br>ACGGCCAGGACTACCGGGGTATCTAATCCCGTTCGCTCCCCTGGCTTTCGCGCCTCAGTGTCAGGTGCAATCCAGGAAGCCGCCTTCGCCACTGGTGTTCCTCCCGATATCTACGCATTTCACCGCTAC<br>ACCGGGAATTCCGCTTCCCTCTCTTGCCCTCTAGCCCCCCAGTTTTAGGCGCACCCCCATAGTTGAGCTATGGGATTTCCCACCTAACTTAAGAGACCACCTGCACGCCCTTTACGCCCAGTAATTCCGG<br>ACAACGCTCGCCCCCTACGTATTACCGCGGCTGCTGGCACGTAGTTAGCCGGGGCTTCCTCCTGGGGTACTGTCCTTTCTCATCCCCCAAGACAGAGGTTTACAATCCGAAGACCTTCTTCCCTCACGC<br>GGCGTCGCTCGGTCAGGCTTCCGCCCATTGCCGAAGATTCCTACTGCTGCCCCCGTAGAATCTGGACCGTGTCTCAGTTCCGT |
| **PM69 culture 2** | >primer 27-forward (1098 bp)<br>TAACGAAGTAGTTTACTATGGAGTTACCCTAGCGGCGGACGGGTGAGTAATGCGTGAACAATCTACCTCAAAGACTGGGATAACAGCTCGAAAGGGCTGCTAATACCGGATATGCTCAAGGTTCCGC<br>ATGGGGCATTGAGGAAAGGGGGGACCCGCTTTGAGATGGGTTCACGTCCCATCAGCTAGTTGGTGAGGTAACGGCTCACCAAGGCGACGACGGGTAGCCGGCCTGAGAGGGTGGTCGGCCACACTGG<br>AACTGAGACACGGTCCAGACTCCTACGGGGGGCAGCAGTAGGGAATCTTCGGCAATGGGCGAAAGCCTGACCGAGCGACGCCGCGTGAGGGAAGAAGGTCTTCGGATTGTAAACCTCTGTCTTGGGG<br>GATGAGAAAGGACAGTACCCCAGGAGGAAGCCCCGGCTAACTACGTGCCAGCAGCCGCGGTAATACGTAGGGGGCGAGCGTTGTCCGGAATTACTGGGCGTAAAGGGCGTGCAGGTGGTCTCTTAA<br>GTTAGGTGGGAAATCCCATAGCTCAACTATGGGGGTGCGCCTAAAACTGGGGGGCTAGAGGGCAAGAGAGGGAAGCGGAATTCCCGGTGTAGCGGTGAAATGCGTAGATATCGGGAGGAACACCAG<br>TGGCGAAGGCGGCTTCCTGGATTGCACCTGACACTGAGGCGCGAAAGCCAGGGGAGCGAACGGGATTAGATACCCCGGTAGTCCTGGCCGTAAACGATGGATGCTAGGTGTGGGAGGTATCGACCCC<br>TTCCGTGCCGCAGCTAACGCATTAAGCATCCCGCCTGGGGAGTACGGCCGCAAGGTTGAAACTCAAAGGAATTGACGGGGGCCCGCACAAGCGGTGGAGCATGTGGTTTAATTCGACGCAACGCGAA<br>GGACCTTACCAGGGTTTGACATGCTGGTGGTACTGAACCGAAAGGGGAAGGACCCAGGCAACTGGGAGCCAGCACAGGTGGTGCATGGCTGTCGTCAGCTCGTGTCGTGAGATGTTGGGTTAAGTCC<br>CGCAACGAGCGCAACCCCTACATTCAGTTGCTAACGGGTAGAGCGAGCACTCTGGATGGACTGCCGGGATGACCCGAGAAAAGG |
| | >primer 1492-reverse (1112 bp)<br>TTAAGGTCACCGGCTTCGGGTGTGACAAACTTTCGTGGTGTGACGGGCGGTGTGTACAAGGCCCGGGAACGTATTCACCGCGGTATGCTGACCCGCGATTACTAGCGATTCCGGCTTCACGCAGGCGA<br>GTTGCAGCCTGCGATCCGAACTTAGACCTGCTTTTGGGGATCCGCTCCCCCTCGCGGGTTTGCCTCCCTCTGTACAGGCCATTGTAGCACGTGTGTGGCCCAGGGCATAAAGGGCATGATGACTTGACG<br>TCATCCCCACCTTCCTCCGGGTCATCCCCGGCAGTCCATCCAGAGTGCTCGGCTCTACCCGTTAGCAACTGAATGTAGGGGTTGCGCTCGTTGCGGGACTTAACCCAACATCTCACGACACGAGCTGAC<br>GACAGCCATGCACCACCTGTGCTGGCTCCCAGTTGCCTGGGTCCTTCCCCTTTCGGTTCAGTACCACCAGCATGTCAAACCCTGGTAAGGTCCTTCGCGTTGCGTCGAATTAAACCACATGCTCCACCG<br>CTTGTGCGGGCCCCCGTCAATTCCTTTGAGTTTCAACCTTGCGGCCGTACTCCCCAGGCGGGATGCTTAATGCGTTAGCTGCGGCACGGAAGGGGTCGATACCTCCCACACCTAGCATCCATCGTTTAC<br>GGCCAGGACTACCGGGGTATCTAATCCCGTTCGCTCCCCTGGCTTTCGCGCCTCAGTGTCAGGTGCAATCCAGGAAGCCGCCTTCGCCACTGGTGTTCCTCCCGATATCTACGCATTTCACCGCTACAC<br>CGGGAATTCCGCTTCCCTCTCTTGCCCTCTAGCCCCCCAGTTTTAGGCGCACCCCCATAGTTGAGCTATGGGATTTCCCACCTAACTTAAGAGACCACCTGCACGCCCTTTACGCCCAGTAATTCCGGAC<br>AACGCTCGCCCCCTACGTATTACCGCGGCTGCTGGCACGTAGTTAGCCGGGGCTTCCTCCTGGGGTACTGTCCTTTCTCATCCCCCAAGACAGAGGTTTACAATCCGAAGACCTTCTTCCCTCACGCGG<br>CGTCGCTCGGTCAGGCTTCGCCCATTGCCGAAGATTCCTACTGCTGCCCCCGTAGAATCTGGACCGTGTCTCAGTTCCGT |

| | |
|---|---|
| **PM69 culture 3** | >primer 27-forward (1106 bp)<br>GGACGGGTGAGTAATGCGTGAACAATCTACCTCAAAGACTGGGATAACAGCTCGAAAGGGCTGCTAATACCGGATATGCTCAAGGTTCCGCATGGGGCATTGAGGAAAGGGGGGACCCGCTTTGAGA<br>TGGGTTCACGTCCCATCAGCTAGTTGGTGAGGTAACGGCTCACCAAGGCGACGACGGGTAGCCGGCCTGAGAGGGTGGTCGGCCACACTGGAACTGAGACACGGTCCAGACTCCTACGGGGGGCAGC<br>AGTAGGGAATCTTCGGCAATGGGCGAAAGCCTGACCGAGCGACGCCGCGTGAGGGAAGAAGGTCTTCGGATTGTAAACCTCTGTCTTGGGGGATGAGAAAGGACAGTACCCCAGGAGGAAGCCCCG<br>GCTAACTACGTGCCAGCAGCCGCGGTAATACGTAGGGGGCGAGCGTTGTCCGGAATTACTGGGCGTAAAGGGCGTGCAGGTGGTCTCTTAAGTTAGGTGGGAAATCCCATAGCTCAACTATGGGGGT<br>GCGCCTAAAACTGGGGGGCTAGAGGGCAAGAGAGGGAAGCGGAATTCCCGGTGTAGCGGTGAAATGCGTAGATATCGGGAGGAACACCAGTGGCGAAGGCGGCTTCCTGGATTGCACCTGACACTG<br>AGGCGCGAAAGCCAGGGGAGCGAACGGGATTAGATACCCCGGTAGTCCTGGCCGTAAACGATGGATGCTAGGTGTGGGAGGTATCGACCCCTTCCGTGCCGCAGCTAACGCATTAAGCATCCCGCCT<br>GGGGAGTACGGCCGCAAGGTTGAAACTCAAAGGAATTGACGGGGGCCCGCACAAGCGGTGGAGCATGTGGTTTAATTCGACGCAACGCGAAGGACCTTACCAGGGTTTGACATGCTGGTGGTACTGA<br>ACCGAAAGGGGAAGGACCCAGGCAACTGGGAGCCAGCACAGGTGGTGCATGGCTGTCGTCAGCTCGTGTCGTGAGATGTTGGGTTAAGTCCCGCAACGAGCGCAACCCCTACATTCAGTTGCTAACG<br>GGTAGAGCCGAGCACTCTGGATGGACTGCCGGGATGACCCGGAGAAAGGTGGGGATGACGTCAAGTCATCATGCCCTTTATGCCCGGGGCCA |
| | >primer 1492-reverse (1074 bp)<br>GGACGGGCGGTGTGTACAAGGCCCGGGAACGTATTCACCGCGGTATGCTGACCCGCGATTACTAGCGATTCCGGCTTCACGCAGGCGAGTTGCAGCCTGCGATCCGAACTTAGACCTGCTTTTGGGGA<br>TCCGCTCCCCCTCGCGGGTTTGCCTCCCTCTGTACAGGCCATTGTAGCACGTGTGTGGCCCAGGGCATAAAGGGCATGATGACTTGACGTCATCCCCACCTTCCTCCGGGTCATCCCCGGCAGTCCATC<br>CAGAGTGCTCGGCTCTACCCGTTAGCAACTGAATGTAGGGGTTGCGCTCGTTGCGGGACTTAACCCAACATCTCACGACACGAGCTGACGACAGCCATGCACCACCTGTGCTGGCTCCCAGTTGCCTG<br>GGTCCTTCCCCTTTCGGTTCAGTACCACCAGCATGTCAAACCCTGGTAAGGTCCTTCGCGTTGCGTCGAATTAAACCACATGCTCCACCGCTTGTGCGGGCCCCCGTCAATTCCTTTGAGTTTCAACCTT<br>GCGGCCGTACTCCCCAGGCGGGATGCTTAATGCGTTAGCTGCGGCACGGAAGGGGTCGATACCTCCCACACCTAGCATCCATCGTTTACGGCCAGGACTACCGGGGTATCTAATCCCGTTCGCTCCCCT<br>GGCTTTCGCGCCTCAGTGTCAGGTGCAATCCAGGAAGCCGCCTTCGCCACTGGTGTTCCTCCCGATATCTACGCATTTCACCGCTACACCGGGAATTCCGCTTCCCTCTCTTGCCCTCTAGCCCCCCAGT<br>TTTAGGCGCACCCCCATAGTTGAGCTATGGGATTTCCCACCTAACTTAAGAGACCACCTGCACGCCCTTTACGCCCAGTAATTCCGGACAACGCTCGCCCCCTACGTATTACCGCGGCTGCTGGCACGT<br>AGTTAGCCGGGGCTTCCTCCTGGGGTACTGTCCTTTCTCATCCCCCAAGACAGAGGTTTACAATCCGAAGACCTTCTTCCCTCACGCGGCGTCGCTCGGTCAGGCTTCCGCCCATTGCCGAAGATTCCTA<br>CTGCTGCCCCCGTAGGAATCTGGACCGTGTCTCAGTTCCGT |

**Table S9.** *GUNC* results for the PM69 assembly using the databases *proGenome* and *GTDB* with or without contigs <1kb

| database | genome | n_genes_called | n_genes_mapped | n_contigs | taxonomic_level | proportion_genes_retained_in_major_clades | genes_retained_index | clade_separation_score | contamination_portion | n_effective_surplus_clades | mean_hit_identity | reference_re-presentation_score | **pass.GUNC** |
|---|---|---|---|---|---|---|---|---|---|---|---|---|---|
| proGenomes | assembly | 3260 | 2849 | 45 | kingdom | 1 | 0.87 | 0.02 | 0.02 | 0.05 | 0.53 | 0.47 | **True** |
| proGenomes | assembly≥1kb | 3210 | 2822 | 18 | kingdom | 1 | 0.88 | 0.03 | 0.02 | 0.05 | 0.53 | 0.47 | **True** |
| GTDB | assembly | 3260 | 2899 | 49 | family | 0.67 | 0.6 | 0.04 | 0.55 | 3.79 | 0.59 | 0.35 | **True** |
| GTDB | assembly≥1kb | 3210 | 2868 | 18 | order | 0.77 | 0.69 | 0.03 | 0.61 | 5.02 | 0.58 | 0.4 | **True** |

### KEGG Mapper Reconstruction Result

| Pathway (194) | Brite (34) | Brite Table (6) | Module (34) |
|---|---|---|---|

View ◉ complete only ○ including 1 block missing ○ including any incomplete [Exec]

Show matched objects

**Pathway modules**

**Carbohydrate metabolism**
Central carbohydrate metabolism
M00001 Glycolysis (Embden-Meyerhof pathway), glucose => pyruvate (11) (complete 9/9)
M00002 Glycolysis, core module involving three-carbon compounds (6) (complete 5/5)
M00307 Pyruvate oxidation, pyruvate => acetyl-CoA (7) (complete 1/1)
M00005 PRPP biosynthesis, ribose-5P => PRPP (1) (complete 1/1)
Other carbohydrate metabolism
M00632 Galactose degradation, Leloir pathway, galactose => alpha-D-glucose-1P (4) (complete 4/4)
**Energy metabolism**
ATP synthesis
M00157 F-type ATPase, prokaryotes and chloroplasts (8) (complete 1/1)
M00159 V/A-type ATPase, prokaryotes (9) (complete 1/1)
**Lipid metabolism**
Fatty acid metabolism
M00082 Fatty acid biosynthesis, initiation (6) (complete 2/2)
**Nucleotide metabolism**
Purine metabolism
M00048 De novo purine biosynthesis, PRPP + glutamine => IMP (11) (complete 8/8)
M00049 Adenine ribonucleotide biosynthesis, IMP => ADP,ATP (4) (complete 4/4)
M00050 Guanine ribonucleotide biosynthesis, IMP => GDP,GTP (4) (complete 4/4)
Pyrimidine metabolism
M00052 Pyrimidine ribonucleotide biosynthesis, UMP => UDP/UTP,CDP/CTP (3) (complete 3/3)
**Amino acid metabolism**
Serine and threonine metabolism
M00018 Threonine biosynthesis, aspartate => homoserine => threonine (5) (complete 5/5)
M00621 Glycine cleavage system (4) (complete 3/3)
Cysteine and methionine metabolism
M00021 Cysteine biosynthesis, serine => cysteine (2) (complete 2/2)
Branched-chain amino acid metabolism
M00019 Valine/isoleucine biosynthesis, pyruvate => valine / 2-oxobutanoate => isoleucine (5) (complete 4/4)
M00432 Leucine biosynthesis, 2-oxoisovalerate => 2-oxoisocaproate (4) (complete 3/3)
Lysine metabolism
M00526 Lysine biosynthesis, DAP dehydrogenase pathway, aspartate => lysine (6) (complete 6/6)
M00527 Lysine biosynthesis, DAP aminotransferase pathway, aspartate => lysine (7) (complete 7/7)
Arginine and proline metabolism
M00028 Ornithine biosynthesis, glutamate => ornithine (5) (complete 4/4)
M00844 Arginine biosynthesis, ornithine => arginine (3) (complete 3/3)
M00015 Proline biosynthesis, glutamate => proline (3) (complete 2/2)
Polyamine biosynthesis
M00133 Polyamine biosynthesis, arginine => agmatine => putrescine => spermidine (5) (complete 4/4)
**Glycan metabolism**
Nucleotide sugar biosynthesis
M00909 UDP-GlcNAc biosynthesis, prokaryotes, Fru-6P => UDP-GlcNAc (4) (complete 3/3)
M00995 UDP-MurNAc biosynthesis, Fru-6P => UDP-MurNAc (5) (complete 5/5)
M00549 UDP-Glc biosynthesis, Glc => UDP-Glc (3) (complete 3/3)
M00554 UDP-Gal biosynthesis, Gal => UDP-Gal (2) (complete 2/2)
M01000 GDP-Man biosynthesis, Fru-6P => GDP-Man (6) (complete 3/3)
M01014 UDP-ManNAcA biosynthesis, UDP-GlcNAc => UDP-ManNAcA (2) (complete 2/2)
M00793 dTDP-L-Rha biosynthesis, Glc-1P => dTDP-L-Rha (4) (complete 3/3)
**Metabolism of cofactors and vitamins**
Cofactor and vitamin metabolism
M00916 Pyridoxal phosphate biosynthesis, R5P + glyceraldehyde-3P + glutamine => pyridoxal-P (2) (complete 1/1)
M00115 NAD biosynthesis, aspartate => quinolinate => NAD (6) (complete 5/5)
M00120 Coenzyme A biosynthesis, pantothenate => CoA (4) (complete 3/3)
M00140 C1-unit interconversion, prokaryotes (3) (complete 3/3)

**Figure S1.** *KEGG Mapper Reconstruct* (https://www.genome.jp/kegg/mapper/reconstruct.html) results: Overview of all complete modules for genome PM69

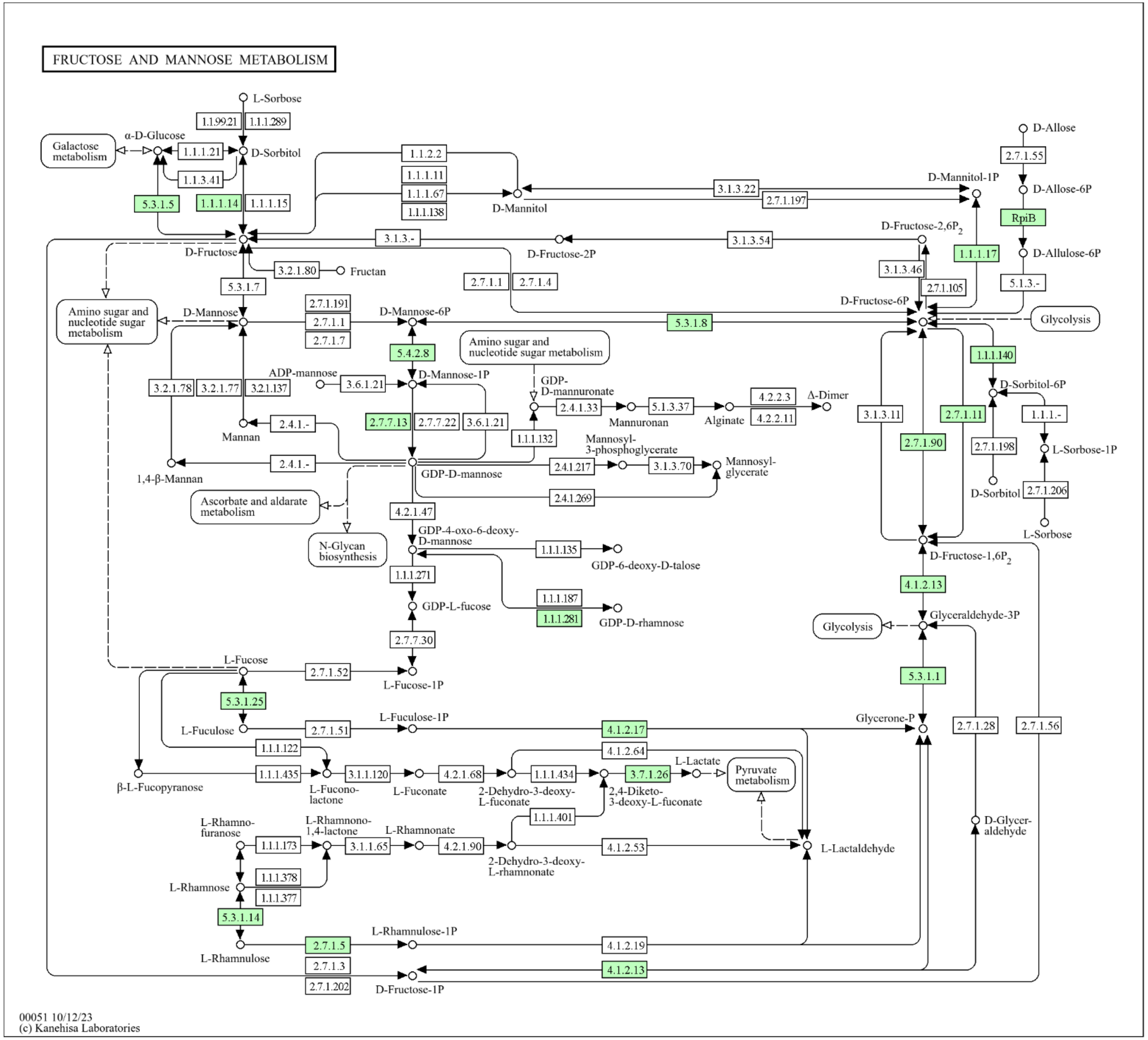


**Figure S2.** *KEGG Mapper Reconstruct* results for Fructose and Mannose Metabolism (map00051). Green EC boxes show that enzyme is present in genome of PM69.

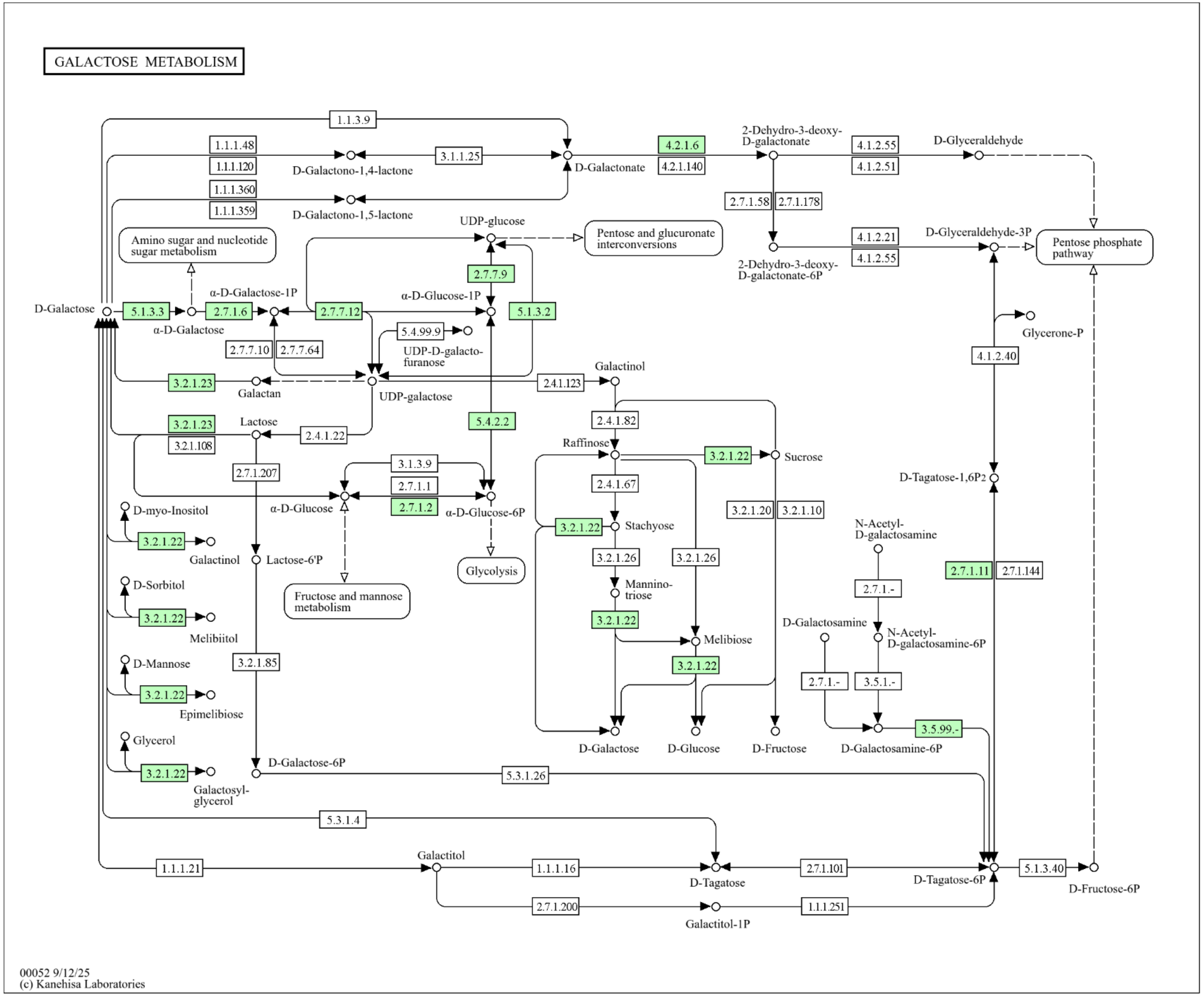


**Figure S3.** *KEGG Mapper Reconstruct* results for Galactose Metabolism (map00052). Green EC boxes show that enzyme is present in genome of PM69.

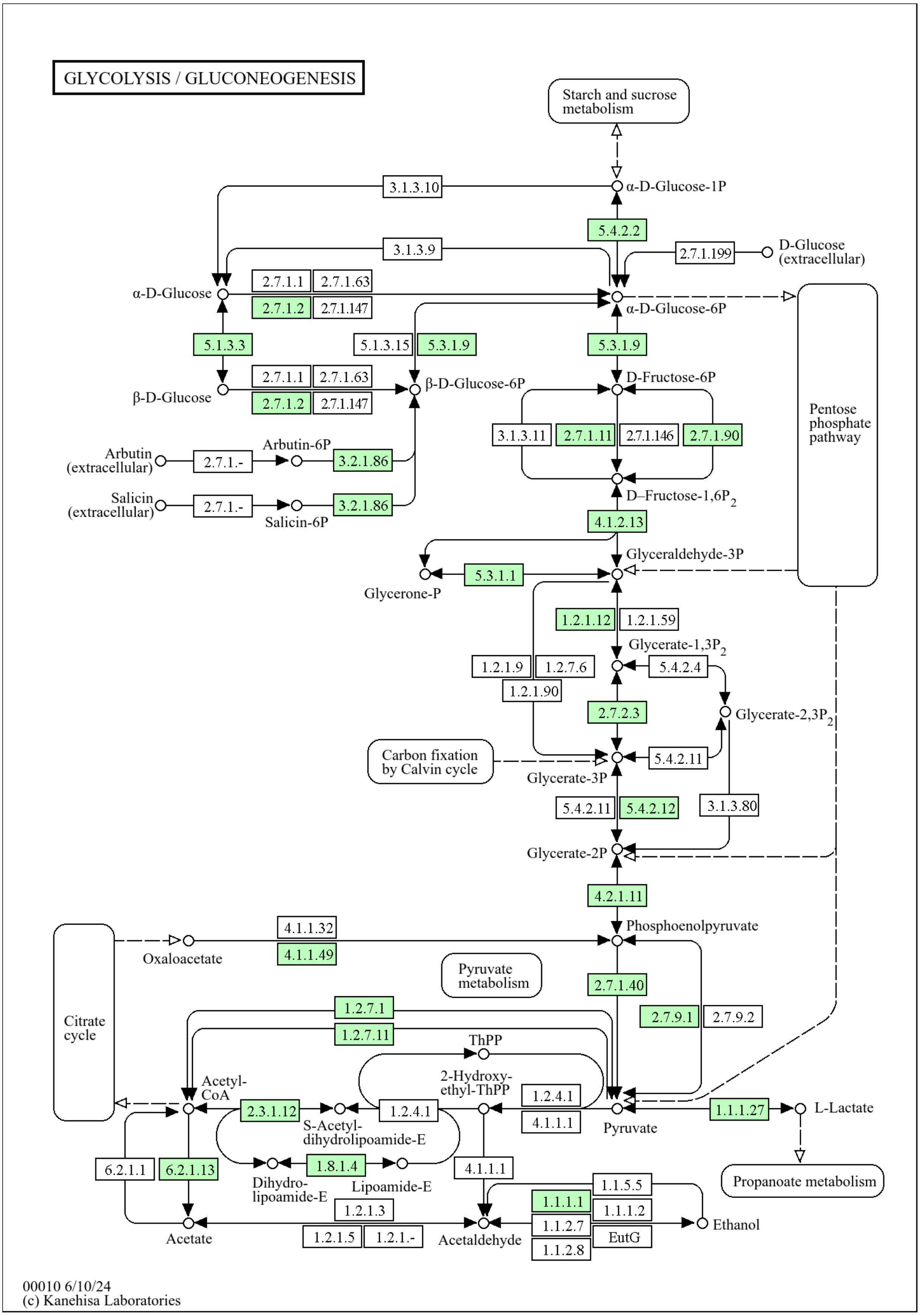


**Figure S4.** *KEGG Mapper Reconstruct* results for Glycolysis (map00010). Green EC boxes show that enzyme is present in genome of PM69.
