## Supplementary Results 2 for "*Thermoaminiphila catenidiffluenda* gen. nov., sp. nov.: A novel thermophilic, strictly anaerobic bacterium representing *Thermoaminiphilia* class nov., a newly described class thriving in hydrocarbon-rich habitats and biogas fermenters"

| Query | Reference | ANI | AF Query | AF Reference |
|---|---|---|---|---|
| GB_GCA_012512055_1.fna | GB_GCA_012512055_1.fna | 100.00 | 617 | 627 |
| GB_GCA_012512055_1.fna | PM003_bin.28.strict.fa | 97.73 | 294 | 627 |
| GB_GCA_012512055_1.fna | PM004_bin.45.permissive.fa | 88.20 | 359 | 627 |
| GB_GCA_012512055_1.fna | s__UBA4971_sp002398515.fna | 87.77 | 465 | 627 |
| GB_GCA_012512055_1.fna | PM006_bin.39.orig.fa | 81.95 | 459 | 627 |
| GB_GCA_012512055_1.fna | s__UBA4971_sp012518595.fna | 81.79 | 364 | 627 |
| GB_GCA_012512055_1.fna | PM003_bin.81.permissive.fa | 81.73 | 318 | 627 |
| GB_GCA_012512055_1.fna | s__UBA4971_sp002424305.fna | 79.33 | 319 | 627 |
| GB_GCA_012512055_1.fna | s__UBA4971_sp012841305.fna | 79.31 | 320 | 627 |
| GB_GCA_012512055_1.fna | PM008_bin.17.strict.fa | 78.99 | 275 | 627 |
| GB_GCA_012512055_1.fna | s__UBA4971_sp900019985.fna | 78.94 | 225 | 627 |
| GB_GCA_012512055_1.fna | s__UBA4971_sp012689375.fna | 76.53 | 65 | 627 |
| PM003_bin.28.strict.fa | PM003_bin.28.strict.fa | 100.00 | 352 | 359 |
| PM003_bin.28.strict.fa | GB_GCA_012512055_1.fna | 98.41 | 301 | 359 |
| PM003_bin.28.strict.fa | s__UBA4971_sp002398515.fna | 87.21 | 289 | 359 |
| PM003_bin.28.strict.fa | PM004_bin.45.permissive.fa | 86.63 | 231 | 359 |
| PM003_bin.28.strict.fa | PM006_bin.39.orig.fa | 82.31 | 288 | 359 |
| PM003_bin.28.strict.fa | s__UBA4971_sp012518595.fna | 81.10 | 225 | 359 |
| PM003_bin.28.strict.fa | PM003_bin.81.permissive.fa | 80.87 | 203 | 359 |
| PM003_bin.28.strict.fa | s__UBA4971_sp012841305.fna | 78.96 | 178 | 359 |
| PM003_bin.28.strict.fa | s__UBA4971_sp002424305.fna | 78.96 | 178 | 359 |
| PM003_bin.28.strict.fa | PM008_bin.17.strict.fa | 78.58 | 161 | 359 |
| PM003_bin.28.strict.fa | s__UBA4971_sp900019985.fna | 78.54 | 137 | 359 |
| PM003_bin.81.permissive.fa | PM003_bin.81.permissive.fa | 100.00 | 535 | 542 |
| PM003_bin.81.permissive.fa | PM006_bin.39.orig.fa | 98.22 | 513 | 542 |
| PM003_bin.81.permissive.fa | s__UBA4971_sp012518595.fna | 83.78 | 340 | 542 |
| PM003_bin.81.permissive.fa | s__UBA4971_sp002398515.fna | 83.52 | 384 | 542 |
| PM003_bin.81.permissive.fa | PM004_bin.45.permissive.fa | 82.40 | 263 | 542 |
| PM003_bin.81.permissive.fa | GB_GCA_012512055_1.fna | 81.78 | 323 | 542 |
| PM003_bin.81.permissive.fa | PM003_bin.28.strict.fa | 80.92 | 203 | 542 |

| | | | | |
|---|---|---|---|---|
| PM003_bin.81.permissive.fa | s__UBA4971_sp002424305.fna | 80.07 | 356 | 542 |
| PM003_bin.81.permissive.fa | s__UBA4971_sp012841305.fna | 80.07 | 356 | 542 |
| PM003_bin.81.permissive.fa | s__UBA4971_sp900019985.fna | 78.08 | 133 | 542 |
| PM003_bin.81.permissive.fa | PM008_bin.17.strict.fa | 77.78 | 166 | 542 |
| PM003_bin.81.permissive.fa | s__UBA4971_sp012689375.fna | 76.77 | 67 | 542 |
| PM004_bin.15.strict.fa | PM004_bin.15.strict.fa | 100.00 | 60 | 61 |
| PM004_bin.45.permissive.fa | PM004_bin.45.permissive.fa | 100.00 | 719 | 733 |
| PM004_bin.45.permissive.fa | s__UBA4971_sp002398515.fna | 90.38 | 394 | 733 |
| PM004_bin.45.permissive.fa | GB_GCA_012512055_1.fna | 88.41 | 355 | 733 |
| PM004_bin.45.permissive.fa | PM003_bin.28.strict.fa | 86.21 | 235 | 733 |
| PM004_bin.45.permissive.fa | PM006_bin.39.orig.fa | 82.74 | 420 | 733 |
| PM004_bin.45.permissive.fa | s__UBA4971_sp012518595.fna | 82.65 | 324 | 733 |
| PM004_bin.45.permissive.fa | PM003_bin.81.permissive.fa | 82.35 | 264 | 733 |
| PM004_bin.45.permissive.fa | s__UBA4971_sp012841305.fna | 80.06 | 301 | 733 |
| PM004_bin.45.permissive.fa | s__UBA4971_sp002424305.fna | 80.06 | 301 | 733 |
| PM004_bin.45.permissive.fa | s__UBA4971_sp900019985.fna | 79.82 | 210 | 733 |
| PM004_bin.45.permissive.fa | PM008_bin.17.strict.fa | 79.40 | 273 | 733 |
| PM004_bin.45.permissive.fa | s__UBA4971_sp012689375.fna | 76.73 | 62 | 733 |
| PM006_bin.39.orig.fa | PM006_bin.39.orig.fa | 100.00 | 923 | 924 |
| PM006_bin.39.orig.fa | PM003_bin.81.permissive.fa | 97.90 | 516 | 924 |
| PM006_bin.39.orig.fa | s__UBA4971_sp012518595.fna | 83.93 | 510 | 924 |
| PM006_bin.39.orig.fa | s__UBA4971_sp002398515.fna | 83.56 | 572 | 924 |
| PM006_bin.39.orig.fa | PM004_bin.45.permissive.fa | 82.67 | 421 | 924 |
| PM006_bin.39.orig.fa | PM003_bin.28.strict.fa | 82.08 | 286 | 924 |
| PM006_bin.39.orig.fa | GB_GCA_012512055_1.fna | 81.99 | 448 | 924 |
| PM006_bin.39.orig.fa | s__UBA4971_sp012841305.fna | 80.55 | 537 | 924 |
| PM006_bin.39.orig.fa | s__UBA4971_sp002424305.fna | 80.55 | 537 | 924 |
| PM006_bin.39.orig.fa | s__UBA4971_sp900019985.fna | 77.96 | 208 | 924 |
| PM006_bin.39.orig.fa | PM008_bin.17.strict.fa | 77.71 | 258 | 924 |
| PM006_bin.39.orig.fa | s__UBA4971_sp012689375.fna | 76.80 | 100 | 924 |
| PM007_bin.27.permissive.fa | PM007_bin.27.permissive.fa | 100.00 | 231 | 236 |

| | | | | |
|---|---|---|---|---|
| PM007_bin.27.permissive.fa | s__DUSO01_sp012842565.fna | 81.55 | 148 | 236 |
| PM008_bin.17.strict.fa | PM008_bin.17.strict.fa | 100.00 | 604 | 606 |
| PM008_bin.17.strict.fa | s__UBA4971_sp900019985.fna | 99.32 | 410 | 606 |
| PM008_bin.17.strict.fa | PM004_bin.45.permissive.fa | 79.60 | 266 | 606 |
| PM008_bin.17.strict.fa | s__UBA4971_sp002398515.fna | 79.30 | 325 | 606 |
| PM008_bin.17.strict.fa | GB_GCA_012512055_1.fna | 78.93 | 277 | 606 |
| PM008_bin.17.strict.fa | PM003_bin.28.strict.fa | 78.50 | 171 | 606 |
| PM008_bin.17.strict.fa | PM006_bin.39.orig.fa | 77.81 | 252 | 606 |
| PM008_bin.17.strict.fa | s__UBA4971_sp012518595.fna | 77.72 | 222 | 606 |
| PM008_bin.17.strict.fa | PM003_bin.81.permissive.fa | 77.46 | 179 | 606 |
| PM008_bin.17.strict.fa | s__UBA4971_sp012841305.fna | 77.40 | 149 | 606 |
| PM008_bin.17.strict.fa | s__UBA4971_sp002424305.fna | 77.40 | 149 | 606 |
| PM008_bin.17.strict.fa | s__UBA4971_sp012689375.fna | 76.78 | 57 | 606 |
| PM008_bin.51.permissive.fa | PM008_bin.51.permissive.fa | 100.00 | 307 | 313 |
| PM008_bin.53.orig.fa | PM008_bin.53.orig.fa | 100.00 | 384 | 391 |
| PM008_bin.53.orig.fa | s__UBA6256_sp012516125.fna | 98.54 | 377 | 391 |
| PM008_bin.53.orig.fa | PM012_bin.13.permissive.fa | 95.49 | 322 | 391 |
| PM008_bin.53.orig.fa | s__UBA6256_sp014360195.fna | 94.70 | 349 | 391 |
| PM008_bin.53.orig.fa | s__UBA6256_sp002452295.fna | 83.10 | 303 | 391 |
| PM008_bin.53.orig.fa | s__UBA6256_sp012842395.fna | 80.69 | 278 | 391 |
| PM008_bin.53.orig.fa | PM009_bin.22.orig.fa | 80.62 | 261 | 391 |
| PM008_bin.53.orig.fa | s__UBA6256_sp024653485.fna | 80.18 | 267 | 391 |
| PM008_bin.65.orig.fa | PM008_bin.65.orig.fa | 100.00 | 679 | 682 |
| PM009_bin.22.orig.fa | PM009_bin.22.orig.fa | 100.00 | 842 | 848 |
| PM009_bin.22.orig.fa | s__UBA6256_sp012842395.fna | 99.23 | 785 | 848 |
| PM009_bin.22.orig.fa | PM008_bin.53.orig.fa | 80.71 | 251 | 848 |
| PM009_bin.22.orig.fa | PM012_bin.13.permissive.fa | 80.61 | 483 | 848 |
| PM009_bin.22.orig.fa | s__UBA6256_sp012516125.fna | 80.61 | 492 | 848 |
| PM009_bin.22.orig.fa | s__UBA6256_sp014360195.fna | 80.26 | 521 | 848 |
| PM009_bin.22.orig.fa | s__UBA6256_sp024653485.fna | 78.65 | 367 | 848 |
| PM009_bin.22.orig.fa | s__UBA6256_sp002452295.fna | 78.52 | 366 | 848 |

| | | | | |
|---|---|---|---|---|
| PM012_bin.13.permissive.fa | PM012_bin.13.permissive.fa | 100.00 | 1157 | 1162 |
| PM012_bin.13.permissive.fa | PM008_bin.53.orig.fa | 95.21 | 314 | 1162 |
| PM012_bin.13.permissive.fa | s__UBA6256_sp012516125.fna | 94.49 | 743 | 1162 |
| PM012_bin.13.permissive.fa | s__UBA6256_sp014360195.fna | 94.03 | 766 | 1162 |
| PM012_bin.13.permissive.fa | s__UBA6256_sp002452295.fna | 83.22 | 536 | 1162 |
| PM012_bin.13.permissive.fa | s__UBA6256_sp012842395.fna | 81.01 | 555 | 1162 |
| PM012_bin.13.permissive.fa | PM009_bin.22.orig.fa | 80.73 | 465 | 1162 |
| PM012_bin.13.permissive.fa | s__UBA6256_sp024653485.fna | 80.34 | 496 | 1162 |
| s__Ch115_sp013178415.fna | s__Ch115_sp013178415.fna | 100.00 | 1163 | 1166 |
| s__Ch115_sp013178415.fna | strain.pm69.vs2.0.gt1kb.fa | 98.21 | 942 | 1166 |
| s__Ch115_sp013178415.fna | s__JABLXL01_sp013178125.fna | 81.28 | 78 | 1166 |
| s__Ch115_sp013178415.fna | s__JABLXL01_sp012839815.fna | 80.27 | 63 | 1166 |
| s__Ch115_sp013178415.fna | s__DUSO01_sp012842565.fna | 78.51 | 67 | 1166 |
| s__DUSO01_sp012842565.fna | s__DUSO01_sp012842565.fna | 100.00 | 738 | 745 |
| s__DUSO01_sp012842565.fna | PM007_bin.27.permissive.fa | 81.47 | 142 | 745 |
| s__DUSO01_sp012842565.fna | s__Ch115_sp013178415.fna | 78.15 | 69 | 745 |
| s__DUSO01_sp012842565.fna | s__JABLXL01_sp012839815.fna | 78.06 | 53 | 745 |
| s__DUSO01_sp012842565.fna | strain.pm69.vs2.0.gt1kb.fa | 76.94 | 63 | 745 |
| s__JABLXL01_sp012839815.fna | s__JABLXL01_sp012839815.fna | 100.00 | 850 | 854 |
| s__JABLXL01_sp012839815.fna | s__JABLXL01_sp013178125.fna | 83.53 | 613 | 854 |
| s__JABLXL01_sp012839815.fna | strain.pm69.vs2.0.gt1kb.fa | 79.62 | 54 | 854 |
| s__JABLXL01_sp012839815.fna | s__Ch115_sp013178415.fna | 79.34 | 73 | 854 |
| s__JABLXL01_sp013178125.fna | s__JABLXL01_sp013178125.fna | 100.00 | 1133 | 1137 |
| s__JABLXL01_sp013178125.fna | s__JABLXL01_sp012839815.fna | 83.68 | 598 | 1137 |
| s__JABLXL01_sp013178125.fna | strain.pm69.vs2.0.gt1kb.fa | 83.39 | 80 | 1137 |
| s__JABLXL01_sp013178125.fna | s__Ch115_sp013178415.fna | 80.90 | 82 | 1137 |
| s__JABLXL01_sp013178125.fna | s__DUSO01_sp012842565.fna | 79.06 | 50 | 1137 |
| s__UBA4971_sp002398515.fna | s__UBA4971_sp002398515.fna | 100.00 | 725 | 736 |
| s__UBA4971_sp002398515.fna | PM004_bin.45.permissive.fa | 89.93 | 406 | 736 |
| s__UBA4971_sp002398515.fna | GB_GCA_012512055_1.fna | 87.87 | 445 | 736 |
| s__UBA4971_sp002398515.fna | PM003_bin.28.strict.fa | 87.11 | 282 | 736 |

| | | | | |
|---|---|---|---|---|
| s__UBA4971_sp002398515.fna | PM006_bin.39.orig.fa | 83.54 | 572 | 736 |
| s__UBA4971_sp002398515.fna | PM003_bin.81.permissive.fa | 83.36 | 384 | 736 |
| s__UBA4971_sp002398515.fna | s__UBA4971_sp012518595.fna | 83.33 | 453 | 736 |
| s__UBA4971_sp002398515.fna | s__UBA4971_sp012841305.fna | 80.36 | 445 | 736 |
| s__UBA4971_sp002398515.fna | s__UBA4971_sp002424305.fna | 80.35 | 446 | 736 |
| s__UBA4971_sp002398515.fna | s__UBA4971_sp900019985.fna | 79.40 | 257 | 736 |
| s__UBA4971_sp002398515.fna | PM008_bin.17.strict.fa | 79.25 | 316 | 736 |
| s__UBA4971_sp002398515.fna | s__UBA4971_sp012689375.fna | 77.29 | 85 | 736 |
| s__UBA4971_sp002424305.fna | s__UBA4971_sp012841305.fna | 100.00 | 757 | 765 |
| s__UBA4971_sp002424305.fna | s__UBA4971_sp002424305.fna | 100.00 | 757 | 765 |
| s__UBA4971_sp002424305.fna | s__UBA4971_sp002398515.fna | 80.62 | 423 | 765 |
| s__UBA4971_sp002424305.fna | s__UBA4971_sp012518595.fna | 80.42 | 415 | 765 |
| s__UBA4971_sp002424305.fna | PM006_bin.39.orig.fa | 80.37 | 538 | 765 |
| s__UBA4971_sp002424305.fna | PM003_bin.81.permissive.fa | 80.07 | 341 | 765 |
| s__UBA4971_sp002424305.fna | PM004_bin.45.permissive.fa | 79.95 | 317 | 765 |
| s__UBA4971_sp002424305.fna | GB_GCA_012512055_1.fna | 79.14 | 322 | 765 |
| s__UBA4971_sp002424305.fna | PM003_bin.28.strict.fa | 78.78 | 197 | 765 |
| s__UBA4971_sp002424305.fna | PM008_bin.17.strict.fa | 77.43 | 161 | 765 |
| s__UBA4971_sp002424305.fna | s__UBA4971_sp900019985.fna | 77.42 | 129 | 765 |
| s__UBA4971_sp002424305.fna | s__UBA4971_sp012689375.fna | 76.62 | 84 | 765 |
| s__UBA4971_sp012518595.fna | s__UBA4971_sp012518595.fna | 100.00 | 634 | 643 |
| s__UBA4971_sp012518595.fna | PM003_bin.81.permissive.fa | 83.99 | 339 | 643 |
| s__UBA4971_sp012518595.fna | PM006_bin.39.orig.fa | 83.99 | 517 | 643 |
| s__UBA4971_sp012518595.fna | s__UBA4971_sp002398515.fna | 83.61 | 459 | 643 |
| s__UBA4971_sp012518595.fna | PM004_bin.45.permissive.fa | 82.33 | 334 | 643 |
| s__UBA4971_sp012518595.fna | GB_GCA_012512055_1.fna | 81.47 | 375 | 643 |
| s__UBA4971_sp012518595.fna | PM003_bin.28.strict.fa | 81.04 | 238 | 643 |
| s__UBA4971_sp012518595.fna | s__UBA4971_sp012841305.fna | 80.38 | 414 | 643 |
| s__UBA4971_sp012518595.fna | s__UBA4971_sp002424305.fna | 80.38 | 414 | 643 |
| s__UBA4971_sp012518595.fna | s__UBA4971_sp900019985.fna | 77.78 | 185 | 643 |
| s__UBA4971_sp012518595.fna | PM008_bin.17.strict.fa | 77.67 | 221 | 643 |

| | | | | |
|---|---|---|---|---|
| s__UBA4971_sp012518595.fna | s__UBA4971_sp012689375.fna | 76.72 | 102 | 643 |
| s__UBA4971_sp012689375.fna | s__UBA4971_sp012689375.fna | 100.00 | 533 | 535 |
| s__UBA4971_sp012689375.fna | PM006_bin.39.orig.fa | 77.13 | 85 | 535 |
| s__UBA4971_sp012689375.fna | s__UBA4971_sp002398515.fna | 77.11 | 92 | 535 |
| s__UBA4971_sp012689375.fna | PM004_bin.45.permissive.fa | 76.99 | 56 | 535 |
| s__UBA4971_sp012689375.fna | PM003_bin.81.permissive.fa | 76.98 | 58 | 535 |
| s__UBA4971_sp012689375.fna | s__UBA4971_sp002424305.fna | 76.97 | 78 | 535 |
| s__UBA4971_sp012689375.fna | s__UBA4971_sp012841305.fna | 76.97 | 78 | 535 |
| s__UBA4971_sp012689375.fna | GB_GCA_012512055_1.fna | 76.94 | 69 | 535 |
| s__UBA4971_sp012689375.fna | PM008_bin.17.strict.fa | 76.94 | 50 | 535 |
| s__UBA4971_sp012689375.fna | s__UBA4971_sp012518595.fna | 76.84 | 94 | 535 |
| s__UBA4971_sp012841305.fna | s__UBA4971_sp012841305.fna | 100.00 | 757 | 765 |
| s__UBA4971_sp012841305.fna | s__UBA4971_sp002424305.fna | 100.00 | 757 | 765 |
| s__UBA4971_sp012841305.fna | s__UBA4971_sp002398515.fna | 80.62 | 423 | 765 |
| s__UBA4971_sp012841305.fna | s__UBA4971_sp012518595.fna | 80.42 | 415 | 765 |
| s__UBA4971_sp012841305.fna | PM006_bin.39.orig.fa | 80.37 | 538 | 765 |
| s__UBA4971_sp012841305.fna | PM003_bin.81.permissive.fa | 80.07 | 341 | 765 |
| s__UBA4971_sp012841305.fna | PM004_bin.45.permissive.fa | 79.95 | 317 | 765 |
| s__UBA4971_sp012841305.fna | GB_GCA_012512055_1.fna | 79.14 | 322 | 765 |
| s__UBA4971_sp012841305.fna | PM003_bin.28.strict.fa | 78.78 | 197 | 765 |
| s__UBA4971_sp012841305.fna | PM008_bin.17.strict.fa | 77.43 | 161 | 765 |
| s__UBA4971_sp012841305.fna | s__UBA4971_sp900019985.fna | 77.42 | 129 | 765 |
| s__UBA4971_sp012841305.fna | s__UBA4971_sp012689375.fna | 76.62 | 84 | 765 |
| s__UBA4971_sp900019985.fna | s__UBA4971_sp900019985.fna | 100.00 | 450 | 454 |
| s__UBA4971_sp900019985.fna | PM008_bin.17.strict.fa | 99.62 | 417 | 454 |
| s__UBA4971_sp900019985.fna | PM004_bin.45.permissive.fa | 79.80 | 207 | 454 |
| s__UBA4971_sp900019985.fna | s__UBA4971_sp002398515.fna | 79.61 | 248 | 454 |
| s__UBA4971_sp900019985.fna | GB_GCA_012512055_1.fna | 78.96 | 220 | 454 |
| s__UBA4971_sp900019985.fna | PM003_bin.28.strict.fa | 78.47 | 135 | 454 |
| s__UBA4971_sp900019985.fna | PM006_bin.39.orig.fa | 78.05 | 199 | 454 |
| s__UBA4971_sp900019985.fna | PM003_bin.81.permissive.fa | 77.94 | 129 | 454 |

| | | | | |
|---|---|---|---|---|
| s__UBA4971_sp900019985.fna | s__UBA4971_sp012518595.fna | 77.83 | 172 | 454 |
| s__UBA4971_sp900019985.fna | s__UBA4971_sp012841305.fna | 77.72 | 115 | 454 |
| s__UBA4971_sp900019985.fna | s__UBA4971_sp002424305.fna | 77.72 | 115 | 454 |
| s__UBA6256_sp002452295.fna | s__UBA6256_sp002452295.fna | 100.00 | 854 | 865 |
| s__UBA6256_sp002452295.fna | PM008_bin.53.orig.fa | 83.22 | 290 | 865 |
| s__UBA6256_sp002452295.fna | PM012_bin.13.permissive.fa | 83.00 | 543 | 865 |
| s__UBA6256_sp002452295.fna | s__UBA6256_sp014360195.fna | 82.79 | 605 | 865 |
| s__UBA6256_sp002452295.fna | s__UBA6256_sp012516125.fna | 82.77 | 607 | 865 |
| s__UBA6256_sp002452295.fna | s__UBA6256_sp012842395.fna | 78.75 | 405 | 865 |
| s__UBA6256_sp002452295.fna | PM009_bin.22.orig.fa | 78.64 | 361 | 865 |
| s__UBA6256_sp002452295.fna | s__UBA6256_sp024653485.fna | 78.56 | 333 | 865 |
| s__UBA6256_sp012516125.fna | s__UBA6256_sp012516125.fna | 100.00 | 1022 | 1028 |
| s__UBA6256_sp012516125.fna | PM008_bin.53.orig.fa | 98.02 | 373 | 1028 |
| s__UBA6256_sp012516125.fna | PM012_bin.13.permissive.fa | 94.41 | 743 | 1028 |
| s__UBA6256_sp012516125.fna | s__UBA6256_sp014360195.fna | 93.57 | 832 | 1028 |
| s__UBA6256_sp012516125.fna | s__UBA6256_sp002452295.fna | 82.79 | 590 | 1028 |
| s__UBA6256_sp012516125.fna | s__UBA6256_sp012842395.fna | 81.08 | 584 | 1028 |
| s__UBA6256_sp012516125.fna | PM009_bin.22.orig.fa | 80.53 | 514 | 1028 |
| s__UBA6256_sp012516125.fna | s__UBA6256_sp024653485.fna | 80.18 | 523 | 1028 |
| s__UBA6256_sp012842395.fna | s__UBA6256_sp012842395.fna | 100.00 | 1015 | 1016 |
| s__UBA6256_sp012842395.fna | PM009_bin.22.orig.fa | 99.12 | 770 | 1016 |
| s__UBA6256_sp012842395.fna | s__UBA6256_sp012516125.fna | 81.00 | 580 | 1016 |
| s__UBA6256_sp012842395.fna | PM012_bin.13.permissive.fa | 80.96 | 550 | 1016 |
| s__UBA6256_sp012842395.fna | PM008_bin.53.orig.fa | 80.68 | 267 | 1016 |
| s__UBA6256_sp012842395.fna | s__UBA6256_sp014360195.fna | 80.65 | 591 | 1016 |
| s__UBA6256_sp012842395.fna | s__UBA6256_sp024653485.fna | 79.19 | 438 | 1016 |
| s__UBA6256_sp012842395.fna | s__UBA6256_sp002452295.fna | 78.59 | 415 | 1016 |
| s__UBA6256_sp014360195.fna | s__UBA6256_sp014360195.fna | 100.00 | 1052 | 1059 |
| s__UBA6256_sp014360195.fna | PM012_bin.13.permissive.fa | 94.07 | 774 | 1059 |
| s__UBA6256_sp014360195.fna | PM008_bin.53.orig.fa | 93.89 | 353 | 1059 |
| s__UBA6256_sp014360195.fna | s__UBA6256_sp012516125.fna | 93.49 | 825 | 1059 |

| | | | | |
|---|---|---|---|---|
| s__UBA6256_sp014360195.fna | s__UBA6256_sp002452295.fna | 82.72 | 599 | 1059 |
| s__UBA6256_sp014360195.fna | s__UBA6256_sp012842395.fna | 80.77 | 595 | 1059 |
| s__UBA6256_sp014360195.fna | PM009_bin.22.orig.fa | 80.48 | 509 | 1059 |
| s__UBA6256_sp014360195.fna | s__UBA6256_sp024653485.fna | 80.18 | 539 | 1059 |
| s__UBA6256_sp024653485.fna | s__UBA6256_sp024653485.fna | 100.00 | 983 | 985 |
| s__UBA6256_sp024653485.fna | PM012_bin.13.permissive.fa | 80.43 | 489 | 985 |
| s__UBA6256_sp024653485.fna | s__UBA6256_sp012516125.fna | 80.32 | 518 | 985 |
| s__UBA6256_sp024653485.fna | PM008_bin.53.orig.fa | 80.30 | 256 | 985 |
| s__UBA6256_sp024653485.fna | s__UBA6256_sp014360195.fna | 80.26 | 542 | 985 |
| s__UBA6256_sp024653485.fna | s__UBA6256_sp012842395.fna | 79.36 | 422 | 985 |
| s__UBA6256_sp024653485.fna | PM009_bin.22.orig.fa | 78.97 | 347 | 985 |
| s__UBA6256_sp024653485.fna | s__UBA6256_sp002452295.fna | 78.37 | 351 | 985 |
| strain.pm69.vs2.0.gt1kb.fa | strain.pm69.vs2.0.gt1kb.fa | 100.00 | 1093 | 1094 |
| strain.pm69.vs2.0.gt1kb.fa | s__Ch115_sp013178415.fna | 98.22 | 947 | 1094 |
| strain.pm69.vs2.0.gt1kb.fa | s__JABLXL01_sp013178125.fna | 83.56 | 74 | 1094 |
| strain.pm69.vs2.0.gt1kb.fa | s__JABLXL01_sp012839815.fna | 79.39 | 56 | 1094 |
| strain.pm69.vs2.0.gt1kb.fa | s__DUSO01_sp012842565.fna | 77.00 | 64 | 1094 |
