## Supplementary Results 3 for "*Thermoaminiphila catenidiffluenda* gen. nov., sp. nov.: A novel thermophilic, strictly anaerobic bacterium representing *Thermoaminiphilia* class nov., a newly described class thriving in hydrocarbon-rich habitats and biogas fermenters"

| ID | Name | PM69_Predicted | GCA_013178415.1_Prediction | GCA_012842565.1_Prediction | GCA_012839815.1_Prediction | GCA_013178125.1_Prediction | GCA_002398515.1_Prediction | GCA_002424305.1_Prediction | GCA_012518595.1_Prediction | GCA_012689375.1_Prediction | GCA_035593725.1_Prediction | GCA_900019985.1_Prediction | GCA_002452295.1_Prediction | GCA_012516125.1_Prediction | GCA_012842395.1_Prediction | GCA_014360195.1_Prediction | GCA_024653485.1_Prediction | GCA_030024085.1_Prediction |
|---|---|---|---|---|---|---|---|---|---|---|---|---|---|---|---|---|---|---|
| \|P401-PWY\| | cyanide degradation | true | false | false | false | false | false | false | false | false | false | false | false | false | false | false | false | false |
| \|PWY-6689\| | tRNA splicing I | true | false | false | false | false | false | false | false | false | false | false | false | false | false | false | false | false |
| \|PWY-7581\| | N-acetylneuraminate and N-acetylmannosamine degradation II | true | false | false | false | false | false | false | false | false | false | false | false | false | false | false | false | false |
| \|DTDPRHAMSYN-PWY\| | dTDP-β-L-rhamnose biosynthesis | true | false | false | true | false | false | false | false | false | false | false | false | false | false | false | false | false |
| \|PWY-7976\| | sulfur reduction III | true | false | false | false | false | false | true | false | false | false | false | false | false | false | false | false | false |
| \|PWY-5653\| | NAD biosynthesis from 2-amino-3-carboxymuconate semialdehyde | true | false | false | false | true | false | false | false | false | false | false | false | false | false | false | false | false |
| PYR-TO-BUT-NADPH | 3-hydroxybutyryl-CoA dehydrogenase | true | false | false | false | false | true | true | true | true | true | true | false | false | false | false | false | false |
| \|BGALACT-PWY\| | lactose degradation III | true | true | false | false | false | false | false | false | false | false | false | false | false | false | false | false | false |
| \|ETOH-ACETYLCOA-ANA-PWY\| | ethanol degradation I | true | true | false | false | false | false | false | false | false | false | false | false | false | false | false | false | false |
| \|PHESYN\| | L-phenylalanine biosynthesis I | true | true | false | false | false | false | false | false | false | false | false | false | false | false | false | false | false |
| \|PWY-6890\| | 4-amino-2-methyl-5-diphosphomethylpyrimidine biosynthesis I | true | true | false | false | false | false | false | false | false | false | false | false | false | false | false | false | false |
| \|PWY-7432\| | L-phenylalanine biosynthesis III (cytosolic, plants) | true | true | false | false | false | false | false | false | false | false | false | false | false | false | false | false | false |
| \|PWY-5041\| | S-adenosyl-L-methionine salvage II | true | true | false | true | false | false | false | false | false | false | false | true | false | false | false | false | false |
| \|PWY-5436\| | L-threonine degradation IV | true | true | false | false | false | false | false | false | true | false | false | false | false | false | false | false | false |
| \|PWY-7180\| | 2-deoxy-α-D-ribose 1-phosphate degradation | true | true | false | false | false | false | false | false | true | false | false | false | false | false | false | false | false |
| \|PWY-7335\| | UDP-N-acetyl-α-D-mannosaminouronate biosynthesis | true | true | false | true | false | false | true | false | false | false | false | false | true | true | true | true | true |
| \|PWY-7761\| | NAD salvage pathway II (PNC IV cycle) | true | true | false | false | true | false | false | false | false | false | false | false | true | true | true | true | true |
| \|PWY-8277\| | NAD de novo biosynthesis IV (anaerobic) | true | true | false | false | true | false | false | false | false | false | false | false | true | true | true | true | true |
| \|PYRIDNUCSYN-PWY\| | NAD de novo biosynthesis I | true | true | false | false | true | false | false | false | false | false | false | false | false | false | false | false | false |
| \|TCA-1\| | TCA cycle VIII (Chlamydia) | true | true | false | false | true | false | false | false | false | false | false | false | false | false | true | false | true |
| IMPSYN-HIGH-HCO3 | inosine-5'-phosphate biosynthesis at high carbonate levels | true | true | false | false | true | false | false | false | false | false | false | false | false | false | false | false | false |
| \|GLUCONEO-PWY\| | gluconeogenesis I | true | true | false | true | true | false | false | false | false | false | false | false | false | false | false | false | false |
| \|LARABITOLUTIL-PWY\| | xylitol degradation I | true | true | false | true | true | false | false | false | false | false | false | false | false | false | false | true | false |
| \|MALATE-ASPARTATE-SHUTTLE-PWY\| | malate/L-aspartate shuttle pathway | true | true | false | true | true | false | false | false | false | false | false | false | false | false | false | false | false |
| \|MANNIDEG-PWY\| | mannitol degradation I | true | true | false | true | true | false | false | false | false | false | false | true | false | false | false | false | false |
| \|PWY0-1301\| | melibiose degradation | true | true | false | true | true | false | false | false | false | false | false | false | false | false | false | false | false |
| \|PWY-5079\| | L-phenylalanine degradation III | true | true | false | true | true | false | false | false | false | false | false | false | false | false | false | false | false |
| \|PWY-5516\| | D-xylose degradation II | true | true | false | true | true | false | false | false | false | false | false | false | false | false | false | false | false |
| \|PWY-5659\| | GDP-mannose biosynthesis | true | true | false | true | true | false | false | false | false | false | false | false | false | false | false | false | false |
| \|PWY-5938\| | pyruvate fermentation to (R)-acetoin I | true | true | false | true | true | false | false | false | false | false | false | true | false | false | false | false | false |
| \|PWY-6121\| | 5-aminoimidazole ribonucleotide biosynthesis I | true | true | false | true | true | false | false | false | false | false | false | false | false | false | false | false | false |
| \|PWY-6122\| | 5-aminoimidazole ribonucleotide biosynthesis II | true | true | false | true | true | false | false | false | false | false | false | false | false | false | false | false | false |
| \|PWY-6527\| | stachyose degradation | true | true | false | true | true | false | false | false | false | false | false | false | false | false | false | false | false |
| \|PWY-6871\| | 3-methylbutanol biosynthesis (engineered) | true | true | false | true | true | false | false | false | false | false | false | false | false | false | false | false | false |
| \|PWY-7111\| | pyruvate fermentation to isobutanol (engineered) | true | true | false | true | true | false | false | false | false | false | false | false | true | true | true | true | true |
| \|PWY-7115\| | C4 photosynthetic carbon assimilation cycle, NAD-ME type | true | true | false | true | true | false | false | false | false | false | false | true | true | true | true | true | true |
| \|PWY-8097\| | 5-hydroxybenzimidazole biosynthesis (anaerobic) | true | true | false | true | true | false | false | false | false | false | false | true | false | false | false | false | false |
| \|PWY-8289\| | dZTP biosynthesis | true | true | false | true | true | false | false | false | false | false | false | false | false | false | false | false | false |
| TRPSYN-PWY2 | L-tryptophan biosynthesis (complex) | true | true | false | true | true | false | false | false | false | false | false | true | false | false | false | false | false |
| \|PWY-6599\| | guanine and guanosine salvage II | true | true | false | false | true | false | false | false | false | true | false | true | false | false | false | false | false |
| \|PWY66-21\| | ethanol degradation II | true | true | false | false | true | false | false | false | false | true | false | true | false | false | false | false | false |
| \|PWY0-1315\| | L-lactaldehyde degradation (anaerobic) | true | true | false | true | true | false | false | false | false | true | false | true | false | false | false | false | false |
| \|PWY-6333\| | acetaldehyde biosynthesis I | true | true | false | true | true | false | false | false | false | true | false | true | false | false | false | false | false |
| \|GALACTITOLCAT-PWY\| | galactitol degradation | true | true | false | true | true | false | false | true | false | false | false | true | false | false | false | false | false |
| \|PWY-3821\| | D-galactose detoxification | true | true | false | true | true | false | false | true | false | false | false | true | false | false | false | false | false |
| \|PWY-6317\| | D-galactose degradation I (Leloir pathway) | true | true | false | true | true | false | false | true | false | false | false | true | false | false | false | false | false |
| \|ARGSYNBSUB-PWY\| | L-arginine biosynthesis II (acetyl cycle) | true | true | false | true | true | false | false | true | true | true | true | true | false | false | false | false | false |
| \|PWY-2942\| | L-lysine biosynthesis III | true | true | false | false | false | true | true | true | true | true | true | true | false | false | false | false | false |
| \|PWY-5484\| | glycolysis II (from fructose 6-phosphate) | true | true | false | false | false | true | true | true | true | true | true | true | false | false | false | false | false |
| \|PYRUVDEHYD-PWY\| | pyruvate decarboxylation to acetyl CoA I | true | true | false | true | true | true | false | true | false | false | true | true | false | false | false | false | false |
| \|PWY-8088\| | dipicolinate biosynthesis | true | true | false | true | true | true | false | true | false | true | true | true | false | false | false | false | false |
| \|THREONINE-DEG2-PWY\| | L-threonine degradation II | true | true | false | false | true | true | true | true | true | true | false | true | false | false | true | false | true |
| \|GLYCOGENSYNTH-PWY\| | glycogen biosynthesis I (from ADP-D-Glucose) | true | true | false | true | true | true | true | true | true | true | true | false | false | false | false | false | false |
| \|PWY-5785\| | di-trans,poly-cis-undecaprenyl phosphate biosynthesis | true | true | false | true | true | true | true | true | true | true | true | true | false | false | false | false | false |
| \|PWY-7343\| | UDP-α-D-glucose biosynthesis | true | true | false | true | true | true | true | true | true | true | true | true | false | false | false | false | false |
| \|ALANINE-VALINESYN-PWY\| | L-alanine biosynthesis I | true | true | true | false | false | false | false | false | false | false | false | true | false | false | false | false | false |
| \|PWY-5101\| | L-isoleucine biosynthesis II | true | true | true | false | false | false | false | false | false | false | false | true | false | false | false | false | false |
| \|PWY-5108\| | L-isoleucine biosynthesis V | true | true | true | false | false | false | false | false | false | false | false | false | false | false | false | false | false |
| \|PWY-6910\| | hydroxymethylpyrimidine salvage | true | true | true | false | false | false | false | false | false | false | false | true | false | false | false | false | false |
| \|PWY-8325\| | glycine betaine degradation III | true | true | true | false | false | false | false | false | false | false | false | true | false | false | false | false | false |
| \|SERDEG-PWY\| | L-serine degradation | true | true | true | false | false | false | false | false | false | false | false | true | false | false | false | false | false |
| \|VALSYN-PWY\| | L-valine biosynthesis | true | true | true | false | false | false | false | false | false | false | false | false | false | false | false | false | false |
| \|PROSYN-PWY\| | L-proline biosynthesis I (from L-glutamate) | true | true | true | true | false | false | false | false | false | false | false | true | true | false | false | true | false |
| \|PWY-3341\| | L-proline biosynthesis III (from L-ornithine) | true | true | true | true | false | false | false | false | false | false | false | true | false | false | false | false | false |
| \|PWY-6970\| | pyruvate decarboxylation to acetyl CoA II | true | true | true | true | false | false | false | false | false | false | false | false | false | false | false | false | false |
| PYR-ACCOA-FROX | Pyruvate-ferredoxin oxidorectase to acetyl-CoA | true | true | true | true | false | false | false | false | false | false | false | true | false | false | false | false | false |
| \|PWY-5512\| | UDP-N-acetyl-D-galactosamine biosynthesis I | true | true | true | true | false | false | true | true | true | true | true | false | false | false | false | false | false |
| \|PWY-63\| | UDP-β-L-arabinose biosynthesis I (from UDP-α-D-xylose) | true | true | true | true | false | false | true | true | true | true | true | false | false | false | false | false | false |
| \|ALANINE-DEG3-PWY\| | L-alanine degradation III | true | true | true | true | true | false | false | false | false | false | false | true | false | false | false | false | false |
| \|ALANINE-SYN2-PWY\| | L-alanine biosynthesis II | true | true | true | true | true | false | false | false | false | false | false | true | false | false | false | false | false |
| \|AMMASSIM-PWY\| | ammonia assimilation cycle III | true | true | true | true | true | false | false | false | false | false | false | true | false | false | false | false | false |
| \|ARGININE-SYN4-PWY\| | L-ornithine biosynthesis II | true | true | true | true | true | false | false | false | false | false | false | true | false | false | false | false | false |
| \|ASPARTATE-DEG1-PWY\| | L-aspartate degradation I | true | true | true | true | true | false | false | false | false | false | false | true | false | false | false | false | false |
| \|ASPARTATESYN-PWY\| | L-aspartate biosynthesis | true | true | true | true | true | false | false | false | false | false | false | true | false | false | false | false | false |
| \|GLNSYN-PWY\| | L-glutamine biosynthesis I | true | true | true | true | true | false | false | false | false | false | false | false | false | false | false | false | false |
| \|GLUTAMINDEG-PWY\| | L-glutamine degradation I | true | true | true | true | true | false | false | false | false | false | false | true | false | false | false | false | false |
| \|GLUTAMINEFUM-PWY\| | L-glutamine degradation II | true | true | true | true | true | false | false | false | false | false | false | true | false | false | false | false | false |
| \|GLUTSYN-PWY\| | L-glutamate biosynthesis I | true | true | true | true | true | false | false | false | false | false | false | false | false | true | true | true | true |
| \|GLYCOLATEMET-PWY\| | glycolate and glyoxylate degradation I | true | true | true | true | true | false | false | false | false | false | false | true | false | false | false | false | false |
| \|HISTSYN-PWY\| | L-histidine biosynthesis | true | true | true | true | true | false | false | false | false | false | false | true | false | false | false | false | false |
| \|HOMOSER-THRESYN-PWY\| | L-threonine biosynthesis | true | true | true | true | true | false | false | false | false | false | false | true | false | false | false | false | false |
| \|LEUSYN-PWY\| | L-leucine biosynthesis | true | true | true | true | true | false | false | false | false | false | false | false | false | false | false | false | false |
| \|PWY0-901\| | L-selenocysteine biosynthesis I (bacteria) | true | true | true | true | true | false | false | false | false | false | false | true | false | false | false | false | false |
| \|PWY-5097\| | L-lysine biosynthesis VI | true | true | true | true | true | false | false | false | false | false | false | false | false | false | false | false | false |
| \|PWY-5104\| | L-isoleucine biosynthesis IV | true | true | true | true | true | false | false | false | false | false | false | true | true | true | true | true | true |
| \|PWY-5481\| | pyruvate fermentation to (S)-lactate | true | true | true | true | true | false | false | false | false | false | false | true | false | false | false | false | false |
| \|PWY-5886\| | 3-(4-hydroxyphenyl)pyruvate biosynthesis | true | true | true | true | true | false | false | false | false | false | false | true | true | false | false | false | true |
| \|PWY-5921\| | glutaminyl-tRNAgln biosynthesis via transamidation | true | true | true | true | true | false | false | false | false | false | false | true | false | false | false | false | false |
| \|PWY-6164\| | 3-dehydroquinate biosynthesis I | true | true | true | true | true | false | false | false | false | false | false | false | true | true | true | true | true |
| \|PWY-6466\| | pyridoxal 5'-phosphate biosynthesis II | true | true | true | true | true | false | false | false | false | false | false | true | true | false | false | false | true |
| \|PWY-6578\| | 8-amino-7-oxononanoate biosynthesis III | true | true | true | true | true | false | false | false | false | false | false | true | true | false | false | false | true |
| \|PWY-7219\| | adenosine ribonucleotides de novo biosynthesis | true | true | true | true | true | false | false | false | false | false | false | false | true | true | true | true | true |
| \|PWY-7268\| | cytosolic NADPH production (yeast) | true | true | true | true | true | false | false | false | false | false | false | true | false | false | false | false | false |
| \|PWY-7395\| | D-galactosamine and N-acetyl-D-galactosamine degradation | true | true | true | true | true | false | false | false | false | false | false | true | false | false | false | false | false |
| \|PWY-7977\| | L-methionine biosynthesis IV | true | true | true | true | true | false | false | false | false | false | false | false | true | false | true | false | true |
| NA-F0F1ATPSYN | Na(+)-translocating ATPase | true | true | true | true | true | false | false | false | false | false | false | true | false | false | false | false | false |
| \|HOMOSERSYN-PWY\| | L-homoserine biosynthesis | true | true | true | true | true | false | false | true | false | false | false | true | false | false | false | false | false |
| THRBIOSYN-GS | Threonine biosynthis from aspartate via homoserine (gapseq) | true | true | true | true | true | false | false | true | false | false | false | false | false | false | false | false | false |
| \|PWY-7685\| | L-malate degradation I | true | true | true | true | true | false | false | true | true | true | true | true | false | false | false | false | false |
| \|PWY-7686\| | L-malate degradation II | true | true | true | true | true | false | false | true | true | true | true | true | false | false | false | false | false |
| \|PWY0-1465\| | D-malate degradation | true | true | true | true | true | false | true | false | true | true | true | true | false | false | false | false | false |
| \|PWY0-823\| | L-arginine degradation III (arginine decarboxylase/agmatinase pathway) | true | true | true | true | true | false | true | false | true | true | true | true | false | false | false | false | false |
| \|PWY-40\| | putrescine biosynthesis I | true | true | true | true | true | false | true | false | false | true | false | true | false | false | false | false | false |
| \|PWY-5535\| | acetate and ATP formation from acetyl-CoA II | true | true | true | true | true | false | true | false | false | true | false | true | false | false | false | false | false |
| \|CYSTSYN-PWY\| | L-cysteine biosynthesis I | true | true | true | true | true | false | true | true | true | true | true | true | false | false | false | false | false |
| \|PWY1-2\| | L-alanine degradation IV | true | true | true | true | true | false | true | true | false | true | true | false | false | false | false | false | false |
| \|PWY-7870\| | L-cysteine biosynthesis VII (from S-sulfo-L-cysteine) | true | true | true | true | true | false | true | true | false | true | true | true | false | false | false | false | false |
| \|GLUCONSUPER-PWY\| | D-gluconate degradation | true | true | true | true | true | false | true | true | true | true | true | true | false | false | false | false | false |
| \|PWY-7344\| | UDP-α-D-galactose biosynthesis | true | true | true | true | false | false | true | true | true | true | true | true | false | false | false | false | false |
| \|LEU-DEG2-PWY\| | L-leucine degradation I | true | true | true | false | false | true | true | false | false | true | true | true | false | false | false | false | true |
| \|PWY-6694\| | oxalate degradation I | true | true | true | false | false | true | true | true | true | true | true | false | false | false | false | false | false |
| \|GLYCINE-SYN2-PWY\| | glycine biosynthesis II | true | true | true | true | false | true | true | true | true | true | true | false | false | false | false | false | false |
| \|SAM-PWY\| | S-adenosyl-L-methionine biosynthesis | true | true | true | true | false | true | true | true | true | true | false | false | true | true | true | false | false |
| \|PWY-7566\| | L-gulonate degradation | true | true | true | true | true | true | false | false | false | false | false | false | true | true | true | true | false |
| \|GLYSYN-PWY\| | glycine biosynthesis I | true | true | true | true | true | true | true | false | true | true | true | false | false | false | false | false | false |
| \|GLYSYN-THR-PWY\| | glycine biosynthesis IV | true | true | true | true | true | true | true | false | true | true | true | true | false | false | false | false | false |
| \|ARGDEG-V-PWY\| | L-arginine degradation X (arginine monooxygenase pathway) | true | true | true | true | true | true | true | true | false | true | true | true | true | true | true | true | true |
| \|PWY0-1299\| | arginine dependent acid resistance | true | true | true | true | true | true | true | true | false | true | true | true | false | false | false | false | false |
| PI-ABC-TRANS | Phosphate ABC transporter | true | true | true | true | true | true | true | true | false | true | true | true | false | false | false | false | false |
| LDH-ETF-PWY | Electron bifurcating LDH/Etf complex | true | true | true | false | true | true | true | true | true | true | true | false | false | false | false | false | false |
| \|GLUGLNSYN-PWY\| | L-glutamate biosynthesis IV | true | true | true | true | true | true | true | true | true | true | true | true | false | false | false | false | false |
| \|GLUTAMATE-DEG1-PWY\| | L-glutamate degradation I | true | true | true | true | true | true | true | true | true | true | true | true | false | false | false | false | false |
| \|GLUTAMATE-SYN2-PWY\| | L-glutamate biosynthesis II | true | true | true | true | true | true | true | true | true | true | true | false | false | true | false | true | true |
| \|GLUTSYNIII-PWY\| | L-glutamate biosynthesis III | true | true | true | true | true | true | true | true | true | true | true | true | true | true | true | false | true |
| \|GLYCLEAV-PWY\| | glycine cleavage | true | true | true | true | true | true | true | true | true | true | true | true | false | false | false | false | false |
| \|GLYSYN-ALA-PWY\| | glycine biosynthesis III | true | true | true | true | true | true | true | true | true | true | true | true | false | false | false | false | false |
| \|LCYSDEG-PWY\| | L-cysteine degradation II | true | true | true | true | true | true | true | true | true | true | true | false | false | false | false | false | false |
| \|NONMEVIPP-PWY\| | methylerythritol phosphate pathway I | true | true | true | true | true | true | true | true | true | true | true | true | false | false | false | false | false |
| \|PWY0-1021\| | L-alanine biosynthesis III | true | true | true | true | true | true | true | true | true | true | true | true | false | false | false | false | false |
| \|PWY0-1296\| | purine ribonucleosides degradation | true | true | true | true | true | true | true | true | true | true | true | true | false | false | false | false | false |
| \|PWY0-1517\| | sedoheptulose bisphosphate bypass | true | true | true | true | true | true | true | true | true | true | true | true | false | false | false | false | false |
| \|PWY0-1587\| | N6-L-threonylcarbamoyladenosine37-modified tRNA biosynthesis | true | true | true | true | true | true | true | true | true | true | true | true | false | false | false | false | false |
| \|PWY0-662\| | PRPP biosynthesis | true | true | true | true | true | true | true | true | true | true | true | false | false | false | false | false | false |
| \|PWY-1722\| | formate assimilation into 5,10-methylenetetrahydrofolate | true | true | true | true | true | true | true | true | true | true | true | true | false | false | false | false | false |
| \|PWY-2161\| | folate polyglutamylation | true | true | true | true | true | true | true | true | true | true | true | true | false | false | false | false | false |
| \|PWY3O-355\| | stearate biosynthesis III (fungi) | true | true | true | true | true | true | true | true | true | true | true | true | false | false | false | false | false |
| \|PWY-4381\| | fatty acid biosynthesis initiation (type II) | true | true | true | true | true | true | true | true | true | true | true | true | false | false | false | false | false |
| \|PWY490-4\| | L-asparagine biosynthesis III (tRNA-dependent) | true | true | true | true | true | true | true | true | true | true | true | false | false | false | false | false | false |
| \|PWY-5382\| | hydrogen oxidation II (aerobic, NAD) | true | true | true | true | true | true | true | true | true | true | true | true | true | false | true | false | true |
| \|PWY-5766\| | L-glutamate degradation X | true | true | true | true | true | true | true | true | true | true | true | true | false | false | false | false | false |
| \|PWY-5966-1\| | fatty acid biosynthesis initiation (type I) | true | true | true | true | true | true | true | true | true | true | true | false | false | false | false | false | false |
| \|PWY-5994\| | palmitate biosynthesis I (type I fatty acid synthase) | true | true | true | true | true | true | true | true | true | true | true | true | false | false | false | false | false |
| \|PWY-6028\| | acetoin degradation | true | true | true | true | true | true | true | true | true | true | true | true | false | false | false | false | false |
| \|PWY-6386\| | UDP-N-acetylmuramoyl-pentapeptide biosynthesis II (lysine-containing) | true | true | true | true | true | true | true | true | true | true | true | true | false | false | false | false | false |
| \|PWY-6387\| | UDP-N-acetylmuramoyl-pentapeptide biosynthesis I (meso-diaminopimelate containing) | true | true | true | true | true | true | true | true | true | true | true | true | false | false | false | false | false |
| \|PWY-6610\| | adenine salvage | true | true | true | true | true | true | true | true | true | true | true | false | false | false | false | false | false |
| \|PWY-6620\| | guanine and guanosine salvage I | true | true | true | true | true | true | true | true | true | true | true | true | false | false | false | false | false |
| \|PWY-6697\| | oxalate degradation IV | true | true | true | true | true | true | true | true | true | true | false | true | false | false | false | false | false |
| \|PWY-6744\| | hydrogen production I | true | true | true | true | true | true | true | true | true | true | true | true | false | false | false | false | false |
| \|PWY-6758\| | hydrogen production II | true | true | true | true | true | true | true | true | true | true | true | false | false | false | false | false | false |
| \|PWY-6938-1\| | NADH repair (prokaryotes) | true | true | true | true | true | true | true | true | true | true | true | true | false | false | false | false | false |
| \|PWY-7183\| | pyrimidine nucleobases salvage I | true | true | true | true | true | true | true | true | true | true | true | true | false | false | false | false | false |
| \|PWY-7560\| | methylerythritol phosphate pathway II | true | true | true | true | true | true | true | true | true | true | true | true | false | false | false | false | false |
| \|PWY-7790\| | UMP biosynthesis II | true | true | true | true | true | true | true | true | true | true | true | true | false | false | false | false | false |
| \|PWY-7791\| | UMP biosynthesis III | true | true | true | true | true | true | true | true | true | true | true | true | false | false | false | false | false |
| \|PWY-7980\| | ATP biosynthesis | true | true | true | true | true | true | true | true | true | true | true | false | false | false | false | false | false |
| \|PWY-8072\| | alanine racemization | true | true | true | true | true | true | true | true | true | true | true | true | false | false | false | false | false |
| \|PWY-8105\| | queuosine biosynthesis II (queuine salvage) | true | true | true | true | true | true | true | true | true | true | true | true | false | false | false | false | false |
| \|PWY-8136\| | NADPH repair (prokaryotes) | true | true | true | true | true | true | true | true | true | true | true | true | false | false | false | false | false |
| \|PWY-8148\| | NADP biosynthesis | true | true | true | true | true | true | true | true | true | true | true | true | false | false | false | false | false |
| \|PWY-8404\| | glycolysis VI (from fructose) | true | true | true | true | true | true | true | true | true | true | true | true | false | false | false | false | false |
| \|TRNA-CHARGING-PWY\| | tRNA charging | true | true | true | true | true | true | true | true | true | true | true | true | false | false | false | false | false |
| \|UDPNAGSYN-PWY\| | UDP-N-acetyl-D-glucosamine biosynthesis I | true | true | true | true | true | true | true | true | true | true | true | true | false | false | false | false | false |
| NADP-RED-FD-NADH | Coupled NADP reduction via ferredoxin and NADH | true | true | true | true | true | true | true | true | true | true | true | true | false | false | false | false | false |
| PEPTIDOGLYCANSYN-PWY2 | peptidoglycan biosynthesis I (meso-diaminopimelate containing) | true | true | true | true | true | true | true | true | true | true | true | true | false | false | false | false | false |
| NA-TRANSLOCATING-RNF | Na(+)-translocating ferredoxin:NAD(+) oxidoreductase complex | true | true | true | true | true | true | true | true | true | true | true | false | false | false | false | false | false |
| \|12DICHLORETHDEG-PWY\| | 1,2-dichloroethane degradation | false | false | false | false | false | false | false | false | false | false | false | false | false | false | false | false | true |
| \|2AMINOBENZDEG-PWY\| | anthranilate degradation III (anaerobic) | false | false | false | false | false | false | false | false | false | false | false | false | false | false | false | false | true |
| \|ALKANEMONOX-PWY\| | two-component alkanesulfonate monooxygenase | false | false | false | false | false | false | false | false | false | false | false | true | false | false | false | false | false |
| \|AMMOXID-PWY\| | ammonia oxidation I (aerobic) | false | false | false | false | false | false | false | false | false | false | false | true | false | false | false | false | false |
| \|ANAGLYCOLYSIS-PWY\| | glycolysis III (from glucose) | false | false | false | false | false | false | false | false | false | false | false | true | false | false | false | false | false |
| \|ARGASEDEG-PWY\| | L-arginine degradation I (arginase pathway) | false | false | false | false | false | false | false | false | false | false | false | true | false | false | false | false | false |
| \|CATECHOL-ORTHO-CLEAVAGE-PWY\| | catechol degradation to β-ketoadipate | false | false | false | false | false | false | false | false | false | false | false | true | false | false | false | false | false |
| \|CENTBENZCOA-PWY\| | benzoyl-CoA degradation II (anaerobic) | false | false | false | false | false | false | false | false | false | false | false | true | false | false | false | false | false |
| \|DESULFONATION-PWY\| | benzenesulfonate degradation | false | false | false | false | false | false | false | false | false | false | false | true | false | false | false | false | false |
| \|ENTNER-DOUDOROFF-PWY\| | Entner-Doudoroff shunt | false | false | false | false | false | false | false | false | false | false | false | true | false | false | false | true | false |
| \|FASYN-ELONG-PWY\| | fatty acid elongation -- saturated | false | false | false | false | false | false | false | false | false | false | false | false | false | false | false | true | true |
| \|HEMESYN2-PWY\| | heme b biosynthesis II (oxygen-independent) | false | false | false | false | false | false | false | false | false | false | false | true | false | false | false | false | false |
| \|HOMOCYSDEGR-PWY\| | L-cysteine biosynthesis III (from L-homocysteine) | false | false | false | false | false | false | false | false | false | false | false | true | false | false | false | false | false |
| \|HOMOSER-METSYN-PWY\| | L-methionine biosynthesis I | false | false | false | false | false | false | false | false | false | false | false | true | false | false | false | false | false |
| \|KDOSYN-PWY\| | Kdo transfer to lipid IVA (E. coli) | false | false | false | false | false | false | false | false | false | false | false | true | false | false | false | false | false |
| \|METH-ACETATE-PWY\| | methanogenesis from acetate | false | false | false | false | false | false | false | false | false | false | false | true | false | false | false | false | false |
| \|METHIONINE-DEG1-PWY\| | L-methionine degradation I (to L-homocysteine) | false | false | false | false | false | false | false | false | false | false | false | true | false | false | false | false | false |
| \|METHYLGALLATE-DEGRADATION-PWY\| | methylgallate degradation | false | false | false | false | false | false | false | false | false | false | false | true | false | false | false | false | false |
| \|NAD-BIOSYNTHESIS-III\| | NAD biosynthesis from nicotinamide | false | false | false | false | false | false | false | false | false | false | false | true | false | false | false | false | false |
| \|OCTOPINEDEG-PWY\| | octopine degradation | false | false | false | false | false | false | false | false | false | false | false | true | false | false | false | false | false |
| \|ORN-AMINOPENTANOATE-CAT-PWY\| | L-ornithine degradation I (L-proline biosynthesis) | false | false | false | false | false | false | false | false | false | false | false | true | false | false | false | false | false |
| \|P203-PWY\| | sulfur disproportionation I (anaerobic) | false | false | false | false | false | false | false | false | false | false | false | true | false | false | false | false | false |
| \|P581-PWY\| | thiocyanate degradation I | false | false | false | false | false | false | false | false | false | false | false | false | true | false | true | false | true |
| \|PWY0-1565\| | D-lactate to cytochrome bo oxidase electron transfer | false | false | false | false | false | false | false | false | false | false | false | true | false | false | false | false | false |
| \|PWY18C3-23\| | guaiacol biosynthesis | false | false | false | false | false | false | false | false | false | false | false | false | false | false | false | true | false |
| \|PWY18C3-7\| | cembratrienediol biosynthesis | false | false | false | false | false | false | false | false | false | false | false | false | true | true | true | true | true |
| \|PWY1G-170\| | formaldehyde oxidation III (mycothiol-dependent) | false | false | false | false | false | false | false | false | false | false | false | false | true | false | false | false | false |
| \|PWY1YI0-1\| | arsenate detoxification V | false | false | false | false | false | false | false | false | false | false | false | true | false | false | false | false | false |
| \|PWY-2661\| | trehalose biosynthesis V | false | false | false | false | false | false | false | false | false | false | false | false | false | false | true | false | false |
| \|PWY-2881\| | cytokinins 7-N-glucoside biosynthesis | false | false | false | false | false | false | false | false | false | false | false | false | false | false | false | true | false |
| \|PWY-361\| | monolignol biosynthesis | false | false | false | false | false | false | false | false | false | false | false | false | false | false | false | true | false |
| \|PWY-3841\| | folate transformations II (plants) | false | false | false | false | false | false | false | false | false | false | false | false | true | true | true | true | true |
| \|PWY-3901\| | berberine biosynthesis | false | false | false | false | false | false | false | false | false | false | false | false | false | true | false | false | false |
| \|PWY-3941\| | β-alanine biosynthesis II | false | false | false | false | false | false | false | false | false | false | false | false | false | false | false | false | false |
| \|PWY-3961\| | phosphopantothenate biosynthesis II | false | false | false | false | false | false | false | false | false | false | false | false | false | false | false | false | false |
| \|PWY-3981\| | β-alanine biosynthesis I | false | false | false | false | false | false | false | false | false | false | false | false | false | false | false | false | false |
| \|PWY3DJ-11281\| | sphingomyelin metabolism | false | false | false | false | false | false | false | false | false | false | false | false | true | true | true | true | true |
| \|PWY3DJ-11470\| | sphingosine and sphingosine-1-phosphate metabolism | false | false | false | false | false | false | false | false | false | false | false | false | false | false | false | false | false |
| \|PWY3DJ-12\| | ceramide de novo biosynthesis | false | false | false | false | false | false | false | false | false | false | false | false | false | false | false | false | false |
| \|PWY3DJ-35471\| | L-ascorbate biosynthesis IV (animals, D-glucuronate pathway) | false | false | false | false | false | false | false | false | false | false | false | false | false | false | false | false | false |
| \|PWY3O-1743\| | D-mannose degradation II | false | false | false | false | false | false | false | false | false | false | false | false | false | false | false | false | false |
| \|PWY3O-1801\| | palmitoleate biosynthesis IV (fungi and animals) | false | false | false | false | false | false | false | false | false | false | false | false | false | false | false | false | false |
| \|PWY3O-19\| | ubiquinol-6 biosynthesis (late decarboxylation) | false | false | false | false | false | false | false | false | false | false | false | false | false | false | false | false | false |
| \|PWY3O-246\| | (R,R)-butanediol degradation | false | false | false | false | false | false | false | false | false | false | false | false | false | false | false | false | false |
| \|PWY3O-4106\| | NAD salvage pathway IV (from nicotinamide riboside) | false | false | false | false | false | false | false | false | false | false | false | false | false | false | false | false | false |
| \|PWY3O-4108\| | L-tyrosine degradation III | false | false | false | false | false | false | false | false | false | false | false | false | false | false | false | false | false |
| \|PWY3O-440\| | pyruvate fermentation to acetoin | false | false | false | false | false | false | false | false | false | false | false | false | false | false | false | false | false |
| \|PWY3O-450\| | phosphatidylcholine biosynthesis I | false | false | false | false | false | false | false | false | false | false | false | false | false | false | false | false | false |
| \|PWY3O-6\| | dehydro-D-arabinono-1,4-lactone biosynthesis | false | false | false | false | false | false | false | false | false | false | false | false | false | false | false | false | false |
| \|PWY-4002\| | L-asparagine degradation II | false | false | false | false | false | false | false | false | false | false | false | false | false | false | false | false | false |
| \|PWY-401\| | galactolipid biosynthesis I | false | false | false | false | false | false | false | false | false | false | false | false | false | false | false | false | false |
| \|PWY-4021\| | β-alanine betaine biosynthesis | false | false | false | false | false | false | false | false | false | false | false | false | true | false | true | false | false |
| \|PWY-4061\| | glutathione-mediated detoxification I | false | false | false | false | false | false | false | false | false | false | false | false | false | false | false | false | false |
| \|PWY-4081\| | glutathione-peroxide redox reactions | false | false | false | false | false | false | false | false | false | false | false | false | false | false | false | false | false |
| \|PWY-4101\| | D-sorbitol degradation I | false | false | false | false | false | false | false | false | false | false | false | false | false | false | false | false | false |
| \|PWY-4121\| | glutathionylspermidine biosynthesis | false | false | false | false | false | false | false | false | false | false | false | false | false | false | false | false | false |
| \|PWY-4181\| | glutathione amide metabolism | false | false | false | false | false | false | false | false | false | false | false | false | false | false | false | false | false |
| \|PWY-4201\| | volatile cinnamoic ester biosynthesis | false | false | false | false | false | false | false | false | false | false | false | false | false | false | false | false | false |
| \|PWY-4202\| | arsenic detoxification (mammals) | false | false | false | false | false | false | false | false | false | false | false | false | false | false | false | false | false |
| \|PWY-4203\| | volatile benzenoid biosynthesis I (ester formation) | false | false | false | false | false | false | false | false | false | false | false | false | false | false | false | false | false |
| \|PWY-4261\| | glycerol degradation I | false | false | false | false | false | false | false | false | false | false | false | false | false | false | false | false | false |
| \|PWY-4281\| | L-proline biosynthesis IV | false | false | false | false | false | false | false | false | false | false | false | false | false | false | false | false | false |
| \|PWY-4302\| | aerobic respiration III (alternative oxidase pathway) | false | false | false | false | false | false | false | false | false | false | false | false | false | false | false | false | false |
| \|PWY-4341\| | L-glutamate biosynthesis V | false | false | false | false | false | false | false | false | false | false | false | false | true | true | true | true | false |
| \|PWY-4361\| | S-methyl-5-thio-α-D-ribose 1-phosphate degradation I | false | false | false | false | false | false | false | false | false | false | false | false | false | false | false | false | false |
| \|PWY-43\| | putrescine biosynthesis II | false | false | false | false | false | false | false | false | false | false | false | false | false | false | false | false | false |
| \|PWY-4421\| | curcumin glucoside biosynthesis | false | false | false | false | false | false | false | false | false | false | false | false | false | false | false | false | false |
| \|PWY-4441\| | DIMBOA-glucoside activation | false | false | false | false | false | false | false | false | false | false | false | false | false | false | false | false | false |
| \|PWY-4502\| | wighteone and luteone biosynthesis | false | false | false | false | false | false | false | false | false | false | false | false | false | false | false | false | false |
| \|PWY-4521\| | arsenite to oxygen electron transfer | false | false | false | false | false | false | false | false | false | false | false | false | false | false | false | false | false |
| \|PWY-4601\| | arsenate reduction (respiratory) | false | false | false | false | false | false | false | false | false | false | false | true | false | false | false | false | false |
| \|PWY-4621\| | arsenic detoxification (yeast) | false | false | false | false | false | false | false | false | false | false | false | false | false | false | false | false | false |
| \|PWY-4661\| | 1D-myo-inositol hexakisphosphate biosynthesis III (Spirodela polyrrhiza) | false | false | false | false | false | false | false | false | false | false | false | false | false | false | false | false | false |
| \|PWY-4681\| | kievitone biosynthesis | false | false | false | false | false | false | false | false | false | false | false | false | false | false | false | false | false |
| \|PWY-46\| | putrescine biosynthesis III | false | false | false | false | false | false | false | false | false | false | false | false | false | false | false | false | false |
| \|PWY-4702\| | phytate degradation I | false | false | false | false | false | false | false | false | false | false | false | false | false | false | false | false | false |
| \|PWY-4722\| | creatinine degradation II | false | false | false | false | false | false | false | false | false | false | false | false | false | false | false | false | false |
| \|PWY-4741\| | creatinine degradation III | false | false | false | false | false | false | false | false | false | false | false | false | false | false | false | false | false |
| \|PWY-4781\| | phytate degradation II | false | false | false | false | false | false | false | false | false | false | false | false | false | false | false | false | false |
| \|PWY-4801\| | aloesone biosynthesis I | false | false | false | false | false | false | false | false | false | false | false | false | false | false | false | false | false |
| \|PWY-481\| | ethylbenzene degradation (anaerobic) | false | false | false | false | false | false | false | false | false | false | false | false | false | false | false | false | false |
| \|PWY-4841\| | UDP-α-D-glucuronate biosynthesis (from myo-inositol) | false | false | false | false | false | false | false | false | false | false | false | false | false | false | false | false | false |
| \|PWY490-3\| | nitrate reduction VI (assimilatory) | false | false | false | false | false | false | false | false | false | false | false | false | false | false | false | false | false |
| \|PWY-4921\| | protein citrullination | false | false | false | false | false | false | false | false | false | false | false | false | false | false | false | false | false |
| \|PWY-4922\| | 6-methoxymellein biosynthesis | false | false | false | false | false | false | false | false | false | false | false | false | false | false | false | false | false |
| \|PWY-4942\| | sterculate biosynthesis | false | false | false | false | false | false | false | false | false | false | false | false | false | false | false | false | false |
| \|PWY-4961\| | β-pyrazole-1-ylalanine biosynthesis | false | false | false | false | false | false | false | false | false | false | false | false | false | false | false | false | false |
| \|PWY-4981\| | L-proline biosynthesis II (from arginine) | false | false | false | false | false | false | false | false | false | false | false | false | false | false | false | false | false |
| \|PWY-4983\| | nitric oxide biosynthesis II (mammals) | false | false | false | false | false | false | false | false | false | false | false | false | false | false | false | false | false |
| \|PWY-4984\| | urea cycle | false | false | false | false | false | false | false | false | false | false | false | false | false | false | false | false | false |
| \|PWY-4985\| | mimosine biosynthesis | false | false | false | false | false | false | false | false | false | false | false | false | false | false | false | false | false |
| \|PWY4FS-11\| | L-ascorbate biosynthesis II (plants, L-gulose pathway) | false | false | false | false | false | false | false | false | false | false | false | false | false | false | false | false | false |
| \|PWY4FS-12\| | VTC2 cycle | false | false | false | false | false | false | false | false | false | false | false | false | false | false | false | false | false |
| \|PWY4FS-13\| | extended VTC2 cycle | false | false | false | false | false | false | false | false | false | false | false | false | false | false | false | false | false |
| \|PWY4FS-17\| | abscisic acid biosynthesis shunt | false | false | false | false | false | false | false | false | false | false | false | false | false | false | false | false | false |
| \|PWY4FS-3\| | phosphatidylcholine biosynthesis III | false | false | false | false | false | false | false | false | false | false | false | false | false | false | false | false | false |
| \|PWY4FS-4\| | phosphatidylcholine biosynthesis IV | false | false | false | false | false | false | false | false | false | false | false | false | false | false | false | false | false |
| \|PWY4FS-6\| | phosphatidylethanolamine biosynthesis II | false | false | false | false | false | false | false | false | false | false | false | false | false | false | false | false | false |
| \|PWY-4\| | UDP-α-D-galacturonate biosynthesis II (from D-galacturonate) | false | false | false | false | false | false | false | false | false | false | false | false | false | false | false | false | false |
| \|PWY-5001\| | 1,3,6,7-tetrahydroxyxanthone biosynthesis (from benzoyl-CoA) | false | false | false | false | false | false | false | false | false | false | false | false | false | false | false | false | false |
| \|PWY-5002\| | tetrahydroxyxanthone biosynthesis (from 3-hydroxybenzoate) | false | false | false | false | false | false | false | false | false | false | false | false | false | false | false | false | false |
| \|PWY-5021\| | willardiine and isowillardiine biosynthesis | false | false | false | false | false | false | false | false | false | false | false | false | false | false | false | false | false |
| \|PWY-5022\| | 4-aminobutanoate degradation V | false | false | false | false | false | false | false | false | false | false | false | false | false | false | false | false | false |
| \|PWY-5024\| | L-arginine degradation XI | false | false | false | false | false | false | false | false | false | false | false | false | false | false | false | false | false |
| \|PWY-5025\| | indole-3-acetate biosynthesis IV (bacteria) | false | false | false | false | false | false | false | false | false | false | false | false | false | false | false | false | false |
| \|PWY-5027\| | phylloquinol biosynthesis | false | false | false | false | false | false | false | false | false | false | false | false | false | false | false | false | false |
| \|PWY-5028\| | L-histidine degradation II | false | false | false | false | false | false | false | false | false | false | false | false | false | false | false | false | false |
| \|PWY-5029\| | 3-(imidazol-5-yl)lactate salvage | false | false | false | false | false | false | false | false | false | false | false | false | false | false | false | false | false |
| \|PWY-5031\| | L-histidine degradation V | false | false | false | false | false | false | false | false | false | false | false | false | false | false | false | false | false |
| \|PWY-5032\| | ent-kaurene biosynthesis I | false | false | false | false | false | false | false | false | false | false | false | false | false | false | false | false | false |
| \|PWY-5033\| | nicotinate degradation II | false | false | false | false | false | false | false | false | false | false | false | false | false | false | false | false | false |
| \|PWY-5034\| | gibberellin A12 biosynthesis | false | false | false | false | false | false | false | false | false | false | false | false | false | false | false | false | false |
| \|PWY-5035\| | gibberellin biosynthesis III (early C-13 hydroxylation) | false | false | false | false | false | false | false | false | false | false | false | false | false | false | false | false | false |
| \|PWY-5036\| | gibberellin biosynthesis II (early C-3 hydroxylation) | false | false | false | false | false | false | false | false | false | false | false | false | false | false | false | false | false |
| \|PWY-5037\| | caffeine biosynthesis I | false | false | false | false | false | false | false | false | false | false | false | false | false | false | false | false | false |
| \|PWY-5038\| | caffeine biosynthesis II (via paraxanthine) | false | false | false | false | false | false | false | false | false | false | false | false | false | false | false | false | false |
| \|PWY-5039\| | theobromine biosynthesis I | false | false | false | false | false | false | false | false | false | false | false | false | false | false | false | false | false |
| \|PWY-5040\| | theobromine biosynthesis II (via xanthine) | false | false | false | false | false | false | false | false | false | false | false | false | false | false | false | false | false |
| \|PWY-5045\| | pinosylvin metabolism | false | false | false | false | false | false | false | false | false | false | false | false | false | false | false | false | false |
| \|PWY-5046\| | 2-oxoisovalerate decarboxylation to isobutanoyl-CoA | false | false | false | false | false | false | false | false | false | false | false | false | false | false | false | false | false |
| \|PWY-5047\| | gibberellin biosynthesis IV (Gibberella fujikuroi) | false | false | false | false | false | false | false | false | false | false | false | false | false | false | false | false | false |
| \|PWY-5048\| | rosmarinic acid biosynthesis I | false | false | false | false | false | false | false | false | false | false | false | false | false | false | false | false | false |
| \|PWY-5049\| | rosmarinic acid biosynthesis II | false | false | false | false | false | false | false | false | false | false | false | false | false | false | false | false | false |
| \|PWY-5054\| | D-sorbitol biosynthesis I | false | false | false | false | false | false | false | false | false | false | false | false | false | false | false | false | false |
| \|PWY-5055\| | nicotinate degradation III | false | false | false | false | false | false | false | false | false | false | false | false | false | false | false | false | false |
| \|PWY-5057\| | L-valine degradation II | false | false | false | false | false | false | false | false | false | false | false | false | false | false | false | false | false |
| \|PWY-5059\| | pinobanksin biosynthesis | false | false | false | false | false | false | false | false | false | false | false | false | false | false | false | false | false |
| \|PWY-5060\| | luteolin biosynthesis | false | false | false | false | false | false | false | false | false | false | false | false | false | false | false | false | false |
| \|PWY-5061\| | 6,7,4'-trihydroxyisoflavone biosynthesis | false | false | false | false | false | false | false | false | false | false | false | false | false | false | false | false | false |
| \|PWY-5063\| | phytyl diphosphate biosynthesis | false | false | false | false | false | false | false | false | false | false | false | false | false | false | false | false | false |
| \|PWY-5064\| | chlorophyll a biosynthesis II | false | false | false | false | false | false | false | false | false | false | false | false | false | false | false | false | false |
| \|PWY-5067\| | glycogen biosynthesis II (from UDP-D-Glucose) | false | false | false | false | false | false | false | false | false | false | false | false | false | false | false | false | false |
| \|PWY-5068\| | chlorophyll cycle | false | false | false | false | false | false | false | false | false | false | false | false | false | false | false | false | false |
| \|PWY-5070\| | gibberellin biosynthesis I (non C-3, non C-13 hydroxylation) | false | false | false | false | false | false | false | false | false | false | false | false | false | false | false | false | false |
| \|PWY-5074\| | mevalonate degradation | false | false | false | false | false | false | false | false | false | false | false | false | false | false | false | false | false |
| \|PWY-5075\| | L-leucine degradation II | false | false | false | false | false | false | false | false | false | false | false | false | false | false | false | false | false |
| \|PWY-5076\| | L-leucine degradation III | false | false | false | false | false | false | false | false | false | false | false | false | false | false | false | false | false |
| \|PWY-5078\| | L-isoleucine degradation II | false | false | false | false | false | false | false | false | false | false | false | false | false | false | false | false | false |
| \|PWY-5080\| | very long chain fatty acid biosynthesis I | false | false | false | false | false | false | false | false | false | false | false | false | false | false | false | false | false |
| \|PWY-5081\| | L-tryptophan degradation VIII (to tryptophol) | false | false | false | false | false | false | false | false | false | false | false | false | false | false | false | false | false |
| \|PWY-5082\| | L-methionine degradation III | false | false | false | false | false | false | false | false | false | false | false | false | false | false | false | false | false |
| \|PWY-5083\| | NAD(P)/NADPH interconversion | false | false | false | false | false | false | false | false | false | false | false | false | false | false | false | false | false |
| \|PWY-5084\| | 2-oxoglutarate decarboxylation to succinyl-CoA | false | false | false | false | false | false | false | false | false | false | false | false | false | false | false | false | false |
| \|PWY-5086\| | chlorophyll a biosynthesis I | false | false | false | false | false | false | false | false | false | false | false | false | false | false | false | false | false |
| \|PWY-5087\| | L-glutamate degradation VI (to pyruvate) | false | false | false | false | false | false | false | false | false | false | false | false | false | false | false | false | false |
| \|PWY-5094\| | naringenin glycoside biosynthesis | false | false | false | false | false | false | false | false | false | false | false | false | false | false | false | false | false |
| \|PWY-5098\| | chlorophyll a degradation I | false | false | false | false | false | false | false | false | false | false | false | false | false | false | false | false | false |
| \|PWY-5103\| | L-isoleucine biosynthesis III | false | false | false | false | false | false | false | false | false | false | false | false | false | false | false | false | false |
| \|PWY-5105\| | hesperitin glycoside biosynthesis | false | false | false | false | false | false | false | false | false | false | false | false | false | false | false | false | false |
| \|PWY-5107\| | phytol salvage pathway | false | false | false | false | false | false | false | false | false | false | false | false | false | false | false | false | false |
| \|PWY-5110\| | trigonelline biosynthesis | false | false | false | false | false | false | false | false | false | false | false | false | false | false | false | false | false |
| \|PWY-5113\| | UDP-D-apiose biosynthesis (from UDP-D-glucuronate) | false | false | false | false | false | false | false | false | false | false | false | false | false | false | false | false | false |
| \|PWY-5115\| | GDP-L-galactose biosynthesis | false | false | false | false | false | false | false | false | false | false | false | false | false | false | false | false | false |
| \|PWY-5116\| | sakuranetin biosynthesis | false | false | false | false | false | false | false | false | false | false | false | false | false | false | false | false | false |
| \|PWY-5118\| | ponciretin biosynthesis | false | false | false | false | false | false | false | false | false | false | false | false | false | false | false | false | false |
| \|PWY-5119\| | acacetin biosynthesis | false | false | false | false | false | false | false | false | false | false | false | false | false | false | false | false | false |
| \|PWY-5120\| | geranylgeranyl diphosphate biosynthesis | false | false | false | false | false | false | false | false | false | false | false | false | false | false | false | false | false |
| \|PWY-5122\| | geranyl diphosphate biosynthesis | false | false | false | false | false | false | false | false | false | false | false | false | false | false | false | false | false |
| \|PWY-5123\| | trans, trans-farnesyl diphosphate biosynthesis | false | false | false | false | false | false | false | false | false | false | false | false | false | false | false | false | false |
| \|PWY-5125\| | anthocyanin biosynthesis | false | false | false | false | false | false | false | false | false | false | false | false | false | false | false | false | false |
| \|PWY-5129\| | sphingolipid biosynthesis (plants) | false | false | false | false | false | false | false | false | false | false | false | false | false | false | false | false | false |
| \|PWY-5132\| | lupulone and humulone biosynthesis | false | false | false | false | false | false | false | false | false | false | false | false | false | false | false | false | false |
| \|PWY-5133\| | colupulone and cohumulone biosynthesis | false | false | false | false | false | false | false | false | false | false | false | false | false | false | false | false | false |
| \|PWY-5135\| | xanthohumol biosynthesis | false | false | false | false | false | false | false | false | false | false | false | false | false | false | false | false | false |
| \|PWY-5136\| | fatty acid β-oxidation II (plant peroxisome) | false | false | false | false | false | false | false | false | false | false | false | false | false | false | false | false | false |
| \|PWY-5137\| | fatty acid β-oxidation III (unsaturated, odd number) | false | false | false | false | false | false | false | false | false | false | false | false | true | false | true | false | true |
| \|PWY-5138\| | fatty acid β-oxidation IV (unsaturated, even number) | false | false | false | false | false | false | false | false | false | false | false | false | false | false | false | false | false |
| \|PWY-5139\| | pelargonidin conjugates biosynthesis | false | false | false | false | false | false | false | false | false | false | false | false | false | false | false | false | false |
| \|PWY-5140\| | cannabinoid biosynthesis | false | false | false | false | false | false | false | false | false | false | false | false | false | false | false | false | false |
| \|PWY-5142\| | acyl-[acyl-carrier protein] thioesterase pathway | false | false | false | false | false | false | false | false | false | false | false | false | false | false | false | false | false |
| \|PWY-5143\| | long-chain fatty acid activation | false | false | false | false | false | false | false | false | false | false | false | false | false | false | false | false | false |
| \|PWY-5147\| | oleate biosynthesis I (plants) | false | false | false | false | false | false | false | false | false | false | false | false | false | false | false | false | false |
| \|PWY-5148\| | acyl-CoA hydrolysis | false | false | false | false | false | false | false | false | false | false | false | false | false | false | false | false | false |

| | | | | | | | | | | | | | | | | | | |
|---|---|---|---|---|---|---|---|---|---|---|---|---|---|---|---|---|---|---|
| \|PWY-5151\| | L-tyrosine degradation II | false | false | false | false | false | false | false | false | false | false | false | false | false | false | false | false | false |
| \|PWY-5152\| | leucodelphinidin biosynthesis | false | false | false | false | false | false | false | false | false | false | false | false | false | false | false | false | false |
| \|PWY-5153\| | anthocyanin biosynthesis (delphinidin 3-O-glucoside) | false | false | false | false | false | false | false | false | false | false | false | false | false | false | false | false | false |
| \|PWY-5154\| | L-arginine biosynthesis III (via N-acetyl-L-citrulline) | false | false | false | false | false | false | false | false | false | false | false | false | false | false | false | false | false |
| \|PWY-5155\| | β-alanine biosynthesis III | false | false | false | false | false | false | false | false | false | false | false | false | false | false | false | false | false |
| \|PWY-5159\| | trans-4-hydroxy-L-proline degradation II | false | false | false | false | false | false | false | false | false | false | false | false | false | false | false | false | false |
| \|PWY-5160\| | rose anthocyanin biosynthesis I (via cyanidin 5-O-β-D-glucoside) | false | false | false | false | false | false | false | false | false | false | false | false | false | false | false | false | false |
| \|PWY-5161\| | 6'-deoxychalcone metabolism | false | false | false | false | false | false | false | false | false | false | false | false | false | false | false | false | false |
| \|PWY-5162\| | 2-hydroxypenta-2,4-dienoate degradation | false | false | false | false | false | false | false | false | false | false | false | false | false | false | false | false | false |
| \|PWY-5163\| | p-cumate degradation to 2-hydroxypentadienoate | false | false | false | false | false | false | false | false | false | false | false | false | false | false | false | false | false |
| \|PWY-5165\| | 4-toluenesulfonate degradation II | false | false | false | false | false | false | false | false | false | false | false | false | false | false | false | false | false |
| \|PWY-5168\| | ferulate and sinapate biosynthesis | false | false | false | false | false | false | false | false | false | false | false | false | false | false | false | false | false |
| \|PWY-5169\| | cyanuric acid degradation II | false | false | false | false | false | false | false | false | false | false | false | false | false | false | false | false | false |
| \|PWY-5170\| | melamine degradation | false | false | false | false | false | false | false | false | false | false | false | false | false | false | false | false | false |
| \|PWY-5171\| | N-cyclopropylmelamine degradation | false | false | false | false | false | false | false | false | false | false | false | false | false | false | false | false | false |
| \|PWY-5172\| | acetyl-CoA biosynthesis from citrate | false | false | false | false | false | false | false | false | false | false | false | false | false | false | false | false | false |
| \|PWY-5174\| | capsanthin and capsorubin biosynthesis | false | false | false | false | false | false | false | false | false | false | false | false | false | false | false | false | false |
| \|PWY-5175\| | lactucaxanthin biosynthesis | false | false | false | false | false | false | false | false | false | false | false | false | false | false | false | false | false |
| \|PWY-5176\| | coumarin biosynthesis (via 2-coumarate) | false | false | false | false | false | false | false | false | false | false | false | false | false | false | false | false | false |
| \|PWY-5189\| | tetrapyrrole biosynthesis II (from glycine) | false | false | false | false | false | false | false | false | false | false | false | false | false | false | false | false | false |
| \|PWY-5194\| | siroheme biosynthesis | false | false | false | false | false | false | false | false | false | false | false | false | false | false | false | false | false |
| \|PWY-5195\| | artemisinin and arteannuin B biosynthesis | false | false | false | false | false | false | false | false | false | false | false | false | false | false | false | false | false |
| \|PWY-5196\| | factor 430 biosynthesis | false | false | false | false | false | false | false | false | false | false | false | true | false | false | false | false | false |
| \|PWY-5197\| | lactate biosynthesis (archaea) | false | false | false | false | false | false | false | false | false | false | false | false | false | false | false | false | false |
| \|PWY-5198\| | factor 420 biosynthesis II (mycobacteria) | false | false | false | false | false | false | false | false | false | false | false | false | false | false | false | false | false |
| \|PWY-5199\| | factor 420 polyglutamylation | false | false | false | false | false | false | false | false | false | false | false | false | false | false | false | false | false |
| \|PWY-5203\| | soybean saponin I biosynthesis | false | false | false | false | false | false | false | false | false | false | false | false | false | false | false | false | false |
| \|PWY-5207\| | coenzyme B/coenzyme M regeneration I (methanophenazine-dependent) | false | false | false | false | false | false | false | false | false | false | false | false | false | false | false | false | false |
| \|PWY-5209\| | methyl-coenzyme M oxidation to CO2 | false | false | false | false | false | false | false | false | false | false | false | false | false | false | false | false | false |
| \|PWY-5247\| | methanogenesis from methylamine | false | false | false | false | false | false | false | false | false | false | false | true | false | false | false | false | false |
| \|PWY-5248\| | methanogenesis from dimethylamine | false | false | false | false | false | false | false | false | false | false | false | false | false | false | false | false | false |
| \|PWY-5250\| | methanogenesis from trimethylamine | false | false | false | false | false | false | false | false | false | false | false | false | false | false | false | false | false |
| \|PWY-5254\| | methanofuran biosynthesis | false | false | false | false | false | false | false | false | false | false | false | false | false | false | false | false | false |
| \|PWY-5258\| | methanogenesis from dimethylsulfide | false | false | false | false | false | false | false | false | false | false | false | false | false | false | false | false | false |
| \|PWY-5259\| | methanogenesis from methanethiol | false | false | false | false | false | false | false | false | false | false | false | false | false | false | false | false | false |
| \|PWY-5260\| | methanogenesis from methylthiopropanoate | false | false | false | false | false | false | false | false | false | false | false | false | false | false | false | false | false |
| \|PWY-5261\| | methanogenesis from tetramethylammonium | false | false | false | false | false | false | false | false | false | false | false | false | false | false | false | false | false |
| \|PWY-5267\| | glucosinolate activation | false | false | false | false | false | false | false | false | false | false | false | false | false | false | false | false | false |
| \|PWY-5268\| | salvianin biosynthesis | false | false | false | false | false | false | false | false | false | false | false | false | false | false | false | false | false |
| \|PWY-5269\| | cardiolipin biosynthesis II | false | false | false | false | false | false | false | false | false | false | false | false | false | false | false | false | false |
| \|PWY-5270\| | morphine biosynthesis | false | false | false | false | false | false | false | false | false | false | false | false | false | false | false | false | false |
| \|PWY-5271\| | abscisic acid degradation to phaseic acid | false | false | false | false | false | false | false | false | false | false | false | false | false | false | false | false | false |
| \|PWY-5272\| | abscisic acid degradation by glucosylation | false | false | false | false | false | false | false | false | false | false | false | false | false | false | false | false | false |
| \|PWY-5274\| | sulfide oxidation II (flavocytochrome c) | false | false | false | false | false | false | false | false | false | false | false | false | false | false | false | false | false |
| \|PWY-5276\| | sulfite oxidation I | false | false | false | false | false | false | false | false | false | false | false | false | false | false | false | false | false |
| \|PWY-5277\| | thiosulfate disproportionation I (thiol-dependent) | false | false | false | false | false | false | false | false | false | false | false | false | false | false | false | false | false |
| \|PWY-5278\| | sulfite oxidation III | false | false | false | false | false | false | false | false | false | false | false | false | false | false | false | false | false |
| \|PWY-5279\| | sulfite oxidation II | false | false | false | false | false | false | false | false | false | false | false | false | false | false | false | false | false |
| \|PWY-5280\| | L-lysine degradation IV | false | false | false | false | false | false | false | false | false | false | false | false | false | false | false | false | false |
| \|PWY-5283\| | L-lysine degradation V | false | false | false | false | false | false | false | false | false | false | false | false | false | false | false | false | false |
| \|PWY-5284\| | shisonin biosynthesis | false | false | false | false | false | false | false | false | false | false | false | false | false | false | false | false | false |
| \|PWY-5285\| | sulfide oxidation III (to sulfite) | false | false | false | false | false | false | false | false | false | false | false | false | false | false | false | false | false |
| \|PWY-5286\| | anthocyanidin sophoroside metabolism | false | false | false | false | false | false | false | false | false | false | false | false | false | false | false | false | false |
| \|PWY-5287\| | sanguinarine and macarpine biosynthesis | false | false | false | false | false | false | false | false | false | false | false | false | false | false | false | false | false |
| \|PWY-5288\| | astaxanthin biosynthesis (bacteria, fungi, algae) | false | false | false | false | false | false | false | false | false | false | false | false | false | false | false | false | false |
| \|PWY-5290\| | secologanin and strictosidine biosynthesis | false | false | false | false | false | false | false | false | false | false | false | false | false | false | false | false | false |
| \|PWY-5291\| | canthaxanthin biosynthesis | false | false | false | false | false | false | false | false | false | false | false | true | false | false | false | false | false |
| \|PWY-5292\| | vindoline, vindorosine and vinblastine biosynthesis | false | false | false | false | false | false | false | false | false | false | false | false | false | false | false | false | false |
| \|PWY-5295\| | ternatin C5 biosynthesis | false | false | false | false | false | false | false | false | false | false | false | false | false | false | false | false | false |
| \|PWY-5296\| | thiosulfate oxidation III (multienzyme complex) | false | false | false | false | false | false | false | false | false | false | false | false | false | false | false | false | false |
| \|PWY-5297\| | siroheme amide biosynthesis | false | false | false | false | false | false | false | false | false | false | false | false | false | false | false | false | false |
| \|PWY-5298\| | L-lysine degradation VI | false | false | false | false | false | false | false | false | false | false | false | false | false | false | false | false | false |
| \|PWY-5301\| | ajmaline and sarpagine biosynthesis | false | false | false | false | false | false | false | false | false | false | false | false | false | false | false | false | false |
| \|PWY-5302\| | sulfur disproportionation II (aerobic) | false | false | false | false | false | false | false | false | false | false | false | false | false | false | false | false | false |
| \|PWY-5303\| | thiosulfate oxidation II (via tetrathionate) | false | false | false | false | false | false | false | false | false | false | false | false | false | false | false | false | false |
| \|PWY-5305\| | bixin biosynthesis | false | false | false | false | false | false | false | false | false | false | false | false | false | false | false | false | false |
| \|PWY-5307\| | gentiodelphin biosynthesis | false | false | false | false | false | false | false | false | false | false | false | false | false | false | false | false | false |
| \|PWY-5311\| | L-lysine degradation VII | false | false | false | false | false | false | false | false | false | false | false | false | false | false | false | false | false |
| \|PWY-5314\| | L-lysine degradation VIII | false | false | false | false | false | false | false | false | false | false | false | false | false | false | false | false | false |
| \|PWY-5315\| | 1-methylpyrrolinium biosynthesis | false | false | false | false | false | false | false | false | false | false | false | false | false | false | false | false | false |
| \|PWY-5316\| | nicotine biosynthesis | false | false | false | false | false | false | false | false | false | false | false | false | false | false | false | false | false |
| \|PWY-5317\| | tropane alkaloids biosynthesis | false | false | false | false | false | false | false | false | false | false | false | false | false | false | false | false | false |
| \|PWY-5318\| | calystegine biosynthesis | false | false | false | false | false | false | false | false | false | false | false | false | false | false | false | false | false |
| \|PWY-5319\| | coumarin metabolism (to melilotic acid) | false | false | false | false | false | false | false | false | false | false | false | false | false | false | false | false | false |
| \|PWY-5320\| | kaempferol glycoside biosynthesis (Arabidopsis) | false | false | false | false | false | false | false | false | false | false | false | false | false | false | false | false | false |
| \|PWY-5321\| | quercetin glycoside biosynthesis (Arabidopsis) | false | false | false | false | false | false | false | false | false | false | false | false | false | false | false | false | false |
| \|PWY-5324\| | L-lysine degradation IX | false | false | false | false | false | false | false | false | false | false | false | false | false | false | false | false | false |
| \|PWY-5326\| | sulfite oxidation IV (sulfite oxidase) | false | false | false | false | false | false | false | false | false | false | false | false | false | false | false | false | false |
| \|PWY-5329\| | L-cysteine degradation III | false | false | false | false | false | false | false | false | false | false | false | false | false | false | false | false | false |
| \|PWY-5331\| | taurine biosynthesis I | false | false | false | false | false | false | false | false | false | false | false | false | false | false | false | false | false |
| \|PWY-5336\| | carbon disulfide oxidation II (aerobic) | false | false | false | false | false | false | false | false | false | false | false | false | false | false | false | false | false |
| \|PWY-5337\| | stachyose biosynthesis | false | false | false | false | false | false | false | false | false | false | false | false | false | false | false | false | false |
| \|PWY-5338\| | galactosylcyclitol biosynthesis | false | false | false | false | false | false | false | false | false | false | false | false | false | false | false | false | false |
| \|PWY-5339\| | chalcone 2'-O-glucoside biosynthesis | false | false | false | false | false | false | false | false | false | false | false | false | false | false | false | false | false |
| \|PWY-5340\| | sulfate activation for sulfonation | false | false | false | false | false | false | false | false | false | false | false | false | false | false | false | false | false |
| \|PWY-5342\| | ajugose biosynthesis I (galactinol-dependent) | false | false | false | false | false | false | false | false | false | false | false | false | false | false | false | false | false |
| \|PWY-5343\| | ajugose biosynthesis II (galactinol-independent) | false | false | false | false | false | false | false | false | false | false | false | false | false | false | false | false | false |
| \|PWY-5344\| | L-homocysteine biosynthesis | false | false | false | false | false | false | false | false | false | false | false | false | false | true | false | false | true |
| \|PWY-5348\| | kaempferol triglucoside biosynthesis | false | false | false | false | false | false | false | false | false | false | false | false | false | false | false | false | false |
| \|PWY-5349\| | esculetin biosynthesis | false | false | false | false | false | false | false | false | false | false | false | false | false | false | false | false | false |
| \|PWY-5350\| | thiosulfate disproportionation IV (rhodanese) | false | false | false | false | false | false | false | false | false | false | false | false | false | false | false | false | false |
| \|PWY-5352\| | thiosulfate disproportionation II (cytochrome) | false | false | false | false | false | false | false | false | false | false | false | false | false | false | false | false | false |
| \|PWY-5353\| | arachidonate biosynthesis I (6-desaturase, lower eukaryotes) | false | false | false | false | false | false | false | false | false | false | false | false | false | false | false | false | false |
| \|PWY-5355\| | nitroethane degradation | false | false | false | false | false | false | false | false | false | false | false | false | false | false | false | false | false |
| \|PWY-5358\| | tetrathionate reduction I (to thiosulfate) | false | false | false | false | false | false | false | false | false | false | false | false | false | false | false | false | false |
| \|PWY-5359\| | tetrathionate reductiuon II (to trithionate) | false | false | false | false | false | false | false | false | false | false | false | false | false | false | false | false | false |
| \|PWY-5361\| | (5Z)-icosenoate biosynthesis | false | false | false | false | false | false | false | false | false | false | false | false | false | false | false | false | false |
| \|PWY-5362\| | sapienate biosynthesis | false | false | false | false | false | false | false | false | false | false | false | false | false | false | false | false | false |
| \|PWY-5363\| | chrysin biosynthesis | false | false | false | false | false | false | false | false | false | false | false | false | false | false | false | false | false |
| \|PWY-5365\| | linear furanocoumarin biosynthesis | false | false | false | false | false | false | false | false | false | false | false | false | false | false | false | false | false |
| \|PWY-5366\| | palmitoleate biosynthesis II (plants and bacteria) | false | false | false | false | false | false | false | false | false | false | false | false | false | false | false | false | false |
| \|PWY-5367\| | petroselinate biosynthesis | false | false | false | false | false | false | false | false | false | false | false | false | false | false | false | false | false |
| \|PWY-5368\| | dimorphecolate biosynthesis | false | false | false | false | false | false | false | false | false | false | false | false | false | false | false | false | false |
| \|PWY-5370\| | carbon tetrachloride degradation I | false | false | false | false | false | false | false | false | false | false | false | false | false | false | false | false | false |
| \|PWY-5373\| | calendate biosynthesis | false | false | false | false | false | false | false | false | false | false | false | false | false | false | false | false | false |
| \|PWY-5374\| | punicate biosynthesis | false | false | false | false | false | false | false | false | false | false | false | false | false | false | false | false | false |
| \|PWY-5375\| | α-eleostearate biosynthesis | false | false | false | false | false | false | false | false | false | false | false | false | false | false | false | false | false |
| \|PWY-5377\| | α-amyrin biosynthesis | false | false | false | false | false | false | false | false | false | false | false | false | false | false | false | false | false |
| \|PWY-5379\| | B series fagopyritols biosynthesis | false | false | false | false | false | false | false | false | false | false | false | false | false | false | true | true | true |
| \|PWY-5380\| | A series fagopyritols biosynthesis | false | false | false | false | false | false | false | false | false | false | false | false | false | false | false | false | false |
| \|PWY-5381\| | NAD salvage (plants) | false | false | false | false | false | false | false | false | false | false | false | false | false | false | false | false | false |
| \|PWY-5384\| | sucrose degradation IV (sucrose phosphorylase) | false | false | false | false | false | false | false | false | false | false | false | false | false | false | false | false | false |
| \|PWY-5386-1\| | methylglyoxal degradation VIII | false | false | false | false | false | false | false | false | false | false | false | false | false | false | false | false | false |
| \|PWY-5386\| | methylglyoxal degradation I | false | false | false | false | false | false | false | false | false | false | false | false | false | false | false | false | false |
| \|PWY-5388\| | N-glucosylnicotinate metabolism | false | false | false | false | false | false | false | false | false | false | false | false | false | false | false | false | false |
| \|PWY-5389\| | 3-methylthiopropanoate biosynthesis | false | false | false | false | false | false | false | false | false | false | false | false | false | false | false | false | false |
| \|PWY-5390\| | rutin biosynthesis | false | false | false | false | false | false | false | false | false | false | false | false | false | false | false | false | false |
| \|PWY-5391\| | syringetin biosynthesis | false | false | false | false | false | false | false | false | false | false | false | false | false | false | false | false | false |
| \|PWY-5392\| | reductive TCA cycle II | false | false | false | false | false | false | false | false | false | false | false | false | false | false | false | false | false |
| \|PWY-5393\| | raspberry ketone biosynthesis | false | false | false | false | false | false | false | false | false | false | false | false | false | false | false | false | false |
| \|PWY-5394\| | betalamic acid biosynthesis | false | false | false | false | false | false | false | false | false | false | false | false | false | false | false | false | false |
| \|PWY-5397\| | crocetin biosynthesis | false | false | false | false | false | false | false | false | false | false | false | false | false | false | false | false | false |
| \|PWY-5398\| | crocetin esters biosynthesis | false | false | false | false | false | false | false | false | false | false | false | false | false | false | false | false | false |
| \|PWY-5399\| | betacyanin biosynthesis | false | false | false | false | false | false | false | false | false | false | false | false | false | false | false | false | false |
| \|PWY-5400\| | amaranthin biosynthesis | false | false | false | false | false | false | false | false | false | false | false | false | false | false | false | false | false |
| \|PWY-5403\| | betaxanthin biosynthesis (via dopamine) | false | false | false | false | false | false | false | false | false | false | false | false | true | true | true | false | true |
| \|PWY-5404\| | betaxanthin biosynthesis (via dopaxanthin) | false | false | false | false | false | false | false | false | false | false | false | false | false | false | false | false | false |
| \|PWY-5406\| | divinyl ether biosynthesis I | false | false | false | false | false | false | false | false | false | false | false | false | false | false | false | false | false |
| \|PWY-5407\| | 9-lipoxygenase and 9-allene oxide synthase pathway | false | false | false | false | false | false | false | false | false | false | false | false | false | false | false | false | false |
| \|PWY-5408\| | 9-lipoxygenase and 9-hydroperoxide lyase pathway | false | false | false | false | false | false | false | false | false | false | false | false | false | false | false | false | false |
| \|PWY-5409\| | divinyl ether biosynthesis II | false | false | false | false | false | false | false | false | false | false | false | false | false | false | false | false | false |
| \|PWY-5410\| | traumatin and (Z)-3-hexen-1-yl acetate biosynthesis | false | false | false | false | false | false | false | false | false | false | false | false | false | false | false | false | false |
| \|PWY-5411\| | abietic acid biosynthesis | false | false | false | false | false | false | false | false | false | false | false | false | false | false | false | false | false |
| \|PWY-5412\| | levopimaric acid biosynthesis | false | false | false | false | false | false | false | false | false | false | false | false | false | false | false | false | false |
| \|PWY-5413\| | neoabietic acid biosynthesis | false | false | false | false | false | false | false | false | false | false | false | false | false | false | false | false | false |
| \|PWY-5414\| | palustric acid biosynthesis | false | false | false | false | false | false | false | false | false | false | false | false | false | false | false | false | false |
| \|PWY-5418\| | phenol degradation I (aerobic) | false | false | false | false | false | false | false | false | false | false | false | false | false | false | false | false | false |
| \|PWY-5419\| | catechol degradation to 2-hydroxypentadienoate II | false | false | false | false | false | false | false | false | false | false | false | false | false | false | false | false | false |
| \|PWY-5421\| | dehydroabietic acid biosynthesis | false | false | false | false | false | false | false | false | false | false | false | false | false | false | false | false | false |
| \|PWY-5422\| | isopimaric acid biosynthesis | false | false | false | false | false | false | false | false | false | false | false | false | false | false | false | false | false |
| \|PWY-5423\| | oleoresin monoterpene volatiles biosynthesis | false | false | false | false | false | false | false | false | false | false | false | false | false | false | false | false | false |
| \|PWY-5425\| | oleoresin sesquiterpene volatiles biosynthesis | false | false | false | false | false | false | false | false | false | false | false | false | false | false | false | false | false |
| \|PWY-5426\| | betaxanthin biosynthesis | false | false | false | false | false | false | false | false | false | false | false | false | false | false | false | false | false |
| \|PWY-5427\| | naphthalene degradation (aerobic) | false | false | false | false | false | false | false | false | false | false | false | false | false | false | false | false | false |
| \|PWY-5428\| | 1,3-dimethylbenzene degradation to 3-methylbenzoate | false | false | false | false | false | false | false | false | false | false | false | false | false | false | false | false | false |
| \|PWY-5429\| | 1,4-dimethylbenzene degradation to 4-methylbenzoate | false | false | false | false | false | false | false | false | false | false | false | false | false | false | false | false | false |
| \|PWY-5434\| | (3E)-4,8-dimethylnona-1,3,7-triene biosynthesis I | false | false | false | false | false | false | false | false | false | false | false | false | false | false | false | false | false |
| \|PWY-5437\| | L-threonine degradation I | false | false | false | false | false | false | false | false | false | false | false | false | false | false | false | false | false |
| \|PWY-5439\| | betacyanin biosynthesis (via dopamine) | false | false | false | false | false | false | false | false | false | false | false | false | false | false | false | false | false |
| \|PWY-5441\| | S-methyl-L-methionine cycle | false | false | false | false | false | false | false | false | false | false | false | false | false | false | false | false | false |
| \|PWY-5443\| | aminopropanol phosphate biosynthesis I | false | false | false | false | false | false | false | false | false | false | false | false | false | false | false | false | false |
| \|PWY-5450\| | benzene degradation I (aerobic) | false | false | false | false | false | false | false | false | false | false | false | false | false | false | false | false | false |
| \|PWY-5451\| | acetone degradation I (to methylglyoxal) | false | false | false | false | false | false | false | false | false | false | false | false | false | false | false | false | false |
| \|PWY-5453\| | methylglyoxal degradation III | false | false | false | false | false | false | false | false | false | false | false | false | false | false | false | false | false |
| \|PWY-5456\| | methylglyoxal degradation VII | false | false | false | false | false | false | false | false | false | false | false | false | false | false | false | false | false |
| \|PWY-5458\| | methylglyoxal degradation V | false | false | false | false | false | false | false | false | false | false | false | false | false | false | false | false | false |
| \|PWY-5461\| | betanidin degradation | false | false | false | false | false | false | false | false | false | false | false | false | false | false | false | false | false |
| \|PWY-5462\| | methylglyoxal degradation II | false | false | false | false | false | false | false | false | false | false | false | false | false | false | false | false | false |
| \|PWY-5466\| | matairesinol biosynthesis | false | false | false | false | false | false | false | false | false | false | false | true | false | false | false | false | false |
| \|PWY-5467\| | gramine biosynthesis | false | false | false | false | false | false | false | false | false | false | false | false | false | false | false | false | false |
| \|PWY-5468\| | lupanine biosynthesis | false | false | false | false | false | false | false | false | false | false | false | false | false | false | false | false | false |
| \|PWY-5469\| | sesamin biosynthesis | false | false | false | false | false | false | false | false | false | false | false | false | false | false | false | false | false |
| \|PWY-5470\| | palmatine biosynthesis | false | false | false | false | false | false | false | false | false | false | false | false | false | false | false | false | false |
| \|PWY-5472\| | bisbenzylisoquinoline alkaloid biosynthesis | false | false | false | false | false | false | false | false | false | false | false | false | false | false | false | false | false |
| \|PWY-5473\| | hydroxycinnamic acid serotonin amides biosynthesis | false | false | false | false | false | false | false | false | false | false | false | false | false | false | false | false | false |
| \|PWY-5474\| | hydroxycinnamic acid tyramine amides biosynthesis | false | false | false | false | false | false | false | false | false | false | false | false | false | false | false | false | false |
| \|PWY-5475\| | pentagalloylglucose biosynthesis | false | false | false | false | false | false | false | false | false | false | false | false | false | false | false | false | false |
| \|PWY-5476\| | cornusiin E biosynthesis | false | false | false | false | false | false | false | false | false | false | false | false | false | false | false | false | false |
| \|PWY-5477\| | gallotannin biosynthesis | false | false | false | false | false | false | false | false | false | false | false | false | false | false | false | false | false |
| \|PWY-5479\| | 6-methoxypodophyllotoxin biosynthesis | false | false | false | false | false | false | false | false | false | false | false | false | false | false | false | false | false |
| \|PWY-5480\| | pyruvate fermentation to ethanol I | false | false | false | false | false | false | false | false | false | false | false | false | false | false | false | false | false |
| \|PWY-5486\| | pyruvate fermentation to ethanol II | false | false | false | false | false | false | false | false | false | false | false | false | false | false | false | false | false |
| \|PWY-5487\| | 4-nitrophenol degradation I | false | false | false | false | false | false | false | false | false | false | false | false | false | false | false | false | false |
| \|PWY-5488\| | 4-nitrophenol degradation II | false | false | false | false | false | false | false | false | false | false | false | false | false | false | false | false | false |
| \|PWY-5489\| | methyl parathion degradation | false | false | false | false | false | false | false | false | false | false | false | false | false | false | false | false | false |
| \|PWY-5490\| | paraoxon degradation | false | false | false | false | false | false | false | false | false | false | false | false | false | false | false | false | false |
| \|PWY-5491\| | diethylphosphate degradation | false | false | false | false | false | false | false | false | false | false | false | false | false | false | false | false | false |
| \|PWY-5493\| | reductive monocarboxylic acid cycle | false | false | false | false | false | false | false | false | false | false | false | false | false | false | false | false | false |
| \|PWY-5494\| | (R)-lactate fermentation to propanoate II (acrylate pathway) | false | false | false | false | false | false | false | false | false | false | false | false | false | false | false | false | false |
| \|PWY-5497\| | purine nucleobases degradation II (anaerobic) | false | false | false | false | false | false | false | false | false | false | false | false | false | false | false | false | false |
| \|PWY-5499\| | vitamin B6 degradation I | false | false | false | false | false | false | false | false | false | false | false | false | false | false | false | false | false |
| \|PWY-5506\| | methanol oxidation to formaldehyde IV | false | false | false | false | false | false | false | false | false | false | false | false | false | false | false | false | false |
| \|PWY-5509\| | adenosylcobalamin biosynthesis from adenosylcobinamide-GDP I | false | false | false | false | false | false | false | false | false | false | false | false | false | false | false | false | false |
| \|PWY-5515\| | L-arabinose degradation II | false | false | false | false | false | false | false | false | false | false | false | false | false | false | false | false | false |
| \|PWY-5517\| | L-arabinose degradation III | false | false | false | false | false | false | false | false | false | false | false | false | false | false | false | false | false |
| \|PWY-5519\| | D-arabinose degradation III | false | false | false | false | false | false | false | false | false | false | false | false | false | false | false | false | false |
| \|PWY-5521\| | L-ascorbate biosynthesis III (D-sorbitol pathway) | false | false | false | false | false | false | false | false | false | false | false | false | false | false | false | false | false |
| \|PWY-5523\| | 5,6-dimethylbenzimidazole biosynthesis I (aerobic) | false | false | false | false | false | false | false | false | false | false | false | false | false | false | false | false | false |
| \|PWY-5525\| | D-glucuronate degradation I | false | false | false | false | false | false | false | false | false | false | false | false | false | false | false | false | false |
| \|PWY-5526\| | bacteriochlorophyll a biosynthesis | false | false | false | false | false | false | false | false | false | false | false | false | false | false | false | false | false |
| \|PWY-5530\| | sorbitol biosynthesis II | false | false | false | false | false | false | false | false | false | false | false | false | false | false | false | false | false |
| \|PWY-5531\| | 3,8-divinyl-chlorophyllide a biosynthesis II (anaerobic) | false | false | false | false | false | false | false | false | false | false | false | false | false | false | false | false | false |
| \|PWY-5532\| | nucleoside and nucleotide degradation (archaea) | false | false | false | false | false | false | false | false | false | false | false | false | false | false | false | false | false |
| \|PWY-5533\| | acetone degradation II (to acetoacetate) | false | false | false | false | false | false | false | false | false | false | false | false | false | false | false | false | false |
| \|PWY-5534\| | propene degradation | false | false | false | false | false | false | false | false | false | false | false | false | false | false | false | false | false |
| \|PWY-5536\| | acetate formation from acetyl-CoA (succinate) | false | false | false | false | false | false | false | false | false | false | false | false | false | false | false | false | false |
| \|PWY-5629\| | isopenicillin N biosynthesis | false | false | false | false | false | false | false | false | false | false | false | false | false | false | false | false | false |
| \|PWY-5630\| | penicillin K biosynthesis | false | false | false | false | false | false | false | false | false | false | false | false | false | false | false | false | false |
| \|PWY-5631\| | deacetylcephalosporin C biosynthesis | false | false | false | false | false | false | false | false | false | false | false | false | false | false | false | false | false |
| \|PWY-5632\| | cephalosporin C biosynthesis | false | false | false | false | false | false | false | false | false | false | false | false | false | false | false | false | false |
| \|PWY-5633\| | cephamycin C biosynthesis | false | false | false | false | false | false | false | false | false | false | false | false | false | false | false | false | false |
| \|PWY-5636\| | 2-nitrophenol degradation | false | false | false | false | false | false | false | false | false | false | false | false | false | false | false | false | false |
| \|PWY-5637\| | nitrobenzene degradation I | false | false | false | false | false | false | false | false | false | false | false | false | false | false | false | false | false |
| \|PWY-5640\| | nitrobenzene degradation II | false | false | false | false | false | false | false | false | false | false | false | false | false | false | false | false | false |
| \|PWY-5641\| | 2-nitrotoluene degradation | false | false | false | false | false | false | false | false | false | false | false | false | false | false | false | false | false |
| \|PWY-5642\| | 2,4-dinitrotoluene degradation | false | false | false | false | false | false | false | false | false | false | false | false | false | false | false | false | false |
| \|PWY-5643\| | 2,6-dinitrotoluene degradation | false | false | false | false | false | false | false | false | false | false | false | false | false | false | false | false | false |
| \|PWY-5644\| | 4-nitrotoluene degradation II | false | false | false | false | false | false | false | false | false | false | false | false | false | false | false | false | false |
| \|PWY-5645\| | 4-chloronitrobenzene degradation | false | false | false | false | false | false | false | false | false | false | false | false | false | false | false | false | false |
| \|PWY-5648\| | 2-nitrobenzoate degradation II | false | false | false | false | false | false | false | false | false | false | false | false | false | false | false | false | false |
| \|PWY-5651\| | L-tryptophan degradation to 2-amino-3-carboxymuconate semialdehyde | false | false | false | false | false | false | false | false | false | false | false | false | false | false | false | false | false |
| \|PWY-5652\| | 2-amino-3-carboxymuconate semialdehyde degradation to glutaryl-CoA | false | false | false | false | false | false | false | false | false | false | false | false | false | false | false | false | false |
| \|PWY-5654\| | 2-amino-3-carboxymuconate semialdehyde degradation to 2-hydroxypentadienoate | false | false | false | false | false | false | false | false | false | false | false | false | false | false | false | false | false |
| \|PWY-5656\| | mannosylglycerate biosynthesis I | false | false | false | false | false | false | false | false | false | false | false | false | false | false | false | false | false |
| \|PWY-5658\| | mannosylglycerate biosynthesis II | false | false | false | false | false | false | false | false | false | false | false | false | false | false | false | false | false |
| \|PWY-5660\| | taxol biosynthesis | false | false | false | false | false | false | false | false | false | false | false | false | false | false | false | false | false |
| \|PWY-5661\| | GDP-α-D-glucose biosynthesis | false | false | false | false | false | false | false | false | false | false | false | false | false | false | false | false | false |
| \|PWY-5662\| | glucosylglycerate biosynthesis I | false | false | false | false | false | false | false | false | false | false | false | false | false | false | false | false | false |
| \|PWY-5663\| | erythro-tetrahydrobiopterin biosynthesis I | false | false | false | false | false | false | false | false | false | false | false | false | false | false | false | false | false |
| \|PWY-5665\| | vanillin biosynthesis I | false | false | false | false | false | false | false | false | false | false | false | false | false | false | false | false | false |
| \|PWY-5666\| | α-solanine/α-chaconine biosynthesis | false | false | false | false | false | false | false | false | false | false | false | false | false | false | false | false | false |
| \|PWY-5667\| | CDP-diacylglycerol biosynthesis I | false | false | false | false | false | false | false | false | false | false | false | false | false | false | false | false | false |
| \|PWY-5668\| | cardiolipin biosynthesis I | false | false | false | false | false | false | false | false | false | false | false | false | false | false | false | false | false |
| \|PWY-5669\| | phosphatidylserine and phosphatidylethanolamine biosynthesis I | false | false | false | false | false | false | false | false | false | false | false | false | false | false | false | false | false |
| \|PWY-5670\| | epoxysqualene biosynthesis | false | false | false | false | false | false | false | false | false | false | false | false | false | false | false | false | false |
| \|PWY-5672\| | ginsenosides biosynthesis | false | false | false | false | false | false | false | false | false | false | false | false | false | false | false | false | false |
| \|PWY-5674\| | nitrate reduction IV (dissimilatory) | false | false | false | false | false | false | false | false | false | false | false | false | false | false | false | false | false |
| \|PWY-5675\| | nitrate reduction V (assimilatory) | false | false | false | false | false | false | false | false | false | false | false | false | false | false | false | false | false |
| \|PWY-5677\| | succinate fermentation to butanoate | false | false | false | false | false | false | false | false | false | false | false | false | false | false | false | false | false |
| \|PWY-5679\| | clavulanate biosynthesis | false | false | false | false | false | false | false | false | false | false | false | false | false | false | false | false | false |
| \|PWY-5686\| | UMP biosynthesis I | false | false | false | false | false | false | false | false | false | false | false | false | false | false | false | false | false |
| \|PWY-5691\| | urate conversion to allantoin I | false | false | false | false | false | false | false | false | false | false | false | false | false | false | false | false | false |
| \|PWY-5695\| | inosine 5'-phosphate degradation | false | false | false | false | false | false | false | false | false | false | false | false | false | false | false | false | false |
| \|PWY-5697\| | allantoin degradation to ureidoglycolate I (urea producing) | false | false | false | false | false | false | false | false | false | false | false | false | false | false | false | false | false |
| \|PWY-5698\| | allantoin degradation to ureidoglycolate II (ammonia producing) | false | false | false | false | false | false | false | false | false | false | false | false | false | false | false | false | false |
| \|PWY-5701\| | shikonin biosynthesis | false | false | false | false | false | false | false | false | false | false | false | false | false | false | false | false | false |
| \|PWY-5703\| | urea degradation I | false | false | false | false | false | false | false | false | false | false | false | false | false | false | false | false | false |
| \|PWY-5704\| | urea degradation II | false | false | false | false | false | false | false | false | false | false | false | false | false | false | false | false | false |
| \|PWY-5706\| | alliin metabolism | false | false | false | false | false | false | false | false | false | false | false | false | true | true | true | true | true |
| \|PWY-5707\| | propanethial S-oxide biosynthesis | false | false | false | false | false | false | false | false | false | false | false | false | false | false | false | false | false |
| \|PWY-5708\| | ethiin metabolism | false | false | false | false | false | false | false | false | false | false | false | false | false | false | false | false | false |
| \|PWY-5710\| | capsaicin biosynthesis | false | false | false | false | false | false | false | false | false | false | false | false | false | false | false | false | false |
| \|PWY-5723\| | Rubisco shunt | false | false | false | false | false | false | false | false | false | false | false | false | false | false | false | false | false |
| \|PWY-5725\| | farnesene biosynthesis | false | false | false | false | false | false | false | false | false | false | false | false | false | false | false | false | false |
| \|PWY-5726\| | deethylsimazine degradation | false | false | false | false | false | false | false | false | false | false | false | false | false | false | false | false | false |
| \|PWY-5727\| | atrazine degradation II | false | false | false | false | false | false | false | false | false | false | false | false | false | false | false | false | false |
| \|PWY-5729\| | vestitol and sativan biosynthesis | false | false | false | false | false | false | false | false | false | false | false | false | false | false | false | false | false |
| \|PWY-5731\| | atrazine degradation III | false | false | false | false | false | false | false | false | false | false | false | false | false | false | false | false | false |
| \|PWY-5733\| | germacrene biosynthesis | false | false | false | false | false | false | false | false | false | false | false | false | false | false | false | false | false |
| \|PWY-5736\| | isopropylamine degradation | false | false | false | false | false | false | false | false | false | false | false | false | false | false | false | false | false |
| \|PWY-5737\| | (5R)-carbapenem carboxylate biosynthesis | false | false | false | false | false | false | false | false | false | false | false | false | false | false | false | false | false |
| \|PWY-5738\| | GDP-6-deoxy-D-talose biosynthesis | false | false | false | false | false | false | false | false | false | false | false | false | false | false | false | false | false |
| \|PWY-5739\| | GDP-D-perosamine biosynthesis | false | false | false | false | false | false | false | false | false | false | false | false | false | false | false | false | false |
| \|PWY-5740\| | GDP-L-colitose biosynthesis | false | false | false | false | false | false | false | false | false | false | false | true | false | false | false | false | false |
| \|PWY-5741\| | ethylmalonyl-CoA pathway | false | false | false | false | false | false | false | false | false | false | false | false | true | true | true | true | true |
| \|PWY-5742\| | L-arginine degradation IX (arginine:pyruvate transaminase pathway) | false | false | false | false | false | false | false | false | false | false | false | false | false | false | false | false | false |
| \|PWY-5743\| | 3-hydroxypropanoate cycle | false | false | false | false | false | false | false | false | false | false | false | false | false | false | false | false | false |
| \|PWY-5744\| | glyoxylate assimilation | false | false | false | false | false | false | false | false | false | false | false | false | false | false | false | false | false |
| \|PWY-5747\| | 2-methylcitrate cycle II | false | false | false | false | false | false | false | false | false | false | false | false | false | false | false | false | false |
| \|PWY-5748\| | γ-coniciene and coniine biosynthesis | false | false | false | false | false | false | false | false | false | false | false | false | false | false | false | false | false |
| \|PWY-5749\| | itaconate degradation | false | false | false | false | false | false | false | false | false | false | false | false | false | false | false | false | false |
| \|PWY-5750\| | itaconate biosynthesis I | false | false | false | false | false | false | false | false | false | false | false | false | false | false | false | false | false |
| \|PWY-5751\| | phenylethanol biosynthesis | false | false | false | false | false | false | false | false | false | false | false | false | false | false | false | false | false |
| \|PWY-5752\| | piperine biosynthesis | false | false | false | false | false | false | false | false | false | false | false | false | false | false | false | false | false |
| \|PWY-5754\| | 4-hydroxybenzoate biosynthesis I (eukaryotes) | false | false | false | false | false | false | false | false | false | false | false | false | false | false | false | false | false |
| \|PWY-5755\| | 4-hydroxybenzoate biosynthesis II (bacteria) | false | false | false | false | false | false | false | false | false | false | false | false | false | false | false | false | false |
| \|PWY-5756\| | saponin biosynthesis II | false | false | false | false | false | false | false | false | false | false | false | false | false | false | false | false | false |
| \|PWY-5757\| | fosfomycin biosynthesis | false | false | false | false | false | false | false | false | false | false | false | false | false | false | false | false | false |
| \|PWY-5759\| | saponin biosynthesis III | false | false | false | false | false | false | false | false | false | false | false | false | false | false | false | false | false |
| \|PWY-5760\| | β-alanine biosynthesis IV | false | false | false | false | false | false | false | false | false | false | false | false | false | false | false | false | false |
| \|PWY-5765\| | 1,3,5-trimethoxybenzene biosynthesis | false | false | false | false | false | false | false | false | false | false | false | false | false | false | false | false | false |
| \|PWY-5768\| | pyruvate fermentation to acetate VIII | false | false | false | false | false | false | false | false | false | false | false | false | false | false | false | false | false |
| \|PWY-5770\| | phenazine-1-carboxylate biosynthesis | false | false | false | false | false | false | false | false | false | false | false | false | false | false | false | false | false |
| \|PWY-5773\| | gossypol biosynthesis | false | false | false | false | false | false | false | false | false | false | false | false | false | false | false | false | false |
| \|PWY-5774\| | saponin biosynthesis IV | false | false | false | false | false | false | false | false | false | false | false | false | false | false | false | false | false |
| \|PWY-5775\| | rotenoid biosynthesis I | false | false | false | false | false | false | false | false | false | false | false | false | false | false | false | false | false |
| \|PWY-5776\| | 2-hydroxyphenazine biosynthesis | false | false | false | false | false | false | false | false | false | false | false | false | false | false | false | false | false |
| \|PWY-5780\| | hypericin biosynthesis | false | false | false | false | false | false | false | false | false | false | false | false | false | false | false | false | false |
| \|PWY-5782\| | 2-keto-L-gulonate biosynthesis | false | false | false | false | false | false | false | false | false | false | false | false | false | false | false | false | false |
| \|PWY-5783\| | octaprenyl diphosphate biosynthesis | false | false | false | false | false | false | false | false | false | false | false | false | false | false | false | false | false |
| \|PWY-5784\| | indole-3-acetate inactivation VIII | false | false | false | false | false | false | false | false | false | false | false | false | false | false | false | false | false |
| \|PWY-5787\| | oligomeric urushiol biosynthesis | false | false | false | false | false | false | false | false | false | false | false | false | false | false | false | false | false |
| \|PWY-5788\| | indole-3-acetate inactivation V | false | false | false | false | false | false | false | false | false | false | false | false | false | false | false | false | false |
| \|PWY-5789\| | 3-hydroxypropanoate/4-hydroxybutanate cycle | false | false | false | false | false | false | false | false | false | false | false | false | false | false | false | false | false |
| \|PWY-5793\| | maysin biosynthesis | false | false | false | false | false | false | false | false | false | false | false | false | false | false | false | false | false |
| \|PWY-5794\| | malonate degradation I (biotin-independent) | false | false | false | false | false | false | false | false | false | false | false | false | false | false | false | false | false |
| \|PWY-5795\| | juglone biosynthesis | false | false | false | false | false | false | false | false | false | false | false | false | false | false | false | false | false |
| \|PWY-5796\| | malonate decarboxylase activation | false | false | false | false | false | false | false | false | false | false | false | false | false | false | false | false | false |
| \|PWY-5797\| | indole-3-acetate inactivation VI | false | false | false | false | false | false | false | false | false | false | false | false | false | false | false | false | false |
| \|PWY-5800\| | xylan biosynthesis | false | false | false | false | false | false | false | false | false | false | false | false | false | false | false | false | false |
| \|PWY-5801\| | lawsone biosynthesis | false | false | false | false | false | false | false | false | false | false | false | false | false | false | false | false | false |
| \|PWY-5802\| | alizarin biosynthesis | false | false | false | false | false | false | false | false | false | false | false | false | false | false | false | false | false |
| \|PWY-5805\| | nonaprenyl diphosphate biosynthesis I | false | false | false | false | false | false | false | false | false | false | false | false | false | false | false | false | false |
| \|PWY-5806\| | all-trans-decaprenyl diphosphate biosynthesis | false | false | false | false | false | false | false | false | false | false | false | false | false | false | false | false | false |
| \|PWY-5807\| | heptaprenyl diphosphate biosynthesis | false | false | false | false | false | false | false | false | false | false | false | false | false | false | false | false | false |
| \|PWY-5808\| | hyperforin and adhyperforin biosynthesis | false | false | false | false | false | false | false | false | false | false | false | false | false | false | false | false | false |
| \|PWY-5811\| | indole-3-acetate inactivation VII | false | false | false | false | false | false | false | false | false | false | false | false | false | false | false | false | false |
| \|PWY-5813\| | bornyl diphosphate biosynthesis | false | false | false | false | false | false | false | false | false | false | false | false | false | false | false | false | false |
| \|PWY-5815\| | rubber biosynthesis | false | false | false | false | false | false | false | false | false | false | false | false | false | false | false | false | false |
| \|PWY-5816\| | all trans undecaprenyl diphosphate biosynthesis | false | false | false | false | false | false | false | false | false | false | false | false | false | false | false | false | false |
| \|PWY-5817\| | dodecaprenyl diphosphate biosynthesis | false | false | false | false | false | false | false | false | false | false | false | false | false | false | false | false | false |
| \|PWY-5818\| | validamycin biosynthesis | false | false | false | false | false | false | false | false | false | false | false | false | false | false | false | false | false |
| \|PWY-581\| | indole-3-acetate biosynthesis II | false | false | false | false | false | false | false | false | false | false | false | false | false | false | false | false | false |
| \|PWY-5821\| | dalcochinin biosynthesis | false | false | false | false | false | false | false | false | false | false | false | false | false | false | false | false | false |
| \|PWY-5822\| | trichloroethene degradation | false | false | false | false | false | false | false | false | false | false | false | false | false | false | false | false | false |
| \|PWY-5825\| | dalpatein and dalnigrein biosynthesis | false | false | false | false | false | false | false | false | false | false | false | false | false | false | false | false | false |
| \|PWY-5826\| | hypoglycin biosynthesis | false | false | false | false | false | false | false | false | false | false | false | false | false | false | false | false | false |
| \|PWY-5827\| | heliocides biosynthesis | false | false | false | false | false | false | false | false | false | false | false | false | false | false | false | false | false |
| \|PWY-5828\| | lacinilene C biosynthesis | false | false | false | false | false | false | false | false | false | false | false | false | false | false | false | false | false |
| \|PWY-5829\| | geraniol and geranial biosynthesis | false | false | false | false | false | false | false | false | false | false | false | false | false | false | false | false | false |
| \|PWY-5830\| | CDP-ascarylose biosynthesis | false | false | false | false | false | false | false | false | false | false | false | false | false | false | false | false | false |
| \|PWY-5831\| | CDP-abequose biosynthesis | false | false | false | false | false | false | false | false | false | false | false | false | false | false | false | false | false |
| \|PWY-5832\| | CDP-paratose biosynthesis | false | false | false | false | false | false | false | false | false | false | false | false | false | false | false | false | false |
| \|PWY-5833\| | CDP-4-dehydro-3,6-dideoxy-D-glucose biosynthesis | false | false | false | false | false | false | false | false | false | false | false | false | false | false | false | false | false |
| \|PWY-5834\| | CDP-tyvelose biosynthesis | false | false | false | false | false | false | false | false | false | false | false | false | false | false | false | false | false |
| \|PWY-5835\| | geranyl acetate biosynthesis | false | false | false | false | false | false | false | false | false | false | false | false | false | false | false | false | false |
| \|PWY-5837\| | 2-carboxy-1,4-naphthoquinol biosynthesis | false | false | false | false | false | false | false | false | false | false | false | false | false | false | false | false | false |
| \|PWY-5839\| | menaquinol-7 biosynthesis | false | false | false | false | false | false | false | false | false | false | false | false | false | false | false | false | false |
| \|PWY-5843\| | cocaine biosynthesis | false | false | false | false | false | false | false | false | false | false | false | false | false | false | false | false | false |
| \|PWY-5844\| | menaquinol-9 biosynthesis | false | false | false | false | false | false | false | false | false | false | false | false | false | false | false | false | false |
| \|PWY-5846\| | colchicine biosynthesis | false | false | false | false | false | false | false | false | false | false | false | false | false | false | false | false | false |
| \|PWY-5848\| | cinchona alkaloids biosynthesis | false | false | false | false | false | false | false | false | false | false | false | false | false | false | false | false | false |
| \|PWY-5849\| | menaquinol-6 biosynthesis | false | false | false | false | false | false | false | false | false | false | false | false | false | false | false | false | false |
| \|PWY-5851\| | demethylmenaquinol-9 biosynthesis | false | false | false | false | false | false | false | false | false | false | false | false | false | false | false | false | false |
| \|PWY-5852\| | demethylmenaquinol-8 biosynthesis I | false | false | false | false | false | false | false | false | false | false | false | false | false | false | false | false | false |
| \|PWY-5853\| | demethylmenaquinol-6 biosynthesis I | false | false | false | false | false | false | false | false | false | false | false | false | false | false | false | false | false |
| \|PWY-5855\| | ubiquinol-7 biosynthesis (early decarboxylation) | false | false | false | false | false | false | false | false | false | false | false | false | false | false | false | false | false |
| \|PWY-5856\| | ubiquinol-9 biosynthesis (early decarboxylation) | false | false | false | false | false | false | false | false | false | false | false | false | false | false | false | false | false |
| \|PWY-5857\| | ubiquinol-10 biosynthesis (early decarboxylation) | false | false | false | false | false | false | false | false | false | false | false | false | false | false | false | false | false |
| \|PWY-5859\| | eugenol and isoeugenol biosynthesis | false | false | false | false | false | false | false | false | false | false | false | false | false | false | false | false | false |
| \|PWY-5867\| | t-anethole biosynthesis | false | false | false | false | false | false | false | false | false | false | false | false | false | false | false | false | false |
| \|PWY-5870\| | ubiquinol-8 biosynthesis (late decarboxylation) | false | false | false | false | false | false | false | false | false | false | false | false | false | false | false | false | false |
| \|PWY-5871\| | ubiquinol-9 biosynthesis (late decarboxylation) | false | false | false | false | false | false | false | false | false | false | false | false | false | false | false | false | false |
| \|PWY-5872\| | ubiquinol-10 biosynthesis (late decarboxylation) | false | false | false | false | false | false | false | false | false | false | false | false | false | false | false | false | false |
| \|PWY-5873\| | ubiquinol-7 biosynthesis (late decarboxylation) | false | false | false | false | false | false | false | false | false | false | false | false | false | false | false | false | false |
| \|PWY-5874\| | heme degradation I | false | false | false | false | false | false | false | false | false | false | false | false | false | false | false | false | false |
| \|PWY-5875\| | staphyloxanthin biosynthesis | false | false | false | false | false | false | false | false | false | false | false | false | false | false | false | false | false |
| \|PWY-5876\| | magnoflorine biosynthesis | false | false | false | false | false | false | false | false | false | false | false | false | false | false | false | false | false |
| \|PWY-5877\| | beta-carboline biosynthesis | false | false | false | false | false | false | false | false | false | false | false | false | false | false | false | false | false |
| \|PWY-5882\| | epoxypseudoisoeugenol-2-methylbutanoate biosynthesis | false | false | false | false | false | false | false | false | false | false | false | false | false | false | false | false | false |

| | | | | | | | | | | | | | | | | | | |
|---|---|---|---|---|---|---|---|---|---|---|---|---|---|---|---|---|---|---|
| \|PWY-5883\| | ephedrine biosynthesis | false | false | false | false | false | false | false | false | false | false | false | false | false | false | false | false | false |
| \|PWY-5884\| | wax esters biosynthesis I | false | false | false | false | false | false | false | false | false | false | false | false | false | false | false | false | false |
| \|PWY-5885\| | wax esters biosynthesis II | false | false | false | false | false | false | false | false | false | false | false | false | false | false | false | false | false |
| \|PWY-5887\| | albaflavenone biosynthesis | false | false | false | false | false | false | false | false | false | false | false | false | false | false | false | false | false |
| \|PWY-5888\| | rhodoquinone-10 biosynthesis | false | false | false | false | false | false | false | false | false | false | false | false | false | false | false | false | false |
| \|PWY-5889\| | rhodoquinone-9 biosynthesis | false | false | false | false | false | false | false | false | false | false | false | false | false | false | false | false | false |
| \|PWY-5890\| | menaquinol-10 biosynthesis | false | false | false | false | false | false | false | false | false | false | false | false | false | false | false | false | false |
| \|PWY-5891\| | menaquinol-11 biosynthesis | false | false | false | false | false | false | false | false | false | false | false | false | false | false | false | false | false |
| \|PWY-5892\| | menaquinol-12 biosynthesis | false | false | false | false | false | false | false | false | false | false | false | false | false | false | false | false | false |
| \|PWY-5893\| | tridecaprenyl diphosphate biosynthesis | false | false | false | false | false | false | false | false | false | false | false | false | false | false | false | false | false |
| \|PWY-5895\| | menaquinol-13 biosynthesis | false | false | false | false | false | false | false | false | false | false | false | false | false | false | false | false | false |
| \|PWY-5901\| | 2,3-dihydroxybenzoate biosynthesis | false | false | false | false | false | false | false | false | false | false | false | false | false | false | false | false | false |
| \|PWY-5905\| | hypusine biosynthesis | false | false | false | false | false | false | false | false | false | false | false | false | false | false | false | false | false |
| \|PWY-5907\| | homospermidine biosynthesis I | false | false | false | false | false | false | false | false | false | false | false | false | false | false | false | false | false |
| \|PWY-5912\| | 2'-deoxymugineic acid phytosiderophore biosynthesis | false | false | false | false | false | false | false | false | false | false | false | false | false | false | false | false | false |
| \|PWY-5913\| | partial TCA cycle (obligate autotrophs) | false | false | false | false | false | false | false | false | false | false | false | false | false | false | false | false | false |
| \|PWY-5915\| | phycoerythrobilin biosynthesis I | false | false | false | false | false | false | false | false | false | false | false | false | false | false | false | false | false |
| \|PWY-5917\| | phycocyanobilin biosynthesis | false | false | false | false | false | false | false | false | false | false | false | false | false | false | false | false | false |
| \|PWY-5922\| | (4R)-carveol and (4R)-dihydrocarveol degradation | false | false | false | false | false | false | false | false | false | false | false | false | false | false | false | false | false |
| \|PWY-5923\| | limonene degradation I (D-limonene) | false | false | false | false | false | false | false | false | false | false | false | false | false | false | false | false | false |
| \|PWY-5924\| | limonene degradation II (L-limonene) | false | false | false | false | false | false | false | false | false | false | false | false | false | false | false | false | false |
| \|PWY-5925\| | hydroxylated mugineic acid phytosiderophore biosynthesis | false | false | false | false | false | false | false | false | false | false | false | false | false | false | false | false | false |
| \|PWY-5926\| | afrormosin conjugates interconversion | false | false | false | false | false | false | false | false | false | false | false | false | false | false | false | false | false |
| \|PWY-5927\| | (4S)-carveol and (4S)-dihydrocarveol degradation | false | false | false | false | false | false | false | false | false | false | false | false | false | false | false | false | false |
| \|PWY-5928\| | (4R)-carvone biosynthesis | false | false | false | false | false | false | false | false | false | false | false | false | false | false | false | false | false |
| \|PWY-5929\| | puromycin biosynthesis | false | false | false | false | false | false | false | false | false | false | false | false | false | false | false | false | false |
| \|PWY-5930\| | terpentecin biosynthesis | false | false | false | false | false | false | false | false | false | false | false | false | false | false | false | false | false |
| \|PWY-5934\| | iron reduction and absorption | false | false | false | false | false | false | false | false | false | false | false | false | false | false | false | false | false |
| \|PWY-5935\| | 1-tuberculosinyladenosine biosynthesis | false | false | false | false | false | false | false | false | false | false | false | false | false | false | false | false | false |
| \|PWY-5936\| | xyloglucan biosynthesis | false | false | false | false | false | false | false | false | false | false | false | false | false | false | false | false | false |
| \|PWY-5939\| | pyruvate fermentation to (R)-acetoin II | false | false | false | false | false | false | false | false | false | false | false | false | false | false | false | false | false |
| \|PWY-5940\| | streptomycin biosynthesis | false | false | false | false | false | false | false | false | false | false | false | false | false | false | false | false | false |
| \|PWY-5941\| | glycogen degradation II | false | false | false | false | false | false | false | false | false | false | false | false | false | false | false | false | false |
| \|PWY-5943\| | β-carotene biosynthesis | false | false | false | false | false | false | false | false | false | false | false | false | false | false | false | false | false |
| \|PWY-5944\| | zeaxanthin biosynthesis | false | false | false | false | false | false | false | false | false | false | false | false | false | false | false | false | false |
| \|PWY-5945\| | violaxanthin, antheraxanthin and zeaxanthin interconversion | false | false | false | false | false | false | false | false | false | false | false | false | false | false | false | false | false |
| \|PWY-5946\| | α-carotene biosynthesis | false | false | false | false | false | false | false | false | false | false | false | false | false | false | false | false | false |
| \|PWY-5947\| | lutein biosynthesis | false | false | false | false | false | false | false | false | false | false | false | false | false | false | false | false | false |
| \|PWY-5950\| | geosmin biosynthesis | false | false | false | false | false | false | false | false | false | false | false | false | false | false | false | false | false |
| \|PWY-5951\| | (R,R)-butanediol biosynthesis | false | false | false | false | false | false | false | false | false | false | false | false | false | false | false | false | false |
| \|PWY-5954\| | (1'S,5'S)-averufin biosynthesis | false | false | false | false | false | false | false | false | false | false | false | false | false | false | false | false | false |
| \|PWY-5955\| | versicolorin B biosynthesis | false | false | false | false | false | false | false | false | false | false | false | false | false | false | false | false | false |
| \|PWY-5956\| | sterigmatocystin biosynthesis | false | false | false | false | false | false | false | false | false | false | false | false | false | false | false | false | false |
| \|PWY-5957\| | L-nicotianamine biosynthesis | false | false | false | false | false | false | false | false | false | false | false | false | false | false | false | false | false |
| \|PWY-5958\| | acridone alkaloid biosynthesis | false | false | false | false | false | false | false | false | false | false | false | false | false | false | false | false | false |
| \|PWY-5959\| | aflatoxins B1 and G1 biosynthesis | false | false | false | false | false | false | false | false | false | false | false | false | false | false | false | false | false |
| \|PWY-5960\| | aflatoxins B2 and G2 biosynthesis | false | false | false | false | false | false | false | false | false | false | false | false | false | false | false | false | false |
| \|PWY-5963\| | molybdenum cofactor sulfulylation (eukaryotes) | false | false | false | false | false | false | false | false | false | false | false | false | false | false | false | false | false |
| \|PWY-5964\| | guanylyl molybdenum cofactor biosynthesis | false | false | false | false | false | false | false | false | false | false | false | false | false | false | false | false | false |
| \|PWY-5967\| | lupinate biosynthesis | false | false | false | false | false | false | false | false | false | false | false | false | false | false | false | false | false |
| \|PWY-5968\| | cinnamate esters biosynthesis | false | false | false | false | false | false | false | false | false | false | false | false | false | false | false | false | false |
| \|PWY-5971\| | palmitate biosynthesis II (type II fatty acid synthase) | false | false | false | false | false | false | false | false | false | false | false | false | false | false | false | false | false |
| \|PWY-5972\| | stearate biosynthesis I (animals) | false | false | false | false | false | false | false | false | false | false | false | false | false | false | false | false | false |
| \|PWY-5973\| | cis-vaccenate biosynthesis | false | false | false | false | false | false | false | false | false | false | false | false | false | false | false | false | false |
| \|PWY-5975\| | furaneol and mesifurane biosynthesis | false | false | false | false | false | false | false | false | false | false | false | false | false | false | false | false | false |
| \|PWY-5976\| | dhurrin degradation | false | false | false | false | false | false | false | false | false | false | false | false | true | true | true | true | true |
| \|PWY-5978\| | kanosamine biosynthesis I | false | false | false | false | false | false | false | false | false | false | false | false | true | true | true | true | true |
| \|PWY-5979\| | 3-amino-5-hydroxybenzoate biosynthesis | false | false | false | false | false | false | false | false | false | false | false | false | false | false | false | false | false |
| \|PWY-5980\| | xylogalacturonan biosynthesis | false | false | false | false | false | false | false | false | false | false | false | false | false | false | false | false | false |
| \|PWY-5981\| | CDP-diacylglycerol biosynthesis III | false | false | false | false | false | false | false | false | false | false | false | false | false | false | false | false | false |
| \|PWY-5984\| | rifamycin B biosynthesis | false | false | false | false | false | false | false | false | false | false | false | false | false | false | false | false | false |
| \|PWY-5987\| | sorgoleone biosynthesis | false | false | false | false | false | false | false | false | false | false | false | false | false | false | false | false | false |
| \|PWY-5989\| | stearate biosynthesis II (bacteria and plants) | false | false | false | false | false | false | false | false | false | false | false | false | false | false | false | false | false |
| \|PWY-5990\| | lotaustralin biosynthesis | false | false | false | false | false | false | false | false | false | false | false | false | false | false | false | false | false |
| \|PWY-5992\| | thalianol and derivatives biosynthesis | false | false | false | false | false | false | false | false | false | false | false | true | false | false | false | false | false |
| \|PWY-5995\| | linoleate biosynthesis I (plants) | false | false | false | false | false | false | false | false | false | false | false | false | false | false | false | false | false |
| \|PWY-5996\| | oleate biosynthesis II (animals and fungi) | false | false | false | false | false | false | false | false | false | false | false | false | false | false | false | false | false |
| \|PWY-5997\| | α-linolenate biosynthesis I (plants and red algae) | false | false | false | false | false | false | false | false | false | false | false | false | false | false | false | false | false |
| \|PWY-5998\| | γ-linolenate biosynthesis I (plants) | false | false | false | false | false | false | false | false | false | false | false | false | false | false | false | false | false |
| \|PWY-5\| | canavanine biosynthesis | false | false | false | false | false | false | false | false | false | false | false | false | false | false | false | false | false |
| \|PWY5F9-12\| | biphenyl degradation | false | false | false | false | false | false | false | false | false | false | false | false | false | false | false | false | false |
| \|PWY5F9-3233\| | phthalate degradation (aerobic) | false | false | false | false | false | false | false | false | false | false | false | false | false | false | false | false | false |
| \|PWY-6000\| | γ-linolenate biosynthesis II (animals) | false | false | false | false | false | false | false | false | false | false | false | false | false | false | false | false | false |
| \|PWY-6001\| | linoleate biosynthesis II (animals) | false | false | false | false | false | false | false | false | false | false | false | false | false | false | false | false | false |
| \|PWY-6002\| | lotaustralin degradation | false | false | false | false | false | false | false | false | false | false | false | false | false | false | false | false | false |
| \|PWY-6003\| | gramicidin S biosynthesis | false | false | false | false | false | false | false | false | false | false | false | false | false | false | false | false | false |
| \|PWY-6005\| | marneral biosynthesis | false | false | false | false | false | false | false | false | false | false | false | false | false | false | false | false | false |
| \|PWY-6007\| | (3E)-4,8-dimethylnona-1,3,7-triene biosynthesis II | false | false | false | false | false | false | false | false | false | false | false | false | false | false | false | false | false |
| \|PWY-6008\| | baruol biosynthesis | false | false | false | false | false | false | false | false | false | false | false | false | false | false | false | false | false |
| \|PWY-6010\| | apigenin glycosides biosynthesis | false | false | false | false | false | false | false | false | false | false | false | false | false | false | false | false | false |
| \|PWY-6011\| | amygdalin and prunasin degradation | false | false | false | false | false | false | false | false | false | false | false | true | true | true | true | true | true |
| \|PWY-6012-1\| | acyl carrier protein activation | false | false | false | false | false | false | false | false | false | false | false | false | false | false | false | false | false |
| \|PWY-6012\| | acyl carrier protein metabolism | false | false | false | false | false | false | false | false | false | false | false | false | false | false | false | false | false |
| \|PWY-6013\| | crepenynate biosynthesis | false | false | false | false | false | false | false | false | false | false | false | false | false | false | false | false | false |
| \|PWY-6014\| | vernolate biosynthesis I | false | false | false | false | false | false | false | false | false | false | false | false | false | false | false | false | false |
| \|PWY-6015\| | vitexin and derivatives biosynthesis | false | false | false | false | false | false | false | false | false | false | false | false | false | false | false | false | false |
| \|PWY-6019\| | pseudouridine degradation | false | false | false | false | false | false | false | false | false | false | false | false | false | false | false | false | false |
| \|PWY-601\| | glucosinolate biosynthesis from tryptophan | false | false | false | false | false | false | false | false | false | false | false | false | false | false | false | false | false |
| \|PWY-6021\| | nitrilotriacetate degradation | false | false | false | false | false | false | false | false | false | false | false | false | false | false | false | false | false |
| \|PWY-6024\| | isovitexin glycosides biosynthesis | false | false | false | false | false | false | false | false | false | false | false | false | false | false | false | false | false |
| \|PWY-6027\| | capsiconiate biosynthesis | false | false | false | false | false | false | false | false | false | false | false | false | false | false | false | false | false |
| \|PWY-6029\| | 2,3-trans-flavanols biosynthesis | false | false | false | false | false | false | false | false | false | false | false | false | false | false | false | false | false |
| \|PWY-6030\| | serotonin and melatonin biosynthesis | false | false | false | false | false | false | false | false | false | false | false | false | false | false | false | false | false |
| \|PWY-6032\| | digitoxigenin biosynthesis | false | false | false | false | false | false | false | false | false | false | false | false | false | false | false | false | false |
| \|PWY-6035\| | 2,3-cis-flavanols biosynthesis | false | false | false | false | false | false | false | false | false | false | false | false | false | false | false | false | false |
| \|PWY-6036\| | cardenolide glucosides biosynthesis | false | false | false | false | false | false | false | false | false | false | false | false | false | false | false | false | false |
| \|PWY-6038\| | citrate degradation | false | false | false | false | false | false | false | false | false | false | false | false | false | false | false | false | false |
| \|PWY-6039\| | chlorogenic acid biosynthesis I | false | false | false | false | false | false | false | false | false | false | false | false | false | false | false | false | false |
| \|PWY-6040\| | chlorogenic acid biosynthesis II | false | false | false | false | false | false | false | false | false | false | false | false | false | false | false | false | false |
| \|PWY-6041\| | 4-sulfocatechol degradation | false | false | false | false | false | false | false | false | false | false | false | false | false | false | false | false | false |
| \|PWY-6043\| | ethanedisulfonate degradation | false | false | false | false | false | false | false | false | false | false | false | false | false | false | false | false | false |
| \|PWY-6044\| | methanesulfonate degradation | false | false | false | false | false | false | false | false | false | false | false | false | false | false | false | false | false |
| \|PWY-6045\| | methylthiopropanonate degradation II (demethylation) | false | false | false | false | false | false | false | false | false | false | false | false | false | false | false | false | false |
| \|PWY-6046\| | dimethylsulfoniopropanoate degradation I (cleavage) | false | false | false | false | false | false | false | false | false | false | false | false | false | false | false | false | false |
| \|PWY-6047\| | dimethyl sulfide degradation I | false | false | false | false | false | false | false | false | false | false | false | false | false | false | false | false | false |
| \|PWY-6048\| | methylthiopropanoate degradation I (cleavage) | false | false | false | false | false | false | false | false | false | false | false | false | false | false | false | false | false |
| \|PWY-6050\| | dimethyl sulfoxide degradation | false | false | false | false | false | false | false | false | false | false | false | false | false | false | false | false | false |
| \|PWY-6051\| | 2,4,6-trinitrotoluene degradation | false | false | false | false | false | false | false | false | false | false | false | false | false | false | false | false | false |
| \|PWY-6052\| | dimethylsulfoniopropanoate degradation III (demethylation) | false | false | false | false | false | false | false | false | false | false | false | false | false | false | false | false | false |
| \|PWY-6053\| | dimethylsulfoniopropanoate biosynthesis III (algae and phytoplankton) | false | false | false | false | false | false | false | false | false | false | false | false | false | false | false | false | false |
| \|PWY-6054\| | dimethylsulfoniopropanoate biosynthesis I (Wollastonia) | false | false | false | false | false | false | false | false | false | false | false | false | false | false | false | false | false |
| \|PWY-6055\| | dimethylsulfoniopropanoate biosynthesis II (Spartina) | false | false | false | false | false | false | false | false | false | false | false | false | false | false | false | false | false |
| \|PWY-6056\| | dimethylsulfoniopropanoate degradation II (cleavage) | false | false | false | false | false | false | false | false | false | false | false | false | false | false | false | false | false |
| \|PWY-6057\| | dimethylsulfide to cytochrome c2 electron transfer | false | false | false | false | false | false | false | false | false | false | false | false | false | false | false | false | false |
| \|PWY-6058\| | dimethyl sulfone degradation | false | false | false | false | false | false | false | false | false | false | false | false | false | false | false | false | false |
| \|PWY-6059\| | dimethyl sulfide degradation II (oxidation) | false | false | false | false | false | false | false | false | false | false | false | false | false | false | false | false | false |
| \|PWY-6060\| | malonate degradation II (biotin-dependent) | false | false | false | false | false | false | false | false | false | false | false | false | false | false | false | false | false |
| \|PWY-6061\| | bile acid biosynthesis, neutral pathway | false | false | false | false | false | false | false | false | false | false | false | false | false | false | false | false | false |
| \|PWY-6064\| | methylquercetin biosynthesis | false | false | false | false | false | false | false | false | false | false | false | false | false | false | false | false | false |
| \|PWY-6068\| | indican biosynthesis | false | false | false | false | false | false | false | false | false | false | false | false | false | false | false | false | false |
| \|PWY-6069\| | indigo biosynthesis | false | false | false | false | false | false | false | false | false | false | false | false | false | false | false | false | false |
| \|PWY-6073\| | alginate biosynthesis I (algal) | false | false | false | false | false | false | false | false | false | false | false | false | false | false | false | false | false |
| \|PWY-6074\| | zymosterol biosynthesis | false | false | false | false | false | false | false | false | false | false | false | false | false | false | false | false | false |
| \|PWY-6075\| | ergosterol biosynthesis I | false | false | false | false | false | false | false | false | false | false | false | false | false | false | false | false | false |
| \|PWY-6076\| | vitamin D3 biosynthesis | false | false | false | false | false | false | false | false | false | false | false | false | false | false | false | false | false |
| \|PWY-6077\| | anthranilate degradation II (aerobic) | false | false | false | false | false | false | false | false | false | false | false | false | false | false | false | false | false |
| \|PWY-6079\| | anthranilate degradation I (aerobic) | false | false | false | false | false | false | false | false | false | false | false | false | false | false | false | false | false |
| \|PWY-6080\| | 4-ethylphenol degradation (anaerobic) | false | false | false | false | false | false | false | false | false | false | false | false | false | false | false | false | false |
| \|PWY-6081\| | 1,3-dichlorobenzene degradation | false | false | false | false | false | false | false | false | false | false | false | false | false | false | false | false | false |
| \|PWY-6082\| | alginate biosynthesis II (bacterial) | false | false | false | false | false | false | false | false | false | false | false | false | false | false | false | false | false |
| \|PWY-6083\| | chlorobenzene degradation | false | false | false | false | false | false | false | false | false | false | false | false | false | false | false | false | false |
| \|PWY-6084\| | 3,5-dichlorocatechol degradation | false | false | false | false | false | false | false | false | false | false | false | false | false | false | false | false | false |
| \|PWY-6085\| | 2,4-dichlorophenoxyacetate degradation | false | false | false | false | false | false | false | false | false | false | false | false | false | false | false | false | false |
| \|PWY-6086\| | 4-chloro-2-methylphenoxyacetate degradation | false | false | false | false | false | false | false | false | false | false | false | false | false | false | false | false | false |
| \|PWY-6087\| | 4-chlorocatechol degradation | false | false | false | false | false | false | false | false | false | false | false | false | false | false | false | false | false |
| \|PWY-6088\| | 3-chlorobenzoate degradation I (via chlorocatechol) | false | false | false | false | false | false | false | false | false | false | false | false | false | false | false | false | false |
| \|PWY-6089\| | 3-chlorocatechol degradation I (ortho) | false | false | false | false | false | false | false | false | false | false | false | false | false | false | false | false | false |
| \|PWY-6090\| | 1,2-dichlorobenzene degradation | false | false | false | false | false | false | false | false | false | false | false | false | false | false | false | false | false |
| \|PWY-6091\| | 1,2,4-trichlorobenzene degradation | false | false | false | false | false | false | false | false | false | false | false | false | false | false | false | false | false |
| \|PWY-6093\| | 4,5-dichlorocatechol degradation | false | false | false | false | false | false | false | false | false | false | false | false | false | false | false | false | false |
| \|PWY-6094\| | 3,4,6-trichlorocatechol degradation | false | false | false | false | false | false | false | false | false | false | false | false | false | false | false | false | false |
| \|PWY-6095\| | dammara-20,24-diene biosynthesis | false | false | false | false | false | false | false | false | false | false | false | false | false | false | false | false | false |
| \|PWY-6098\| | diploterol biosynthesis | false | false | false | false | false | false | false | false | false | false | false | false | false | false | false | false | false |
| \|PWY-6099\| | 1,2,4,5-tetrachlorobenzene degradation | false | false | false | false | false | false | false | false | false | false | false | false | false | false | false | false | false |
| \|PWY-6100\| | L-carnitine biosynthesis | false | false | false | false | false | false | false | false | false | false | false | false | false | false | false | false | false |
| \|PWY-6102\| | 5-chloro-3-methyl-catechol degradation | false | false | false | false | false | false | false | false | false | false | false | false | false | false | false | false | false |
| \|PWY-6103\| | 3-chlorotoluene degradation I | false | false | false | false | false | false | false | false | false | false | false | false | false | false | false | false | false |
| \|PWY-6104\| | 3-chlorotoluene degradation II | false | false | false | false | false | false | false | false | false | false | false | false | false | false | false | false | false |
| \|PWY-6105\| | botryococcenes and methylated squalene biosynthesis | false | false | false | false | false | false | false | false | false | false | false | false | false | false | false | false | false |
| \|PWY-6107\| | chlorosalicylate degradation | false | false | false | false | false | false | false | false | false | false | false | false | false | false | false | false | false |
| \|PWY-6109\| | mangrove triterpenoid biosynthesis | false | false | false | false | false | false | false | false | false | false | false | false | true | true | true | true | true |
| \|PWY-6111\| | mitochondrial L-carnitine shuttle | false | false | false | false | false | false | false | false | false | false | false | false | false | false | false | false | false |
| \|PWY-6116\| | mannosylfructose biosynthesis | false | false | false | false | false | false | false | false | false | false | false | false | false | false | false | false | false |
| \|PWY-6117\| | spermine and spermidine degradation I | false | false | false | false | false | false | false | false | false | false | false | false | false | false | false | true | false |
| \|PWY-6118\| | glycerol-3-phosphate shuttle | false | false | false | false | false | false | false | false | false | false | false | false | false | false | false | false | false |
| \|PWY-6120\| | L-tyrosine biosynthesis III | false | false | false | false | false | false | false | false | false | false | false | false | false | false | false | false | false |
| \|PWY-6123\| | inosine-5'-phosphate biosynthesis I | false | false | false | false | false | false | false | false | false | false | false | false | false | false | false | false | false |
| \|PWY-6124\| | inosine-5'-phosphate biosynthesis II | false | false | false | false | false | false | false | false | false | false | false | false | false | false | false | false | false |
| \|PWY-6128\| | cis-calamenene related sesquiterpenoids biosynthesis | false | false | false | false | false | false | false | false | false | false | false | false | false | false | false | false | false |
| \|PWY-6129\| | dolichol and dolichyl phosphate biosynthesis | false | false | false | false | false | false | false | false | false | false | false | false | false | false | false | false | false |
| \|PWY-6130\| | glycerol degradation III | false | false | false | false | false | false | false | false | false | false | false | false | false | false | false | true | false |
| \|PWY-6131\| | glycerol degradation II | false | false | false | false | false | false | false | false | false | false | false | true | false | false | false | false | false |
| \|PWY-6132\| | lanosterol biosynthesis | false | false | false | false | false | false | false | false | false | false | false | false | false | false | false | false | false |
| \|PWY-6133\| | (S)-reticuline biosynthesis II | false | false | false | false | false | false | false | false | false | false | false | false | false | false | false | false | false |
| \|PWY-6134\| | L-tyrosine biosynthesis IV | false | false | false | false | false | false | false | false | false | false | false | false | false | false | false | false | false |
| \|PWY-6138\| | CMP-N-acetylneuraminate biosynthesis I (eukaryotes) | false | false | false | false | false | false | false | false | false | false | false | false | false | false | false | false | false |
| \|PWY-6139\| | CMP-N-acetylneuraminate biosynthesis II (bacteria) | false | false | false | false | false | false | false | false | false | false | false | false | false | false | false | false | false |
| \|PWY-6140\| | CMP-2-keto-3-deoxy-D-glycero-D-galacto-nononate biosynthesis | false | false | false | false | false | false | false | false | false | false | false | false | false | false | false | false | false |
| \|PWY-6143\| | CMP-pseudaminate biosynthesis | false | false | false | false | false | false | false | false | false | false | false | false | false | false | false | false | false |
| \|PWY-6144\| | CMP-N-glycoloylneuraminate biosynthesis | false | false | false | false | false | false | false | false | false | false | false | false | false | false | false | false | false |
| \|PWY-6147\| | 6-hydroxymethyl-dihydropterin diphosphate biosynthesis I | false | false | false | false | false | false | false | false | false | false | false | false | false | false | false | false | false |
| \|PWY-6148\| | tetrahydromethanopterin biosynthesis | false | false | false | false | false | false | false | false | false | false | false | false | false | false | false | false | false |
| \|PWY-6151\| | S-adenosyl-L-methionine salvage I | false | false | false | false | false | false | false | false | false | false | false | false | false | false | false | false | false |
| \|PWY-6153\| | autoinducer AI-2 biosynthesis I | false | false | false | false | false | false | false | false | false | false | false | false | false | false | false | false | false |
| \|PWY-6154\| | autoinducer AI-2 biosynthesis II (Vibrio) | false | false | false | false | false | false | false | false | false | false | false | false | false | false | false | false | false |
| \|PWY-6157\| | autoinducer AI-1 biosynthesis | false | false | false | false | false | false | false | false | false | false | false | false | false | false | false | false | false |
| \|PWY-6158\| | creatine phosphate biosynthesis | false | false | false | false | false | false | false | false | false | false | false | false | false | false | false | false | false |
| \|PWY-6160\| | 3-dehydroquinate biosynthesis II (archaea) | false | false | false | false | false | false | false | false | false | false | false | false | false | false | false | false | false |
| \|PWY-6167\| | flavin biosynthesis II (archaea) | false | false | false | false | false | false | false | false | false | false | false | false | false | false | false | false | false |
| \|PWY-6168\| | flavin biosynthesis III (fungi) | false | false | false | false | false | false | false | false | false | false | false | false | false | false | false | false | false |
| \|PWY-6173\| | histamine biosynthesis | false | false | false | false | false | false | false | false | false | false | false | false | false | false | false | false | false |
| \|PWY-6174\| | mevalonate pathway II (haloarchaea) | false | false | false | false | false | false | false | false | false | false | false | false | false | false | false | false | false |
| \|PWY-6178\| | 2,4,6-trichlorophenol degradation | false | false | false | false | false | false | false | false | false | false | false | false | false | false | false | false | false |
| \|PWY-6181\| | histamine degradation | false | false | false | false | false | false | false | false | false | false | false | false | false | false | false | false | false |
| \|PWY-6183\| | salicylate degradation I | false | false | false | false | false | false | false | false | false | false | false | false | false | false | false | false | false |
| \|PWY-6184\| | methylsalicylate degradation | false | false | false | false | false | false | false | false | false | false | false | false | false | false | false | false | false |
| \|PWY-6185\| | 4-methylcatechol degradation (ortho cleavage) | false | false | false | false | false | false | false | false | false | false | false | false | false | false | false | false | false |
| \|PWY-6190\| | 2,4-dichlorotoluene degradation | false | false | false | false | false | false | false | false | false | false | false | false | false | false | false | false | false |
| \|PWY-6191\| | 2,5-dichlorotoluene degradation | false | false | false | false | false | false | false | false | false | false | false | false | false | false | false | false | false |
| \|PWY-6192\| | 3,4-dichlorotoluene degradation | false | false | false | false | false | false | false | false | false | false | false | false | false | false | false | false | false |
| \|PWY-6193\| | 3-chlorocatechol degradation II (ortho) | false | false | false | false | false | false | false | false | false | false | false | false | false | false | false | false | false |
| \|PWY-6197\| | chlorinated phenols degradation | false | false | false | false | false | false | false | false | false | false | false | false | false | false | false | false | false |
| \|PWY-6199\| | quercetin sulfate biosynthesis | false | false | false | false | false | false | false | false | false | false | false | false | false | false | false | false | false |
| \|PWY-6200\| | 2,4,5-trichlorophenoxyacetate degradation | false | false | false | false | false | false | false | false | false | false | false | false | false | false | false | false | false |
| \|PWY-6210\| | 2-aminophenol degradation | false | false | false | false | false | false | false | false | false | false | false | false | false | false | false | false | false |
| \|PWY-6214\| | 3-chlorocatechol degradation III (meta pathway) | false | false | false | false | false | false | false | false | false | false | false | false | false | false | false | false | false |
| \|PWY-6215\| | 4-chlorobenzoate degradation | false | false | false | false | false | false | false | false | false | false | false | false | false | false | false | false | false |
| \|PWY-6216\| | 3-chlorobenzoate degradation II (via protocatechuate) | false | false | false | false | false | false | false | false | false | false | false | false | false | false | false | false | false |
| \|PWY-6217\| | 3,4-dichlorobenzoate degradation | false | false | false | false | false | false | false | false | false | false | false | false | false | false | false | false | false |
| \|PWY-6219\| | indole-3-acetate inactivation VIII | false | false | false | false | false | false | false | false | false | false | false | false | false | false | false | false | false |
| \|PWY-621\| | sucrose degradation III (sucrose invertase) | false | false | false | false | false | false | false | false | false | false | false | false | false | false | false | false | false |
| \|PWY-6220\| | jasmonoyl-amino acid conjugates biosynthesis I | false | false | false | false | false | false | false | false | false | false | false | false | false | false | false | false | false |
| \|PWY-6221\| | 2-chlorobenzoate degradation | false | false | false | false | false | false | false | false | false | false | false | false | false | false | false | false | false |
| \|PWY-6223\| | gentisate degradation I | false | false | false | false | false | false | false | false | false | false | false | false | false | false | false | false | false |
| \|PWY-6224\| | salicylate degradation II | false | false | false | false | false | false | false | false | false | false | false | false | false | false | false | false | false |
| \|PWY-6228\| | 3-chlorobenzoate degradation III (via gentisate) | false | false | false | false | false | false | false | false | false | false | false | false | false | false | false | false | false |
| \|PWY-622\| | starch biosynthesis | false | false | false | false | false | false | false | false | false | false | false | false | false | false | false | false | false |
| \|PWY-6232\| | chrysoeriol biosynthesis | false | false | false | false | false | false | false | false | false | false | false | false | false | false | false | false | false |
| \|PWY-6233\| | jasmonoyl-amino acid conjugates biosynthesis II | false | false | false | false | false | false | false | false | false | false | false | false | false | false | false | false | false |
| \|PWY-6239\| | luteolin glycosides biosynthesis | false | false | false | false | false | false | false | false | false | false | false | false | false | false | false | false | false |
| \|PWY-6241\| | thyroid hormone biosynthesis | false | false | false | false | false | false | false | false | false | false | false | false | false | false | false | false | false |
| \|PWY-6243\| | bergamotene biosynthesis I | false | false | false | false | false | false | false | false | false | false | false | false | false | false | false | false | false |
| \|PWY-6244\| | bergamotene biosynthesis II | false | false | false | false | false | false | false | false | false | false | false | false | false | false | false | false | false |
| \|PWY-6254\| | santalene biosynthesis I | false | false | false | false | false | false | false | false | false | false | false | false | false | false | false | false | false |
| \|PWY-6257\| | curcumene biosynthesis | false | false | false | false | false | false | false | false | false | false | false | false | false | false | false | false | false |
| \|PWY-6258\| | patchoulol biosynthesis | false | false | false | false | false | false | false | false | false | false | false | false | false | false | false | false | false |
| \|PWY-6260\| | thyroid hormone metabolism I (via deiodination) | false | false | false | false | false | false | false | false | false | false | false | false | false | false | false | false | false |
| \|PWY-6261\| | thyroid hormone metabolism II (via conjugation and/or degradation) | false | false | false | false | false | false | false | false | false | false | false | false | false | false | false | false | false |
| \|PWY-6262\| | demethylmenaquinol-8 biosynthesis II | false | false | false | false | false | false | false | false | false | false | false | false | false | false | false | false | false |
| \|PWY-6265\| | zerumbone biosynthesis | false | false | false | false | false | false | false | false | false | false | false | false | false | false | false | false | false |
| \|PWY-6268\| | adenosylcobalamin salvage from cobalamin | false | false | false | false | false | false | false | false | false | false | false | false | false | false | false | false | false |
| \|PWY-6271\| | eudesmol biosynthesis | false | false | false | false | false | false | false | false | false | false | false | true | false | false | false | false | false |
| \|PWY-6273\| | phosphatidylethanolamine biosynthesis III | false | false | false | false | false | false | false | false | false | false | false | true | false | false | false | false | false |
| \|PWY-6275\| | β-caryophyllene biosynthesis | false | false | false | false | false | false | false | false | false | false | false | false | false | false | false | false | false |
| \|PWY-6278\| | botrydial biosynthesis | false | false | false | false | false | false | false | false | false | false | false | false | false | false | false | false | false |
| \|PWY-6279\| | myxol-2' fucoside biosynthesis | false | false | false | false | false | false | false | false | false | false | false | false | false | false | false | false | false |
| \|PWY-6280\| | synechoxanthin biosynthesis | false | false | false | false | false | false | false | false | false | false | false | false | false | false | false | false | false |
| \|PWY-6282\| | palmitoleate biosynthesis I (from (5Z)-dodec-5-enoate) | false | false | false | false | false | false | false | false | false | false | false | false | false | false | false | false | false |
| \|PWY-6286\| | spheroidene and spheroidenone biosynthesis | false | false | false | false | false | false | false | false | false | false | false | false | false | false | false | false | false |
| \|PWY-6287\| | neurosporene biosynthesis | false | false | false | false | false | false | false | false | false | false | false | false | false | false | false | false | false |
| \|PWY-6288\| | zeaxanthin-β-D-diglucoside biosynthesis | false | false | false | false | false | false | false | false | false | false | false | false | false | false | false | false | false |
| \|PWY-6289\| | petrobactin biosynthesis | false | false | false | false | false | false | false | false | false | false | false | false | false | false | false | false | false |
| \|PWY-6290\| | β-cubebene biosynthesis | false | false | false | false | false | false | false | false | false | false | false | false | false | false | false | false | false |
| \|PWY-6291\| | valencene and 7-epi-α-selinene biosynthesis | false | false | false | false | false | false | false | false | false | false | false | false | false | false | false | false | false |
| \|PWY-6294\| | selinene biosynthesis | false | false | false | false | false | false | false | false | false | false | false | false | false | false | false | false | false |
| \|PWY-6297\| | tuberonate glucoside biosynthesis | false | false | false | false | false | false | false | false | false | false | false | false | false | false | false | false | false |
| \|PWY-6302\| | dihydroconiferyl alcohol biosynthesis | false | false | false | false | false | false | false | false | false | false | false | false | false | false | false | false | false |
| \|PWY-6303\| | methyl indole-3-acetate interconversion | false | false | false | false | false | false | false | false | false | false | false | false | false | false | false | false | false |
| \|PWY-6304\| | casbene biosynthesis | false | false | false | false | false | false | false | false | false | false | false | false | false | false | false | false | false |
| \|PWY-6307\| | L-tryptophan degradation X (mammalian, via tryptamine) | false | false | false | false | false | false | false | false | false | false | false | false | false | false | false | false | false |
| \|PWY-6308\| | L-cysteine biosynthesis II (tRNA-dependent) | false | false | false | false | false | false | false | false | false | false | false | false | false | false | false | false | false |
| \|PWY-6310\| | aloesone biosynthesis II | false | false | false | false | false | false | false | false | false | false | false | false | false | false | false | false | false |
| \|PWY-6312\| | aloin biosynthesis | false | false | false | false | false | false | false | false | false | false | false | false | false | false | false | false | false |
| \|PWY-6313\| | serotonin degradation | false | false | false | false | false | false | false | false | false | false | false | false | false | false | false | false | false |
| \|PWY-6314\| | plumbagin biosynthesis | false | false | false | false | false | false | false | false | false | false | false | false | false | false | false | false | false |
| \|PWY-6316\| | aromatic polyketides biosynthesis | false | false | false | false | false | false | false | false | false | false | false | false | false | false | false | false | false |
| \|PWY-6318\| | L-phenylalanine degradation IV (mammalian, via side chain) | false | false | false | false | false | false | false | false | false | false | false | true | false | false | false | false | false |
| \|PWY-6320\| | phaselate biosynthesis | false | false | false | false | false | false | false | false | false | false | false | false | false | false | false | false | false |
| \|PWY-6322\| | phosphinothricin tripeptide biosynthesis | false | false | false | false | false | false | false | false | false | false | false | false | true | true | true | false | true |
| \|PWY-6323\| | benzoylanthranilate biosynthesis | false | false | false | false | false | false | false | false | false | false | false | false | false | false | false | false | false |
| \|PWY-6324\| | rebeccamycin biosynthesis | false | false | false | false | false | false | false | false | false | false | false | false | false | false | false | false | false |
| \|PWY-6325\| | echinatin biosynthesis | false | false | false | false | false | false | false | false | false | false | false | false | false | false | false | false | false |
| \|PWY-6326\| | camptothecin biosynthesis | false | false | false | false | false | false | false | false | false | false | false | false | false | false | false | false | false |
| \|PWY-6327\| | tetrathionate oxidation | false | false | false | false | false | false | false | false | false | false | false | false | false | false | false | false | false |
| \|PWY-6328\| | L-lysine degradation X | false | false | false | false | false | false | false | false | false | false | true | false | true | true | true | true | true |
| \|PWY-6330\| | acetaldehyde biosynthesis II | false | false | false | false | false | false | false | false | false | false | false | false | false | false | false | false | false |
| \|PWY-6332\| | coumestrol biosynthesis | false | false | false | false | false | false | false | false | false | false | false | false | false | false | false | false | false |
| \|PWY-6334\| | L-dopa degradation I (mammalian) | false | false | false | false | false | false | false | false | false | false | false | false | false | false | false | false | false |
| \|PWY-6336\| | protocatechuate degradation III (para-cleavage pathway) | false | false | false | false | false | false | false | false | false | false | false | false | false | false | false | false | false |
| \|PWY-6337\| | dehydroscoulerine biosynthesis | false | false | false | false | false | false | false | false | false | false | false | false | false | false | false | false | false |
| \|PWY-6340\| | 5,5'-dehydrodivanillate degradation | false | false | false | false | false | false | false | false | false | false | false | false | false | false | false | false | false |
| \|PWY-6341\| | guaiacylglycerol-β-guaiacyl ether degradation | false | false | false | false | false | false | false | false | false | false | false | false | false | false | false | false | false |
| \|PWY-6342\| | noradrenaline and adrenaline degradation | false | false | false | false | false | false | false | false | false | false | false | false | false | false | false | false | false |
| \|PWY-6343\| | ferulate degradation | false | false | false | false | false | false | false | false | false | false | false | false | false | false | false | false | false |
| \|PWY-6344\| | L-arginine degradation XIV (oxidative Stickland reaction) | false | false | false | false | false | false | false | false | false | false | false | false | false | false | false | false | false |
| \|PWY-6345\| | K-252 biosynthesis | false | false | false | false | false | false | false | false | false | false | false | false | false | false | false | false | false |
| \|PWY-6346\| | staurosporine biosynthesis | false | false | false | false | false | false | false | false | false | false | false | false | false | false | false | false | false |
| \|PWY-6349\| | C20,20 CDP-archaeol biosynthesis | false | false | false | false | false | false | false | false | false | false | false | false | false | false | false | false | false |
| \|PWY-6351\| | D-myo-inositol (1,4,5)-trisphosphate biosynthesis | false | false | false | false | false | false | false | false | false | false | false | false | false | false | false | false | false |
| \|PWY-6352\| | 3-phosphoinositide biosynthesis | false | false | false | false | false | false | false | false | false | false | false | false | false | false | false | false | false |
| \|PWY-6361\| | 1D-myo-inositol hexakisphosphate biosynthesis I (from Ins(1,4,5)P3) | false | false | false | false | false | false | false | false | false | false | false | false | false | false | false | false | false |
| \|PWY-6362\| | 1D-myo-inositol hexakisphosphate biosynthesis II (mammalian) | false | false | false | false | false | false | false | false | false | false | false | false | false | false | false | false | false |
| \|PWY-6363\| | D-myo-inositol (1,4,5)-trisphosphate degradation | false | false | false | false | false | false | false | false | false | false | false | false | false | false | false | false | false |
| \|PWY-6364\| | D-myo-inositol (1,3,4)-trisphosphate biosynthesis | false | false | false | false | false | false | false | false | false | false | false | false | false | false | false | false | false |
| \|PWY-6365\| | D-myo-inositol (3,4,5,6)-tetrakisphosphate biosynthesis | false | false | false | false | false | false | false | false | false | false | false | false | false | false | false | false | false |
| \|PWY-6366\| | D-myo-inositol (1,4,5,6)-tetrakisphosphate biosynthesis | false | false | false | false | false | false | false | false | false | false | false | false | false | false | false | false | false |
| \|PWY-6367\| | D-myo-inositol-5-phosphate metabolism | false | false | false | false | false | false | false | false | false | false | false | false | false | false | false | false | false |
| \|PWY-6368\| | 3-phosphoinositide degradation | false | false | false | false | false | false | false | false | false | false | false | false | false | false | false | false | false |
| \|PWY-6369\| | inositol diphosphates biosynthesis | false | false | false | false | false | false | false | false | false | false | false | false | false | false | false | false | false |
| \|PWY-6370\| | ascorbate recycling (cytosolic) | false | false | false | false | false | false | false | false | false | false | false | false | false | false | false | false | false |
| \|PWY-6372\| | 1D-myo-inositol hexakisphosphate biosynthesis IV (Dictyostelium) | false | false | false | false | false | false | false | false | false | false | false | false | false | false | false | false | false |
| \|PWY-6373\| | acrylate degradation I | false | false | false | false | false | false | false | false | false | false | false | false | false | false | false | false | false |
| \|PWY-6375\| | desferrioxamine E biosynthesis | false | false | false | false | false | false | false | false | false | false | false | false | false | false | false | false | false |
| \|PWY-6376\| | desferrioxamine B biosynthesis | false | false | false | false | false | false | false | false | false | false | false | false | false | false | false | false | false |
| \|PWY-6377\| | α-tocopherol degradation | false | false | false | false | false | false | false | false | false | false | false | false | false | false | false | false | false |
| \|PWY-6378\| | putrebactin biosynthesis | false | false | false | false | false | false | false | false | false | false | false | false | false | false | false | false | false |
| \|PWY-6379\| | alcaligin biosynthesis | false | false | false | false | false | false | false | false | false | false | false | false | false | true | false | false | true |
| \|PWY-6381\| | bisucaberin biosynthesis | false | false | false | false | false | false | false | false | false | false | false | false | false | false | false | false | false |
| \|PWY-6383\| | mono-trans, poly-cis decaprenyl phosphate biosynthesis | false | false | false | false | false | false | false | false | false | false | false | false | false | false | false | false | false |
| \|PWY-6388\| | (S,S)-butanediol degradation | false | false | false | false | false | false | false | false | false | false | false | false | false | false | false | false | false |
| \|PWY-6389\| | pyruvate fermentation to (S)-acetoin | false | false | false | false | false | false | false | false | false | false | false | false | false | false | false | false | false |
| \|PWY-6390\| | (S,S)-butanediol biosynthesis | false | false | false | false | false | false | false | false | false | false | false | false | false | false | false | false | false |
| \|PWY-6391\| | meso-butanediol biosynthesis I | false | false | false | false | false | false | false | false | false | false | false | false | false | false | false | false | false |
| \|PWY-6392\| | meso-butanediol biosynthesis II | false | false | false | false | false | false | false | false | false | false | false | false | false | false | false | false | false |
| \|PWY-6397\| | mycolyl-arabinogalactan-peptidoglycan complex biosynthesis | false | false | false | false | false | false | false | false | false | false | false | false | false | false | false | false | false |
| \|PWY-6398\| | melatonin degradation I | false | false | false | false | false | false | false | false | false | false | false | false | false | false | false | false | false |
| \|PWY-6399\| | melatonin degradation II | false | false | false | false | false | false | false | false | false | false | false | false | false | false | false | false | false |
| \|PWY-6400\| | melatonin degradation III | false | false | false | false | false | false | false | false | false | false | false | false | false | false | false | false | false |
| \|PWY-6401\| | hispidol and hispidol 4'-O-β-D-glucoside biosynthesis | false | false | false | false | false | false | false | false | false | false | false | false | false | false | false | false | false |
| \|PWY-6403\| | carrageenan biosynthesis | false | false | false | false | false | false | false | false | false | false | false | false | false | false | false | false | false |
| \|PWY-6405\| | Rapoport-Luebering glycolytic shunt | false | false | false | false | false | false | false | false | false | false | false | false | false | false | false | false | false |
| \|PWY-6406\| | salicylate biosynthesis I | false | false | false | false | false | false | false | false | false | false | false | false | false | false | false | false | false |
| \|PWY-6407\| | yersiniabactin biosynthesis | false | false | false | false | false | false | false | false | false | false | false | false | false | false | false | false | false |
| \|PWY-6408\| | pyochelin biosynthesis | false | false | false | false | false | false | false | false | false | false | false | false | false | false | false | false | false |
| \|PWY-6409\| | pyoverdine I biosynthesis | false | false | false | false | false | false | false | false | false | false | false | false | false | false | false | false | false |
| \|PWY-6411\| | ginsenoside degradation I | false | false | false | false | false | false | false | false | false | false | false | false | false | false | false | false | false |
| \|PWY-6412\| | ginsenoside degradation II | false | false | false | false | false | false | false | false | false | false | false | false | false | false | false | false | false |
| \|PWY-6413\| | ginsenoside degradation III | false | false | false | false | false | false | false | false | false | false | false | false | false | false | false | false | false |
| \|PWY-6415\| | L-ascorbate biosynthesis V (euglena, D-galacturonate pathway) | false | false | false | false | false | false | false | false | false | false | false | false | false | false | false | false | false |
| \|PWY-6416\| | quinate degradation II | false | false | false | false | false | false | false | false | false | false | false | false | false | false | false | false | false |
| \|PWY-6418\| | 4-hydroxycoumarin and dicoumarol biosynthesis | false | false | false | false | false | false | false | false | false | false | false | false | false | false | false | false | false |
| \|PWY-6419\| | shikimate degradation II | false | false | false | false | false | false | false | false | false | false | false | false | false | false | false | false | false |
| \|PWY-641\| | proanthocyanidins biosynthesis from flavanols | false | false | false | false | false | false | false | false | false | false | false | false | false | false | false | false | false |
| \|PWY-6420\| | pyrroloquinoline quinone biosynthesis | false | false | false | false | false | false | false | false | false | false | false | false | false | false | false | false | false |
| \|PWY-6421\| | arsenate detoxification IV (mycothiol) | false | false | false | false | false | false | false | false | false | false | false | false | false | false | false | false | false |
| \|PWY-6422\| | D-arginine degradation | false | false | false | false | false | false | false | false | false | false | false | false | false | false | false | false | false |
| \|PWY-6423\| | hemoglobin degradation | false | false | false | false | false | false | false | false | false | false | false | true | false | false | false | false | false |
| \|PWY-6425\| | rotenoid biosynthesis II | false | false | false | false | false | false | false | false | false | false | false | false | false | false | false | false | false |
| \|PWY-6426\| | uracil degradation II (oxidative) | false | false | false | false | false | false | false | false | false | false | false | false | false | false | false | false | false |
| \|PWY-6427\| | rot-2'-enonate biosynthesis | false | false | false | false | false | false | false | false | false | false | false | false | false | false | false | false | false |
| \|PWY-6431\| | 4-hydroxybenzoate biosynthesis IV (plants) | false | false | false | false | false | false | false | false | false | false | false | false | false | false | false | false | false |
| \|PWY-6432\| | curcuminoid biosynthesis | false | false | false | false | false | false | false | false | false | false | false | false | false | false | false | false | false |
| \|PWY-6433\| | hydroxylated fatty acid biosynthesis (plants) | false | false | false | false | false | false | false | false | false | false | false | false | false | false | false | false | false |
| \|PWY-6435\| | 4-hydroxybenzoate biosynthesis III (plants) | false | false | false | false | false | false | false | false | false | false | false | false | false | false | false | false | false |
| \|PWY-6436\| | perillyl aldehyde biosynthesis | false | false | false | false | false | false | false | false | false | false | false | false | false | false | false | false | false |
| \|PWY-6437\| | fenchol biosynthesis I | false | false | false | false | false | false | false | false | false | false | false | false | false | false | false | false | false |
| \|PWY-6438\| | phenylphenalenone biosynthesis | false | false | false | false | false | false | false | false | false | false | false | false | false | false | false | false | false |
| \|PWY-6440\| | spermine and spermidine degradation II | false | false | false | false | false | false | false | false | false | false | false | false | false | false | false | false | false |
| \|PWY-6441\| | spermine and spermidine degradation III | false | false | false | false | false | false | false | false | false | false | false | false | false | false | false | false | false |
| \|PWY-6442\| | spermidine hydroxycinnamic acid conjugates biosynthesis | false | false | false | false | false | false | false | false | false | false | false | false | false | false | false | false | false |
| \|PWY-6444\| | benzoate biosynthesis II (CoA-independent, non-β-oxidative) | false | false | false | false | false | false | false | false | false | false | false | false | false | true | true | true | true |
| \|PWY-6445\| | fenchol biosynthesis II | false | false | false | false | false | false | false | false | false | false | false | false | false | false | false | false | false |
| \|PWY-6447\| | trichome monoterpenes biosynthesis | false | false | false | false | false | false | false | false | false | false | false | false | false | false | false | false | false |
| \|PWY-6448\| | hordatine biosynthesis | false | false | false | false | false | false | false | false | false | false | false | false | false | false | false | false | false |
| \|PWY-6449\| | fenchone biosynthesis | false | false | false | false | false | false | false | false | false | false | false | false | false | false | false | false | false |
| \|PWY-6451\| | 3-carene biosynthesis | false | false | false | false | false | false | false | false | false | false | false | false | false | false | false | false | false |
| \|PWY-6453\| | stigma estolide biosynthesis | false | false | false | false | false | false | false | false | false | false | false | false | false | false | false | false | false |
| \|PWY-6454\| | vancomycin resistance I | false | false | false | false | false | false | false | false | false | false | false | false | false | false | false | false | false |
| \|PWY-6455\| | vancomycin resistance II | false | false | false | false | false | false | false | false | false | false | false | false | false | false | false | false | false |
| \|PWY-6456\| | serinol biosynthesis | false | false | false | false | false | false | false | false | false | false | false | false | false | false | false | false | false |

| | | | | | | | | | | | | | | | | | | |
|---|---|---|---|---|---|---|---|---|---|---|---|---|---|---|---|---|---|---|
| \|PWY-6457\| | cinnamoyl-CoA biosynthesis | false | false | false | false | false | false | false | false | false | false | false | false | false | false | false | false | false |
| \|PWY-6458\| | benzoyl-CoA biosynthesis | false | false | false | false | false | false | false | false | false | false | false | false | false | false | false | false | false |
| \|PWY-6459\| | peptidoglycan cross-bridge biosynthesis I (S. aureus) | false | false | false | false | false | false | false | false | false | false | false | false | false | false | false | false | false |
| \|PWY-6461\| | peptidoglycan cross-bridge biosynthesis II (E. faecium) | false | false | false | false | false | false | false | false | false | false | false | false | false | false | false | false | false |
| \|PWY-6462\| | peptidoglycan cross-bridge biosynthesis III (Enterococcus faecalis) | false | false | false | false | false | false | false | false | false | false | false | false | false | false | false | false | false |
| \|PWY-6463\| | peptidoglycan cross-bridge biosynthesis IV (Weissella viridescens) | false | false | false | false | false | false | false | false | false | false | false | false | false | false | false | false | false |
| \|PWY-6464\| | polyvinyl alcohol degradation | false | false | false | false | false | false | false | false | false | false | false | false | false | false | false | false | false |
| \|PWY-6473\| | 4-aminobutanoate degradation IV | false | false | false | false | false | false | false | false | false | false | false | false | false | false | false | false | false |
| \|PWY-6475\| | trans-lycopene biosynthesis II (oxygenic phototrophs and green sulfur bacteria) | false | false | false | false | false | false | false | false | false | false | false | false | false | false | false | false | false |
| \|PWY-6476\| | cytidylyl molybdenum cofactor biosynthesis | false | false | false | false | false | false | false | false | false | false | false | false | false | false | false | false | false |
| \|PWY-6477\| | gibberellin inactivation II (methylation) | false | false | false | false | false | false | false | false | false | false | false | false | false | false | false | false | false |
| \|PWY-6478\| | GDP-D-glycero-α-D-manno-heptose biosynthesis | false | false | false | false | false | false | false | false | false | false | false | false | false | false | false | false | false |
| \|PWY-6481\| | L-dopa and L-dopachrome biosynthesis | false | false | false | false | false | false | false | false | false | false | false | false | false | false | false | false | false |
| \|PWY-6482\| | diphthamide biosynthesis I (archaea) | false | false | false | false | false | false | false | false | false | false | false | false | false | false | false | false | false |
| \|PWY-6483\| | ceramide degradation (generic) | false | false | false | false | false | false | false | false | false | false | false | false | false | false | false | false | false |
| \|PWY-6486\| | D-galacturonate degradation II | false | false | false | false | false | false | false | false | false | false | false | false | false | false | false | false | false |
| \|PWY-6491\| | D-galacturonate degradation IV | false | false | false | false | false | false | false | false | false | false | false | false | false | false | false | false | false |
| \|PWY-6493\| | chanoclavine I aldehyde biosynthesis | false | false | false | false | false | false | false | false | false | false | false | false | false | false | false | false | false |
| \|PWY-6494\| | gibberellin inactivation III (epoxidation) | false | false | false | false | false | false | false | false | false | false | false | false | false | false | false | false | false |
| \|PWY-6495\| | ergotamine biosynthesis | false | false | false | false | false | false | false | false | false | false | false | false | false | false | false | false | false |
| \|PWY-6497\| | D-galactarate degradation II | false | false | false | false | false | false | false | false | false | false | false | false | false | false | false | false | false |
| \|PWY-6498\| | eumelanin biosynthesis | false | false | false | false | false | false | false | false | false | false | false | false | false | false | false | false | false |
| \|PWY-6499\| | D-glucarate degradation II | false | false | false | false | false | false | false | false | false | false | false | false | false | false | false | false | false |
| \|PWY-6501\| | D-glucuronate degradation II | false | false | false | false | false | false | false | false | false | false | false | false | false | false | false | false | false |
| \|PWY-6502\| | 8-oxo-(d)GTP detoxification I | false | false | false | false | false | false | false | false | false | false | false | false | false | false | false | false | false |
| \|PWY-6504\| | anthranilate degradation IV (aerobic) | false | false | false | false | false | false | false | false | false | false | false | false | false | false | false | false | false |
| \|PWY-6507\| | 4-deoxy-L-threo-hex-4-enopyranuronate degradation | false | false | false | false | false | false | false | false | false | false | false | false | false | false | false | false | false |
| \|PWY-6509\| | methanol oxidation to formaldehyde III | false | false | false | false | false | false | false | false | false | false | false | false | false | false | false | false | false |
| \|PWY-6510\| | methanol oxidation to formaldehyde II | false | false | false | false | false | false | false | false | false | false | false | false | false | false | false | false | false |
| \|PWY-6511\| | 3-methylarginine biosynthesis | false | false | false | false | false | false | false | false | false | false | false | false | false | false | false | false | false |
| \|PWY-6512\| | hydrogen oxidation III (anaerobic, NADP) | false | false | false | false | false | false | false | false | false | false | false | false | false | false | false | false | false |
| \|PWY-6515\| | phloridzin biosynthesis | false | false | false | false | false | false | false | false | false | false | false | false | false | false | false | false | false |
| \|PWY-6519\| | 8-amino-7-oxononanoate biosynthesis I | false | false | false | false | false | false | false | false | false | false | false | false | false | false | false | false | false |
| \|PWY-6520\| | nonaprenyl diphosphate biosynthesis II | false | false | false | false | false | false | false | false | false | false | false | false | false | false | false | false | false |
| \|PWY-6523\| | nitrite-dependent anaerobic methane oxidation | false | false | false | false | false | false | false | false | false | false | false | false | false | false | false | false | false |
| \|PWY-6524\| | lychnose and isolychnose biosynthesis | false | false | false | false | false | false | false | false | false | false | false | false | false | false | false | false | false |
| \|PWY-6525\| | stellariose and mediose biosynthesis | false | false | false | false | false | false | false | false | false | false | false | false | false | false | false | true | false |
| \|PWY-6526\| | limonene degradation III (to perillate) | false | false | false | false | false | false | false | false | false | false | false | false | false | false | false | false | false |
| \|PWY-6529\| | chlorate reduction | false | false | false | false | false | false | false | false | false | false | false | false | true | false | true | false | false |
| \|PWY-6530\| | perchlorate reduction | false | false | false | false | false | false | false | false | false | false | false | false | false | false | false | false | false |
| \|PWY-6531\| | mannitol cycle | false | false | false | false | false | false | false | false | false | false | false | false | false | false | false | false | false |
| \|PWY-6532\| | diphenylamine degradation | false | false | false | false | false | false | false | false | false | false | false | false | false | false | false | false | false |
| \|PWY-6533\| | aniline degradation | false | false | false | false | false | false | false | false | false | false | false | false | false | false | false | false | false |
| \|PWY-6534\| | phenylethylamine degradation II | false | false | false | false | false | false | false | false | false | false | false | false | false | false | false | false | false |
| \|PWY-6535\| | 4-aminobutanoate degradation I | false | false | false | false | false | false | false | false | false | false | false | false | true | true | true | false | true |
| \|PWY-6536\| | 4-aminobutanoate degradation III | false | false | false | false | false | false | false | false | false | false | false | false | false | false | false | false | false |
| \|PWY-6537\| | 4-aminobutanoate degradation II | false | false | false | false | false | false | false | false | false | false | false | false | false | false | false | false | false |
| \|PWY-6538\| | caffeine degradation III (bacteria, via demethylation) | false | false | false | false | false | false | false | false | false | false | false | false | false | false | false | false | false |
| \|PWY-6539\| | (Z)-phenylmethanethial S-oxide biosynthesis | false | false | false | false | false | false | false | false | false | false | false | false | false | false | false | false | false |
| \|PWY-6540\| | costunolide biosynthesis | false | false | false | false | false | false | false | false | false | false | false | false | false | false | false | false | false |
| \|PWY-6545\| | pyrimidine deoxyribonucleotides de novo biosynthesis III | false | false | false | false | false | false | false | false | false | false | false | false | false | false | false | false | false |
| \|PWY-6546\| | brassinosteroids inactivation | false | false | false | false | false | false | false | false | false | false | false | false | false | false | false | false | false |
| \|PWY-6549\| | nitrogen remobilization from senescing leaves | false | false | false | false | false | false | false | false | false | false | false | false | false | false | false | false | false |
| \|PWY-6550\| | carbazole degradation | false | false | false | false | false | false | false | false | false | false | false | false | false | false | false | false | false |
| \|PWY-6552\| | caffeine degradation I (main, plants) | false | false | false | false | false | false | false | false | false | false | false | false | false | false | false | false | false |
| \|PWY-6553\| | caffeine degradation II | false | false | false | false | false | false | false | false | false | false | false | false | false | false | false | false | false |
| \|PWY-6554\| | 1D-myo-inositol hexakisphosphate biosynthesis V (from Ins(1,3,4)P3) | false | false | false | false | false | false | false | false | false | false | false | false | false | false | false | false | false |
| \|PWY-6556\| | pyrimidine ribonucleosides salvage II | false | false | false | false | false | false | false | false | false | false | false | false | false | false | false | false | false |
| \|PWY-6557\| | glycosaminoglycan-protein linkage region biosynthesis | false | false | false | false | false | false | false | false | false | false | false | false | false | false | false | false | false |
| \|PWY-6558\| | heparan sulfate biosynthesis | false | false | false | false | false | false | false | false | false | false | false | false | false | false | false | false | false |
| \|PWY-6559\| | spermidine biosynthesis II | false | false | false | false | false | false | false | false | false | false | false | false | false | false | false | false | false |
| \|PWY-6562\| | norspermidine biosynthesis | false | false | false | false | false | false | false | false | false | false | false | false | false | false | false | false | false |
| \|PWY-6566\| | chondroitin biosynthesis | false | false | false | false | false | false | false | false | false | false | false | false | false | false | false | false | false |
| \|PWY-6567\| | chondroitin sulfate biosynthesis | false | false | false | false | false | false | false | false | false | false | false | false | false | false | false | false | false |
| \|PWY-6568\| | dermatan sulfate biosynthesis | false | false | false | false | false | false | false | false | false | false | false | false | false | false | false | false | false |
| \|PWY-6572\| | chondroitin sulfate degradation I (bacterial) | false | false | false | false | false | false | false | false | false | false | false | false | false | false | false | false | false |
| \|PWY-6573\| | chondroitin sulfate degradation (metazoa) | false | false | false | false | false | false | false | false | false | false | false | false | false | false | false | false | false |
| \|PWY-6574\| | achromobactin biosynthesis | false | false | false | false | false | false | false | false | false | false | false | false | false | false | false | false | false |
| \|PWY-6575\| | juvenile hormone III biosynthesis I | false | false | false | false | false | false | false | false | false | false | false | false | false | false | false | false | false |
| \|PWY-6576\| | dermatan sulfate degradation (metazoa) | false | false | false | false | false | false | false | false | false | false | false | false | false | false | false | false | false |
| \|PWY-6577\| | farnesylcysteine salvage pathway | false | false | false | false | false | false | false | false | false | false | false | false | false | false | false | false | false |
| \|PWY-6580\| | phosphatidylinositol biosynthesis I (bacteria) | false | false | false | false | false | false | false | false | false | false | false | false | false | false | false | false | false |
| \|PWY-6581\| | spirilloxanthin and 2,2'-diketo-spirilloxanthin biosynthesis | false | false | false | false | false | false | false | false | false | false | false | false | false | false | false | false | false |
| \|PWY-6583\| | pyruvate fermentation to butanol I | false | false | false | false | false | false | false | false | false | false | false | false | false | false | false | false | false |
| \|PWY-6585\| | 2-methylketone biosynthesis | false | false | false | false | false | false | false | false | false | false | false | true | false | false | false | false | false |
| \|PWY-6587\| | pyruvate fermentation to ethanol III | false | false | false | false | false | false | false | false | false | false | false | false | false | false | false | false | false |
| \|PWY-6588\| | pyruvate fermentation to acetone | false | false | false | false | false | false | false | false | false | false | false | false | false | false | false | false | false |
| \|PWY-6591\| | manganese oxidation I | false | false | false | false | false | false | false | false | false | false | false | false | false | false | false | false | false |
| \|PWY-6592\| | manganese oxidation II | false | false | false | false | false | false | false | false | false | false | false | false | false | false | false | false | false |
| \|PWY-6593\| | sulfoacetate degradation | false | false | false | false | false | false | false | false | false | false | false | false | false | false | false | false | false |
| \|PWY-6596\| | adenosine nucleotides degradation I | false | false | false | false | false | false | false | false | false | false | false | true | false | false | false | false | false |
| \|PWY-6598\| | sciadonate biosynthesis | false | false | false | false | false | false | false | false | false | false | false | false | false | false | false | false | false |
| \|PWY-6602\| | naringenin C-glucosylation | false | false | false | false | false | false | false | false | false | false | false | false | false | false | false | false | false |
| \|PWY-6603\| | dicranin biosynthesis | false | false | false | false | false | false | false | false | false | false | false | false | false | false | false | false | false |
| \|PWY-6605\| | adenine and adenosine salvage II | false | false | false | false | false | false | false | false | false | false | false | false | false | false | false | false | false |
| \|PWY-6606\| | guanosine nucleotides degradation II | false | false | false | false | false | false | false | false | false | false | false | false | false | false | false | false | false |
| \|PWY-6607\| | guanosine nucleotides degradation I | false | false | false | false | false | false | false | false | false | false | false | false | false | false | false | false | false |
| \|PWY-6608\| | guanosine nucleotides degradation III | false | false | false | false | false | false | false | false | false | false | false | false | false | false | false | false | false |
| \|PWY-6609\| | adenine and adenosine salvage III | false | false | false | false | false | false | false | false | false | false | false | false | false | false | false | false | false |
| \|PWY-6611\| | adenine and adenosine salvage V | false | false | false | false | false | false | false | false | false | false | false | false | false | false | false | false | false |
| \|PWY-6613\| | tetrahydrofolate salvage from 5,10-methenyltetrahydrofolate | false | false | false | false | false | false | false | false | false | false | false | false | false | false | false | false | false |
| \|PWY-6614\| | tetrahydrofolate biosynthesis I | false | false | false | false | false | false | false | false | false | false | false | false | false | false | false | false | true |
| \|PWY66-161\| | ethanol degradation III | false | false | false | false | false | false | false | false | false | false | false | false | false | false | false | false | false |
| \|PWY66-162\| | ethanol degradation IV | false | false | false | false | false | false | false | false | false | false | false | false | false | false | false | false | false |
| \|PWY-6616\| | sulfolactate degradation I | false | false | false | false | false | false | false | false | false | false | false | false | false | false | false | false | false |
| \|PWY-6617\| | adenosine nucleotides degradation III | false | false | false | false | false | false | false | false | false | false | false | false | false | false | false | false | false |
| \|PWY-6618\| | guanine and guanosine salvage III | false | false | false | false | false | false | false | false | false | false | false | false | false | false | false | false | false |
| \|PWY-6619\| | adenine and adenosine salvage VI | false | false | false | false | false | false | false | false | false | false | false | false | false | false | false | false | false |
| \|PWY66-201\| | nicotine degradation IV | false | false | false | false | false | false | false | false | false | false | false | false | false | false | false | false | false |
| \|PWY66-221\| | nicotine degradation V | false | false | false | false | false | false | false | false | false | false | false | false | false | false | false | false | false |
| \|PWY-6622\| | heptadecane biosynthesis | false | false | false | false | false | false | false | false | false | false | false | false | false | false | false | false | false |
| \|PWY-6623\| | salicylate glucosides biosynthesis II | false | false | false | false | false | false | false | false | false | false | false | false | false | false | false | false | false |
| \|PWY66-241\| | bupropion degradation | false | false | false | false | false | false | false | false | false | false | false | false | false | false | false | false | false |
| \|PWY-6624\| | salicylate glucosides biosynthesis III | false | false | false | false | false | false | false | false | false | false | false | false | false | false | false | false | false |
| \|PWY-6626\| | cytidine-5'-diphosphate-glycerol biosynthesis | false | false | false | false | false | false | false | false | false | false | false | false | false | false | false | false | false |
| \|PWY-6627\| | salinosporamide A biosynthesis | false | false | false | false | false | false | false | false | false | false | false | false | false | false | false | false | false |
| \|PWY66-301\| | catecholamine biosynthesis | false | false | false | false | false | false | false | false | false | false | false | false | false | false | false | false | false |
| \|PWY-6631\| | tricetin methylation | false | false | false | false | false | false | false | false | false | false | false | false | false | false | false | false | false |
| \|PWY-6633\| | caffeine degradation V (bacteria, via trimethylurate) | false | false | false | false | false | false | false | false | false | false | false | false | false | false | false | false | false |
| \|PWY-6634\| | 3-sulfopropanediol degradation I | false | false | false | false | false | false | false | false | false | false | false | false | false | false | false | false | false |
| \|PWY66-367\| | ketogenesis | false | false | false | false | false | false | false | false | false | false | false | false | false | false | false | false | false |
| \|PWY66-368\| | ketolysis | false | false | false | false | false | false | false | false | false | false | false | false | false | false | false | false | false |
| \|PWY-6636\| | salicylate degradation III | false | false | false | false | false | false | false | false | false | false | false | true | false | false | false | false | false |
| \|PWY66-373\| | sucrose degradation V (sucrose α-glucosidase) | false | false | false | false | false | false | false | false | false | false | false | false | false | false | false | false | false |
| \|PWY66-374\| | C20 prostanoid biosynthesis | false | false | false | false | false | false | false | false | false | false | false | false | false | false | false | false | false |
| \|PWY66-375\| | leukotriene biosynthesis | false | false | false | false | false | false | false | false | false | false | false | false | false | false | false | false | false |
| \|PWY66-377\| | pregnenolone biosynthesis | false | false | false | false | false | false | false | false | false | false | false | false | false | false | false | false | false |
| \|PWY66-378\| | androgen biosynthesis | false | false | false | false | false | false | false | false | false | false | false | false | false | false | false | false | false |
| \|PWY-6637\| | sulfolactate degradation II | false | false | false | false | false | false | false | false | false | false | false | false | false | false | false | false | false |
| \|PWY66-380\| | estradiol biosynthesis I (via estrone) | false | false | false | false | false | false | false | false | false | false | false | false | false | false | false | false | false |
| \|PWY66-381\| | glucocorticoid biosynthesis | false | false | false | false | false | false | false | false | false | false | false | false | false | false | false | false | false |
| \|PWY66-382\| | mineralocorticoid biosynthesis | false | false | false | false | false | false | false | false | false | false | false | false | false | false | false | false | false |
| \|PWY66-385\| | dTMP de novo biosynthesis (mitochondrial) | false | false | false | false | false | false | false | false | false | false | false | false | false | false | false | false | false |
| \|PWY66-387\| | 3-methyl-branched fatty acid α-oxidation | false | false | false | false | false | false | false | false | false | false | false | false | false | false | false | false | false |
| \|PWY66-388\| | ceramide degradation by α-oxidation | false | false | false | false | false | false | false | false | false | false | false | false | false | false | false | false | false |
| \|PWY-6638\| | sulfolactate degradation III | false | false | false | false | false | false | false | false | false | false | false | false | false | false | false | false | false |
| \|PWY66-391\| | fatty acid β-oxidation VI (mammalian peroxisome) | false | false | false | false | false | false | false | false | false | false | false | false | false | false | false | false | false |
| \|PWY66-392\| | lipoxin biosynthesis | false | false | false | false | false | false | false | false | false | false | false | false | false | false | false | false | false |
| \|PWY66-393\| | 15-epi-lipoxin biosynthesis | false | false | false | false | false | false | false | false | false | false | false | false | false | false | false | false | false |
| \|PWY66-394\| | aspirin triggered resolvin E biosynthesis | false | false | false | false | false | false | false | false | false | false | false | false | false | false | false | false | false |
| \|PWY66-395\| | aspirin triggered resolvin D biosynthesis | false | false | false | false | false | false | false | false | false | false | false | false | false | false | false | false | false |
| \|PWY66-397\| | resolvin D biosynthesis | false | false | false | false | false | false | false | false | false | false | false | false | false | false | false | false | false |
| \|PWY66-398\| | TCA cycle III (animals) | false | false | false | false | false | false | false | false | false | false | false | false | false | false | false | false | false |
| \|PWY66-399\| | gluconeogenesis III | false | false | false | false | false | false | false | false | false | false | false | false | false | false | false | false | false |
| \|PWY-6640\| | salicylate degradation IV | false | false | false | false | false | false | false | false | false | false | false | false | false | false | false | false | false |
| \|PWY66-420\| | carnosine biosynthesis | false | false | false | false | false | false | false | false | false | false | false | false | false | false | false | false | false |
| \|PWY66-421\| | homocarnosine biosynthesis | false | false | false | false | false | false | false | false | false | false | false | false | false | false | false | false | false |
| \|PWY66-423\| | fructose 2,6-bisphosphate biosynthesis | false | false | false | false | false | false | false | false | false | false | false | false | false | false | false | false | false |
| \|PWY66-425\| | L-lysine degradation II (L-pipecolate pathway) | false | false | false | false | false | false | false | false | false | false | false | false | false | false | false | false | false |
| \|PWY66-426\| | hydrogen sulfide biosynthesis II (mammalian) | false | false | false | false | false | false | false | false | false | false | false | false | false | false | false | false | false |
| \|PWY66-428\| | L-threonine degradation V | false | false | false | false | false | false | false | false | false | false | false | false | false | false | false | false | false |
| \|PWY66-429\| | fatty acid biosynthesis initiation (mitochondria) | false | false | false | false | false | false | false | false | false | false | false | false | false | false | false | false | false |
| \|PWY-6642\| | (R)-cysteate degradation | false | false | false | false | false | false | false | false | false | false | false | false | false | false | false | false | false |
| \|PWY66-430\| | tetradecanoate biosynthesis (mitochondria) | false | false | false | false | false | false | false | false | false | false | false | false | false | false | false | false | false |
| \|PWY66-432\| | blood coagulation - tissue factor pathway | false | false | false | false | false | false | false | false | false | false | false | false | false | false | false | false | false |
| \|PWY-6643\| | coenzyme M biosynthesis II | false | false | false | false | false | false | false | false | false | false | false | false | false | false | false | false | false |
| \|PWY-6644\| | fluoroacetate and fluorothreonine biosynthesis | false | false | false | false | false | false | false | false | false | false | false | false | false | false | false | false | false |
| \|PWY-6645\| | labdane-type diterpenes biosynthesis | false | false | false | false | false | false | false | false | false | false | false | false | false | false | false | false | false |
| \|PWY-6646\| | fluoroacetate degradation | false | false | false | false | false | false | false | false | false | false | false | false | false | false | false | false | false |
| \|PWY-6648\| | rhamnolipid biosynthesis | false | false | false | false | false | false | false | false | false | false | false | false | false | false | false | false | false |
| \|PWY-6649\| | glycolate and glyoxylate degradation III | false | false | false | false | false | false | false | false | false | false | false | false | false | false | false | false | false |
| \|PWY-6650\| | juvenile hormone III biosynthesis II | false | false | false | false | false | false | false | false | false | false | false | false | false | false | false | false | false |
| \|PWY-6653\| | ent -kaurene biosynthesis II | false | false | false | false | false | false | false | false | false | false | false | false | false | false | false | false | false |
| \|PWY-6654\| | phosphopantothenate biosynthesis III (archaea) | false | false | false | false | false | false | false | false | false | false | false | false | false | false | false | false | false |
| \|PWY-6655\| | xanthan biosynthesis | false | false | false | false | false | false | false | false | false | false | false | false | false | false | false | false | false |
| \|PWY-6657\| | polyhydroxydecanoate biosynthesis | false | false | false | false | false | false | false | false | false | false | false | false | false | false | false | false | false |
| \|PWY-6658\| | acetan biosynthesis | false | false | false | false | false | false | false | false | false | false | false | true | false | false | false | false | false |
| \|PWY-6659\| | fusicoccin A biosynthesis | false | false | false | false | false | false | false | false | false | false | false | false | false | false | false | false | false |
| \|PWY-6660\| | 2-heptyl-3-hydroxy-4(1H)-quinolone biosynthesis | false | false | false | false | false | false | false | false | false | false | false | false | false | false | false | false | false |
| \|PWY-6661\| | 4-hydroxy-2(1H)-quinolone biosynthesis | false | false | false | false | false | false | false | false | false | false | false | false | false | false | false | false | false |
| \|PWY-6664\| | di-myo-inositol phosphate biosynthesis | false | false | false | false | false | false | false | false | false | false | false | false | false | false | false | false | false |
| \|PWY-6665\| | pterostilbene biosynthesis | false | false | false | false | false | false | false | false | false | false | false | false | false | false | false | false | false |
| \|PWY6666-1\| | anandamide degradation | false | false | false | false | false | false | false | false | false | false | false | false | false | false | false | false | false |
| \|PWY6666-2\| | dopamine degradation | false | false | false | false | false | false | false | false | false | false | false | false | false | false | false | false | false |
| \|PWY-6666\| | pyocyanin biosynthesis | false | false | false | false | false | false | false | false | false | false | false | false | false | false | false | false | false |
| \|PWY-6667\| | resveratrol degradation | false | false | false | false | false | false | false | false | false | false | false | false | false | false | false | false | false |
| \|PWY-6668\| | (E,E)-4,8,12-trimethyltrideca-1,3,7,11-tetraene biosynthesis | false | false | false | false | false | false | false | false | false | false | false | false | false | false | false | false | false |
| \|PWY-6669\| | δ-guaiene biosynthesis | false | false | false | false | false | false | false | false | false | false | false | false | false | false | false | false | false |
| \|PWY-6670\| | citronellol degradation | false | false | false | false | false | false | false | false | false | false | false | false | false | false | false | false | false |
| \|PWY-6672\| | cis-geranyl-CoA degradation | false | false | false | false | false | false | false | false | false | false | false | false | false | false | false | false | false |
| \|PWY-6673\| | caffeoylglucarate biosynthesis | false | false | false | false | false | false | false | false | false | false | false | false | false | false | false | false | false |
| \|PWY-6675\| | sulfur oxidation IV (intracellular sulfur) | false | false | false | false | false | false | false | false | false | false | false | false | false | false | false | false | false |
| \|PWY-6677\| | thiosulfate oxidation IV (multienzyme complex) | false | false | false | false | false | false | false | false | false | false | false | false | false | false | false | false | false |
| \|PWY-6678\| | geraniol and nerol degradation | false | false | false | false | false | false | false | false | false | false | false | false | false | false | false | false | false |
| \|PWY-6679\| | jadomycin biosynthesis | false | false | false | false | false | false | false | false | false | false | false | false | false | false | false | false | false |
| \|PWY-6681\| | neurosporaxanthin biosynthesis | false | false | false | false | false | false | false | false | false | false | false | false | false | false | false | false | false |
| \|PWY-6682\| | dehydrophos biosynthesis | false | false | false | false | false | false | false | false | false | false | false | false | false | false | false | false | false |
| \|PWY-6683\| | assimilatory sulfate reduction III | false | false | false | false | false | false | false | false | false | false | false | false | false | false | false | false | false |
| \|PWY-6684\| | aromatic glucosinolate activation | false | false | false | false | false | false | false | false | false | false | false | false | false | false | false | false | false |
| \|PWY-6685\| | glucosylglycerate biosynthesis II | false | false | false | false | false | false | false | false | false | false | false | false | false | false | false | false | false |
| \|PWY-6686\| | mannosylglucosylglycerate biosynthesis I | false | false | false | false | false | false | false | false | false | false | false | false | false | false | false | false | false |
| \|PWY-6687\| | mannosylglucosylglycerate biosynthesis II | false | false | false | false | false | false | false | false | false | false | false | false | false | false | false | false | false |
| \|PWY-6688\| | thyronamine and iodothyronamine metabolism | false | false | false | false | false | false | false | false | false | false | false | false | false | false | false | false | false |
| \|PWY-6690\| | cinnamate and 3-hydroxycinnamate degradation to 2-hydroxypentadienoate | false | false | false | false | false | false | false | false | false | false | false | false | false | false | false | false | false |
| \|PWY-6691\| | plaunotol biosynthesis | false | false | false | false | false | false | false | false | false | false | false | false | false | false | false | false | false |
| \|PWY-6692\| | Fe(II) oxidation | false | false | false | false | false | false | false | false | false | false | false | false | false | false | false | false | false |
| \|PWY-6693\| | D-galactose degradation IV | false | false | false | false | false | false | false | false | false | false | false | false | false | false | false | false | false |
| \|PWY-6695\| | oxalate degradation II | false | false | false | false | false | false | false | false | false | false | false | false | false | false | false | false | false |
| \|PWY-6696\| | oxalate degradation III | false | false | false | false | false | false | false | false | false | false | false | false | false | false | false | false | false |
| \|PWY-6698\| | oxalate degradation V | false | false | false | false | false | false | false | false | false | false | false | false | false | false | false | false | false |
| \|PWY-6699\| | oxalate biosynthesis | false | false | false | false | false | false | false | false | false | false | false | false | false | false | false | false | false |
| \|PWY-66\| | GDP-L-fucose biosynthesis I (from GDP-D-mannose) | false | false | false | false | false | false | false | false | false | false | false | false | false | false | false | false | false |
| \|PWY-6700\| | queuosine biosynthesis I (de novo) | false | false | false | false | false | false | false | false | false | false | false | false | false | false | false | false | false |
| \|PWY-6703\| | preQ0 biosynthesis | false | false | false | false | false | false | false | false | false | false | false | false | false | false | false | false | false |
| \|PWY-6704\| | L-ascorbate degradation IV | false | false | false | false | false | false | false | false | false | false | false | false | false | false | false | false | false |
| \|PWY-6707\| | gallate biosynthesis | false | false | false | false | false | false | false | false | false | false | false | false | false | false | false | false | false |
| \|PWY-6708\| | ubiquinol-8 biosynthesis (early decarboxylation) | false | false | false | false | false | false | false | false | false | false | false | false | false | false | false | false | false |
| \|PWY-6710\| | poly-hydroxy fatty acids biosynthesis | false | false | false | false | false | false | false | false | false | false | false | false | false | false | false | false | false |
| \|PWY-6711\| | archaeosine biosynthesis I | false | false | false | false | false | false | false | false | false | false | false | false | false | false | false | false | false |
| \|PWY-6713\| | L-rhamnose degradation II | false | false | false | false | false | false | false | false | false | false | false | false | false | false | false | false | false |
| \|PWY-6714\| | L-rhamnose degradation III | false | false | false | false | false | false | false | false | false | false | false | false | false | false | false | false | false |
| \|PWY-6717\| | (1,4)-β-D-xylan degradation | false | false | false | false | false | false | false | false | false | false | false | false | false | false | false | false | false |
| \|PWY-6718\| | sulfoacetaldehyde degradation III | false | false | false | false | false | false | false | false | false | false | false | false | false | false | false | false | false |
| \|PWY-6720\| | toyocamycin biosynthesis | false | false | false | false | false | false | false | false | false | false | false | false | true | false | false | true | false |
| \|PWY-6721\| | sangivamycin biosynthesis | false | false | false | false | false | false | false | false | false | false | false | false | false | false | false | false | false |
| \|PWY-6722\| | candicidin biosynthesis | false | false | false | false | false | false | false | false | false | false | false | false | false | false | false | false | false |
| \|PWY-6724\| | starch degradation II | false | false | false | false | false | false | false | false | false | false | false | false | false | false | false | false | false |
| \|PWY-6728\| | methylaspartate cycle | false | false | false | false | false | false | false | false | false | false | false | false | false | false | false | false | false |
| \|PWY-6730\| | methylhalides biosynthesis (plants) | false | false | false | false | false | false | false | false | false | false | false | false | false | false | false | false | false |
| \|PWY-6731\| | starch degradation III | false | false | false | false | false | false | false | false | false | false | false | false | false | false | true | false | false |
| \|PWY-6733\| | sporopollenin precursors biosynthesis | false | false | false | false | false | false | false | false | false | false | false | false | false | false | false | false | false |
| \|PWY-6735\| | starch degradation IV | false | false | false | false | false | false | false | false | false | false | false | false | false | false | false | false | false |
| \|PWY-6736\| | sulfur volatiles biosynthesis | false | false | false | false | false | false | false | false | false | false | false | false | false | false | false | false | false |
| \|PWY-6737\| | starch degradation V | false | false | false | false | false | false | false | false | false | false | false | false | false | false | false | false | false |
| \|PWY-6738\| | pinitol biosynthesis I | false | false | false | false | false | false | false | false | false | false | false | false | false | false | false | false | false |
| \|PWY-6739\| | pinitol biosynthesis II | false | false | false | false | false | false | false | false | false | false | false | false | false | false | false | false | false |
| \|PWY-6742\| | methane oxidation to methanol II | false | false | false | false | false | false | false | false | false | false | false | false | false | false | false | false | false |
| \|PWY-6745\| | phytochelatins biosynthesis | false | false | false | false | false | false | false | false | false | false | false | false | false | false | false | false | false |
| \|PWY-6748\| | nitrate reduction VII (denitrification) | false | false | false | false | false | false | false | false | false | false | false | false | false | false | false | false | false |
| \|PWY-6749\| | CMP-legionaminate biosynthesis I | false | false | false | false | false | false | false | false | false | false | false | false | false | false | false | false | false |
| \|PWY-6752\| | ortho-quinones biosynthesis | false | false | false | false | false | false | false | false | false | false | false | false | false | false | false | false | false |
| \|PWY-6754\| | S-methyl-5'-thioadenosine degradation I | false | false | false | false | false | false | false | false | false | false | false | false | false | false | false | false | false |
| \|PWY-6756\| | S-methyl-5'-thioadenosine degradation II | false | false | false | false | false | false | false | false | false | false | false | false | false | false | false | false | false |
| \|PWY-6759\| | hydrogen production III | false | false | false | false | false | false | false | false | false | false | false | false | false | false | false | false | false |
| \|PWY-6760\| | D-xylose degradation III | false | false | false | false | false | false | false | false | false | false | false | false | false | false | false | false | false |
| \|PWY-6762\| | salicylate glucosides biosynthesis IV | false | false | false | false | false | false | false | false | false | false | false | false | false | false | false | false | false |
| \|PWY-6763\| | salicortin biosynthesis | false | false | false | false | false | false | false | false | false | false | false | false | false | false | false | false | false |
| \|PWY-6765\| | hydrogen production IV | false | false | false | false | false | false | false | false | false | false | false | false | false | false | false | false | false |
| \|PWY-6766\| | salicin biosynthesis | false | false | false | false | false | false | false | false | false | false | false | false | false | false | false | false | false |
| \|PWY-6767\| | 4,4'-diapolycopenedioate biosynthesis | false | false | false | false | false | false | false | false | false | false | false | false | false | false | false | false | false |
| \|PWY-6769\| | rhamnogalacturonan type I degradation I | false | false | false | false | false | false | false | false | false | false | false | false | false | false | false | false | false |
| \|PWY-6771\| | rhamnogalacturonan type I degradation II (bacteria) | false | false | false | false | false | false | false | false | false | false | false | false | false | false | false | false | false |
| \|PWY-6772\| | hydrogen production V | false | false | false | false | false | false | false | false | false | false | false | false | false | false | false | false | false |
| \|PWY-6773\| | 1,3-β-D-glucan biosynthesis | false | false | false | false | false | false | false | false | false | false | false | false | false | false | false | false | false |
| \|PWY-6778\| | laminaribiose degradation | false | false | false | false | false | false | false | false | false | false | false | false | false | false | false | false | false |
| \|PWY-6780\| | hydrogen production VI | false | false | false | false | false | false | false | false | false | false | false | false | false | false | false | false | false |
| \|PWY-6781\| | chlorogenic acid degradation | false | false | false | false | false | false | false | false | false | false | false | false | false | false | false | false | false |
| \|PWY-6784\| | cellulose and hemicellulose degradation (cellulolosome) | false | false | false | false | false | false | false | false | false | false | false | false | false | false | false | false | false |
| \|PWY-6785\| | hydrogen production VIII | false | false | false | false | false | false | false | false | false | false | false | false | false | false | false | false | false |
| \|PWY-6786\| | detoxification of reactive carbonyls in chloroplasts | false | false | false | false | false | false | false | false | false | false | false | false | false | false | false | false | false |
| \|PWY-6787\| | flavonoid biosynthesis (in equisetum) | false | false | false | false | false | false | false | false | false | false | false | false | false | false | false | false | false |
| \|PWY-6788\| | cellulose degradation II (fungi) | false | false | false | false | false | false | false | false | false | false | false | false | false | false | false | false | false |
| \|PWY-6789\| | (1,3)-β-D-xylan degradation | false | false | false | false | false | false | false | false | false | false | false | false | true | true | true | true | true |
| \|PWY-6790\| | L-arabinan degradation | false | false | false | false | false | false | false | false | false | false | false | false | false | false | false | false | false |
| \|PWY-6791\| | xyloglucan degradation I (endoglucanase) | false | false | false | false | false | false | false | false | false | false | false | false | false | false | false | false | false |
| \|PWY-6792\| | scopoletin biosynthesis | false | false | false | false | false | false | false | false | false | false | false | false | false | false | false | false | false |
| \|PWY-6794\| | adenosine 5'-phosphoramidate biosynthesis | false | false | false | false | false | false | false | false | false | false | false | false | false | false | false | false | false |
| \|PWY-6795\| | diacylglyceryl-N,N,N-trimethylhomoserine biosynthesis | false | false | false | false | false | false | false | false | false | false | false | false | false | false | false | false | false |
| \|PWY-6797\| | 6-hydroxymethyl-dihydropterin diphosphate biosynthesis II (Methanocaldococcus) | false | false | false | false | false | false | false | false | false | false | false | false | false | false | false | false | false |
| \|PWY-6799\| | fatty acid biosynthesis initiation (plant mitochondria) | false | false | false | false | false | false | false | false | false | false | false | false | false | false | false | false | false |
| \|PWY-6801\| | volatile esters biosynthesis (during fruit ripening) | false | false | false | false | false | false | false | false | false | false | false | false | false | false | false | false | false |
| \|PWY-6802\| | salidroside biosynthesis | false | false | false | false | false | false | false | false | false | false | false | false | false | false | false | false | false |
| \|PWY-6803\| | phosphatidylcholine acyl editing | false | false | false | false | false | false | false | false | false | false | false | false | false | false | false | false | false |
| \|PWY-6804\| | diacylglycerol biosynthesis (PUFA enrichment in oilseed) | false | false | false | false | false | false | false | false | false | false | false | false | false | false | false | false | false |
| \|PWY-6805\| | cellulose degradation I (cellulosome) | false | false | false | false | false | false | false | false | false | false | false | false | false | false | false | false | false |
| \|PWY-6806\| | carotenoid cleavage | false | false | false | false | false | false | false | false | false | false | false | false | false | false | false | false | false |
| \|PWY-6807\| | xyloglucan degradation II (exoglucanase) | false | false | false | false | false | false | false | false | false | false | false | false | false | false | false | false | false |
| \|PWY-6808\| | dTDP-α-D-forosamine biosynthesis | false | false | false | false | false | false | false | false | false | false | false | true | false | false | false | false | false |
| \|PWY-6809\| | neoxanthin biosynthesis | false | false | false | false | false | false | false | false | false | false | false | false | false | false | false | false | false |
| \|PWY-6812\| | xyloglucan degradation III (cellobiohydrolase) | false | false | false | false | false | false | false | false | false | false | false | false | false | false | false | false | false |
| \|PWY-6813\| | glucuronoarabinoxylan degradation | false | false | false | false | false | false | false | false | false | false | false | false | false | false | false | false | false |
| \|PWY-6815\| | porphyran degradation | false | false | false | false | false | false | false | false | false | false | false | false | false | false | false | false | false |
| \|PWY-6816\| | agarose degradation | false | false | false | false | false | false | false | false | false | false | false | false | false | false | false | false | false |
| \|PWY-6817\| | λ-carrageenan degradation | false | false | false | false | false | false | false | false | false | false | false | false | false | false | false | false | false |
| \|PWY-6818\| | ornithine lipid biosynthesis | false | false | false | false | false | false | false | false | false | false | false | false | false | false | false | false | false |
| \|PWY-681\| | dibenzothiophene desulfurization | false | false | false | false | false | false | false | false | false | false | false | false | false | false | false | false | false |
| \|PWY-6820\| | diphyllin biosynthesis | false | false | false | false | false | false | false | false | false | false | false | false | false | false | false | false | false |
| \|PWY-6821\| | κ-carrageenan degradation | false | false | false | false | false | false | false | false | false | false | false | false | false | false | false | false | false |
| \|PWY-6822\| | ι-carrageenan degradation | false | false | false | false | false | false | false | false | false | false | false | false | false | false | false | false | false |
| \|PWY-6823\| | molybdopterin biosynthesis | false | false | false | false | false | false | false | false | false | false | false | false | false | false | false | false | false |
| \|PWY-6824\| | justicidin B biosynthesis | false | false | false | false | false | false | false | false | false | false | false | false | false | false | false | false | false |
| \|PWY-6825\| | phosphatidylcholine biosynthesis V | false | false | false | false | false | false | false | false | false | false | false | false | false | false | false | false | false |
| \|PWY-6826\| | phosphatidylcholine biosynthesis VI | false | false | false | false | false | false | false | false | false | false | false | false | false | false | false | false | false |
| \|PWY-6827\| | gellan degradation | false | false | false | false | false | false | false | false | false | false | false | false | false | false | false | false | false |
| \|PWY-6829\| | tRNA methylation (yeast) | false | false | false | false | false | false | false | false | false | false | false | false | false | false | false | false | false |
| \|PWY-6831\| | pyrrolnitrin biosynthesis | false | false | false | false | false | false | false | false | false | false | false | false | false | false | false | false | false |
| \|PWY-6832\| | 2-aminoethylphosphonate degradation II | false | false | false | false | false | false | false | false | false | false | false | false | false | false | false | false | false |
| \|PWY-6834\| | spermidine biosynthesis III | false | false | false | false | false | false | false | false | false | false | false | false | false | false | false | false | false |
| \|PWY-6835\| | 6-gingerol biosynthesis | false | false | false | false | false | false | false | false | false | false | false | false | false | false | false | false | false |
| \|PWY-6836\| | santalene biosynthesis II | false | false | false | false | false | false | false | false | false | false | false | false | false | false | false | false | false |
| \|PWY-6837\| | fatty acid β-oxidation V (unsaturated, odd number, di-isomerase-dependent) | false | false | false | false | false | false | false | false | false | false | false | false | false | false | false | false | false |
| \|PWY-6839\| | 2-aminoethylphosphonate biosynthesis | false | false | false | false | false | false | false | false | false | false | false | false | false | false | false | false | false |
| \|PWY-6840\| | homoglutathione biosynthesis | false | false | false | false | false | false | false | false | false | false | false | false | false | false | false | false | false |
| \|PWY-6841\| | homophytochelatin biosynthesis | false | false | false | false | false | false | false | false | false | false | false | false | false | false | false | false | false |
| \|PWY-6842\| | glutathione-mediated detoxification II | false | false | false | false | false | false | false | false | false | false | false | false | false | false | false | false | false |
| \|PWY-6845\| | nitric oxide biosynthesis I (plants) | false | false | false | false | false | false | false | false | false | false | false | false | false | false | false | false | false |
| \|PWY-6848\| | rutin degradation | false | false | false | false | false | false | false | false | false | false | false | false | false | false | false | false | false |
| \|PWY-6852\| | senecionine N-oxide biosynthesis | false | false | false | false | false | false | false | false | false | false | false | false | false | false | false | false | false |
| \|PWY-6853\| | ethene biosynthesis II (microbes) | false | false | false | false | false | false | false | false | false | false | false | false | false | false | false | false | false |
| \|PWY-6854\| | ethene biosynthesis III (microbes) | false | false | false | false | false | false | false | false | false | false | false | false | false | false | false | false | false |
| \|PWY-6855\| | chitin degradation I (archaea) | false | false | false | false | false | false | false | false | false | false | false | false | false | false | false | false | false |
| \|PWY-6857\| | retinol biosynthesis | false | false | false | false | false | false | false | false | false | false | false | false | false | false | false | false | false |
| \|PWY-6859\| | all-trans-farnesol biosynthesis | false | false | false | false | false | false | false | false | false | false | false | false | false | false | false | false | false |
| \|PWY-6861\| | the visual cycle I (vertebrates) | false | false | false | false | false | false | false | false | false | false | false | false | false | false | false | false | false |
| \|PWY-6863\| | pyruvate fermentation to hexanol (engineered) | false | false | false | false | false | false | false | false | false | false | false | false | false | false | false | false | false |
| \|PWY-6872\| | retinoate biosynthesis I | false | false | false | false | false | false | false | false | false | false | false | false | false | false | false | false | false |
| \|PWY-6873\| | long chain fatty acid ester synthesis (engineered) | false | false | false | false | false | false | false | false | false | false | false | false | false | false | false | false | false |
| \|PWY-6875\| | retinoate biosynthesis II | false | false | false | false | false | false | false | false | false | false | false | false | false | false | false | false | false |
| \|PWY-6876\| | isopropanol biosynthesis (engineered) | false | false | false | false | false | false | false | false | false | false | false | false | false | false | false | false | false |
| \|PWY-6883\| | pyruvate fermentation to butanol II (engineered) | false | false | false | false | false | false | false | false | false | false | false | false | false | false | false | false | false |
| \|PWY-6888\| | zealexin biosynthesis | false | false | false | false | false | false | false | false | false | false | false | false | false | false | false | false | false |
| \|PWY-6891\| | thiazole component of thiamine diphosphate biosynthesis II | false | false | false | false | false | false | false | false | false | false | false | false | false | false | false | false | false |
| \|PWY-6892\| | thiazole component of thiamine diphosphate biosynthesis I | false | false | false | false | false | false | false | false | false | false | false | false | false | false | false | false | false |
| \|PWY-6893\| | thiamine diphosphate biosynthesis II (Bacillus) | false | false | false | false | false | false | false | false | false | false | false | false | false | false | false | false | false |
| \|PWY-6894\| | thiamine diphosphate biosynthesis I (E. coli) | false | false | false | false | false | false | false | false | false | false | false | false | false | false | false | false | false |
| \|PWY-6896\| | thiamine diphosphate salvage I | false | false | false | false | false | false | false | false | false | false | false | false | false | false | false | false | false |
| \|PWY-6898\| | thiamine diphosphate salvage III | false | false | false | false | false | false | false | false | false | false | false | false | false | false | false | false | false |
| \|PWY-6899\| | base-degraded thiamine salvage | false | false | false | false | false | false | false | false | false | false | false | false | false | false | false | false | false |
| \|PWY-6900\| | (Z)-butanethial-S-oxide biosynthesis | false | false | false | false | false | false | false | false | false | false | false | false | false | false | false | false | false |
| \|PWY-6902\| | chitin degradation II (Vibrio) | false | false | false | false | false | false | false | false | false | false | false | false | true | true | true | true | true |
| \|PWY-6906\| | chitin derivatives degradation | false | false | false | false | false | false | false | false | false | false | false | false | false | false | false | false | false |
| \|PWY-6907\| | thiamine diphosphate biosynthesis III (Staphylococcus) | false | false | false | false | false | false | false | false | false | false | false | false | false | false | false | false | false |
| \|PWY-6908\| | thiamine diphosphate biosynthesis IV (eukaryotes) | false | false | false | false | false | false | false | false | false | false | false | false | false | false | false | false | false |
| \|PWY-6909\| | thiazole component of thiamine diphosphate biosynthesis III | false | false | false | false | false | false | false | false | false | false | false | false | false | false | false | false | false |
| \|PWY-6913\| | methylbutenol biosynthesis | false | false | false | false | false | false | false | false | false | false | false | false | false | false | false | false | false |
| \|PWY-6914\| | sophoraflavanone G biosynthesis | false | false | false | false | false | false | false | false | false | false | false | false | false | false | false | false | false |
| \|PWY-6915\| | pentalenolactone biosynthesis | false | false | false | false | false | false | false | false | false | false | false | false | true | true | true | true | false |
| \|PWY-6917\| | vernolate biosynthesis III | false | false | false | false | false | false | false | false | false | false | false | false | false | false | false | false | false |
| \|PWY-6919\| | neopentalenoketolactone and pentalenate biosynthesis | false | false | false | false | false | false | false | false | false | false | false | false | false | false | false | false | false |
| \|PWY-6920\| | 6-gingerol analog biosynthesis (engineered) | false | false | false | false | false | false | false | false | false | false | false | false | false | false | false | false | false |
| \|PWY-6922\| | L-Nδ-acetylornithine biosynthesis | false | false | false | false | false | false | false | false | false | false | false | false | false | false | false | false | false |
| \|PWY-6923\| | ricinine degradation | false | false | false | false | false | false | false | false | false | false | false | false | false | false | false | false | false |
| \|PWY-6926\| | pyrethrin I biosynthesis | false | false | false | false | false | false | false | false | false | false | false | false | false | false | false | false | false |
| \|PWY-6927\| | chlorophyll a degradation II | false | false | false | false | false | false | false | false | false | false | false | false | false | false | false | false | false |
| \|PWY-6930\| | phenolic malonylglucosides biosynthesis | false | false | false | false | false | false | false | false | false | false | false | false | false | false | false | false | false |
| \|PWY-6932\| | selenate reduction | false | false | false | false | false | false | false | false | false | false | false | false | false | false | false | false | false |
| \|PWY-6933\| | seleno-amino acid detoxification and volatilization III | false | false | false | false | false | false | false | false | false | false | false | false | false | false | false | false | false |
| \|PWY-6935\| | seleno-amino acid detoxification and volatilization II | false | false | false | false | false | false | false | false | false | false | false | false | false | false | false | false | false |
| \|PWY-6938\| | NADH repair (eukaryotes) | false | false | false | false | false | false | false | false | false | false | false | false | false | false | false | false | false |
| \|PWY-6940\| | icosapentaenoate biosynthesis VI (fungi) | false | false | false | false | false | false | false | false | false | false | false | false | false | false | false | false | false |
| \|PWY-6941\| | styrene degradation | false | false | false | false | false | false | false | false | false | false | false | false | false | false | false | false | false |
| \|PWY-6942\| | dTDP-D-desosamine biosynthesis | false | false | false | false | false | false | false | false | false | false | false | false | false | false | false | false | false |
| \|PWY-6943\| | testosterone and androsterone degradation to androstendione (aerobic) | false | false | false | false | false | false | false | false | false | false | false | false | false | false | false | false | false |
| \|PWY-6944\| | androstenedione degradation I (aerobic) | false | false | false | false | false | false | false | false | false | false | false | false | false | false | false | false | false |
| \|PWY-6945\| | cholesterol degradation to androstenedione I (cholesterol oxidase) | false | false | false | false | false | false | false | false | false | false | false | false | false | false | false | false | false |
| \|PWY-6946\| | cholesterol degradation to androstenedione II (cholesterol dehydrogenase) | false | false | false | false | false | false | false | false | false | false | false | false | false | false | false | false | false |
| \|PWY-6948\| | sitosterol degradation to androstenedione | false | false | false | false | false | false | false | false | false | false | false | false | false | false | false | false | false |
| \|PWY-6949\| | DIBOA-glucoside biosynthesis | false | false | false | false | false | false | false | false | false | false | false | false | false | false | false | false | false |
| \|PWY-6950\| | DIMBOA-glucoside biosynthesis | false | false | false | false | false | false | false | false | false | false | false | false | false | false | false | false | false |
| \|PWY-6952\| | glycerophosphodiester degradation | false | false | false | false | false | false | false | false | false | false | false | false | false | false | false | false | false |

| | | | | | | | | | | | | | | | | | | |
|---|---|---|---|---|---|---|---|---|---|---|---|---|---|---|---|---|---|---|
| \|PWY-6953\| | dTDP-3-acetamido-α-D-fucose biosynthesis | false | false | false | false | false | false | false | false | false | false | false | false | false | false | false | false | false |
| \|PWY-6955\| | lincomycin A biosynthesis | false | false | false | false | false | false | false | false | false | false | false | false | false | false | false | false | false |
| \|PWY-6958\| | icosapentaenoate biosynthesis I (lower eukaryotes) | false | false | false | false | false | false | false | false | false | false | false | false | false | false | false | false | false |
| \|PWY-6959\| | L-ascorbate degradation V | false | false | false | false | false | false | false | false | false | false | false | false | false | false | false | false | false |
| \|PWY-695\| | abscisic acid biosynthesis | false | false | false | false | false | false | false | false | false | false | false | false | false | false | false | false | false |
| \|PWY-6960\| | L-ascorbate degradation III | false | false | false | false | false | false | false | false | false | false | false | false | false | false | false | false | false |
| \|PWY-6961\| | L-ascorbate degradation II (bacterial, aerobic) | false | false | false | false | false | false | false | false | false | false | false | false | false | false | false | false | false |
| \|PWY-6964\| | ammonia assimilation cycle II | false | false | false | false | false | false | false | false | false | false | false | false | false | false | false | false | false |
| \|PWY-6965\| | methylamine degradation II | false | false | false | false | false | false | false | false | false | false | false | false | false | false | false | false | false |
| \|PWY-6966\| | methanol oxidation to formaldehyde I | false | false | false | false | false | false | false | false | false | false | false | false | false | false | false | false | false |
| \|PWY-6967\| | methylamine degradation I | false | false | false | false | false | false | false | false | false | false | false | false | false | false | false | false | false |
| \|PWY-6968\| | trimethylamine degradation | false | false | false | false | false | false | false | false | false | false | false | false | false | false | false | false | false |
| \|PWY-6969\| | TCA cycle V (2-oxoglutarate synthase) | false | false | false | false | false | false | false | false | false | false | false | false | false | false | false | false | false |
| \|PWY-6971\| | oleandomycin biosynthesis | false | false | false | false | false | false | false | false | false | false | false | false | false | false | false | false | false |
| \|PWY-6972\| | oleandomycin activation/inactivation | false | false | false | false | false | false | false | false | false | false | false | false | false | false | false | false | false |
| \|PWY-6973\| | dTDP-α-D-olivose, dTDP-α-D-oliose and dTDP-α-D-mycarose biosynthesis | false | false | false | false | false | false | false | false | false | false | false | false | false | false | false | false | false |
| \|PWY-6974\| | dTDP-β-L-olivose biosynthesis | false | false | false | false | false | false | false | false | false | false | false | false | false | false | false | false | false |
| \|PWY-6976\| | dTDP-β-L-mycarose biosynthesis | false | false | false | false | false | false | false | false | false | false | false | false | false | false | false | false | false |
| \|PWY-6978\| | plastoquinol-9 biosynthesis II | false | false | false | false | false | false | false | false | false | false | false | false | false | false | false | false | false |
| \|PWY-6982\| | umbelliferone biosynthesis | false | false | false | false | false | false | false | false | false | false | false | false | false | false | false | false | false |
| \|PWY-6983\| | threo-tetrahydrobiopterin biosynthesis | false | false | false | false | false | false | false | false | false | false | false | false | false | false | false | false | false |
| \|PWY-6984\| | lipoate salvage II | false | false | false | false | false | false | false | false | false | false | false | false | false | false | false | false | false |
| \|PWY-6986\| | alginate degradation | false | false | false | false | false | false | false | false | false | false | false | false | false | false | false | false | false |
| \|PWY-6987\| | lipoate biosynthesis and incorporation III (Bacillus) | false | false | false | false | false | false | false | false | false | false | false | false | false | false | false | false | false |
| \|PWY-6989\| | (S)-camphor degradation | false | false | false | false | false | false | false | false | false | false | false | false | false | false | false | false | false |
| \|PWY-6990\| | (+)-camphor biosynthesis | false | false | false | false | false | false | false | false | false | false | false | false | false | false | false | false | false |
| \|PWY-6991\| | (-)-camphor biosynthesis | false | false | false | false | false | false | false | false | false | false | false | false | false | false | false | false | false |
| \|PWY-6992\| | 1,5-anhydrofructose degradation | false | false | false | false | false | false | false | false | false | false | false | false | false | false | false | false | false |
| \|PWY-6993\| | nicotine degradation II (pyrrolidine pathway) | false | false | false | false | false | false | false | false | false | false | false | false | false | false | false | false | false |
| \|PWY-6994\| | L-pyrrolysine biosynthesis | false | false | false | false | false | false | false | false | false | false | false | false | false | false | false | false | false |
| \|PWY-6995\| | 5-hydroxymethylfurfural degradation | false | false | false | false | false | false | false | false | false | false | false | false | false | false | false | false | false |
| \|PWY-6996\| | daidzin and daidzein degradation | false | false | false | false | false | false | false | false | false | false | false | false | false | false | false | false | false |
| \|PWY-6997\| | furfural degradation | false | false | false | false | false | false | false | false | false | false | false | false | false | false | false | false | false |
| \|PWY-6998\| | CDP-D-arabitol biosynthesis | false | false | false | false | false | false | false | false | false | false | false | false | false | false | false | false | false |
| \|PWY-6999\| | theophylline degradation | false | false | false | false | false | false | false | false | false | false | false | false | false | false | false | false | false |
| \|PWY-699\| | brassinolide biosynthesis I | false | false | false | false | false | false | false | false | false | false | false | false | false | false | false | false | false |
| \|PWY-6\| | GDP-L-fucose biosynthesis II (from L-fucose) | false | false | false | false | false | false | false | false | false | false | false | false | false | false | false | false | false |
| \|PWY-7002\| | 4-hydroxyacetophenone degradation | false | false | false | false | false | false | false | false | false | false | false | false | false | false | false | false | false |
| \|PWY-7006\| | 4-amino-3-hydroxybenzoate degradation | false | false | false | false | false | false | false | false | false | false | false | false | false | false | false | false | false |
| \|PWY-7007\| | methyl ketone biosynthesis (engineered) | false | false | false | false | false | false | false | false | false | false | false | false | false | false | false | false | false |
| \|PWY-7008\| | 2-hydroxybiphenyl degradation | false | false | false | false | false | false | false | false | false | false | false | false | false | false | false | false | false |
| \|PWY-7009\| | 2,2'-dihydroxybiphenyl degradation | false | false | false | false | false | false | false | false | false | false | false | false | false | false | false | false | false |
| \|PWY-7010\| | 2-propylphenol degradation | false | false | false | false | false | false | false | false | false | false | false | false | false | false | false | false | false |
| \|PWY-7011\| | 2-isopropylphenol degradation | false | false | false | false | false | false | false | false | false | false | false | false | false | false | false | false | false |
| \|PWY-7013\| | (S)-propane-1,2-diol degradation | false | false | false | false | false | false | false | false | false | false | false | false | false | false | false | false | false |
| \|PWY-7014\| | paromamine biosynthesis I | false | false | false | false | false | false | false | false | false | false | false | false | false | false | false | false | false |
| \|PWY-7015\| | ribostamycin biosynthesis | false | false | false | false | false | false | false | false | false | false | false | false | false | false | false | false | false |
| \|PWY-7016\| | neomycin biosynthesis | false | false | false | false | false | false | false | false | false | false | false | false | false | false | false | false | false |
| \|PWY-7018\| | paromomycin biosynthesis | false | false | false | false | false | false | false | false | false | false | false | false | false | false | false | false | false |
| \|PWY-7019\| | butirosin biosynthesis | false | false | false | false | false | false | false | false | false | false | false | false | false | false | false | false | false |
| \|PWY-7022\| | paromamine biosynthesis II | false | false | false | false | false | false | false | false | false | false | false | false | false | false | false | false | false |
| \|PWY-7025\| | gentamicin biosynthesis | false | false | false | false | false | false | false | false | false | false | false | false | false | false | false | false | false |
| \|PWY-7026\| | terminal olefins biosynthesis I | false | false | false | false | false | false | false | false | false | false | false | false | false | false | false | false | false |
| \|PWY-7027\| | hentriaconta-3,6,9,12,15,19,22,25,28-nonaene biosynthesis | false | false | false | false | false | false | false | false | false | false | false | false | false | false | false | false | false |
| \|PWY-7028\| | UDP-N,N'-diacetylbacillosamine biosynthesis | false | false | false | false | false | false | false | false | false | false | false | false | false | false | false | false | false |
| \|PWY-7029\| | terminal olefins biosynthesis II | false | false | false | false | false | false | false | false | false | false | false | false | false | false | false | false | false |
| \|PWY-7031\| | protein N-glycosylation (bacterial) | false | false | false | false | false | false | false | false | false | false | false | false | false | false | false | false | false |
| \|PWY-7032\| | alkane biosynthesis I | false | false | false | false | false | false | false | false | false | false | false | false | false | false | false | false | false |
| \|PWY-7033\| | alkane biosynthesis II | false | false | false | false | false | false | false | false | false | false | false | false | false | false | false | false | false |
| \|PWY-7035\| | (9Z)-tricosene biosynthesis | false | false | false | false | false | false | false | false | false | false | false | false | false | false | false | false | false |
| \|PWY-7036\| | very long chain fatty acid biosynthesis II | false | false | false | false | false | false | false | false | false | false | false | false | false | false | false | false | false |
| \|PWY-7037\| | protein O-glycosylation (Neisseria) | false | false | false | false | false | false | false | false | false | false | false | false | false | false | false | false | false |
| \|PWY-7039\| | phosphatidate metabolism, as a signaling molecule | false | false | false | false | false | false | false | false | false | false | false | false | false | false | false | false | false |
| \|PWY-7040\| | violacein biosynthesis | false | false | false | false | false | false | false | false | false | false | false | false | false | false | false | false | false |
| \|PWY-7041\| | the visual cycle II (molluscs) | false | false | false | false | false | false | false | false | false | false | false | false | false | false | false | false | false |
| \|PWY-7042\| | the visual cycle (insects) | false | false | false | false | false | false | false | false | false | false | false | false | false | false | false | false | false |
| \|PWY-7043\| | 11-cis-3-hydroxyretinal biosynthesis | false | false | false | false | false | false | false | false | false | false | false | false | false | false | false | false | false |
| \|PWY-7044\| | 5-nitroanthranilate degradation | false | false | false | false | false | false | false | false | false | false | false | false | false | false | false | false | false |
| \|PWY-7045\| | mithramycin biosynthesis | false | false | false | false | false | false | false | false | false | false | false | true | false | false | false | false | false |
| \|PWY-7046\| | 4-coumarate degradation (anaerobic) | false | false | false | false | false | false | false | false | false | false | false | false | false | false | false | false | false |
| \|PWY-7049\| | icosapentaenoate biosynthesis II (6-desaturase, mammals) | false | false | false | false | false | false | false | false | false | false | false | false | false | false | false | false | false |
| \|PWY-7050\| | icosapentaenoate biosynthesis IV (bacteria) | false | false | false | false | false | false | false | false | false | false | false | false | false | false | false | false | false |
| \|PWY-7052\| | cyanophycin metabolism | false | false | false | false | false | false | false | false | false | false | false | false | false | false | false | false | false |
| \|PWY-7053\| | docosahexaenoate biosynthesis I (lower eukaryotes) | false | false | false | false | false | false | false | false | false | false | false | false | false | false | false | false | false |
| \|PWY-7054\| | N-acetylglutaminylglutamine amide biosynthesis | false | false | false | false | false | false | false | false | false | false | false | false | false | false | false | false | false |
| \|PWY-7055\| | daphnetin modification | false | false | false | false | false | false | false | false | false | false | false | false | false | false | false | false | false |
| \|PWY-7056\| | daphnin interconversion | false | false | false | false | false | false | false | false | false | false | false | false | false | false | false | false | false |
| \|PWY-7057\| | cichoriin interconversion | false | false | false | false | false | false | false | false | false | false | false | false | false | false | false | false | false |
| \|PWY-7058\| | esculetin modification | false | false | false | false | false | false | false | false | false | false | false | false | false | false | false | false | false |
| \|PWY-7059\| | fumigaclavine biosynthesis | false | false | false | false | false | false | false | false | false | false | false | false | false | false | false | false | false |
| \|PWY-7065\| | 2α,7β-dihydroxylation of taxusin | false | false | false | false | false | false | false | false | false | false | false | false | false | false | false | false | false |
| \|PWY-7066\| | glycyrrhetinate biosynthesis | false | false | false | false | false | false | false | false | false | false | false | false | false | false | false | false | false |
| \|PWY-7067\| | betulinate biosynthesis | false | false | false | false | false | false | false | false | false | false | false | false | false | false | false | false | false |
| \|PWY-7068\| | ursolate biosynthesis | false | false | false | false | false | false | false | false | false | false | false | false | false | false | false | false | false |
| \|PWY-7069\| | oleanolate biosynthesis | false | false | false | false | false | false | false | false | false | false | false | false | false | false | false | false | false |
| \|PWY-7070\| | steviol biosynthesis | false | false | false | false | false | false | false | false | false | false | false | false | false | false | false | false | false |
| \|PWY-7071\| | steviol glucoside biosynthesis (rebaudioside A biosynthesis) | false | false | false | false | false | false | false | false | false | false | false | false | false | false | false | false | false |
| \|PWY-7072\| | hopanoid biosynthesis (bacteria) | false | false | false | false | false | false | false | false | false | false | false | false | false | false | false | false | false |
| \|PWY-7074\| | phenylethanol glycoconjugate biosynthesis | false | false | false | false | false | false | false | false | false | false | false | false | false | false | false | false | false |
| \|PWY-7075\| | phenylethyl acetate biosynthesis | false | false | false | false | false | false | false | false | false | false | false | false | false | false | false | false | false |
| \|PWY-7076\| | 3,5-dimethoxytoluene biosynthesis | false | false | false | false | false | false | false | false | false | false | false | false | false | false | false | false | false |
| \|PWY-7077\| | N-acetyl-D-galactosamine degradation | false | false | false | false | false | false | false | false | false | false | false | false | false | false | false | false | false |
| \|PWY-7079\| | geodin biosynthesis | false | false | false | false | false | false | false | false | false | false | false | false | false | false | false | false | false |
| \|PWY-7080\| | asterrate biosynthesis | false | false | false | false | false | false | false | false | false | false | false | false | false | false | false | false | false |
| \|PWY-7081\| | 4-aminophenol degradation | false | false | false | false | false | false | false | false | false | false | false | false | false | false | false | false | false |
| \|PWY-7085\| | triethylamine degradation | false | false | false | false | false | false | false | false | false | false | false | false | false | false | false | false | false |
| \|PWY-7087\| | 2-methylisoborneol biosynthesis | false | false | false | false | false | false | false | false | false | false | false | false | false | false | false | false | false |
| \|PWY-7088\| | taxiphyllin biosynthesis | false | false | false | false | false | false | false | false | false | false | false | false | false | false | false | false | false |
| \|PWY-7089\| | taxiphyllin bioactivation | false | false | false | false | false | false | false | false | false | false | false | false | false | false | false | false | false |
| \|PWY-7091\| | linustatin bioactivation | false | false | false | false | false | false | false | false | false | false | false | false | false | false | false | false | false |
| \|PWY-7092\| | neolinustatin bioactivation | false | false | false | false | false | false | false | false | false | false | false | false | false | false | false | false | false |
| \|PWY-7093\| | vicianin bioactivation | false | false | false | false | false | false | false | false | false | false | false | false | false | false | false | false | false |
| \|PWY-7094\| | fatty acid salvage | false | false | false | false | false | false | false | false | false | false | false | false | false | false | false | false | false |
| \|PWY-7095\| | 3,4-dihydroxymandelonitrile β-D-glucose biosynthesis | false | false | false | false | false | false | false | false | false | false | false | false | false | false | false | false | false |
| \|PWY-7096\| | triclosan resistance | false | false | false | false | false | false | false | false | false | false | false | false | false | false | false | false | false |
| \|PWY-7097\| | vanillin and vanillate degradation I | false | false | false | false | false | false | false | false | false | false | false | false | false | false | false | false | false |
| \|PWY-7098\| | vanillin and vanillate degradation II | false | false | false | false | false | false | false | false | false | false | false | false | false | false | false | false | false |
| \|PWY-7100\| | spinosyn biosynthesis | false | false | false | false | false | false | false | false | false | false | false | false | false | false | false | false | false |
| \|PWY-7101\| | 5-deoxystrigol biosynthesis | false | false | false | false | false | false | false | false | false | false | false | false | false | false | false | false | false |
| \|PWY-7102\| | bisabolene biosynthesis (engineered) | false | false | false | false | false | false | false | false | false | false | false | false | false | false | false | false | false |
| \|PWY-7104\| | dTDP-β-L-megosamine biosynthesis | false | false | false | false | false | false | false | false | false | false | false | false | false | false | false | false | false |
| \|PWY-7105\| | olivetol biosynthesis | false | false | false | false | false | false | false | false | false | false | false | false | false | false | false | false | false |
| \|PWY-7106\| | erythromycin D biosynthesis | false | false | false | false | false | false | false | false | false | false | false | false | false | false | false | false | false |
| \|PWY-7108\| | erythromycin A biosynthesis | false | false | false | false | false | false | false | false | false | false | false | false | false | false | false | false | false |
| \|PWY-7109\| | megalomicin A biosynthesis | false | false | false | false | false | false | false | false | false | false | false | false | false | false | false | false | false |
| \|PWY-7112\| | 4-hydroxy-2-nonenal detoxification | false | false | false | false | false | false | false | false | false | false | false | false | false | false | false | false | false |
| \|PWY-7113\| | furcatin degradation | false | false | false | false | false | false | false | false | false | false | false | false | true | true | true | true | true |
| \|PWY-7114\| | tea aroma glycosidic precursor bioactivation | false | false | false | false | false | false | false | false | false | false | false | false | false | false | false | false | false |
| \|PWY-7118\| | chitin deacetylation | false | false | false | false | false | false | false | false | false | false | false | false | false | false | false | false | false |
| \|PWY-7119\| | ceramide and sphingolipid recycling and degradation (yeast) | false | false | false | false | false | false | false | false | false | false | false | false | false | false | false | false | false |
| \|PWY-7120\| | esterified suberin biosynthesis | false | false | false | false | false | false | false | false | false | false | false | false | false | false | false | false | false |
| \|PWY-7126\| | ethene biosynthesis IV (engineered) | false | false | false | false | false | false | false | false | false | false | false | false | false | false | false | false | false |
| \|PWY-7127\| | CDP-D-mannitol biosynthesis | false | false | false | false | false | false | false | false | false | false | false | false | false | false | false | false | false |
| \|PWY-7128\| | nicotine degradation III (VPP pathway) | false | false | false | false | false | false | false | false | false | false | false | false | false | false | false | false | false |
| \|PWY-7129\| | quercetin glucoside biosynthesis (Allium) | false | false | false | false | false | false | false | false | false | false | false | false | false | false | false | false | false |
| \|PWY-7130\| | L-glucose degradation | false | false | false | false | false | false | false | false | false | false | false | false | false | false | false | false | false |
| \|PWY-7131\| | CMP-legionaminate biosynthesis II | false | false | false | false | false | false | false | false | false | false | false | false | false | false | false | false | false |
| \|PWY-7133\| | quercetin glucoside degradation (Allium) | false | false | false | false | false | false | false | false | false | false | false | false | false | false | false | false | false |
| \|PWY-7134\| | rutin degradation (plants) | false | false | false | false | false | false | false | false | false | false | false | false | false | false | false | false | false |
| \|PWY-7135\| | emetine biosynthesis | false | false | false | false | false | false | false | false | false | false | false | false | false | false | false | false | false |
| \|PWY-7136\| | β myrcene degradation | false | false | false | false | false | false | false | false | false | false | false | false | false | false | false | false | false |
| \|PWY-7137\| | quercetin gentiotetraside biosynthesis | false | false | false | false | false | false | false | false | false | false | false | false | false | false | false | false | false |
| \|PWY-7138\| | noscapine biosynthesis | false | false | false | false | false | false | false | false | false | false | false | false | false | false | false | false | false |
| \|PWY-7139\| | sesaminol glucoside biosynthesis | false | false | false | false | false | false | false | false | false | false | false | false | false | false | false | false | false |
| \|PWY-7140\| | myricetin gentiobioside biosynthesis | false | false | false | false | false | false | false | false | false | false | false | false | false | false | false | false | false |
| \|PWY-7141\| | (3S)-linalool biosynthesis | false | false | false | false | false | false | false | false | false | false | false | true | false | false | false | false | false |
| \|PWY-7142\| | cyanide detoxification II | false | false | false | false | false | false | false | false | false | false | false | false | false | true | false | false | true |
| \|PWY-7143\| | kaempferol gentiobioside biosynthesis | false | false | false | false | false | false | false | false | false | false | false | false | false | false | false | false | false |
| \|PWY-7145\| | genistin gentiobioside biosynthesis | false | false | false | false | false | false | false | false | false | false | false | false | false | false | false | false | false |
| \|PWY-7147\| | 8-amino-7-oxononanoate biosynthesis II | false | false | false | false | false | false | false | false | false | false | false | false | false | false | false | false | false |
| \|PWY-7150\| | polymethylated quercetin glucoside biosynthesis I - quercetin series (Chrysosplenium) | false | false | false | false | false | false | false | false | false | false | false | false | false | false | false | false | false |
| \|PWY-7151\| | polymethylated quercetin glucoside biosynthesis II - quercetagetin series (Chrysosplenium) | false | false | false | false | false | false | false | false | false | false | false | false | false | false | false | false | false |
| \|PWY-7152\| | pinolenate and coniferonate biosynthesis | false | false | false | false | false | false | false | false | false | false | false | true | false | false | false | false | false |
| \|PWY-7153\| | grixazone biosynthesis | false | false | false | false | false | false | false | false | false | false | false | false | false | false | false | false | false |
| \|PWY-7154\| | ergosterol biosynthesis II | false | false | false | false | false | false | false | false | false | false | false | false | false | false | false | false | false |
| \|PWY-7155\| | 7-dehydroporiferasterol biosynthesis | false | false | false | false | false | false | false | false | false | false | false | false | false | false | true | true | false |
| \|PWY-7157\| | eupatolitin 3-O-glucoside biosynthesis | false | false | false | false | false | false | false | false | false | false | false | false | false | false | false | false | false |
| \|PWY-7158\| | L-phenylalanine degradation V | false | false | false | false | false | false | false | false | false | false | false | false | false | false | false | false | false |
| \|PWY-7159\| | 3,8-divinyl-chlorophyllide a biosynthesis III (aerobic, light independent) | false | false | false | false | false | false | false | false | false | false | false | false | false | false | false | false | false |
| \|PWY-7160\| | polymethylated myricetin biosynthesis (tomato) | false | false | false | false | false | false | false | false | false | false | false | false | false | false | false | false | false |
| \|PWY-7161\| | polymethylated quercetin biosynthesis | false | false | false | false | false | false | false | false | false | false | false | false | false | false | false | false | false |
| \|PWY-7163\| | polymethylated kaempferol biosynthesis | false | false | false | false | false | false | false | false | false | false | false | false | false | false | false | false | false |
| \|PWY-7164\| | chlorophyll a degradation III | false | false | false | false | false | false | false | false | false | false | false | false | false | false | false | false | false |
| \|PWY-7165\| | L-ascorbate biosynthesis VIII (engineered pathway) | false | false | false | false | false | false | false | false | false | false | false | false | false | false | false | false | false |
| \|PWY-7166\| | kaempferide triglycoside biosynthesis | false | false | false | false | false | false | false | false | false | false | false | false | false | false | false | false | false |
| \|PWY-7168\| | flavonol acylglucoside biosynthesis I - kaempferol derivatives | false | false | false | false | false | false | false | false | false | false | false | false | false | false | false | false | false |
| \|PWY-7170\| | phytochromobilin biosynthesis | false | false | false | false | false | false | false | false | false | false | false | false | false | false | false | false | false |
| \|PWY-7171\| | flavonol acylglucoside biosynthesis II - isorhamnetin derivatives | false | false | false | false | false | false | false | false | false | false | false | false | false | false | false | false | false |
| \|PWY-7172\| | flavonol acylglucoside biosynthesis III - quercetin derivatives | false | false | false | false | false | false | false | false | false | false | false | false | false | false | false | false | false |
| \|PWY-7173\| | quercetin triglucoside biosynthesis | false | false | false | false | false | false | false | false | false | false | false | false | false | false | false | false | false |
| \|PWY-7174\| | S-methyl-5-thio-α-D-ribose 1-phosphate degradation II | false | false | false | false | false | false | false | false | false | false | false | false | false | false | false | false | false |
| \|PWY-7175\| | nostoxanthin biosynthesis | false | false | false | false | false | false | false | false | false | false | false | false | false | false | false | false | false |
| \|PWY-7176\| | UTP and CTP de novo biosynthesis | false | false | false | false | false | false | false | false | false | false | false | false | false | false | false | false | false |
| \|PWY-7177\| | UTP and CTP dephosphorylation II | false | false | false | false | false | false | false | false | false | false | false | false | false | false | false | false | false |
| \|PWY-7178\| | D-xylose degradation to ethylene glycol (engineered) | false | false | false | false | false | false | false | false | false | false | false | false | false | false | false | false | false |
| \|PWY-7179-1\| | purine deoxyribonucleosides degradation II | false | false | false | false | false | false | false | false | false | false | false | false | false | false | false | false | false |
| \|PWY-7179\| | purine deoxyribonucleosides degradation I | false | false | false | false | false | false | false | false | false | false | false | false | false | false | false | false | false |
| \|PWY-7181\| | pyrimidine deoxyribonucleosides degradation | false | false | false | false | false | false | false | false | false | false | false | false | false | false | false | false | false |
| \|PWY-7182\| | linalool biosynthesis I | false | false | false | false | false | false | false | false | false | false | false | false | false | false | false | false | false |
| \|PWY-7184\| | pyrimidine deoxyribonucleotides de novo biosynthesis I | false | false | false | false | false | false | false | false | false | false | false | false | false | false | false | false | false |
| \|PWY-7185\| | UTP and CTP dephosphorylation I | false | false | false | false | false | false | false | false | false | false | false | false | false | false | false | false | false |
| \|PWY-7187\| | pyrimidine deoxyribonucleotides de novo biosynthesis II | false | false | false | false | false | false | false | false | false | false | false | false | false | false | false | false | false |
| \|PWY-7188\| | orientin and isoorientin biosynthesis I | false | false | false | false | false | false | false | false | false | false | false | false | false | false | false | false | false |
| \|PWY-7189\| | pinocembrin C-glucosylation | false | false | false | false | false | false | false | false | false | false | false | false | false | false | false | false | false |
| \|PWY-7191\| | kaempferol diglycoside biosynthesis (pollen-specific) | false | false | false | false | false | false | false | false | false | false | false | false | false | false | false | false | false |
| \|PWY-7192\| | quercetin diglycoside biosynthesis (pollen-specific) | false | false | false | false | false | false | false | false | false | false | false | false | false | false | false | false | false |
| \|PWY-7193\| | pyrimidine ribonucleosides salvage I | false | false | false | false | false | false | false | false | false | false | false | false | false | false | false | false | false |
| \|PWY-7195\| | pyrimidine ribonucleosides salvage III | false | false | false | false | false | false | false | false | false | false | false | false | false | false | false | false | false |
| \|PWY-7197\| | pyrimidine deoxyribonucleotide phosphorylation | false | false | false | false | false | false | false | false | false | false | false | false | false | false | false | false | false |
| \|PWY-7198\| | pyrimidine deoxyribonucleotides de novo biosynthesis IV | false | false | false | false | false | false | false | false | false | false | false | false | false | false | false | false | false |
| \|PWY-7199\| | pyrimidine deoxyribonucleosides salvage | false | false | false | false | false | false | false | false | false | false | false | false | false | false | false | false | false |
| \|PWY-7204\| | pyridoxal 5'-phosphate salvage II (plants) | false | false | false | false | false | false | false | false | false | false | false | false | false | false | false | false | false |
| \|PWY-7205\| | CMP phosphorylation | false | false | false | false | false | false | false | false | false | false | false | false | false | false | false | false | false |
| \|PWY-7206\| | pyrimidine deoxyribonucleotides dephosphorylation | false | false | false | false | false | false | false | false | false | false | false | false | false | true | true | true | false |
| \|PWY-7210\| | pyrimidine deoxyribonucleotides biosynthesis from CTP | false | false | false | false | false | false | false | false | false | false | false | false | false | false | false | false | false |
| \|PWY-7212\| | baicalein metabolism | false | false | false | false | false | false | false | false | false | false | false | false | false | false | false | false | false |
| \|PWY-721\| | 3-methylquinoline degradation | false | false | false | false | false | false | false | false | false | false | false | false | false | false | false | false | false |
| \|PWY-7213\| | wogonin metabolism | false | false | false | false | false | false | false | false | false | false | false | false | false | false | false | false | false |
| \|PWY-7214\| | baicalein degradation (hydrogen peroxide detoxification) | false | false | false | false | false | false | false | false | false | false | false | false | false | false | false | false | false |
| \|PWY-7216\| | (R)- and (S)-3-hydroxybutanoate biosynthesis (engineered) | false | false | false | false | false | false | false | false | false | false | false | false | false | false | false | false | false |
| \|PWY-7221\| | guanosine ribonucleotides de novo biosynthesis | false | false | false | false | false | false | false | false | false | false | false | false | false | false | false | false | false |
| \|PWY-7224\| | purine deoxyribonucleosides salvage | false | false | false | false | false | false | false | false | false | false | false | false | false | false | false | false | false |
| \|PWY-7226\| | guanosine deoxyribonucleotides de novo biosynthesis I | false | false | false | false | false | false | false | false | false | false | false | false | false | false | false | false | false |
| \|PWY-7227\| | adenosine deoxyribonucleotides de novo biosynthesis I | false | false | false | false | false | false | false | false | false | false | false | false | false | false | false | false | false |
| \|PWY-722\| | nicotinate degradation I | false | false | false | false | false | false | false | false | false | false | false | false | false | false | false | false | false |
| \|PWY-7230\| | ubiquinol-6 biosynthesis from 4-aminobenzoate (yeast) | false | false | false | false | false | false | false | false | false | false | false | false | false | false | false | false | false |
| \|PWY-7232\| | gibberellin biosynthesis V | false | false | false | false | false | false | false | false | false | false | false | false | false | false | false | false | false |
| \|PWY-7234\| | inosine-5'-phosphate biosynthesis III | false | false | false | false | false | false | false | false | false | false | false | false | false | false | false | false | false |
| \|PWY-7236\| | mycocyclosin biosynthesis | false | false | false | false | false | false | false | false | false | false | false | false | false | false | false | false | false |
| \|PWY-7238\| | sucrose biosynthesis II | false | false | false | false | false | false | false | false | false | false | false | false | false | false | false | false | false |
| \|PWY-723\| | alkylnitronates degradation | false | false | false | false | false | false | false | false | false | false | false | false | false | false | false | false | false |
| \|PWY-7242\| | D-fructuronate degradation | false | false | false | false | false | false | false | false | false | false | false | false | false | false | false | false | false |
| \|PWY-7246\| | pectin degradation I | false | false | false | false | false | false | false | false | false | false | false | false | false | false | false | false | false |
| \|PWY-7247\| | β-D-glucuronide and D-glucuronate degradation | false | false | false | false | false | false | false | false | false | false | false | false | false | false | false | false | false |
| \|PWY-7248\| | pectin degradation II | false | false | false | false | false | false | false | false | false | false | false | false | false | false | false | false | false |
| \|PWY-7250\| | [2Fe-2S] iron-sulfur cluster biosynthesis | false | false | false | false | false | false | false | false | false | false | false | false | false | false | false | false | false |
| \|PWY-7251\| | pentacyclic triterpene biosynthesis | false | false | false | false | false | false | false | false | false | false | false | false | false | false | false | false | false |
| \|PWY-7252\| | luteolinidin 5-O-glucoside biosynthesis | false | false | false | false | false | false | false | false | false | false | false | false | false | false | false | false | false |
| \|PWY-7253\| | apigeninidin 5-O-glucoside biosynthesis | false | false | false | false | false | false | false | false | false | false | false | false | false | false | false | false | false |
| \|PWY-7254\| | TCA cycle VII (acetate-producers) | false | false | false | false | false | false | false | false | false | false | false | false | false | false | false | false | false |
| \|PWY-7255\| | ergothioneine biosynthesis I (bacteria) | false | false | false | false | false | false | false | false | false | false | false | false | false | false | false | false | false |
| \|PWY-7256\| | cyanidin diglucoside biosynthesis (acyl-glucose dependent) | false | false | false | false | false | false | false | false | false | false | false | false | false | false | false | false | false |
| \|PWY-7259\| | pelargonidin diglucoside biosynthesis (acyl-glucose dependent) | false | false | false | false | false | false | false | false | false | false | false | false | false | false | false | false | false |
| \|PWY-7260\| | delphinidin diglucoside biosynthesis (acyl-glucose dependent) | false | false | false | false | false | false | false | false | false | false | false | false | false | false | false | false | false |
| \|PWY-7261\| | anthocyanidin 3-malylglucoside biosynthesis (acyl-glucose dependent) | false | false | false | false | false | false | false | false | false | false | false | false | false | false | false | false | false |
| \|PWY-7262\| | rose anthocyanin biosynthesis II (via cyanidin 3-O-β-D-glucoside) | false | false | false | false | false | false | false | false | false | false | false | false | false | false | false | false | false |
| \|PWY-7265\| | lampranthin biosynthesis | false | false | false | false | false | false | false | false | false | false | false | false | false | false | false | false | false |
| \|PWY-7267\| | anthocyanin biosynthesis (pelargonidin 3-O-glucoside) | false | false | false | false | false | false | false | false | false | false | false | false | false | false | false | false | false |
| \|PWY-7274\| | D-cycloserine biosynthesis | false | false | false | false | false | false | false | false | false | false | false | false | false | false | false | false | false |
| \|PWY-7275\| | L-homophenylalanine biosynthesis | false | false | false | false | false | false | false | false | false | false | false | false | false | false | false | false | false |
| \|PWY-7279\| | aerobic respiration II (cytochrome c) (yeast) | false | false | false | false | false | false | false | false | false | false | false | false | false | false | false | false | false |
| \|PWY-7280\| | ternatin C3 biosynthesis | false | false | false | false | false | false | false | false | false | false | false | false | false | false | false | false | false |
| \|PWY-7282\| | 4-amino-2-methyl-5-diphosphomethylpyrimidine biosynthesis II | false | false | false | false | false | false | false | false | false | false | false | false | false | false | false | false | false |
| \|PWY-7285\| | methylwyosine biosynthesis | false | false | false | false | false | false | false | false | false | false | false | false | false | false | false | false | false |
| \|PWY-7286\| | 7-(3-amino-3-carboxypropyl)-wyosine biosynthesis | false | false | false | false | false | false | false | false | false | false | false | false | false | false | false | false | false |
| \|PWY-7287\| | novobiocin biosynthesis | false | false | false | false | false | false | false | false | false | false | false | false | false | false | false | false | false |
| \|PWY-7288\| | fatty acid β-oxidation VII (yeast peroxisome) | false | false | false | false | false | false | false | false | false | false | false | false | false | false | false | false | false |
| \|PWY-7289\| | L-cysteine biosynthesis V (mycobacteria) | false | false | false | false | false | false | false | false | false | false | false | false | false | false | false | false | false |
| \|PWY-7290\| | Escherichia coli serotype O:86 O antigen biosynthesis | false | false | false | false | false | false | false | false | false | false | false | false | false | false | false | false | false |
| \|PWY-7291\| | oleate β-oxidation (isomerase-dependent, yeast) | false | false | false | false | false | false | false | false | false | false | false | false | false | false | false | false | false |
| \|PWY-7292\| | oleate β-oxidation (thioesterase-dependent, yeast) | false | false | false | false | false | false | false | false | false | false | false | false | false | false | false | false | false |
| \|PWY-7294\| | D-xylose degradation IV | false | false | false | false | false | false | false | false | false | false | false | false | false | false | false | false | false |
| \|PWY-7295\| | L-arabinose degradation IV | false | false | false | false | false | false | false | false | false | false | false | false | false | false | false | false | false |
| \|PWY-7297\| | octopamine biosynthesis | false | false | false | false | false | false | false | false | false | false | false | false | false | false | false | false | false |
| \|PWY-7298\| | nevadensin biosynthesis | false | false | false | false | false | false | false | false | false | false | false | false | false | false | false | false | false |
| \|PWY-7299\| | progesterone biosynthesis | false | false | false | false | false | false | false | false | false | false | false | false | false | false | false | false | false |
| \|PWY-7300\| | ecdysone and 20-hydroxyecdysone biosynthesis | false | false | false | false | false | false | false | false | false | false | false | false | false | false | false | false | false |
| \|PWY-7301\| | dTDP-4-O-demethyl-β-L-noviose biosynthesis | false | false | false | false | false | false | false | false | false | false | false | false | false | false | false | false | false |
| \|PWY-7303\| | 4-hydroxy-3-prenylbenzoate biosynthesis | false | false | false | false | false | false | false | false | false | false | false | false | false | false | false | false | false |
| \|PWY-7304\| | 3-amino-4,7-dihydroxy-coumarin biosynthesis | false | false | false | false | false | false | false | false | false | false | false | false | false | false | false | false | false |
| \|PWY-7306\| | estradiol biosynthesis II | false | false | false | false | false | false | false | false | false | false | false | false | false | false | false | false | false |
| \|PWY-7307\| | oleate β-oxidation (reductase-dependent, yeast) | false | false | false | false | false | false | false | false | false | false | false | false | false | false | false | false | false |
| \|PWY-7308\| | acrylonitrile degradation I | false | false | false | false | false | false | false | false | false | false | false | false | false | false | false | false | false |
| \|PWY-7310\| | D-glucosaminate degradation | false | false | false | false | false | false | false | false | false | false | false | false | false | false | false | false | false |
| \|PWY-7312\| | dTDP-β-D-fucofuranose biosynthesis | false | false | false | false | false | false | false | false | false | false | false | false | false | false | false | false | false |
| \|PWY-7315\| | dTDP-N-acetylthomosamine biosynthesis | false | false | false | false | false | false | false | false | false | false | false | false | false | false | false | false | false |
| \|PWY-7316\| | dTDP-N-acetylviosamine biosynthesis | false | false | false | false | false | false | false | false | false | false | false | false | false | false | false | false | false |
| \|PWY-7318\| | dTDP-3-acetamido-3,6-dideoxy-α-D-glucose biosynthesis | false | false | false | false | false | false | false | false | false | false | false | false | false | false | false | false | false |
| \|PWY-7321\| | ecdysteroid metabolism (arthropods) | false | false | false | false | false | false | false | false | false | false | false | false | false | false | false | false | false |
| \|PWY-7325\| | salvigenin biosynthesis | false | false | false | false | false | false | false | false | false | false | false | false | false | false | false | false | false |
| \|PWY-7333\| | UDP-N-acetyl-α-D-fucosamine biosynthesis | false | false | false | false | false | false | false | false | false | false | false | false | false | false | false | false | false |
| \|PWY-7334\| | UDP-N-acetyl-α-D-quinovosamine biosynthesis | false | false | false | false | false | false | false | false | false | false | false | false | false | false | false | false | false |
| \|PWY-7336\| | UDP-N-acetyl-α-D-galactosaminuronate biosynthesis | false | false | false | false | false | false | false | false | false | false | false | false | false | false | false | false | false |
| \|PWY-7337\| | 10-cis-heptadecenoyl-CoA degradation (yeast) | false | false | false | false | false | false | false | false | false | false | false | false | false | false | false | false | false |
| \|PWY-7338\| | 10-trans-heptadecenoyl-CoA degradation (reductase-dependent, yeast) | false | false | false | false | false | false | false | false | false | false | false | false | false | false | false | false | false |
| \|PWY-7339\| | 10-trans-heptadecenoyl-CoA degradation (MFE-dependent, yeast) | false | false | false | false | false | false | false | false | false | false | false | false | false | false | false | false | false |
| \|PWY-7340\| | 9-cis, 11-trans-octadecadienoyl-CoA degradation (isomerase-dependent, yeast) | false | false | false | false | false | false | false | false | false | false | false | false | false | false | false | false | false |
| \|PWY-7347\| | sucrose biosynthesis III | false | false | false | false | false | false | false | false | false | false | false | false | false | false | false | false | false |
| \|PWY-7351\| | pyruvate-derived opine biosynthesis II (marine invertebrates) | false | false | false | false | false | false | false | false | false | false | false | false | false | false | false | false | false |
| \|PWY-7352\| | daunorubicin biosynthesis | false | false | false | false | false | false | false | false | false | false | false | false | false | false | false | false | false |
| \|PWY-7354\| | aclacinomycin biosynthesis | false | false | false | false | false | false | false | false | false | false | false | false | false | false | false | false | false |
| \|PWY-7355\| | doxorubicin biosynthesis | false | false | false | false | false | false | false | false | false | false | false | false | false | false | false | false | false |
| \|PWY-7356\| | thiamine diphosphate salvage IV (yeast) | false | false | false | false | false | false | false | false | false | false | false | false | false | false | false | false | false |
| \|PWY-735\| | jasmonic acid biosynthesis | false | false | false | false | false | false | false | false | false | false | false | false | false | false | false | false | false |
| \|PWY-7363\| | papaverine biosynthesis | false | false | false | false | false | false | false | false | false | false | false | false | false | false | false | false | false |
| \|PWY-7366\| | phosphatidylglycerol degradation | false | false | false | false | false | false | false | false | false | false | false | false | false | false | false | false | false |
| \|PWY-7367\| | phosphatidylcholine resynthesis via glycerophosphocholine | false | false | false | false | false | false | false | false | false | false | false | false | false | false | false | false | false |
| \|PWY-7369\| | thiamine triphosphate metabolism | false | false | false | false | false | false | false | false | false | false | false | false | false | false | false | false | false |
| \|PWY-7371\| | 1,4-dihydroxy-6-naphthoate biosynthesis II | false | false | false | false | false | false | false | false | false | false | false | false | false | false | false | false | false |
| \|PWY-7372\| | demethylmenaquinol-6 biosynthesis II | false | false | false | false | false | false | false | false | false | false | false | false | false | false | false | false | false |
| \|PWY-7374\| | 1,4-dihydroxy-6-naphthoate biosynthesis I | false | false | false | false | false | false | false | false | false | false | false | false | false | false | false | false | false |
| \|PWY-7375\| | mRNA capping I | false | false | false | false | false | false | false | false | false | false | false | false | false | false | false | false | false |
| \|PWY-7376\| | cob(II)yrinate a,c-diamide biosynthesis II (late cobalt incorporation) | false | false | false | false | false | false | false | false | false | false | false | false | false | false | false | false | false |
| \|PWY-7377\| | cob(II)yrinate a,c-diamide biosynthesis I (early cobalt insertion) | false | false | false | false | false | false | false | false | false | false | false | false | false | false | false | false | false |
| \|PWY-7378\| | aminopropanol phosphate biosynthesis II | false | false | false | false | false | false | false | false | false | false | false | false | false | false | false | false | false |
| \|PWY-7380\| | biotin biosynthesis from 8-amino-7-oxononanoate II | false | false | false | false | false | false | false | false | false | false | false | false | false | false | false | false | false |
| \|PWY-7382\| | lipoate biosynthesis and incorporation IV (yeast) | false | false | false | false | false | false | false | false | false | false | false | false | false | false | false | false | false |
| \|PWY-7383\| | anaerobic energy metabolism (invertebrates, cytosol) | false | false | false | false | false | false | false | false | false | false | false | false | false | false | false | false | false |
| \|PWY-7385\| | 1,3-propanediol biosynthesis (engineered) | false | false | false | false | false | false | false | false | false | false | false | false | false | false | false | false | false |
| \|PWY-7388\| | octanoyl-[acyl-carrier protein] biosynthesis (mitochondria, yeast) | false | false | false | false | false | false | false | false | false | false | false | false | false | false | false | false | false |
| \|PWY-7391\| | isoprene biosynthesis II (engineered) | false | false | false | false | false | false | false | false | false | false | false | false | false | false | false | false | false |
| \|PWY-7394\| | urate conversion to allantoin II | false | false | false | false | false | false | false | false | false | false | false | false | false | false | false | false | false |
| \|PWY-7396\| | butanol and isobutanol biosynthesis (engineered) | false | false | false | false | false | false | false | false | false | false | false | true | false | false | false | false | false |
| \|PWY-7397\| | naringenin biosynthesis (engineered) | false | false | false | false | false | false | false | false | false | false | false | false | false | false | false | false | false |
| \|PWY-7398\| | coumarins biosynthesis (engineered) | false | false | false | false | false | false | false | false | false | false | false | false | false | false | false | false | false |
| \|PWY-7399\| | methylphosphonate degradation II | false | false | false | false | false | false | false | false | false | false | false | false | false | false | false | false | false |
| \|PWY-7400\| | L-arginine biosynthesis IV (archaea) | false | false | false | false | false | false | false | false | false | false | false | false | false | false | false | false | false |
| \|PWY-7403\| | tetramethylpyrazine degradation | false | false | false | false | false | false | false | false | false | false | false | false | false | false | false | false | false |
| \|PWY-7404\| | ceramide phosphoethanolamine biosynthesis | false | false | false | false | false | false | false | false | false | false | false | false | false | false | false | false | false |
| \|PWY-7405\| | aurachin RE biosynthesis | false | false | false | false | false | false | false | false | false | false | false | false | false | false | false | false | false |
| \|PWY-7407\| | aurachin A, B, C and D biosynthesis | false | false | false | false | false | false | false | false | false | false | false | false | false | false | false | false | false |
| \|PWY-7409\| | phospholipid remodeling (phosphatidylethanolamine, yeast) | false | false | false | false | false | false | false | false | false | false | false | false | false | false | false | false | false |
| \|PWY-7410\| | ipsdienol biosynthesis | false | false | false | false | false | false | false | false | false | false | false | false | false | false | false | false | false |
| \|PWY-7411\| | phosphatidate biosynthesis (yeast) | false | false | false | false | false | false | false | false | false | false | false | false | false | false | false | false | false |
| \|PWY-7412\| | mycinamicin biosynthesis | false | false | false | false | false | false | false | false | false | false | false | false | false | false | false | false | false |
| \|PWY-7413\| | dTDP-6-deoxy-α-D-allose biosynthesis | false | false | false | false | false | false | false | false | false | false | false | false | false | false | false | false | false |
| \|PWY-7414\| | dTDP-α-D-mycaminose biosynthesis | false | false | false | false | false | false | false | false | false | false | false | false | false | false | false | false | false |
| \|PWY-7415\| | tylosin biosynthesis | false | false | false | false | false | false | false | false | false | false | false | false | false | false | false | false | false |
| \|PWY-7416\| | phospholipid remodeling (phosphatidylcholine, yeast) | false | false | false | false | false | false | false | false | false | false | false | false | false | false | false | false | false |
| \|PWY-7417\| | phospholipid remodeling (phosphatidate, yeast) | false | false | false | false | false | false | false | false | false | false | false | false | false | false | false | false | false |
| \|PWY-7419\| | FR-900098 and FR-33289 antibiotics biosynthesis | false | false | false | false | false | false | false | false | false | false | false | false | false | false | false | false | false |
| \|PWY-741\| | p-cymene degradation to p-cumate | false | false | false | false | false | false | false | false | false | false | false | false | false | false | false | false | false |
| \|PWY-7420\| | monoacylglycerol metabolism (yeast) | false | false | false | false | false | false | false | false | false | false | false | false | false | false | false | false | false |
| \|PWY-7421\| | narbomycin, pikromycin and novapikromycin biosynthesis | false | false | false | false | false | false | false | false | false | false | false | false | false | false | false | false | false |
| \|PWY-7422\| | methymycin, neomethymycin and novamethymycin biosynthesis | false | false | false | false | false | false | false | false | false | false | false | false | false | false | false | false | false |
| \|PWY-7423\| | bombykol biosynthesis | false | false | false | false | false | false | false | false | false | false | false | false | false | false | false | false | false |
| \|PWY-7424\| | sterol:steryl ester interconversion (yeast) | false | false | false | false | false | false | false | false | false | false | false | false | false | false | false | false | false |
| \|PWY-7425\| | 2-chloroacrylate degradation I | false | false | false | false | false | false | false | false | false | false | false | false | false | false | false | false | false |
| \|PWY-7426\| | complex N-linked glycan biosynthesis (vertebrates) | false | false | false | false | false | false | false | false | false | false | false | false | false | false | false | false | false |
| \|PWY-7428\| | 2-chloroacrylate degradation II | false | false | false | false | false | false | false | false | false | false | false | false | false | false | false | false | false |
| \|PWY-7429\| | arsenite to oxygen electron transfer (via azurin) | false | false | false | false | false | false | false | false | false | false | false | false | false | false | false | false | false |
| \|PWY-7430\| | indole degradation to anthranil and anthranilate | false | false | false | false | false | false | false | false | false | false | false | false | false | false | false | false | false |
| \|PWY-7431\| | aromatic biogenic amine degradation (bacteria) | false | false | false | false | false | false | false | false | false | false | false | false | false | false | false | false | false |
| \|PWY-7433\| | mucin core 1 and core 2 O-glycosylation | false | false | false | false | false | false | false | false | false | false | false | false | false | false | false | false | false |
| \|PWY-7435\| | mucin core 3 and core 4 O-glycosylation | false | false | false | false | false | false | false | false | false | false | false | false | false | false | false | false | false |
| \|PWY-7436\| | vitamin E biosynthesis (tocotrienols) | false | false | false | false | false | false | false | false | false | false | false | false | false | false | false | false | false |
| \|PWY-7437\| | protein O-[N-acetyl]-glucosylation | false | false | false | false | false | false | false | false | false | false | false | false | false | false | false | false | false |
| \|PWY-7439\| | dTDP-β-L-evernitrose biosynthesis | false | false | false | false | false | false | false | false | false | false | false | false | false | false | false | false | false |
| \|PWY-743\| | thiocyanate degradation II | false | false | false | false | false | false | false | false | false | false | false | false | false | false | false | false | false |
| \|PWY-7440\| | dTDP-β-L-4-epi-vancosamine biosynthesis | false | false | false | false | false | false | false | false | false | false | false | false | false | false | false | false | false |
| \|PWY-7441\| | polymyxin A biosynthesis | false | false | false | false | false | false | false | false | false | false | false | false | false | false | false | false | false |
| \|PWY-7442\| | drosopterin and aurodrosopterin biosynthesis | false | false | false | false | false | false | false | false | false | false | false | false | false | false | false | false | false |
| \|PWY-7443\| | (4S)-carvone biosynthesis | false | false | false | false | false | false | false | false | false | false | false | false | false | false | false | false | false |
| \|PWY-7444\| | luteolin triglucuronide biosynthesis | false | false | false | false | false | false | false | false | false | false | false | false | false | false | false | false | false |
| \|PWY-7445\| | luteolin triglucuronide degradation | false | false | false | false | false | false | false | false | false | false | false | false | false | false | false | false | false |
| \|PWY-7446\| | sulfoquinovose degradation I | false | false | false | false | false | false | false | false | false | false | false | false | false | false | false | false | false |
| \|PWY-7447\| | 2-aminoethylphosphonate degradation III | false | false | false | false | false | false | false | false | false | false | false | false | false | false | false | false | false |
| \|PWY-7448\| | galloylated catechin biosynthesis | false | false | false | false | false | false | false | false | false | false | false | false | false | false | false | false | false |
| \|PWY-7449\| | acylated cyanidin galactoside biosynthesis | false | false | false | false | false | false | false | false | false | false | false | false | false | false | false | false | false |
| \|PWY-7450\| | anthocyanidin modification (Arabidopsis) | false | false | false | false | false | false | false | false | false | false | false | false | false | false | false | false | false |
| \|PWY-7452\| | cyanidin dimalonylglucoside biosynthesis | false | false | false | false | false | false | false | false | false | false | false | false | false | false | false | false | false |
| \|PWY-7455\| | allopregnanolone biosynthesis | false | false | false | false | false | false | false | false | false | false | false | false | false | false | false | false | false |
| \|PWY-7456\| | β-(1,4)-mannan degradation | false | false | false | false | false | false | false | false | false | false | false | false | false | false | false | false | false |
| \|PWY-7457\| | sulfite oxidation V (SoeABC) | false | false | false | false | false | false | false | false | false | false | false | false | false | false | false | false | false |
| \|PWY-7458\| | violdelphin biosynthesis | false | false | false | false | false | false | false | false | false | false | false | false | false | false | false | false | false |
| \|PWY-7459\| | kojibiose degradation | false | false | false | false | false | false | false | false | false | false | false | false | false | false | false | false | false |
| \|PWY-7460\| | 2-O-acetyl-3-O-trans-coutarate biosynthesis | false | false | false | false | false | false | false | false | false | false | false | false | false | false | false | false | false |
| \|PWY-7461\| | hydroxycinnamate sugar acid ester biosynthesis | false | false | false | false | false | false | false | false | false | false | false | false | false | false | false | false | false |
| \|PWY-7462\| | 3,3'-disulfanediyldipropannoate degradation | false | false | false | false | false | false | false | false | false | false | false | false | false | false | false | false | false |
| \|PWY-7463\| | N-methylanthraniloyl-β-D-glucopyranose biosynthesis | false | false | false | false | false | false | false | false | false | false | false | false | false | false | false | false | false |

| | | | | | | | | | | | | | | | | | | |
|---|---|---|---|---|---|---|---|---|---|---|---|---|---|---|---|---|---|---|
| \|PWY-7464\| | cyanidin 3,7-diglucoside polyacylation biosynthesis | false | false | false | false | false | false | false | false | false | false | false | false | false | false | false | false | false |
| \|PWY-7465\| | 3,3'-thiodipropanoate degradation | false | false | false | false | false | false | false | false | false | false | false | false | false | false | false | false | false |
| \|PWY-7466\| | acetone degradation III (to propane-1,2-diol) | false | false | false | false | false | false | false | false | false | false | false | false | false | false | false | false | false |
| \|PWY-7468\| | benzoyl-β-D-glucopyranose biosynthesis | false | false | false | false | false | false | false | false | false | false | false | false | false | false | false | false | false |
| \|PWY-7469\| | gentisate degradation II | false | false | false | false | false | false | false | false | false | false | false | false | false | false | false | false | false |
| \|PWY-7470\| | phosphatidylcholine biosynthesis VII | false | false | false | false | false | false | false | false | false | false | false | false | false | false | false | false | false |
| \|PWY-7471\| | D-carnitine degradation I | false | false | false | false | false | false | false | false | false | false | false | false | false | false | false | false | false |
| \|PWY-7472\| | D-carnitine degradation II | false | false | false | false | false | false | false | false | false | false | false | false | false | false | false | false | false |
| \|PWY-7473\| | avenacin A-1 biosynthesis | false | false | false | false | false | false | false | false | false | false | false | false | false | false | false | false | false |
| \|PWY-7474\| | avenacin A-2 biosynthesis | false | false | false | false | false | false | false | false | false | false | false | false | false | false | false | false | false |
| \|PWY-7475\| | des-methyl avenacin A-1 biosynthesis | false | false | false | false | false | false | false | false | false | false | false | false | false | false | false | false | false |
| \|PWY-7477\| | momilactone A biosynthesis | false | false | false | false | false | false | false | false | false | false | false | false | false | false | false | false | false |
| \|PWY-7478\| | oryzalexin D and E biosynthesis | false | false | false | false | false | false | false | false | false | false | false | false | false | false | false | false | false |
| \|PWY-7480\| | 2,3-dihydroxybenzoate degradation | false | false | false | false | false | false | false | false | false | false | false | false | false | false | false | false | false |
| \|PWY-7481\| | oryzalide A biosynthesis | false | false | false | false | false | false | false | false | false | false | false | false | false | false | false | false | false |
| \|PWY-7482\| | cyclooctatin biosynthesis | false | false | false | false | false | false | false | false | false | false | false | false | false | true | false | true | false |
| \|PWY-7483\| | elloramycin biosynthesis | false | false | false | false | false | false | false | false | false | false | false | false | false | false | false | false | false |
| \|PWY-7484\| | phytocassanes biosynthesis, shared reactions | false | false | false | false | false | false | false | false | false | false | false | false | false | false | false | false | false |
| \|PWY-7485\| | tetracenomycin C biosynthesis | false | false | false | false | false | false | false | false | false | false | false | false | false | false | false | false | false |
| \|PWY-7487\| | (+)-secoisolariciresinol diglucoside biosynthesis | false | false | false | false | false | false | false | false | false | false | false | false | false | false | false | false | false |
| \|PWY-7489\| | oryzalexin A, B, and C biosynthesis | false | false | false | false | false | false | false | false | false | false | false | false | false | false | false | false | false |
| \|PWY-7490\| | patulin biosynthesis | false | false | false | false | false | false | false | false | false | false | false | false | false | false | false | false | false |
| \|PWY-7491\| | podophyllotoxin glucosides metabolism | false | false | false | false | false | false | false | false | false | false | false | false | false | false | false | false | false |
| \|PWY-7492\| | paspaline biosynthesis | false | false | false | false | false | false | false | false | false | false | false | false | false | false | false | false | false |
| \|PWY-7493\| | paxilline and diprenylpaxilline biosynthesis | false | false | false | false | false | false | false | false | false | false | false | false | false | false | false | false | false |
| \|PWY-7495\| | gossypetin metabolism | false | false | false | false | false | false | false | false | false | false | false | false | false | false | false | false | false |
| \|PWY-7496\| | linuron degradation | false | false | false | false | false | false | false | false | false | false | false | false | false | false | false | false | false |
| \|PWY-7497\| | 3β-hydroxysesquiterpene lactone biosynthesis | false | false | false | false | false | false | false | false | false | false | false | false | false | false | false | false | false |
| \|PWY-7498\| | phenylpropanoids methylation (ice plant) | false | false | false | false | false | false | false | false | false | false | false | false | false | false | false | false | false |
| \|PWY-7501\| | phosphatidylserine biosynthesis I | false | false | false | false | false | false | false | false | false | false | false | false | false | false | false | false | false |
| \|PWY-7506\| | phosphatidylserine biosynthesis II | false | false | false | false | false | false | false | false | false | false | false | false | false | false | false | false | false |
| \|PWY-7507\| | chelerythrine biosynthesis | false | false | false | false | false | false | false | false | false | false | false | false | false | false | false | false | false |
| \|PWY-7510\| | rhizocticin A and B biosynthesis | false | false | false | false | false | false | false | false | false | false | false | false | false | false | false | false | false |
| \|PWY-7511\| | protein ubiquitination | false | false | false | false | false | false | false | false | false | false | false | false | false | false | false | false | false |
| \|PWY-7512\| | 3,5,6-trichloro-2-pyridinol degradation | false | false | false | false | false | false | false | false | false | false | false | false | false | false | false | false | false |
| \|PWY-7513\| | flaviolin dimer and mompain biosynthesis | false | false | false | false | false | false | false | false | false | false | false | false | false | false | false | false | false |
| \|PWY-7514\| | L-tyrosine degradation IV (to 4-methylphenol) | false | false | false | false | false | false | false | false | false | false | false | false | false | false | false | false | false |
| \|PWY-7515\| | trans-3-hydroxy-L-proline degradation | false | false | false | false | false | false | false | false | false | false | false | false | false | false | false | false | false |
| \|PWY-7516\| | L-lyxonate degradation | false | false | false | false | false | false | false | false | false | false | false | false | false | false | false | false | false |
| \|PWY-7517\| | brassicicene C biosynthesis | false | false | false | false | false | false | false | false | false | false | false | false | false | false | false | false | false |
| \|PWY-7518\| | atromentin biosynthesis | false | false | false | false | false | false | false | false | false | false | false | false | false | false | false | false | false |
| \|PWY-7520\| | terrequinone A biosynthesis | false | false | false | false | false | false | false | false | false | false | false | false | false | false | false | false | false |
| \|PWY-7521\| | 3-(4-sulfophenyl)butanoate degradation | false | false | false | false | false | false | false | false | false | false | false | false | false | false | false | false | false |
| \|PWY-7522\| | (R)-canadine biosynthesis | false | false | false | false | false | false | false | false | false | false | false | false | false | false | false | false | false |
| \|PWY-7523\| | L-arginine degradation XII | false | false | false | false | false | false | false | false | false | false | false | false | false | false | false | false | false |
| \|PWY-7524\| | mevalonate pathway III (Thermoplasma) | false | false | false | false | false | false | false | false | false | false | false | false | false | false | false | false | false |
| \|PWY-7525\| | fumitremorgin C biosynthesis | false | false | false | false | false | false | false | false | false | false | false | false | false | false | false | false | false |
| \|PWY-7526\| | fumitremorgin A biosynthesis | false | false | false | false | false | false | false | false | false | false | false | false | false | false | false | false | false |
| \|PWY-7529\| | CMP-N-acetyl-7-O-acetylneuraminate biosynthesis | false | false | false | false | false | false | false | false | false | false | false | false | false | false | false | false | false |
| \|PWY-7530\| | Escherichia coli serotype O:104 O antigen biosynthesis | false | false | false | false | false | false | false | false | false | false | false | false | false | false | false | false | false |
| \|PWY-7531\| | mannojirimycin biosynthesis | false | false | false | false | false | false | false | false | false | false | false | false | false | false | false | false | false |
| \|PWY-7532\| | acetylaszonalenin biosynthesis | false | false | false | false | false | false | false | false | false | false | false | false | false | false | false | false | false |
| \|PWY-7533\| | gliotoxin biosynthesis | false | false | false | false | false | false | false | false | false | false | false | false | false | false | false | false | false |
| \|PWY-7534\| | gliotoxin inactivation | false | false | false | false | false | false | false | false | false | false | false | false | false | false | false | false | false |
| \|PWY-7535\| | lovastatin biosynthesis | false | false | false | false | false | false | false | false | false | false | false | false | false | false | false | false | false |
| \|PWY-7536\| | 2-amino-3-hydroxycyclopent-2-enone biosynthesis | false | false | false | false | false | false | false | false | false | false | false | false | false | false | false | false | false |
| \|PWY-7539\| | 6-hydroxymethyl-dihydropterin diphosphate biosynthesis III (Chlamydia) | false | false | false | false | false | false | false | false | false | false | false | false | false | false | false | false | false |
| \|PWY-7540\| | aflatrem biosynthesis | false | false | false | false | false | false | false | false | false | false | false | false | false | false | false | false | false |
| \|PWY-7541\| | 1,2-propanediol biosynthesis from lactate (engineered) | false | false | false | false | false | false | false | false | false | false | false | false | false | false | false | false | false |
| \|PWY-7542\| | fumiquinazoline D biosynthesis | false | false | false | false | false | false | false | false | false | false | false | false | false | false | false | false | false |
| \|PWY-7543\| | 5-N-acetylardeemin biosynthesis | false | false | false | false | false | false | false | false | false | false | false | false | false | false | false | false | false |
| \|PWY-7544\| | pyruvate to cytochrome bo oxidase electron transfer | false | false | false | false | false | false | false | false | false | false | false | false | false | false | false | false | false |
| \|PWY-7545\| | pyruvate to cytochrome bd oxidase electron transfer | false | false | false | false | false | false | false | false | false | false | false | false | false | false | false | false | false |
| \|PWY-7546\| | diphthamide biosynthesis II (eukaryotes) | false | false | false | false | false | false | false | false | false | false | false | false | false | false | false | false | false |
| \|PWY-7547\| | prodigiosin biosynthesis | false | false | false | false | false | false | false | false | false | false | false | false | false | false | false | false | false |
| \|PWY-7549\| | asperlicin E biosynthesis | false | false | false | false | false | false | false | false | false | false | false | false | false | false | false | false | false |
| \|PWY-7550\| | ergothioneine biosynthesis II (fungi) | false | false | false | false | false | false | false | false | false | false | false | false | false | false | false | false | false |
| \|PWY-7552\| | heme b biosynthesis III (from siroheme) | false | false | false | false | false | false | false | false | false | false | false | false | false | false | false | false | false |
| \|PWY-7554\| | heme d1 biosynthesis | false | false | false | false | false | false | false | false | false | false | false | false | false | false | false | false | false |
| \|PWY-7555\| | α-cyclopiazonate biosynthesis | false | false | false | false | false | false | false | false | false | false | false | false | false | false | false | false | false |
| \|PWY-7556\| | (-)-microperfuranone biosynthesis | false | false | false | false | false | false | false | false | false | false | false | false | false | false | false | false | false |
| \|PWY-7557\| | (-)-dehydrodiconiferyl alcohol degradation | false | false | false | false | false | false | false | false | false | false | false | false | false | false | false | false | false |
| \|PWY-7558\| | α-cyclopiazonate detoxification | false | false | false | false | false | false | false | false | false | false | false | false | false | false | false | false | false |
| \|PWY-7559\| | glutathione degradation (DUG pathway) | false | false | false | false | false | false | false | false | false | false | false | false | false | false | false | false | false |
| \|PWY-7561\| | tenellin biosynthesis | false | false | false | false | false | false | false | false | false | false | false | false | false | false | false | false | false |
| \|PWY-7562\| | 3,6-anhydro-α-L-galactopyranose degradation | false | false | false | false | false | false | false | false | false | false | false | false | false | false | false | false | false |
| \|PWY-7563\| | bassianin and desmethylbassianin biosynthesis | false | false | false | false | false | false | false | false | false | false | false | false | false | false | false | false | false |
| \|PWY-7564\| | bacimethrin and bacimethrin pyrophosphate biosynthesis | false | false | false | false | false | false | false | false | false | false | false | false | false | false | false | false | false |
| \|PWY-7565\| | aspyridone A biosynthesis | false | false | false | false | false | false | false | false | false | false | false | false | false | false | false | false | false |
| \|PWY-7569\| | arginomycin biosynthesis | false | false | false | false | false | false | false | false | false | false | false | false | false | false | false | false | false |
| \|PWY-7570\| | blasticidin S biosynthesis | false | false | false | false | false | false | false | false | false | false | false | false | false | false | false | false | false |
| \|PWY-7571\| | ferrichrome A biosynthesis | false | false | false | false | false | false | false | false | false | false | false | false | false | false | false | false | false |
| \|PWY-7572\| | lolitrem B biosynthesis | false | false | false | false | false | false | false | false | false | false | false | false | false | false | false | false | false |
| \|PWY-7573\| | GDP-mycosamine biosynthesis | false | false | false | false | false | false | false | false | false | false | false | false | false | false | false | false | false |
| \|PWY-7574\| | propanoyl-CoA degradation II | false | false | false | false | false | false | false | false | false | false | false | false | false | false | false | false | false |
| \|PWY-7576\| | nitrogen fixation II (flavodoxin) | false | false | false | false | false | false | false | false | false | false | false | false | false | false | false | false | false |
| \|PWY-7577\| | ferrichrome biosynthesis | false | false | false | false | false | false | false | false | false | false | false | false | false | false | false | false | false |
| \|PWY-7578\| | phycoviolobilin biosynthesis | false | false | false | false | false | false | false | false | false | false | false | false | false | false | false | false | false |
| \|PWY-7579\| | phycourobilin biosynthesis | false | false | false | false | false | false | false | false | false | false | false | false | false | false | false | false | false |
| \|PWY-7580\| | phycoerythrobilin biosynthesis II | false | false | false | false | false | false | false | false | false | false | false | false | false | false | false | false | false |
| \|PWY-7582\| | mercaptosuccinate degradation | false | false | false | false | false | false | false | false | false | false | false | false | false | false | false | false | false |
| \|PWY-7583\| | arachidonate biosynthesis II (bacteria) | false | false | false | false | false | false | false | false | false | false | false | false | false | false | false | false | false |
| \|PWY-7585\| | docosahexaenoate biosynthesis II (bacteria) | false | false | false | false | false | false | false | false | false | false | false | false | false | false | false | false | false |
| \|PWY-7586\| | β-1,4-D-mannosyl-N-acetyl-D-glucosamine degradation | false | false | false | false | false | false | false | false | false | false | false | false | false | false | false | false | false |
| \|PWY-7587\| | oleate biosynthesis III (cyanobacteria) | false | false | false | false | false | false | false | false | false | false | false | false | false | false | false | false | false |
| \|PWY-7588\| | ursodeoxycholate biosynthesis (bacteria) | false | false | false | false | false | false | false | false | false | false | false | false | false | false | false | false | false |
| \|PWY-7589\| | palmitoleate biosynthesis III (cyanobacteria) | false | false | false | false | false | false | false | false | false | false | false | false | false | false | false | false | false |
| \|PWY-7590\| | (7Z,10Z,13Z)-hexadecatrienoate biosynthesis | false | false | false | false | false | false | false | false | false | false | false | false | false | false | false | false | false |
| \|PWY-7591\| | okenone biosynthesis | false | false | false | false | false | false | false | false | false | false | false | false | false | false | false | false | false |
| \|PWY-7593\| | linoleate biosynthesis III (cyanobacteria) | false | false | false | false | false | false | false | false | false | false | false | false | false | false | false | false | false |
| \|PWY-7594\| | γ-linolenate biosynthesis III (cyanobacteria) | false | false | false | false | false | false | false | false | false | false | false | false | false | false | false | false | false |
| \|PWY-7595\| | stearidonate biosynthesis (cyanobacteria) | false | false | false | false | false | false | false | false | false | false | false | false | false | false | false | false | false |
| \|PWY-7598\| | α-linolenate biosynthesis II (cyanobacteria) | false | false | false | false | false | false | false | false | false | false | false | false | false | false | false | false | false |
| \|PWY-7599\| | anditomin biosynthesis | false | false | false | false | false | false | false | false | false | false | false | false | false | false | false | false | false |
| \|PWY-7600\| | peramine biosynthesis | false | false | false | false | false | false | false | false | false | false | false | false | false | false | false | false | false |
| \|PWY-7601\| | arachidonate biosynthesis IV (8-detaturase, lower eukaryotes) | false | false | false | false | false | false | false | false | false | false | false | false | false | false | false | false | false |
| \|PWY-7602\| | icosapentaenoate biosynthesis V (8-desaturase, lower eukaryotes) | false | false | false | false | false | false | false | false | false | false | false | false | false | false | false | false | false |
| \|PWY-7603\| | stephacidin A biosynthesis | false | false | false | false | false | false | false | false | false | false | false | false | false | false | false | false | false |
| \|PWY-7604\| | notoamide C and D biosynthesis | false | false | false | false | false | false | false | false | false | false | false | false | false | false | false | false | false |
| \|PWY-7605\| | roquefortine C biosynthesis | false | false | false | false | false | false | false | false | false | false | false | false | false | false | false | false | false |
| \|PWY-7606\| | docosahexaenoate biosynthesis III (6-desaturase, mammals) | false | false | false | false | false | false | false | false | false | false | false | false | false | false | false | false | false |
| \|PWY-7607\| | meleagrin biosynthesis | false | false | false | false | false | false | false | false | false | false | false | false | false | false | false | false | false |
| \|PWY-7608\| | neoxaline biosynthesis | false | false | false | false | false | false | false | false | false | false | false | false | false | false | false | false | false |
| \|PWY-7610\| | GDP-6-deoxy-D-altro-heptose biosynthesis | false | false | false | false | false | false | false | false | false | false | false | false | false | false | false | false | false |
| \|PWY-7612\| | chaetoglobosin A biosynthesis | false | false | false | false | false | false | false | false | false | false | false | false | false | false | false | false | false |
| \|PWY-7613\| | GDP-6-deoxy-D-manno-heptose biosynthesis | false | false | false | false | false | false | false | false | false | false | false | false | false | false | false | false | false |
| \|PWY-7614\| | methiin metabolism | false | false | false | false | false | false | false | false | false | false | false | false | false | false | false | false | false |
| \|PWY-7615\| | pterocarpan phytoalexins modification (maackiain, medicarpin, pisatin, phaseollin) | false | false | false | false | false | false | false | false | false | false | false | false | false | false | false | false | false |
| \|PWY-7618\| | ricinoleate biosynthesis | false | false | false | false | false | false | false | false | false | false | false | false | false | false | false | false | false |
| \|PWY-7619\| | juniperonate biosynthesis | false | false | false | false | false | false | false | false | false | false | false | false | false | false | false | false | false |
| \|PWY-761\| | rhizobactin 1021 biosynthesis | false | false | false | false | false | false | false | false | false | false | false | false | false | false | false | false | false |
| \|PWY-7620\| | naphthalene degradation (anaerobic) | false | false | false | false | false | false | false | false | false | false | false | false | false | false | false | false | false |
| \|PWY-7621\| | autoinducer CAI-1 biosynthesis | false | false | false | false | false | false | false | false | false | false | false | false | false | false | false | false | false |
| \|PWY-7623\| | astaxanthin dirhamnoside biosynthesis | false | false | false | false | false | false | false | false | false | false | false | false | false | false | false | false | false |
| \|PWY-7624\| | nystatin biosynthesis | false | false | false | false | false | false | false | false | false | false | false | false | false | false | false | false | false |
| \|PWY-7625\| | phosphatidylinositol biosynthesis II (eukaryotes) | false | false | false | false | false | false | false | false | false | false | false | false | false | false | false | false | false |
| \|PWY-7626\| | bacilysin biosynthesis | false | false | false | false | false | false | false | false | false | false | false | false | false | false | false | false | false |
| \|PWY-7627\| | 2,4,6-trinitrophenol and 2,4-dinitrophenol degradation | false | false | false | false | false | false | false | false | false | false | false | false | false | false | false | false | false |
| \|PWY-7628\| | 2,4-dinitroanisole degradation | false | false | false | false | false | false | false | false | false | false | false | false | false | false | false | false | false |
| \|PWY-7629\| | yatein biosynthesis II | false | false | false | false | false | false | false | false | false | false | false | false | false | false | false | false | false |
| \|PWY-762\| | phospholipid desaturation | false | false | false | false | false | false | false | false | false | false | false | false | false | false | false | false | false |
| \|PWY-7630\| | hinokinin biosynthesis | false | false | false | false | false | false | false | false | false | false | false | false | false | false | false | false | false |
| \|PWY-7631\| | arctigenin and isoarctigenin biosynthesis | false | false | false | false | false | false | false | false | false | false | false | false | false | false | false | false | false |
| \|PWY-7632\| | hinokiresinol biosynthesis | false | false | false | false | false | false | false | false | false | false | false | false | false | false | false | false | false |
| \|PWY-7633\| | calycosin 7-O-glucoside biosynthesis | false | false | false | false | false | false | false | false | false | false | false | false | false | false | false | false | false |
| \|PWY-7634\| | vernolate biosynthesis II | false | false | false | false | false | false | false | false | false | false | false | false | false | false | false | false | false |
| \|PWY-7635\| | kievitone detoxification | false | false | false | false | false | false | false | false | false | false | false | false | false | false | false | false | false |
| \|PWY-7636\| | astaxanthin biosynthesis (flowering plants) | false | false | false | false | false | false | false | false | false | false | false | false | false | false | false | false | false |
| \|PWY-7637\| | 2,2'-dihydroxyketocarotenoids biosynthesis | false | false | false | false | false | false | false | false | false | false | false | false | false | false | false | false | false |
| \|PWY-7638\| | echinenone and zeaxanthin biosynthesis (Synechocystis) | false | false | false | false | false | false | false | false | false | false | false | false | false | false | false | false | false |
| \|PWY-7639\| | bis(guanylyl molybdenum cofactor) biosynthesis | false | false | false | false | false | false | false | false | false | false | false | false | false | false | false | false | false |
| \|PWY-7640\| | abscisic acid degradation to neophaseic acid | false | false | false | false | false | false | false | false | false | false | false | false | false | false | false | false | false |
| \|PWY-7641\| | 5-hexynoate biosynthesis | false | false | false | false | false | false | false | false | false | false | false | false | false | false | false | false | false |
| \|PWY-7642\| | abscisic acid degradation to 7'-hydroxyabscisate | false | false | false | false | false | false | false | false | false | false | false | false | false | false | false | false | false |
| \|PWY-7643\| | coniferyl alcohol 9-methyl ester biosynthesis | false | false | false | false | false | false | false | false | false | false | false | false | false | false | false | false | false |
| \|PWY-7644\| | heparin degradation | false | false | false | false | false | false | false | false | false | false | false | false | false | false | false | false | false |
| \|PWY-7645\| | hyaluronan degradation | false | false | false | false | false | false | false | false | false | false | false | false | false | false | false | false | false |
| \|PWY-7646\| | dermatan sulfate degradation I (bacterial) | false | false | false | false | false | false | false | false | false | false | false | false | false | false | false | false | false |
| \|PWY-7647\| | ulvan degradation | false | false | false | false | false | false | false | false | false | false | false | false | false | false | false | false | false |
| \|PWY-7648\| | 3-hydroxy-4-methyl-proline biosynthesis | false | false | false | false | false | false | false | false | false | false | false | false | false | false | false | false | false |
| \|PWY-7649\| | 3-hydroxy-L-homotyrosine biosynthesis | false | false | false | false | false | false | false | false | false | false | false | false | false | false | false | false | false |
| \|PWY-7650\| | echinocandin B biosynthesis | false | false | false | false | false | false | false | false | false | false | false | false | false | false | false | false | false |
| \|PWY-7651\| | heparan sulfate degradation | false | false | false | false | false | false | false | false | false | false | false | false | false | false | false | false | false |
| \|PWY-7652\| | echinocandin B degradation | false | false | false | false | false | false | false | false | false | false | false | false | false | false | false | false | false |
| \|PWY-7653\| | griseofulvin biosynthesis | false | false | false | false | false | false | false | false | false | false | false | false | false | false | false | false | false |
| \|PWY-7654\| | (8E,10E)-dodeca-8,10-dienol biosynthesis | false | false | false | false | false | false | false | false | false | false | false | false | false | false | false | false | false |
| \|PWY-7655\| | dechlorogriseofulvin biosynthesis | false | false | false | false | false | false | false | false | false | false | false | false | false | false | false | false | false |
| \|PWY-7656\| | Spodoptera littoralis pheromone biosynthesis | false | false | false | false | false | false | false | false | false | false | false | false | false | false | false | false | false |
| \|PWY-7657\| | dTDP-β-L-digitoxose biosynthesis | false | false | false | false | false | false | false | false | false | false | false | false | false | false | false | false | false |
| \|PWY-7658\| | protein N-glycosylation (Methanococcus voltae) | false | false | false | false | false | false | false | false | false | false | false | false | false | false | false | false | false |
| \|PWY-7659\| | viridicatumtoxin biosynthesis | false | false | false | false | false | false | false | false | false | false | false | false | false | false | false | false | false |
| \|PWY-7660\| | tryptoquialanine biosynthesis | false | false | false | false | false | false | false | false | false | false | false | false | false | false | false | false | false |
| \|PWY-7661\| | protein N-glycosylation (Haloferax volcanii) | false | false | false | false | false | false | false | false | false | false | false | false | false | false | false | false | false |
| \|PWY-7662\| | glycogen degradation III (via anhydrofructose) | false | false | false | false | false | false | false | false | false | false | false | false | false | false | false | false | false |
| \|PWY-7663\| | gondoate biosynthesis (anaerobic) | false | false | false | false | false | false | false | false | false | false | false | false | false | false | false | false | false |
| \|PWY-7664\| | oleate biosynthesis IV (anaerobic) | false | false | false | false | false | false | false | false | false | false | false | false | false | false | false | false | false |
| \|PWY-7665\| | aureobasidin A biosynthesis | false | false | false | false | false | false | false | false | false | false | false | false | false | false | false | false | false |
| \|PWY-7666\| | galactolipid biosynthesis II | false | false | false | false | false | false | false | false | false | false | false | false | false | false | false | false | false |
| \|PWY-7667\| | apicidin biosynthesis | false | false | false | false | false | false | false | false | false | false | false | false | false | false | false | false | false |
| \|PWY-7668\| | apicidin F biosynthesis | false | false | false | false | false | false | false | false | false | false | false | false | false | false | false | false | false |
| \|PWY-7669\| | equisetin biosynthesis | false | false | false | false | false | false | false | false | false | false | false | false | false | false | false | false | false |
| \|PWY-7670\| | fusaridione A biosynthesis | false | false | false | false | false | false | false | false | false | false | false | false | false | false | false | false | false |
| \|PWY-7671\| | saframycin A biosynthesis | false | false | false | false | false | false | false | false | false | false | false | false | false | false | false | false | false |
| \|PWY-7672\| | fusaric acid biosynthesis | false | false | false | false | false | false | false | false | false | false | false | false | false | false | false | false | false |
| \|PWY-7673\| | fusarin C biosynthesis | false | false | false | false | false | false | false | false | false | false | false | false | false | false | false | false | false |
| \|PWY-7674\| | CMP-8-amino-3,8-dideoxy-D-manno-octulosonate biosynthesis | false | false | false | false | false | false | false | false | false | false | false | false | false | false | false | false | false |
| \|PWY-7675\| | Kdo transfer to lipid IVA (Haemophilus) | false | false | false | false | false | false | false | false | false | false | false | false | false | false | false | false | false |
| \|PWY-7676\| | Kdo8N transfer to lipid IVA | false | false | false | false | false | false | false | false | false | false | false | false | false | false | false | false | false |
| \|PWY-7677\| | rosamicin biosynthesis | false | false | false | false | false | false | false | false | false | false | false | false | false | false | false | false | false |
| \|PWY-7678\| | anthocyanidin sambubioside biosynthesis | false | false | false | false | false | false | false | false | false | false | false | false | false | false | false | false | false |
| \|PWY-7679\| | anthocyanidin acylglucoside and acylsambubioside biosynthesis | false | false | false | false | false | false | false | false | false | false | false | false | false | false | false | false | false |
| \|PWY-7680\| | carnosate bioynthesis | false | false | false | false | false | false | false | false | false | false | false | false | false | false | false | false | false |
| \|PWY-7681\| | 1-chloro-2-nitrobenzene degradation | false | false | false | false | false | false | false | false | false | false | false | false | false | false | false | false | false |
| \|PWY-7682\| | arabidopyrone biosynthesis | false | false | false | false | false | false | false | false | false | false | false | false | false | false | false | false | false |
| \|PWY-7683\| | nitrite reduction (hemoglobin) | false | false | false | false | false | false | false | false | false | false | false | false | false | false | false | false | false |
| \|PWY-7684\| | α-diglucosyldiacylglycerol biosynthesis | false | false | false | false | false | false | false | false | false | false | false | false | false | false | false | false | false |
| \|PWY-7687\| | stipitatate biosynthesis | false | false | false | false | false | false | false | false | false | false | false | false | false | false | false | false | false |
| \|PWY-7688\| | dTDP-α-D-ravidosamine and dTDP-4-acetyl-α-D-ravidosamine biosynthesis | false | false | false | false | false | false | false | false | false | false | false | false | false | false | false | false | false |
| \|PWY-7689\| | 8-O-methylfusarubin biosynthesis | false | false | false | false | false | false | false | false | false | false | false | false | false | false | false | false | false |
| \|PWY-7690\| | holomycin biosynthesis | false | false | false | false | false | false | false | false | false | false | false | false | false | false | false | false | false |
| \|PWY-7691\| | 10,13-epoxy-11-methyl-octadecadienoate biosynthesis | false | false | false | false | false | false | false | false | false | false | false | false | false | false | false | false | false |
| \|PWY-7692\| | bikaverin biosynthesis | false | false | false | false | false | false | false | false | false | false | false | false | false | false | false | false | false |
| \|PWY-7693\| | guadinomine B biosynthesis | false | false | false | false | false | false | false | false | false | false | false | false | true | false | true | false | true |
| \|PWY-7694\| | zwittermicin A biosynthesis | false | false | false | false | false | false | false | false | false | false | false | false | false | false | false | false | false |
| \|PWY-7695\| | aurofusarin biosynthesis | false | false | false | false | false | false | false | false | false | false | false | false | false | false | false | false | false |
| \|PWY-7696\| | citreoisocoumarin and bikisocoumarin biosynthesis | false | false | false | false | false | false | false | false | false | false | false | false | false | false | false | false | false |
| \|PWY-7697\| | geranyl β-primeveroside biosynthesis | false | false | false | false | false | false | false | false | false | false | false | false | false | false | false | false | false |
| \|PWY-7698\| | 2,5-xylenol and 3,5-xylenol degradation | false | false | false | false | false | false | false | false | false | false | false | false | false | false | false | false | false |
| \|PWY-7700\| | 4-methylphenol degradation to protocatechuate | false | false | false | false | false | false | false | false | false | false | false | false | false | false | false | false | false |
| \|PWY-7701\| | 4-hydroxy-4-methyl-L-glutamate biosynthesis | false | false | false | false | false | false | false | false | false | false | false | false | false | false | false | false | false |
| \|PWY-7702\| | sch210971 and sch210972 biosynthesis | false | false | false | false | false | false | false | false | false | false | false | false | false | false | false | false | false |
| \|PWY-7703\| | 2,4-xylenol degradation to protocatechuate | false | false | false | false | false | false | false | false | false | false | false | false | false | false | false | false | false |
| \|PWY-7704\| | viridicatin biosynthesis | false | false | false | false | false | false | false | false | false | false | false | false | false | false | false | false | false |
| \|PWY-7705\| | quinolinone B biosynthesis | false | false | false | false | false | false | false | false | false | false | false | false | false | false | false | false | false |
| \|PWY-7706\| | dapdiamides biosynthesis | false | false | false | false | false | false | false | false | false | false | false | false | false | false | false | false | false |
| \|PWY-7708\| | lyngbyatoxin biosynthesis | false | false | false | false | false | false | false | false | false | false | false | false | false | false | false | false | false |
| \|PWY-7709\| | (3R)-linalool biosynthesis | false | false | false | false | false | false | false | false | false | false | false | false | false | false | false | false | false |
| \|PWY-7710\| | FeMo cofactor biosynthesis | false | false | false | false | false | false | false | false | false | false | false | false | false | false | false | false | false |
| \|PWY-7711\| | calonectrin biosynthesis | false | false | false | false | false | false | false | false | false | false | false | false | false | false | false | false | false |
| \|PWY-7712\| | deoxynivalenol biosynthesis | false | false | false | false | false | false | false | false | false | false | false | false | false | false | false | false | false |
| \|PWY-7713\| | nivalenol biosynthesis | false | false | false | false | false | false | false | false | false | false | false | false | false | false | false | false | false |
| \|PWY-7714\| | harzianum A and trichodermin biosynthesis | false | false | false | false | false | false | false | false | false | false | false | false | false | false | false | false | false |
| \|PWY-7716\| | penicillin G and penicillin V biosynthesis | false | false | false | false | false | false | false | false | false | false | false | false | false | false | false | false | false |
| \|PWY-7717\| | 3-hydroxy-4-methyl-anthranilate biosynthesis I | false | false | false | false | false | false | false | false | false | false | false | false | false | false | false | false | false |
| \|PWY-7718\| | actinomycin D biosynthesis | false | false | false | false | false | false | false | false | false | false | false | false | false | false | false | false | false |
| \|PWY-7719\| | CMP-diacetamido-8-epilegionaminic acid biosynthesis | false | false | false | false | false | false | false | false | false | false | false | false | false | false | false | false | false |
| \|PWY-7720\| | ophiobolin F biosynthesis | false | false | false | false | false | false | false | false | false | false | false | false | false | false | false | false | false |
| \|PWY-7721\| | methyl phomopsenoate biosynthesis | false | false | false | false | false | false | false | false | false | false | false | false | false | false | false | false | false |
| \|PWY-7722\| | sulfoquinovose degradation II | false | false | false | false | false | false | false | false | false | false | false | false | false | false | false | false | false |
| \|PWY-7723\| | bacterial bioluminescence | false | false | false | false | false | false | false | false | false | false | false | false | false | false | false | false | false |
| \|PWY-7724\| | icosapentaenoate biosynthesis III (8-desaturase, mammals) | false | false | false | false | false | false | false | false | false | false | false | false | false | false | false | false | false |
| \|PWY-7725\| | arachidonate biosynthesis V (8-detaturase, mammals) | false | false | false | false | false | false | false | false | false | false | false | false | false | false | false | false | false |
| \|PWY-7726\| | (4Z,7Z,10Z,13Z,16Z)-docosapentaenoate biosynthesis (6-desaturase) | false | false | false | false | false | false | false | false | false | false | false | false | false | false | false | false | false |
| \|PWY-7727\| | docosahexaenoate biosynthesis IV (4-desaturase, mammals) | false | false | false | false | false | false | false | false | false | false | false | false | false | false | false | false | false |
| \|PWY-7728\| | (4Z,7Z,10Z,13Z,16Z)-docosa-4,7,10,13,16-pentaenoate biosynthesis II (4-desaturase) | false | false | false | false | false | false | false | false | false | false | false | false | false | false | false | false | false |
| \|PWY-7729\| | 5,6-dimethylbenzimidazole biosynthesis II (anaerobic) | false | false | false | false | false | false | false | false | false | false | false | false | false | false | false | false | false |
| \|PWY-7730\| | T-2 toxin biosynthesis | false | false | false | false | false | false | false | false | false | false | false | false | false | false | false | false | false |
| \|PWY-7733\| | 3-hydroxyquinaldate biosynthesis | false | false | false | false | false | false | false | false | false | false | false | false | false | false | false | false | false |
| \|PWY-7734\| | quinoxaline-2-carboxylate biosynthesis | false | false | false | false | false | false | false | false | false | false | false | false | false | false | false | false | false |
| \|PWY-7735\| | echinomycin and triostin A biosynthesis | false | false | false | false | false | false | false | false | false | false | false | false | false | false | false | false | false |
| \|PWY-7736\| | stellatic acid biosynthesis | false | false | false | false | false | false | false | false | false | false | false | false | false | false | false | false | false |
| \|PWY-7737\| | thiocoraline biosynthesis | false | false | false | false | false | false | false | false | false | false | false | false | false | false | false | false | false |
| \|PWY-7738\| | polyacyltrehalose biosynthesis | false | false | false | false | false | false | false | false | false | false | false | false | false | false | false | false | false |
| \|PWY-7739\| | aucuparin biosynthesis | false | false | false | false | false | false | false | false | false | false | false | false | false | false | false | false | false |
| \|PWY-7740\| | β-D-mannosyl phosphomycoketide biosynthesis | false | false | false | false | false | false | false | false | false | false | false | false | false | false | false | false | false |
| \|PWY-7741\| | phthiocerol biosynthesis | false | false | false | false | false | false | false | false | false | false | false | false | false | false | false | false | false |
| \|PWY-7742\| | phenolphthiocerol biosynthesis | false | false | false | false | false | false | false | false | false | false | false | false | false | false | false | false | false |
| \|PWY-7743\| | dimycocerosyl triglycosyl phenolphthiocerol biosynthesis | false | false | false | false | false | false | false | false | false | false | false | false | false | false | false | false | false |
| \|PWY-7744\| | dimycocerosyl phthiocerol biosynthesis | false | false | false | false | false | false | false | false | false | false | false | false | false | false | false | false | false |
| \|PWY-7745\| | p-HBAD biosynthesis | false | false | false | false | false | false | false | false | false | false | false | false | false | false | false | false | false |
| \|PWY-7746\| | mycobacterial sulfolipid biosynthesis | false | false | false | false | false | false | false | false | false | false | false | false | false | false | false | false | false |
| \|PWY-7747\| | diphenyl ethers degradation | false | false | false | false | false | false | false | false | false | false | false | false | false | false | false | false | false |
| \|PWY-7748\| | yatein biosynthesis I | false | false | false | false | false | false | false | false | false | false | false | false | false | false | false | false | false |
| \|PWY-7749\| | (-)-4'-demethyl-epipodophyllotoxin biosynthesis | false | false | false | false | false | false | false | false | false | false | false | false | false | false | false | false | false |
| \|PWY-7750\| | carbon monoxide oxidation to CO2 | false | false | false | false | false | false | false | false | false | false | false | false | false | false | false | false | false |
| \|PWY-7751\| | shinorine biosynthesis | false | false | false | false | false | false | false | false | false | false | false | false | false | false | false | false | false |
| \|PWY-7752\| | gadusol biosynthesis | false | false | false | false | false | false | false | false | false | false | false | false | false | false | false | false | false |
| \|PWY-7754\| | bile acid 7α-dehydroxylation | false | false | false | false | false | false | false | false | false | false | false | false | false | false | false | false | false |
| \|PWY-7755\| | iso-bile acids biosynthesis I | false | false | false | false | false | false | false | false | false | false | false | false | false | false | false | false | false |
| \|PWY-7756\| | iso-bile acids biosynthesis II | false | false | false | false | false | false | false | false | false | false | false | false | false | false | false | false | false |
| \|PWY-7757\| | bisphenol A degradation | false | false | false | false | false | false | false | false | false | false | false | false | false | false | false | false | false |
| \|PWY-7758\| | bacteriochlorophyll d biosynthesis | false | false | false | false | false | false | false | false | false | false | false | false | false | false | false | false | true |
| \|PWY-7759\| | bacteriochlorophyll c biosynthesis | false | false | false | false | false | false | false | false | false | false | false | false | false | false | false | false | false |
| \|PWY-7760\| | bacteriochlorophyll e biosynthesis | false | false | false | false | false | false | false | false | false | false | false | false | false | false | false | false | false |
| \|PWY-7762\| | bacteriochlorophyll b biosynthesis | false | false | false | false | false | false | false | false | false | false | false | false | false | false | false | false | false |
| \|PWY-7764\| | chlorophyll a biosynthesis III | false | false | false | false | false | false | false | false | false | false | false | false | false | false | false | false | false |
| \|PWY-7765\| | 3-hydroxy-4-methyl-anthranilate biosynthesis II | false | false | false | false | false | false | false | false | false | false | false | false | false | false | false | false | false |
| \|PWY-7766\| | heme b biosynthesis IV (Gram-positive bacteria) | false | false | false | false | false | false | false | false | false | false | false | false | false | false | false | false | false |
| \|PWY-7767\| | L-leucine degradation IV (reductive Stickland reaction) | false | false | false | false | false | false | false | false | false | false | false | false | false | false | false | false | false |
| \|PWY-7769\| | phosalacine biosynthesis | false | false | false | false | false | false | false | false | false | false | false | false | false | false | false | false | false |
| \|PWY-7770\| | indolmycin biosynthesis | false | false | false | false | false | false | false | false | false | false | false | false | false | false | false | false | false |
| \|PWY-7771\| | butachlor degradation | false | false | false | false | false | false | false | false | false | false | false | false | false | false | false | false | false |
| \|PWY-7772\| | γ-resorcylate degradation II | false | false | false | false | false | false | false | false | false | false | false | false | false | false | false | false | false |
| \|PWY-7773\| | γ-resorcylate degradation I | false | false | false | false | false | false | false | false | false | false | false | false | false | false | false | false | false |
| \|PWY-7774\| | propane degradation I | false | false | false | false | false | false | false | false | false | false | false | false | false | false | false | false | false |
| \|PWY-7775\| | propane degradation II | false | false | false | false | false | false | false | false | false | false | false | false | false | false | false | false | false |
| \|PWY-7776\| | ethene and chloroethene degradation | false | false | false | false | false | false | false | false | false | false | false | false | false | false | false | false | false |
| \|PWY-7777\| | isoprene degradation | false | false | false | false | false | false | false | false | false | false | false | false | false | false | false | false | false |
| \|PWY-7778\| | 2-methylpropene degradation | false | false | false | false | false | false | false | false | false | false | false | false | false | false | false | false | false |
| \|PWY-7779\| | methyl tert-butyl ether degradation | false | false | false | false | false | false | false | false | false | false | false | false | false | false | false | false | false |
| \|PWY-7780\| | butane degradation | false | false | false | false | false | false | false | false | false | false | false | false | false | false | false | false | false |
| \|PWY-7781\| | ω-sulfo-II-dihydromenaquinone-9 biosynthesis | false | false | false | false | false | false | false | false | false | false | false | false | false | false | false | false | false |
| \|PWY-7782\| | plasmalogen biosynthesis I (aerobic) | false | false | false | false | false | false | false | false | false | false | false | false | false | false | false | false | false |
| \|PWY-7783\| | plasmalogen degradation | false | false | false | false | false | false | false | false | false | false | false | false | false | false | false | false | false |
| \|PWY-7784\| | reductive acetyl coenzyme A pathway II (autotrophic methanogens) | false | false | false | false | false | false | false | false | false | false | false | false | false | false | false | false | false |
| \|PWY-7786\| | D-threitol degradation | false | false | false | false | false | false | false | false | false | false | false | false | false | false | false | false | false |
| \|PWY-7787\| | L-threitol degradation | false | false | false | false | false | false | false | false | false | false | false | false | false | false | false | false | false |
| \|PWY-7788\| | erythritol degradation II | false | false | false | false | false | false | false | false | false | false | false | false | false | false | false | false | false |
| \|PWY-7789\| | erythritol degradation I | false | false | false | false | false | false | false | false | false | false | false | false | false | false | false | false | false |
| \|PWY-7793\| | dimethyl sulfide biosynthesis from methionine | false | false | false | false | false | false | false | false | false | false | false | false | false | false | false | false | false |
| \|PWY-7794\| | polyethylene terephthalate degradation | false | false | false | false | false | false | false | false | false | false | false | false | false | false | false | false | false |
| \|PWY-7795\| | terephthalate degradation | false | false | false | false | false | false | false | false | false | false | false | false | false | false | false | false | false |
| \|PWY-7796\| | pentose phosphate pathway (oxidative branch) II | false | false | false | false | false | false | false | false | false | false | false | false | false | false | false | false | false |
| \|PWY-7797\| | nocardicin A biosynthesis | false | false | false | false | false | false | false | false | false | false | false | false | false | false | false | false | false |
| \|PWY-7798\| | protein S-nitrosylation and denitrosylation | false | false | false | false | false | false | false | false | false | false | false | false | false | false | false | false | false |
| \|PWY-7799\| | Arg/N-end rule pathway (eukaryotic) | false | false | false | false | false | false | false | false | false | false | false | false | false | false | false | false | false |
| \|PWY-7800\| | Ac/N-end rule pathway | false | false | false | false | false | false | false | false | false | false | false | false | false | false | false | false | false |
| \|PWY-7801\| | N-end rule pathway I (prokaryotic) | false | false | false | false | false | false | false | false | false | false | false | false | false | false | false | false | false |
| \|PWY-7802\| | N-end rule pathway II (prokaryotic) | false | false | false | false | false | false | false | false | false | false | false | false | false | false | false | false | false |
| \|PWY-7803\| | tRNA splicing II | false | false | false | false | false | false | false | false | false | false | false | false | false | false | false | false | false |
| \|PWY-7804\| | glyphosate degradation I | false | false | false | false | false | false | false | false | false | false | false | false | false | false | false | false | false |
| \|PWY-7805\| | (aminomethyl)phosphonate degradation | false | false | false | false | false | false | false | false | false | false | false | false | false | false | false | false | false |
| \|PWY-7806\| | glyphosate degradation II | false | false | false | false | false | false | false | false | false | false | false | false | false | false | false | false | false |
| \|PWY-7807\| | glyphosate degradation III | false | false | false | false | false | false | false | false | false | false | false | false | false | false | false | false | false |
| \|PWY-7808\| | tetracycline resistance | false | false | false | false | false | false | false | false | false | false | false | false | false | false | false | false | false |
| \|PWY-7810\| | chlorotetracycline biosynthesis | false | false | false | false | false | false | false | false | false | false | false | false | false | false | false | false | false |
| \|PWY-7811\| | 6-methylpretetramide biosynthesis | false | false | false | false | false | false | false | false | false | false | false | false | false | false | false | false | false |
| \|PWY-7812\| | tetracycline and oxytetracycline biosynthesis | false | false | false | false | false | false | false | false | false | false | false | false | false | false | false | false | false |
| \|PWY-7813\| | thiosulfate disproportionation III (quinone) | false | false | false | false | false | false | false | false | false | false | false | false | false | false | false | false | false |
| \|PWY-7814\| | dTDP-L-daunosamine biosynthesis | false | false | false | false | false | false | false | false | false | false | false | false | false | false | false | false | false |
| \|PWY-7815\| | poly(ribitol phosphate) wall teichoic acid biosynthesis I (B. subtilis) | false | false | false | false | false | false | false | false | false | false | false | false | false | false | false | false | false |
| \|PWY-7816\| | poly(ribitol phosphate) wall teichoic acid biosynthesis II (S. aureus) | false | false | false | false | false | false | false | false | false | false | false | false | false | false | false | false | false |
| \|PWY-7817\| | type I lipoteichoic acid biosynthesis (S. aureus) | false | false | false | false | false | false | false | false | false | false | false | false | false | false | false | false | false |
| \|PWY-7818\| | type IV lipoteichoic acid biosynthesis (S. pneumoniae) | false | false | false | false | false | false | false | false | false | false | false | false | false | false | false | false | false |
| \|PWY-7819\| | poly(3-O-β-D-glucopyranosyl-N-acetylgalactosamine 1-phosphate) wall teichoic acid biosynthesis | false | false | false | false | false | false | false | false | false | false | false | false | false | false | false | false | false |
| \|PWY-7820\| | teichuronic acid biosynthesis (B. subtilis 168) | false | false | false | false | false | false | false | false | false | false | false | false | false | false | false | false | false |
| \|PWY-7821\| | tunicamycin biosynthesis | false | false | false | false | false | false | false | false | false | false | false | false | false | false | false | false | false |
| \|PWY-7822\| | chitin degradation III (Serratia) | false | false | false | false | false | false | false | false | false | false | false | false | false | false | false | false | false |
| \|PWY-7823\| | chlorzoxazone degradation | false | false | false | false | false | false | false | false | false | false | false | false | false | false | false | false | false |
| \|PWY-7824\| | prunasin and amygdalin biosynthesis | false | false | false | false | false | false | false | false | false | false | false | false | false | false | false | false | false |
| \|PWY-7825\| | juglone degradation | false | false | false | false | false | false | false | false | false | false | false | false | false | false | false | false | false |
| \|PWY-7826\| | Amaryllidacea alkaloids biosynthesis | false | false | false | false | false | false | false | false | false | false | false | false | false | false | false | false | false |
| \|PWY-782\| | glycolipid desaturation | false | false | false | false | false | false | false | false | false | false | false | false | false | false | false | false | false |
| \|PWY-7831\| | ABH and Lewis epitopes biosynthesis from type 2 precursor disaccharide | false | false | false | false | false | false | false | false | false | false | false | false | false | false | false | false | false |
| \|PWY-7832\| | ABH and Lewis epitopes biosynthesis from type 1 precursor disaccharide | false | false | false | false | false | false | false | false | false | false | false | false | false | false | false | false | false |
| \|PWY-7833\| | biosynthesis of Lewis epitopes (H. pylori) | false | false | false | false | false | false | false | false | false | false | false | false | false | false | false | false | false |
| \|PWY-7836\| | ganglio-series glycosphingolipids biosynthesis | false | false | false | false | false | false | false | false | false | false | false | false | false | false | false | false | false |
| \|PWY-7837\| | i antigen and I antigen biosynthesis | false | false | false | false | false | false | false | false | false | false | false | false | false | false | false | false | false |
| \|PWY-7838\| | globo-series glycosphingolipids biosynthesis | false | false | false | false | false | false | false | false | false | false | false | false | false | false | false | false | false |
| \|PWY-7839\| | lacto-series glycosphingolipids biosynthesis | false | false | false | false | false | false | false | false | false | false | false | false | false | false | false | false | false |
| \|PWY-7840\| | gala-series glycosphingolipids biosynthesis | false | false | false | false | false | false | false | false | false | false | false | false | false | false | false | false | false |
| \|PWY-7841\| | neolacto-series glycosphingolipids biosynthesis | false | false | false | false | false | false | false | false | false | false | false | false | false | false | false | false | false |
| \|PWY-7842\| | UDP-yelosamine biosynthesis | false | false | false | false | false | false | false | false | false | false | false | false | false | false | false | false | false |
| \|PWY-7843\| | heme degradation IV | false | false | false | false | false | false | false | false | false | false | false | false | false | false | false | false | false |
| \|PWY-7844\| | heme degradation III | false | false | false | false | false | false | false | false | false | false | false | false | false | false | false | false | false |
| \|PWY-7845\| | heme degradation II | false | false | false | false | false | false | false | false | false | false | false | false | false | false | false | false | false |
| \|PWY-7846\| | heme degradation V | false | false | false | false | false | false | false | false | false | false | false | false | false | false | false | false | false |
| \|PWY-7847\| | heme degradation VI | false | false | false | false | false | false | false | false | false | false | false | false | false | false | false | false | false |
| \|PWY-7848\| | heme degradation VII | false | false | false | false | false | false | false | false | false | false | false | false | false | false | false | false | false |
| \|PWY-7849\| | urate conversion to allantoin III | false | false | false | false | false | false | false | false | false | false | false | false | false | false | false | false | false |
| \|PWY-7850\| | taurine biosynthesis II | false | false | false | false | false | false | false | false | false | false | false | false | false | false | false | false | false |
| \|PWY-7851\| | coenzyme A biosynthesis II (eukaryotic) | false | false | false | false | false | false | false | false | false | false | false | false | false | false | false | false | false |
| \|PWY-7852\| | 6-hydroxymethyl-dihydropterin diphosphate biosynthesis IV (Plasmodium) | false | false | false | false | false | false | false | false | false | false | false | false | false | false | false | false | false |
| \|PWY-7853\| | 6-hydroxymethyl-dihydropterin diphosphate biosynthesis V (Pyrococcus) | false | false | false | false | false | false | false | false | false | false | false | false | false | false | false | false | false |
| \|PWY-7854\| | crotonyl-CoA/ethylmalonyl-CoA/hydroxybutyryl-CoA cycle (engineered) | false | false | false | false | false | false | false | false | false | false | false | false | false | false | false | false | false |
| \|PWY-7855\| | ectoine degradation | false | false | false | false | false | false | false | false | false | false | false | false | false | false | false | false | false |
| \|PWY-7856\| | heme a biosynthesis | false | false | false | false | false | false | false | false | false | false | false | false | false | false | false | false | false |
| \|PWY-7857\| | adlupulone and adhumulone biosynthesis | false | false | false | false | false | false | false | false | false | false | false | false | false | false | false | false | false |
| \|PWY-7858\| | (5Z)-dodecenoate biosynthesis II | false | false | false | false | false | false | false | false | false | false | false | false | false | false | false | false | false |
| \|PWY-7859\| | jasmonoyl-L-isoleucine inactivation | false | false | false | false | false | false | false | false | false | false | false | false | false | false | false | false | false |

| | | | | | | | | | | | | | | | | | | |
|---|---|---|---|---|---|---|---|---|---|---|---|---|---|---|---|---|---|---|
| \|PWY-7860\| | nitric oxide biosynthesis III (bacteria) | false | false | false | false | false | false | false | false | false | false | false | false | false | false | false | false | false |
| \|PWY-7861\| | N-hydroxy-L-pipecolate biosynthesis | false | false | false | false | false | false | false | false | false | false | false | false | false | false | false | false | false |
| \|PWY-7862\| | D-altritol and galactitol degradation | false | false | false | false | false | false | false | false | false | false | false | false | false | false | false | false | false |
| \|PWY-7863\| | roseoflavin biosynthesis | false | false | false | false | false | false | false | false | false | false | false | false | false | false | false | false | false |
| \|PWY-7864\| | 8-methylmenaquinone biosynthesis | false | false | false | false | false | false | false | false | false | false | false | false | false | false | false | false | false |
| \|PWY-7865\| | coenzyme B/coenzyme M regeneration II (ferredoxin-dependent) | false | false | false | false | false | false | false | false | false | false | false | false | false | false | false | false | false |
| \|PWY-7866\| | coenzyme B/coenzyme M regeneration III (coenzyme F420-dependent) | false | false | false | false | false | false | false | false | false | false | false | false | false | false | false | false | false |
| \|PWY-7867\| | coenzyme B/coenzyme M regeneration IV (H2-dependent) | false | false | false | false | false | false | false | false | false | false | false | false | false | false | false | false | false |
| \|PWY-7868\| | coenzyme B/coenzyme M regeneration V (formate-dependent) | false | false | false | false | false | false | false | false | false | false | false | false | false | false | false | false | false |
| \|PWY-7869\| | cis-alkene biosynthesis | false | false | false | false | false | false | false | false | false | false | false | false | false | false | false | false | false |
| \|PWY-7871\| | D-threonate degradation | false | false | false | false | false | false | false | false | false | false | false | false | false | false | false | false | false |
| \|PWY-7872\| | D-erythronate degradation I | false | false | false | false | false | false | false | false | false | false | false | false | false | false | false | false | false |
| \|PWY-7873\| | D-erythronate degradation II | false | false | false | false | false | false | false | false | false | false | false | false | false | false | false | false | false |
| \|PWY-7874\| | L-threonate degradation | false | false | false | false | false | false | false | false | false | false | false | false | false | false | false | false | false |
| \|PWY-7878\| | A-factor γ-butyrolactone biosynthesis | false | false | false | false | false | false | false | false | false | false | false | false | false | false | false | false | false |
| \|PWY-7880\| | coelimycin P1 biosynthesis | false | false | false | false | false | false | false | false | false | false | false | false | false | false | false | false | false |
| \|PWY-7881\| | virginiae butanolide type γ-butyrolactones biosynthesis | false | false | false | false | false | false | false | false | false | false | false | false | false | false | false | false | false |
| \|PWY-7882\| | IM-2 type γ-butyrolactones biosynthesis | false | false | false | false | false | false | false | false | false | false | false | false | false | false | false | false | false |
| \|PWY-7883\| | peptidoglycan recycling II | false | false | false | false | false | false | false | false | false | false | false | false | false | false | false | false | false |
| \|PWY-7884\| | lipoprotein posttranslational modification | false | false | false | false | false | false | false | false | false | false | false | false | false | false | false | false | false |
| \|PWY-7885\| | phosphatidylinositol mannoside biosynthesis | false | false | false | false | false | false | false | false | false | false | false | false | false | false | false | false | false |
| \|PWY-7886\| | cell-surface glycoconjugate-linked phosphocholine biosynthesis | false | false | false | false | false | false | false | false | false | false | false | false | false | false | false | false | false |
| \|PWY-7887\| | protein SAMPylation and SAMP-mediated thiolation | false | false | false | false | false | false | false | false | false | false | false | false | false | false | false | false | false |
| \|PWY-7888\| | tRNA-uridine 2-thiolation (cytoplasmic) | false | false | false | false | false | false | false | false | false | false | false | false | false | false | false | false | false |
| \|PWY-7889\| | tRNA-uridine 2-thiolation (mammalian mitochondria) | false | false | false | false | false | false | false | false | false | false | false | false | false | false | false | false | false |
| \|PWY-7891\| | tRNA-uridine 2-thiolation (yeast mitochondria) | false | false | false | false | false | false | false | false | false | false | false | false | false | false | false | false | false |
| \|PWY-7892\| | tRNA-uridine 2-thiolation and selenation (bacteria) | false | false | false | false | false | false | false | false | false | false | false | false | false | false | false | false | false |
| \|PWY-7893\| | protein Pupylation and dePupylation | false | false | false | false | false | false | false | false | false | false | false | false | false | false | false | false | false |
| \|PWY-7894\| | procollagen hydroxylation and glycosylation | false | false | false | false | false | false | false | false | false | false | false | false | false | false | false | false | false |
| \|PWY-7895\| | ethionamide activation | false | false | false | false | false | false | false | false | false | false | false | false | false | false | false | false | false |
| \|PWY-7896\| | isoniazid activation | false | false | false | false | false | false | false | false | false | false | false | false | false | false | false | false | false |
| \|PWY-7897\| | flavonoid di-C-glucosylation | false | false | false | false | false | false | false | false | false | false | false | false | false | false | false | false | false |
| \|PWY-7898\| | 4,4'-disulfanediyldibutanoate degradation | false | false | false | false | false | false | false | false | false | false | false | false | false | false | false | false | false |
| \|PWY-7899\| | protein NEDDylation | false | false | false | false | false | false | false | false | false | false | false | false | false | false | false | false | false |
| \|PWY-7900\| | glycogen biosynthesis III (from α-maltose 1-phosphate) | false | false | false | false | false | false | false | false | false | false | false | false | false | false | false | false | false |
| \|PWY-7901\| | glucosinolate biosynthesis from tyrosine | false | false | false | false | false | false | false | false | false | false | false | false | false | false | false | false | false |
| \|PWY-7902\| | glucosylglycerol biosynthesis | false | false | false | false | false | false | false | false | false | false | false | false | false | false | false | false | false |
| \|PWY-7904\| | tRNA-uridine 2-thiolation (thermophilic bacteria) | false | false | false | false | false | false | false | false | false | false | false | false | false | false | false | false | false |
| \|PWY-7905\| | Escherichia coli serotype O:9a O antigen biosynthesis | false | false | false | false | false | false | false | false | false | false | false | false | false | false | false | false | false |
| \|PWY-7906\| | rhizobitoxine biosynthesis | false | false | false | false | false | false | false | false | false | false | false | false | false | false | false | false | false |
| \|PWY-7907\| | (2S,3E)-2-amino-4-methoxy-but-3-enoate biosynthesis | false | false | false | false | false | false | false | false | false | false | false | false | false | false | false | false | false |
| \|PWY-7908\| | formaldehyde oxidation V (bacillithiol-dependent) | false | false | false | false | false | false | false | false | false | false | false | false | false | false | false | false | false |
| \|PWY-7909\| | formaldehyde oxidation VII (THF pathway) | false | false | false | false | false | false | false | false | false | false | false | false | false | false | false | false | false |
| \|PWY-7910\| | homofuraneol biosynthesis | false | false | false | false | false | false | false | false | false | false | false | false | false | false | false | false | false |
| \|PWY-7911\| | 6'-dechloromelleolide F biosynthesis | false | false | false | false | false | false | false | false | false | false | false | false | false | false | false | false | false |
| \|PWY-7912\| | dinoflagellate bioluminescence | false | false | false | false | false | false | false | false | false | false | false | false | false | false | false | false | false |
| \|PWY-7913\| | firefly bioluminescence | false | false | false | false | false | false | false | false | false | false | false | false | false | false | false | false | false |
| \|PWY-7914\| | coral bioluminescence | false | false | false | false | false | false | false | false | false | false | false | false | false | false | true | false | false |
| \|PWY-7915\| | jellyfish bioluminescence | false | false | false | false | false | false | false | false | false | false | false | false | false | false | false | false | false |
| \|PWY-7916\| | squid bioluminescence | false | false | false | false | false | false | false | false | false | false | false | false | false | false | false | false | false |
| \|PWY-7917\| | pheomelanin biosynthesis | false | false | false | false | false | false | false | false | false | false | false | false | false | false | false | false | false |
| \|PWY-7918\| | protein N-glycosylation processing phase (endoplasmic reticulum, yeast) | false | false | false | false | false | false | false | false | false | false | false | false | false | false | false | false | false |
| \|PWY-7919\| | protein N-glycosylation processing phase (plants and animals) | false | false | false | false | false | false | false | false | false | false | false | false | false | false | false | false | false |
| \|PWY-7920\| | complex N-linked glycan biosynthesis (plants) | false | false | false | false | false | false | false | false | false | false | false | false | false | false | false | false | false |
| \|PWY-7921\| | protein O-mannosylation I (yeast) | false | false | false | false | false | false | false | false | false | false | false | false | false | false | false | false | false |
| \|PWY-7922\| | protein O-mannosylation II (mammals, core M1 and core M2) | false | false | false | false | false | false | false | false | false | false | false | false | false | false | false | false | false |
| \|PWY-7923\| | archaeosine biosynthesis II | false | false | false | false | false | false | false | false | false | false | false | false | false | false | false | false | false |
| \|PWY-7924\| | rubber degradation I | false | false | false | false | false | false | false | false | false | false | false | false | false | false | false | false | false |
| \|PWY-7925\| | rubber degradation II | false | false | false | false | false | false | false | false | false | false | false | false | false | false | false | false | false |
| \|PWY-7926\| | carbon disulfide oxidation III (metazoa) | false | false | false | false | false | false | false | false | false | false | false | false | false | false | false | false | false |
| \|PWY-7927\| | sulfide oxidation IV (mitochondria) | false | false | false | false | false | false | false | false | false | false | false | false | false | false | false | false | false |
| \|PWY-7928\| | lactate racemase nickel cofactor biosynthesis | false | false | false | false | false | false | false | false | false | false | false | false | false | false | false | false | false |
| \|PWY-7929\| | polybrominated phenols biosynthesis | false | false | false | false | false | false | false | false | false | false | false | false | false | false | false | false | false |
| \|PWY-7930\| | pyoluteorin biosynthesis | false | false | false | false | false | false | false | false | false | false | false | false | false | false | false | false | false |
| \|PWY-7931\| | brominated pyrroles biosynthesis | false | false | false | false | false | false | false | false | false | false | false | false | false | false | false | false | false |
| \|PWY-7932\| | polybrominated biphenyls and diphenyl ethers biosynthesis | false | false | false | false | false | false | false | false | false | false | false | false | false | false | false | false | false |
| \|PWY-7934\| | polybrominated dihydroxylated diphenyl ethers biosynthesis | false | false | false | false | false | false | false | false | false | false | false | false | false | false | false | false | false |
| \|PWY-7935\| | spongiadioxin C biosynthesis | false | false | false | false | false | false | false | false | false | false | false | false | false | false | false | false | false |
| \|PWY-7936\| | psilocybin biosynthesis | false | false | false | false | false | false | false | false | false | false | false | false | false | false | false | false | false |
| \|PWY-7937\| | fungal bioluminescence | false | false | false | false | false | false | false | false | false | false | false | false | false | false | false | false | false |
| \|PWY-7938\| | isorenieratene biosynthesis I (actinobacteria) | false | false | false | false | false | false | false | false | false | false | false | false | false | false | false | false | false |
| \|PWY-7939\| | chlorobactene biosynthesis | false | false | false | false | false | false | false | false | false | false | false | false | false | false | false | false | false |
| \|PWY-7940\| | lauryl-hydroxychlorobactene glucoside biosynthesis | false | false | false | false | false | false | false | false | false | false | false | false | false | false | false | false | false |
| \|PWY-7941\| | isorenieratene biosynthesis II (Chlorobiaceae) | false | false | false | false | false | false | false | false | false | false | false | false | false | false | false | false | false |
| \|PWY-7942\| | 5-oxo-L-proline metabolism | false | false | false | false | false | false | false | false | false | false | false | false | false | false | false | false | false |
| \|PWY-7943\| | lycopadiene biosynthesis | false | false | false | false | false | false | false | false | false | false | false | false | false | false | false | false | false |
| \|PWY-7944\| | bacterioruberin biosynthesis | false | false | false | false | false | false | false | false | false | false | false | false | false | false | false | false | false |
| \|PWY-7945\| | 2,2'-bis-(4-hydroxy-3-methybut-2-enyl)-β,β-carotene monoglucoside biosynthesis | false | false | false | false | false | false | false | false | false | false | false | true | false | false | false | false | false |
| \|PWY-7946\| | sarcinaxanthin diglucoside biosynthesis | false | false | false | false | false | false | false | false | false | false | false | false | false | false | false | false | false |
| \|PWY-7947\| | flexixanthin biosynthesis | false | false | false | false | false | false | false | false | false | false | false | false | false | false | false | false | false |
| \|PWY-7948\| | 4-oxopentanoate degradation | false | false | false | false | false | false | false | false | false | false | false | false | false | false | false | false | false |
| \|PWY-7949\| | diadinoxanthin and diatoxanthin interconversion | false | false | false | false | false | false | false | false | false | false | false | false | false | false | false | false | false |
| \|PWY-7950\| | diadinoxanthin and fucoxanthin biosynthesis | false | false | false | false | false | false | false | false | false | false | false | false | false | false | false | false | false |
| \|PWY-7951\| | ellagic acid degradation to urolithins | false | false | false | false | false | false | false | false | false | false | false | false | false | false | false | false | false |
| \|PWY-7953\| | UDP-N-acetylmuramoyl-pentapeptide biosynthesis III (meso-diaminopimelate containing) | false | false | false | false | false | false | false | false | false | false | false | false | false | false | false | false | false |
| \|PWY-7954\| | tabtoxinine-β-lactam biosynthesis | false | false | false | false | false | false | false | false | false | false | false | false | false | false | false | false | false |
| \|PWY-7955\| | paerucumarin biosynthesis | false | false | false | false | false | false | false | false | false | false | false | false | false | false | false | false | false |
| \|PWY-7956\| | 3-[(E)-2-isocyanoethenyl]-1H-indole biosynthesis | false | false | false | false | false | false | false | false | false | false | false | false | false | false | false | false | false |
| \|PWY-7957\| | hapalindole H biosynthesis | false | false | false | false | false | false | false | false | false | false | false | false | false | false | false | false | false |
| \|PWY-7958\| | 12-epi-fischerindole biosynthesis | false | false | false | false | false | false | false | false | false | false | false | false | false | false | false | false | false |
| \|PWY-7959\| | 12-epi-hapalindole biosynthesis | false | false | false | false | false | false | false | false | false | false | false | false | false | false | false | false | false |
| \|PWY-7960\| | rhabduscin biosynthesis | false | false | false | false | false | false | false | false | false | false | false | false | false | false | false | false | false |
| \|PWY-7961\| | phenyl adenosylcobamide biosynthesis from adenosylcobinamide-GDP | false | false | false | false | false | false | false | false | false | false | false | false | false | false | false | false | false |
| \|PWY-7962\| | adenosylcobinamide-GDP biosynthesis from cobyrinate a,c-diamide | false | false | false | false | false | false | false | false | false | false | false | false | false | false | false | false | false |
| \|PWY-7963\| | 4-methylphenyl adenosylcobamide biosynthesis from adenosylcobinamide-GDP | false | false | false | false | false | false | false | false | false | false | false | false | false | false | false | false | false |
| \|PWY-7964\| | adeninyl adenosylcobamide biosynthesis from adenosylcobinamide-GDP | false | false | false | false | false | false | false | false | false | false | false | false | false | false | false | false | false |
| \|PWY-7965\| | 2-methyladeninyl adenosylcobamide biosynthesis from adenosylcobinamide-GDP | false | false | false | false | false | false | false | false | false | false | false | false | false | false | false | false | false |
| \|PWY-7966\| | 5-methoxy-6-methylbenzimidazolyl adenosylcobamide biosynthesis from adenosylcobinamide-GDP | false | false | false | false | false | false | false | false | false | false | false | false | false | false | false | false | false |
| \|PWY-7967\| | 5-methoxybenzimidazolyl adenosylcobamide biosynthesis from adenosylcobinamide-GDP | false | false | false | false | false | false | false | false | false | false | false | false | false | false | false | false | false |
| \|PWY-7968\| | 5-methylbenzimidazolyl adenosylcobamide biosynthesis from adenosylcobinamide-GDP | false | false | false | false | false | false | false | false | false | false | false | false | false | false | false | false | false |
| \|PWY-7969\| | 5-hydroxybenzimidazolyl adenosylcobamide biosynthesis from adenosylcobinamide-GDP | false | false | false | false | false | false | false | false | false | false | false | false | false | false | false | false | false |
| \|PWY-7970\| | benzimidazolyl adenosylcobamide biosynthesis from adenosylcobinamide-GDP | false | false | false | false | false | false | false | false | false | false | false | false | false | false | false | false | false |
| \|PWY-7971\| | adenosylcobinamide-GDP salvage from cobinamide I | false | false | false | false | false | false | false | false | false | false | false | false | false | false | false | false | false |
| \|PWY-7972\| | adenosylcobinamide-GDP salvage from cobinamide II | false | false | false | false | false | false | false | false | false | false | false | false | false | false | false | false | false |
| \|PWY-7974\| | cobalamin salvage (eukaryotic) | false | false | false | false | false | false | false | false | false | false | false | false | false | false | false | false | false |
| \|PWY-7975\| | adenosylcobalamin biosynthesis from adenosylcobinamide-GDP II | false | false | false | false | false | false | false | false | false | false | false | false | false | false | false | false | false |
| \|PWY-7978\| | N-methylpyrrolidone degradation | false | false | false | false | false | false | false | false | false | false | false | false | false | false | false | false | false |
| \|PWY-7979\| | protein O-mannosylation III (mammals, core M3) | false | false | false | false | false | false | false | false | false | false | false | false | false | false | false | false | false |
| \|PWY-7981\| | α-dystroglycan glycosylation | false | false | false | false | false | false | false | false | false | false | false | false | false | false | false | false | false |
| \|PWY-7982\| | pinoresinol degradation | false | false | false | false | false | false | false | false | false | false | false | false | false | false | false | false | false |
| \|PWY-7983\| | pseudomonine biosynthesis | false | false | false | false | false | false | false | false | false | false | false | false | false | false | false | false | false |
| \|PWY-7984\| | acinetobactin biosynthesis | false | false | false | false | false | false | false | false | false | false | false | false | false | false | false | false | false |
| \|PWY-7985\| | oxalate degradation VI | false | false | false | false | false | false | false | false | false | false | false | false | false | false | false | false | false |
| \|PWY-7986\| | anguibactin biosynthesis | false | false | false | false | false | false | false | false | false | false | false | false | false | false | false | false | false |
| \|PWY-7987\| | vanchrobactin biosynthesis | false | false | false | false | false | false | false | false | false | false | false | false | false | false | false | false | false |
| \|PWY-7988\| | baumannoferrin biosynthesis | false | false | false | false | false | false | false | false | false | false | false | false | false | false | false | false | false |
| \|PWY-7989\| | acinetoferrin biosynthesis | false | false | false | false | false | false | false | false | false | false | false | false | false | false | false | false | false |
| \|PWY-7990\| | staphyloferrin A biosynthesis | false | false | false | false | false | false | false | false | false | false | false | false | false | false | false | false | false |
| \|PWY-7991\| | toxoflavin biosynthesis | false | false | false | false | false | false | false | false | false | false | false | false | false | false | false | false | false |
| \|PWY-7993\| | β-dihydromenaquinone-9 biosynthesis | false | false | false | false | false | false | false | false | false | false | false | false | false | false | false | false | false |
| \|PWY-7994\| | dolabralexins biosynthesis | false | false | false | false | false | false | false | false | false | false | false | false | false | false | false | false | false |
| \|PWY-7995\| | tricin biosynthesis | false | false | false | false | false | false | false | false | false | false | false | false | false | false | false | false | false |
| \|PWY-7996\| | menaquinol-4 biosynthesis I | false | false | false | false | false | false | false | false | false | false | false | false | false | false | false | false | false |
| \|PWY-7997\| | demethylmenaquinol-4 biosynthesis | false | false | false | false | false | false | false | false | false | false | false | false | false | false | false | false | false |
| \|PWY-7998\| | menaquinol-4 biosynthesis II | false | false | false | false | false | false | false | false | false | false | false | false | false | false | false | false | false |
| \|PWY-7999\| | vitamin K-epoxide cycle | false | false | false | false | false | false | false | false | false | false | false | false | false | false | false | false | false |
| \|PWY-8000\| | vitamin K degradation | false | false | false | false | false | false | false | false | false | false | false | false | false | false | false | false | false |
| \|PWY-8002\| | 4-coumarate degradation (aerobic) | false | false | false | false | false | false | false | false | false | false | false | false | false | false | false | false | false |
| \|PWY-8003\| | trans-caffeate degradation (aerobic) | false | false | false | false | false | false | false | false | false | false | false | false | false | false | false | false | false |
| \|PWY-8006\| | plant arabinogalactan type II degradation | false | false | false | false | false | false | false | false | false | false | false | false | false | false | false | false | false |
| \|PWY-8007\| | staphylopine biosynthesis | false | false | false | false | false | false | false | false | false | false | false | false | false | false | false | false | false |
| \|PWY-8008\| | staphyloferrin B biosynthesis | false | false | false | false | false | false | false | false | false | false | false | false | false | false | false | false | false |
| \|PWY-8009\| | L-cysteine biosynthesis VIII (Thermococcus kodakarensis) | false | false | false | false | false | false | false | false | false | false | false | false | false | false | false | false | false |
| \|PWY-8010\| | L-cysteine biosynthesis IX (Trichomonas vaginalis) | false | false | false | false | false | false | false | false | false | false | false | false | false | false | false | false | false |
| \|PWY-8011\| | L-serine biosynthesis II | false | false | false | false | false | false | false | false | false | false | false | false | false | false | false | false | false |
| \|PWY-8012\| | mupirocin biosynthesis | false | false | false | false | false | false | false | false | false | false | false | false | false | false | false | false | false |
| \|PWY-8013\| | UDP-N-acetyl-D-galactosamine biosynthesis III | false | false | false | false | false | false | false | false | false | false | false | false | false | false | false | false | false |
| \|PWY-8014\| | L-phenylalanine degradation VI (reductive Stickland reaction) | false | false | false | false | false | false | false | false | false | false | false | false | false | false | false | false | false |
| \|PWY-8015\| | glycine degradation (reductive Stickland reaction) | false | false | false | false | false | false | false | false | false | false | false | false | false | false | false | false | false |
| \|PWY-8016\| | L-tyrosine degradation V (reductive Stickland reaction) | false | false | false | false | false | false | false | false | false | false | false | false | false | false | false | false | false |
| \|PWY-8018\| | itaconate biosynthesis II | false | false | false | false | false | false | false | false | false | false | false | false | false | false | false | false | false |
| \|PWY-8019\| | 2-hydroxyparaconate and itatartarate biosynthesis | false | false | false | false | false | false | false | false | false | false | false | false | false | false | false | false | false |
| \|PWY-801\| | homocysteine and cysteine interconversion | false | false | false | false | false | false | false | false | false | false | false | false | false | false | false | false | false |
| \|PWY-8020\| | D-xylose degradation V | false | false | false | false | false | false | false | false | false | false | false | false | false | false | false | false | false |
| \|PWY-8022\| | salmochelin degradation | false | false | false | false | false | false | false | false | false | false | false | false | false | false | false | false | false |
| \|PWY-8023\| | salmochelin biosynthesis | false | false | false | false | false | false | false | false | false | false | false | false | false | false | false | false | true |
| \|PWY-8024\| | 4-hydroxyindole-3-carbonyl nitrile biosynthesis | false | false | false | false | false | false | false | false | false | false | false | false | false | false | false | false | false |
| \|PWY-8025\| | cyanuric acid degradation I | false | false | false | false | false | false | false | false | false | false | false | false | false | false | false | false | false |
| \|PWY-8026\| | sterol biosynthesis (methylotrophs) | false | false | false | false | false | false | false | false | false | false | false | false | false | false | false | false | false |
| \|PWY-8027\| | parkeol biosynthesis | false | false | false | false | false | false | false | false | false | false | false | false | false | false | false | false | false |
| \|PWY-8028\| | cycloartenol biosynthesis | false | false | false | false | false | false | false | false | false | false | false | false | false | false | false | false | false |
| \|PWY-8029\| | limonene degradation IV (anaerobic) | false | false | false | false | false | false | false | false | false | false | false | false | false | false | false | false | false |
| \|PWY-8030\| | coptisine biosynthesis | false | false | false | false | false | false | false | false | false | false | false | false | false | false | false | false | false |
| \|PWY-8031\| | epiberberine biosynthesis | false | false | false | false | false | false | false | false | false | false | false | false | false | false | false | false | false |
| \|PWY-8032\| | chloramphenicol biosynthesis | false | false | false | false | false | false | false | false | false | false | false | false | false | false | false | false | false |
| \|PWY-8033\| | aureothin biosynthesis | false | false | false | false | false | false | false | false | false | false | false | false | false | false | false | false | false |
| \|PWY-8034\| | spectinabilin biosynthesis | false | false | false | false | false | false | false | false | false | false | false | false | false | false | false | false | false |
| \|PWY-8035\| | streptovaricin biosynthesis | false | false | false | false | false | false | false | false | false | false | false | false | false | false | false | false | false |
| \|PWY-8036\| | chaxamycin biosynthesis | false | false | false | false | false | false | false | false | false | false | false | false | false | false | false | false | false |
| \|PWY-8037\| | saliniketal A biosynthesis | false | false | false | false | false | false | false | false | false | false | false | false | false | false | false | false | false |
| \|PWY-8038\| | naphthomycin biosynthesis | false | false | false | false | false | false | false | false | false | false | false | false | false | false | false | false | false |
| \|PWY-8039\| | mitomycin biosynthesis | false | false | false | false | false | false | false | false | false | false | false | false | false | false | false | false | false |
| \|PWY-8040\| | ansatrienin biosynthesis | false | false | false | false | false | false | false | false | false | false | false | false | false | false | false | false | false |
| \|PWY-8041\| | ultra-long-chain fatty acid biosynthesis | false | false | false | false | false | false | false | false | false | false | false | false | false | false | false | false | false |
| \|PWY-8042\| | acylceramide biosynthesis and processing | false | false | false | false | false | false | false | false | false | false | false | false | false | false | false | false | false |
| \|PWY-8043\| | ophthalmate biosynthesis | false | false | false | false | false | false | false | false | false | false | false | false | false | false | false | false | false |
| \|PWY-8044\| | cyclohexane-1-carboxyl-CoA biosynthesis | false | false | false | false | false | false | false | false | false | false | false | false | false | false | false | false | false |
| \|PWY-8045\| | cylindrospermopsin biosynthesis | false | false | false | false | false | false | false | false | false | false | false | false | false | false | false | false | false |
| \|PWY-8047\| | bryostatin biosynthesis | false | false | false | false | false | false | false | false | false | false | false | false | false | false | false | false | false |
| \|PWY-8049\| | pederin biosynthesis | false | false | false | false | false | false | false | false | false | false | false | false | false | false | false | false | false |
| \|PWY-804\| | glycolate degradation II | false | false | false | false | false | false | false | false | false | false | false | false | false | false | false | false | false |
| \|PWY-8051\| | anandamide biosynthesis I | false | false | false | false | false | false | false | false | false | false | false | false | false | false | false | false | false |
| \|PWY-8052\| | 2-arachidonoylglycerol biosynthesis | false | false | false | false | false | false | false | false | false | false | false | false | false | false | false | false | false |
| \|PWY-8053\| | anandamide biosynthesis II | false | false | false | false | false | false | false | false | false | false | false | false | false | false | false | false | false |
| \|PWY-8055\| | palmitoyl ethanolamide biosynthesis | false | false | false | false | false | false | false | false | false | false | false | false | false | false | false | false | false |
| \|PWY-8056\| | anandamide lipoxygenation | false | false | false | false | false | false | false | false | false | false | false | false | false | false | false | false | false |
| \|PWY-8057\| | pyruvoyl group formation from L-serine | false | false | false | false | false | false | false | false | false | false | false | false | false | false | false | false | false |
| \|PWY-8058\| | 2-deoxy-D-ribose degradation II | false | false | false | false | false | false | false | false | false | false | false | false | false | false | false | false | false |
| \|PWY-8061\| | 8-oxo-(d)GTP detoxification II | false | false | false | false | false | false | false | false | false | false | false | false | false | false | false | false | false |
| \|PWY-8062\| | sulfoacetaldehyde degradation IV | false | false | false | false | false | false | false | false | false | false | false | false | false | false | false | false | false |
| \|PWY-8063\| | flavin-N5-oxide biosynthesis | false | false | false | false | false | false | false | false | false | false | false | false | false | false | false | false | false |
| \|PWY-8064\| | sodorifen biosynthesis | false | false | false | false | false | false | false | false | false | false | false | false | false | false | false | false | false |
| \|PWY-8065\| | chlorpyrifos degradation | false | false | false | false | false | false | false | false | false | false | false | false | false | false | false | false | false |
| \|PWY-8066\| | methylphosphonate degradation III | false | false | false | false | false | false | false | false | false | false | false | false | false | false | false | false | false |
| \|PWY-8067\| | methylphosphonate biosynthesis | false | false | false | false | false | false | false | false | false | false | false | false | false | false | false | false | false |
| \|PWY-8069\| | nepetalactone biosynthesis | false | false | false | false | false | false | false | false | false | false | false | false | false | false | false | false | false |
| \|PWY-8071\| | N-3-oxalyl-L-2,3-diaminopropanoate biosynthesis | false | false | false | false | false | false | false | false | false | false | false | false | false | false | false | false | false |
| \|PWY-8073\| | lipid IVA biosynthesis (P. putida) | false | false | false | false | false | false | false | false | false | false | false | false | false | false | false | false | false |
| \|PWY-8074\| | Kdo transfer to lipid IVA (P. putida) | false | false | false | false | false | false | false | false | false | false | false | false | false | false | false | false | false |
| \|PWY-8075\| | (Kdo)2-lipid A biosynthesis (P. putida) | false | false | false | false | false | false | false | false | false | false | false | false | false | false | false | false | false |
| \|PWY-8076\| | 8-O-methylated benzoxazinoid glucoside biosynthesis | false | false | false | false | false | false | false | false | false | false | false | false | false | false | false | false | false |
| \|PWY-8077\| | ginkgotoxin biosynthesis | false | false | false | false | false | false | false | false | false | false | false | false | false | false | false | true | false |
| \|PWY-8078\| | quebrachitol biosynthesis | false | false | false | false | false | false | false | false | false | false | false | false | false | false | false | false | false |
| \|PWY-8080\| | kainate biosynthesis | false | false | false | false | false | false | false | false | false | false | false | false | false | false | false | false | false |
| \|PWY-8081\| | leucine-derived hydroxynitrile glucoside biosynthesis | false | false | false | false | false | false | false | false | false | false | false | false | false | false | false | false | false |
| \|PWY-8082\| | glycine lipid biosynthesis | false | false | false | false | false | false | false | false | false | false | false | false | false | false | false | false | false |
| \|PWY-8083\| | domoic acid biosynthesis | false | false | false | false | false | false | false | false | false | false | false | false | false | false | false | false | false |
| \|PWY-8084\| | daurichromenate biosynthesis | false | false | false | false | false | false | false | false | false | false | false | false | false | false | false | false | false |
| \|PWY-8085\| | picolinate degradation | false | false | false | false | false | false | false | false | false | false | false | false | false | false | false | false | false |
| \|PWY-8086\| | (S)-lactate fermentation to propanoate, acetate and hydrogen | false | false | false | false | false | false | false | false | false | false | false | false | false | false | false | false | false |
| \|PWY-8087\| | indole-3-acetate degradation II | false | false | false | false | false | false | false | false | false | false | false | false | false | false | false | false | false |
| \|PWY-8089\| | D-apiose degradation I | false | false | false | false | false | false | false | false | false | false | false | false | false | false | false | false | false |
| \|PWY-8090\| | D-apionate degradation II (RLP decarboxylase) | false | false | false | false | false | false | false | false | false | false | false | false | false | false | false | false | false |
| \|PWY-8091\| | D-apionate degradation I (xylose isomerase family decarboxylase) | false | false | false | false | false | false | false | false | false | false | false | false | false | false | false | false | false |
| \|PWY-8092\| | D-apionate degradation III (RLP transcarboxylase/hydrolase) | false | false | false | false | false | false | false | false | false | false | false | false | false | false | false | false | false |
| \|PWY-8093\| | D-apiose degradation II (to D-apionate) | false | false | false | false | false | false | false | false | false | false | false | false | false | false | false | false | false |
| \|PWY-8095\| | 5-methoxy-6-methylbenzimidazole biosynthesis (anaerobic) | false | false | false | false | false | false | false | false | false | false | false | false | false | false | false | false | false |
| \|PWY-8096\| | 5-methoxybenzimidazole biosynthesis (anaerobic) | false | false | false | false | false | false | false | false | false | false | false | false | false | false | false | false | false |
| \|PWY-8098\| | cyclic 2,3-bisphosphoglycerate biosynthesis | false | false | false | false | false | false | false | false | false | false | false | false | false | false | false | false | false |
| \|PWY-8099\| | tetrahydropteridine recycling | false | false | false | false | false | false | false | false | false | false | false | false | false | false | false | false | false |
| \|PWY-8100\| | crotonosine biosynthesis | false | false | false | false | false | false | false | false | false | false | false | false | false | false | false | false | false |
| \|PWY-8101\| | arsenate detoxification II | false | false | false | false | false | false | false | false | false | false | false | false | false | false | false | false | false |
| \|PWY-8103\| | kavain biosynthesis | false | false | false | false | false | false | false | false | false | false | false | false | false | false | false | false | false |
| \|PWY-8104\| | yangonin biosynthesis | false | false | false | false | false | false | false | false | false | false | false | false | false | false | false | false | false |
| \|PWY-8106\| | queuosine biosynthesis III (queuosine salvage) | false | false | false | false | false | false | false | false | false | false | false | false | false | false | false | false | false |
| \|PWY-8107\| | methanogenesis from glycine betaine | false | false | false | false | false | false | false | false | false | false | false | false | false | false | false | false | false |
| \|PWY-8108\| | glycine betaine degradation IV | false | false | false | false | false | false | false | false | false | false | false | false | false | false | false | false | false |
| \|PWY-8109\| | proline betaine degradation II | false | false | false | false | false | false | false | false | false | false | false | false | false | false | false | false | false |
| \|PWY-8110\| | L-dopa degradation II (bacterial) | false | false | false | false | false | false | false | false | false | false | false | false | false | false | false | false | false |
| \|PWY-8111\| | carbaryl degradation | false | false | false | false | false | false | false | false | false | false | false | false | false | false | false | false | false |
| \|PWY-8112\| | factor 420 biosynthesis I (archaea) | false | false | false | false | false | false | false | false | false | false | false | false | false | false | false | false | false |
| \|PWY-8113\| | 3PG-factor 420 biosynthesis | false | false | false | false | false | false | false | false | false | false | false | false | false | false | false | false | true |
| \|PWY-8114\| | branched-chain polyamines biosynthesis | false | false | false | false | false | false | false | false | false | false | false | false | false | false | false | false | false |
| \|PWY-8115\| | long-chain polyamine biosynthesis | false | false | false | false | false | false | false | false | false | false | false | false | false | false | false | false | false |
| \|PWY-8116\| | ochratoxin A biosynthesis | false | false | false | false | false | false | false | false | false | false | false | false | false | false | false | false | false |
| \|PWY-8117\| | colibactin biosynthesis | false | false | false | false | false | false | false | false | false | false | false | false | true | true | true | false | true |
| \|PWY-8118\| | patellamide A and C biosynthesis | false | false | false | false | false | false | false | false | false | false | false | false | false | false | false | false | false |
| \|PWY-8119\| | microcin B17 biosynthesis | false | false | false | false | false | false | false | false | false | false | false | false | false | false | false | false | false |
| \|PWY-8120\| | β-alanine degradation III | false | false | false | false | false | false | false | false | false | false | false | false | false | false | false | false | false |
| \|PWY-8121\| | 2-deoxy-D-glucose 6-phosphate degradation | false | false | false | false | false | false | false | false | false | false | false | false | false | false | false | false | false |
| \|PWY-8122\| | mycofactocin biosynthesis | false | false | false | false | false | false | false | false | false | false | false | false | false | false | false | false | false |
| \|PWY-8124\| | (2-trimethylamino)ethylphosphonate degradation | false | false | false | false | false | false | false | false | false | false | false | false | false | false | false | false | false |
| \|PWY-8125\| | mevalonate pathway IV (archaea) | false | false | false | false | false | false | false | false | false | false | false | false | false | false | false | false | false |
| \|PWY-8126\| | chlorophyll a2 biosynthesis | false | false | false | false | false | false | false | false | false | false | false | false | false | false | false | false | false |
| \|PWY-8127\| | chlorophyll b2 biosynthesis | false | false | false | false | false | false | false | false | false | false | false | false | false | false | false | false | false |
| \|PWY-8128\| | biotin biosynthesis from 8-amino-7-oxononanoate III | false | false | false | false | false | false | false | false | false | false | false | false | false | false | false | false | false |
| \|PWY-8129\| | cell-surface glycoconjugate-linked phosphonate biosynthesis | false | false | false | false | false | false | false | false | false | false | false | false | false | false | false | false | false |
| \|PWY-8130\| | 5'-deoxyadenosine degradation I | false | false | false | false | false | false | false | false | false | false | false | false | false | false | false | false | false |
| \|PWY-8131\| | 5'-deoxyadenosine degradation II | false | false | false | false | false | false | false | false | false | false | false | false | false | false | false | false | false |
| \|PWY-8132\| | 5-methyl-5-thio-α-D-ribose 1-phosphate degradation III | false | false | false | false | false | false | false | false | false | false | false | false | false | false | false | false | false |
| \|PWY-8133\| | succinoglycan biosynthesis | false | false | false | false | false | false | false | false | false | false | false | false | false | false | false | false | false |
| \|PWY-8134\| | bile acid 7β-dehydroxylation | false | false | false | false | false | false | false | false | false | false | false | false | false | false | false | false | false |
| \|PWY-8135\| | bile acids deconjugation | false | false | false | false | false | false | false | false | false | false | false | false | false | false | false | false | false |
| \|PWY-8137\| | NADPH repair (eukaryotes) | false | false | false | false | false | false | false | false | false | false | false | false | false | false | false | false | false |
| \|PWY-8138\| | polyphosphate metabolism | false | false | false | false | false | false | false | false | false | false | false | false | false | false | false | false | false |
| \|PWY-8139\| | CDP-6-deoxy-D-gulose biosynthesis | false | false | false | false | false | false | false | false | false | false | false | false | false | false | false | false | false |
| \|PWY-8140\| | serine racemization | false | false | false | false | false | false | false | false | false | false | false | false | false | false | false | false | false |
| \|PWY-8141\| | 9-decynoate biosynthesis | false | false | false | false | false | false | false | false | false | false | false | false | false | false | false | false | false |
| \|PWY-8142\| | L-ascorbate biosynthesis VI (plants, myo-inositol pathway) | false | false | false | false | false | false | false | false | false | false | false | false | false | false | false | false | false |
| \|PWY-8143\| | L-ascorbate biosynthesis VII (plants, D-galacturonate pathway) | false | false | false | false | false | false | false | false | false | false | false | false | false | false | false | false | false |
| \|PWY-8145\| | cytochrome c biogenesis (system III type) | false | false | false | false | false | false | false | false | false | false | false | false | false | false | false | false | false |
| \|PWY-8146\| | cytochrome c biogenesis (system II type) | false | false | false | false | false | false | false | false | false | false | false | false | false | false | false | false | false |
| \|PWY-8147\| | cytochrome c biogenesis (system I type) | false | false | false | false | false | false | false | false | false | false | false | false | false | false | false | false | false |
| \|PWY-8149\| | homospermidine biosynthesis II | false | false | false | false | false | false | false | false | false | false | false | false | false | false | false | false | false |
| \|PWY-8151\| | cholesterol degradation to androstenedione III (anaerobic) | false | false | false | false | false | false | false | false | false | false | false | false | false | false | false | false | false |
| \|PWY-8152\| | androstenedione degradation II (anaerobic) | false | false | false | false | false | false | false | false | false | false | false | false | false | false | false | false | false |
| \|PWY-8155\| | testosterone degradation (anaerobic) | false | false | false | false | false | false | false | false | false | false | false | false | false | false | false | false | false |
| \|PWY-8156\| | omphalotin biosynthesis | false | false | false | false | false | false | false | false | false | false | false | false | false | false | false | false | false |
| \|PWY-8157\| | avenanthramide biosynthesis | false | false | false | false | false | false | false | false | false | false | false | false | false | false | false | false | false |
| \|PWY-8158\| | ibotenate and muscimol biosynthesis | false | false | false | false | false | false | false | false | false | false | false | false | false | false | false | false | false |
| \|PWY-8161\| | D-phenylglycine degradation | false | false | false | false | false | false | false | false | false | false | false | false | false | false | false | false | false |
| \|PWY-8162\| | phenylethanol degradation | false | false | false | false | false | false | false | false | false | false | false | false | false | false | false | false | false |
| \|PWY-8163\| | cytidylyl copper-molybdenum cofactor biosynthesis | false | false | false | false | false | false | false | false | false | false | false | false | false | false | false | false | false |
| \|PWY-8164\| | bis(guanylyl molybdopterin) cofactor sulfurylation | false | false | false | false | false | false | false | false | false | false | false | false | false | false | false | false | false |
| \|PWY-8165\| | cytidylyl molybdenum cofactor sulfurylation | false | false | false | false | false | false | false | false | false | false | false | false | false | false | false | false | false |
| \|PWY-8166\| | emodin biosynthesis | false | false | false | false | false | false | false | false | false | false | false | false | false | false | false | false | false |
| \|PWY-8167\| | bis(tungstenpterin) cofactor biosynthesis | false | false | false | false | false | false | false | false | false | false | false | false | false | false | false | false | false |
| \|PWY-8168\| | bis(guanylyl tungstenpterin) cofactor biosynthesis | false | false | false | false | false | false | false | false | false | false | false | false | false | false | false | false | false |
| \|PWY-8171\| | molybdenum cofactor biosynthesis | false | false | false | false | false | false | false | false | false | false | false | false | false | false | false | false | false |
| \|PWY-8172\| | 10-methylstearate biosynthesis | false | false | false | false | false | false | false | false | false | false | false | false | false | false | false | false | false |
| \|PWY-8173\| | anteiso-branched-chain fatty acid biosynthesis | false | false | false | false | false | false | false | false | false | false | false | false | false | false | false | false | false |
| \|PWY-8174\| | odd iso-branched-chain fatty acid biosynthesis | false | false | false | false | false | false | false | false | false | false | false | false | false | false | false | false | false |
| \|PWY-8175\| | even iso-branched-chain fatty acid biosynthesis | false | false | false | false | false | false | false | false | false | false | false | false | false | false | false | false | false |
| \|PWY-8176\| | geraniol biosynthesis (cytosol) | false | false | false | false | false | false | false | false | false | false | false | false | false | false | false | false | false |
| \|PWY-8177\| | nerol biosynthesis | false | false | false | false | false | false | false | false | false | false | false | false | false | false | false | false | false |
| \|PWY-8179\| | platensimycin biosynthesis | false | false | false | false | false | false | false | false | false | false | false | false | false | false | false | false | false |
| \|PWY-8180\| | acrylate degradation II | false | false | false | false | false | false | false | false | false | false | false | false | false | false | false | false | false |
| \|PWY-8181\| | 2-methyl-branched fatty acid β-oxidation | false | false | false | false | false | false | false | false | false | false | false | false | false | false | false | false | false |
| \|PWY-8182\| | valproate β-oxidation | false | false | false | false | false | false | false | false | false | false | false | false | false | false | false | false | false |
| \|PWY-8183\| | L-valine degradation III (oxidative Stickland reaction) | false | false | false | false | false | false | false | false | false | false | false | false | false | false | false | false | false |
| \|PWY-8184\| | L-isoleucine degradation III (oxidative Stickland reaction) | false | false | false | false | false | false | false | false | false | false | false | false | false | false | false | false | false |
| \|PWY-8185\| | L-leucine degradation V (oxidative Stickland reaction) | false | false | false | false | false | false | false | false | false | false | false | false | false | false | false | false | false |
| \|PWY-8186\| | L-proline degradation II (reductive Stickland reaction) | false | false | false | false | false | false | false | false | false | false | false | false | false | false | false | false | false |
| \|PWY-8187\| | L-arginine degradation XIII (reductive Stickland reaction) | false | false | false | false | false | false | false | false | false | false | false | false | false | false | false | false | false |
| \|PWY-8189\| | L-alanine degradation V (oxidative Stickland reaction) | false | false | false | false | false | false | false | false | false | false | false | false | false | false | false | false | false |
| \|PWY-8190\| | L-glutamate degradation XI (reductive Stickland reaction) | false | false | false | false | false | false | false | false | false | false | false | false | false | false | false | false | false |
| \|PWY-8191\| | cholesterol biosynthesis (algae, late side-chain reductase) | false | false | false | false | false | false | false | false | false | false | false | false | false | false | false | false | false |
| \|PWY-8192\| | Salmonella enterica serotype O:4 O antigen biosynthesis (group B1) | false | false | false | false | false | false | false | false | false | false | false | false | false | false | false | false | false |
| \|PWY-8193\| | Salmonella enterica serotype O:8 O antigen biosynthesis | false | false | false | false | false | false | false | false | false | false | false | false | false | false | false | false | false |
| \|PWY-8195\| | Salmonella enterica serotype O:9 O antigen biosynthesis | false | false | false | false | false | false | false | false | false | false | false | false | false | false | false | false | false |
| \|PWY-8196\| | Salmonella enterica serotype O:3,10 O antigen biosynthesis | false | false | false | false | false | false | false | false | false | false | false | false | false | false | false | false | false |
| \|PWY-8197\| | Salmonella enterica serotype O:9,46 O antigen biosynthesis | false | false | false | false | false | false | false | false | false | false | false | false | false | false | false | false | false |
| \|PWY-8198\| | Salmonella enterica serotype O:2 O antigen biosynthesis | false | false | false | false | false | false | false | false | false | false | false | false | false | false | false | false | false |
| \|PWY-8199\| | Salmonella enterica serotype O:9,46,27 O antigen biosynthesis | false | false | false | false | false | false | false | false | false | false | false | false | false | false | false | false | false |
| \|PWY-81\| | toluene degradation to benzoyl-CoA (anaerobic) | false | false | false | false | false | false | false | false | false | false | false | false | false | false | false | false | false |
| \|PWY-8200\| | backdoor pathway of androgen biosynthesis | false | false | false | false | false | false | false | false | false | false | false | false | false | false | false | false | false |
| \|PWY-8202\| | 11-oxyandrogens biosynthesis | false | false | false | false | false | false | false | false | false | false | false | false | false | false | false | false | false |
| \|PWY-8203\| | 8-amino-7-oxononanoate biosynthesis IV | false | false | false | false | false | false | false | false | false | false | false | false | false | false | false | false | false |
| \|PWY-8204\| | Salmonella enterica serotype O:54 O antigen biosynthesis | false | false | false | false | false | false | false | false | false | false | false | false | false | false | false | false | false |
| \|PWY-8205\| | Escherichia coli serotype O:77/Salmonella enterica serotype O:6,14 O antigen biosynthesis | false | false | false | false | false | false | false | false | false | false | false | false | false | false | false | false | false |
| \|PWY-8206\| | Salmonella enterica serotype O:18 O antigen biosynthesis | false | false | false | false | false | false | false | false | false | false | false | false | false | false | false | false | false |
| \|PWY-8207\| | Escherichia coli serotype O:85/Salmonella enterica serotype O:17 O antigen biosynthesis | false | false | false | false | false | false | false | false | false | false | false | false | false | false | false | false | false |
| \|PWY-8208\| | Salmonella enterica serotype O:6,7 O antigen biosynthesis | false | false | false | false | false | false | false | false | false | false | false | false | false | false | false | false | false |
| \|PWY-8209\| | Escherichia coli serotype O:15 O antigen biosynthesis | false | false | false | false | false | false | false | false | false | false | false | false | false | false | false | false | false |
| \|PWY-8210\| | pyrrolomycin biosynthesis | false | false | false | false | false | false | false | false | false | false | false | false | false | false | false | false | false |
| \|PWY-8211\| | Escherichia coli serotype O:7 O antigen biosynthesis | false | false | false | false | false | false | false | false | false | false | false | false | false | false | false | false | false |
| \|PWY-8212\| | Escherichia coli serotype O:8 O antigen biosynthesis | false | false | false | false | false | false | false | false | false | false | false | false | false | false | false | false | false |
| \|PWY-8213\| | sulfoquinovose degradation III | false | false | false | false | false | false | false | false | false | false | false | false | false | false | false | false | false |
| \|PWY-8216\| | strychnine biosynthesis | false | false | false | false | false | false | false | false | false | false | false | false | false | false | false | false | false |
| \|PWY-8217\| | Escherichia coli serotype O:55/Salmonella enterica serotype O:50 O antigen biosynthesis | false | false | false | false | false | false | false | false | false | false | false | false | false | false | false | false | false |
| \|PWY-8218\| | Escherichia coli serotype O:56 O antigen biosynthesis | false | false | false | false | false | false | false | false | false | false | false | false | false | false | false | false | false |
| \|PWY-8219\| | Escherichia coli serotype O:107 O antigen biosynthesis | false | false | false | false | false | false | false | false | false | false | false | false | false | false | false | false | false |
| \|PWY-8220\| | Escherichia coli serotype O:117 O antigen biosynthesis | false | false | false | false | false | false | false | false | false | false | false | false | false | false | false | false | false |
| \|PWY-8221\| | Escherichia coli serotype O:111/Salmonella enterica serotype O:35 O antigen biosynthesis | false | false | false | false | false | false | false | false | false | false | false | false | false | false | false | false | false |
| \|PWY-8222\| | Escherichia coli serotype O:128 O antigen biosynthesis | false | false | false | false | false | false | false | false | false | false | false | false | false | false | false | false | false |
| \|PWY-8224\| | Escherichia coli serotype O:157/Salmonella enterica serotype O:30 O antigen biosynthesis | false | false | false | false | false | false | false | false | false | false | false | false | false | false | false | false | false |
| \|PWY-8225\| | GDP-N-acetyl-α-D-perosamine biosynthesis | false | false | false | false | false | false | false | false | false | false | false | false | false | false | false | false | false |
| \|PWY-8226\| | Escherichia coli serotype O:152 O antigen biosynthesis | false | false | false | false | false | false | false | false | false | false | false | false | false | false | false | false | false |
| \|PWY-8228\| | Porphyromonas gingivalis O-LPS antigen biosynthesis | false | false | false | false | false | false | false | false | false | false | false | false | false | false | false | false | false |
| \|PWY-8229\| | UDP-3-acetamido-2,3-dideoxy-2-(seryl)amino-α-D-glucuronamide biosynthesis | false | false | false | false | false | false | false | false | false | false | false | false | false | false | false | false | false |
| \|PWY-822\| | fructan biosynthesis | false | false | false | false | false | false | false | false | false | false | false | false | false | false | false | false | false |
| \|PWY-8230\| | Salmonella enterica serotype O:13 O antigen biosynthesis | false | false | false | false | false | false | false | false | false | false | false | false | false | false | false | false | false |
| \|PWY-8231\| | Escherichia coli serotype O:127 O antigen biosynthesis | false | false | false | false | false | false | false | false | false | false | false | false | false | false | false | false | false |
| \|PWY-8232\| | Shigella boydii serotype 6 O antigen biosynthesis | false | false | false | false | false | false | false | false | false | false | false | false | false | false | false | false | false |
| \|PWY-8233\| | Escherichia coli serotype O:169 O antigen biosynthesis | false | false | false | false | false | false | false | false | false | false | false | false | false | false | false | false | false |
| \|PWY-8234\| | Escherichia coli serotype O:183/Shigella boydii serotype O:10 O antigen biosynthesis | false | false | false | false | false | false | false | false | false | false | false | false | true | true | true | true | true |
| \|PWY-8235\| | Escherichia coli serotype O:21/Salmonella enterica serotype O:38 O antigen biosynthesis | false | false | false | false | false | false | false | false | false | false | false | false | false | false | false | false | false |
| \|PWY-8236\| | Salmonella enterica serotype O:39 O antigen biosynthesis | false | false | false | false | false | false | false | false | false | false | false | false | false | false | false | false | false |
| \|PWY-8237\| | Escherichia coli serotype O:1B/Salmonella enterica serotype O:42 O antigen biosynthesis | false | false | false | false | false | false | false | false | false | false | false | false | false | false | false | false | false |
| \|PWY-8238\| | 24-epi-campesterol, fucosterol, and clionasterol biosynthesis (diatoms) | false | false | false | false | false | false | false | false | false | false | false | false | false | false | false | false | false |
| \|PWY-8239\| | cholesterol biosynthesis (diatoms) | false | false | false | false | false | false | false | false | false | false | false | false | false | false | false | false | false |
| \|PWY-8241\| | Escherichia coli serotype O:71/Salmonella enterica serotype O:28ac O antigen biosynthesis | false | false | false | false | false | false | false | false | false | false | false | false | false | false | false | false | false |
| \|PWY-8242\| | Escherichia coli serotype O:51/Salmonella enterica serotype O:57 O antigen biosynthesis | false | false | false | false | false | false | false | false | false | false | false | false | false | false | false | false | false |
| \|PWY-8243\| | colanic acid (Escherichia coli K12) biosynthesis | false | false | false | false | false | false | false | false | false | false | false | false | false | false | false | false | false |
| \|PWY-8244\| | Escherichia coli serotype O:177 O antigen biosynthesis | false | false | false | false | false | false | false | false | false | false | false | false | false | false | false | false | false |
| \|PWY-8245\| | lipid IVA biosynthesis (P. gingivalis) | false | false | false | false | false | false | false | false | false | false | false | false | false | false | false | false | false |
| \|PWY-8246\| | Kdo transfer to lipid IVA (P. gingivalis) | false | false | false | false | false | false | false | false | false | false | false | false | false | false | false | false | false |
| \|PWY-8247\| | (Kdo)2-lipid A biosynthesis (P. gingivalis) | false | false | false | false | false | false | false | false | false | false | false | false | false | false | false | false | false |
| \|PWY-8248\| | lipid A-core biosynthesis (P. gingivalis) | false | false | false | false | false | false | false | false | false | false | false | false | false | false | false | false | false |
| \|PWY-8249\| | xanthommatin biosynthesis | false | false | false | false | false | false | false | false | false | false | false | false | false | false | false | false | false |

| | | | | | | | | | | | | | | | | | | |
|---|---|---|---|---|---|---|---|---|---|---|---|---|---|---|---|---|---|---|
| \|PWY-8250\| | Escherichia coli serotype O:9 O antigen biosynthesis | false | false | false | false | false | false | false | false | false | false | false | false | false | false | false | false | false |
| \|PWY-8251\| | Escherichia coli serotype O:2 O antigen biosynthesis | false | false | false | false | false | false | false | false | false | false | false | false | false | false | false | false | false |
| \|PWY-8252\| | Escherichia coli serotype O:50 O antigen biosynthesis | false | false | false | false | false | false | false | false | false | false | false | false | false | false | false | false | false |
| \|PWY-8253\| | Escherichia coli serotype O:149/Shigella boydii serotype O1 O antigen biosynthesis | false | false | false | false | false | false | false | false | false | false | false | false | false | false | false | false | false |
| \|PWY-8254\| | bile acids 7-O epimerization | false | false | false | false | false | false | false | false | false | false | false | false | false | false | false | false | false |
| \|PWY-8255\| | bile acids 3-O-epimerization | false | false | false | false | false | false | false | false | false | false | false | false | false | false | false | false | false |
| \|PWY-8256\| | bile acids 12-O-epimerization | false | false | false | false | false | false | false | false | false | false | false | false | false | false | false | false | false |
| \|PWY-8257\| | Escherichia coli serotype O:52 O antigen biosynthesis | false | false | false | false | false | false | false | false | false | false | false | false | false | false | false | false | false |
| \|PWY-8259\| | arsenic detoxification (plants) | false | false | false | false | false | false | false | false | false | false | false | false | false | false | false | false | false |
| \|PWY-8260\| | roxarsone degradation I | false | false | false | false | false | false | false | false | false | false | false | false | false | false | false | false | false |
| \|PWY-8261\| | roxarsone degradation II | false | false | false | false | false | false | false | false | false | false | false | false | false | false | false | false | false |
| \|PWY-8262\| | roxarsone (and nitarsone) degradation III | false | false | false | false | false | false | false | false | false | false | false | false | false | false | false | false | false |
| \|PWY-8263\| | arsenate detoxification III | false | false | false | false | false | false | false | false | false | false | false | false | false | false | false | false | false |
| \|PWY-8264\| | arsenate detoxification I | false | false | false | false | false | false | false | false | false | false | false | false | false | false | false | false | false |
| \|PWY-8265\| | roxarsone (and nitarsone) degradation IV | false | false | false | false | false | false | false | false | false | false | false | false | false | false | false | false | false |
| \|PWY-8267\| | methylarsonous acid detoxification I | false | false | false | false | false | false | false | false | false | false | false | false | false | false | false | false | false |
| \|PWY-8268\| | methylarsonous acid detoxification II | false | false | false | false | false | false | false | false | false | false | false | false | false | false | false | false | false |
| \|PWY-8269\| | methylarsonous acid detoxification III | false | false | false | false | false | false | false | false | false | false | false | false | false | false | false | false | false |
| \|PWY-8270\| | cyclic electron flow | false | false | false | false | false | false | false | false | false | false | false | false | false | false | false | false | false |
| \|PWY-8271\| | NADPH to cytochrome c oxidase via plastocyanin | false | false | false | false | false | false | false | false | false | false | false | false | false | false | false | false | false |
| \|PWY-8272\| | Escherichia coli serotype O:49 O antigen biosynthesis | false | false | false | false | false | false | false | false | false | false | false | false | false | false | false | false | false |
| \|PWY-8273\| | UDP-3-acetamido-2-amino-2,3-dideoxy-α-D-glucopyranose biosynthesis | false | false | false | false | false | false | false | false | false | false | false | false | false | false | false | false | false |
| \|PWY-8274\| | pyruvate fermentation to (R)-lactate | false | false | false | false | false | false | false | false | false | false | false | false | false | false | false | false | false |
| \|PWY-8275\| | pyruvate decarboxylation to acetyl CoA III | false | false | false | false | false | false | false | false | false | false | false | false | false | false | false | false | false |
| \|PWY-8279\| | palmitate biosynthesis III | false | false | false | false | false | false | false | false | false | false | false | false | false | false | false | false | false |
| \|PWY-8280\| | stearate biosynthesis IV | false | false | false | false | false | false | false | false | false | false | false | false | false | false | false | false | false |
| \|PWY-8281\| | 4-aminobenzoate biosynthesis III | false | false | false | false | false | false | false | false | false | false | false | false | false | false | false | false | false |
| \|PWY-8282\| | adenosylcobinamide-GDP salvage from assorted adenosylcobamides | false | false | false | false | false | false | false | false | false | false | false | false | false | false | false | false | false |
| \|PWY-8283\| | lipid IVA biosynthesis (generic) | false | false | false | false | false | false | false | false | false | false | false | false | false | false | false | false | false |
| \|PWY-8284-1\| | Kdo transfer to lipid IVA (Vibrio cholerae serogroup O1 El Tor) | false | false | false | false | false | false | false | false | false | false | false | false | false | false | false | false | false |
| \|PWY-8284\| | Kdo transfer to lipid IVA (generic) | false | false | false | false | false | false | false | false | false | false | false | false | false | false | false | false | false |
| \|PWY-8285\| | (Kdo)2-lipid A biosynthesis (generic) | false | false | false | false | false | false | false | false | false | false | false | false | false | false | false | false | false |
| \|PWY-8286\| | carbofuran degradation I | false | false | false | false | false | false | false | false | false | false | false | false | false | false | false | false | false |
| \|PWY-8287\| | carbofuran degradation II | false | false | false | false | false | false | false | false | false | false | false | false | false | false | false | false | false |
| \|PWY-8288\| | carbofuran degradation III | false | false | false | false | false | false | false | false | false | false | false | false | false | false | false | false | false |
| \|PWY-8291\| | L-aspartate degradation II (aerobic) | false | false | false | false | false | false | false | false | false | false | false | false | false | false | false | false | false |
| \|PWY-8292\| | trimethylamine N-oxide biosynthesis | false | false | false | false | false | false | false | false | false | false | false | false | false | false | false | false | false |
| \|PWY-8293\| | alazopeptin biosynthesis | false | false | false | false | false | false | false | false | false | false | false | false | false | false | false | false | false |
| \|PWY-8294\| | L-aspartate degradation III (anaerobic) | false | false | false | false | false | false | false | false | false | false | false | false | false | false | false | false | false |
| \|PWY-8295\| | nitrite biosynthesis | false | false | false | false | false | false | false | false | false | false | false | false | false | false | false | false | false |
| \|PWY-8296\| | cremeomycin biosynthesis | false | false | false | false | false | false | false | false | false | false | false | false | false | false | false | false | false |
| \|PWY-8297\| | 6-diazo-5-oxo-L-norleucine biosynthesis | false | false | false | false | false | false | false | false | false | false | false | false | false | false | false | false | false |
| \|PWY-8298\| | streptozotocin biosynthesis | false | false | false | false | false | false | false | false | false | false | false | false | false | false | false | false | false |
| \|PWY-8299\| | (3S)-diazinane-3-carboxylate biosynthesis | false | false | false | false | false | false | false | false | false | false | false | false | false | false | false | false | false |
| \|PWY-82\| | UDP-β-L-arabinose biosynthesis II (from β-L-arabinose) | false | false | false | false | false | false | false | false | false | false | false | false | false | false | false | false | false |
| \|PWY-8300\| | D-cycloserine inhibition of pyridoxal 5'-phosphate | false | false | false | false | false | false | false | false | false | false | false | false | false | false | false | false | false |
| \|PWY-8301\| | S-(6-hydroxy-4-methylhexan-4-yl)-L-cysteinylglycine biosynthesis | false | false | false | false | false | false | false | false | false | false | false | false | false | false | false | false | false |
| \|PWY-8302\| | S-(6-hydroxy-4-methylhexan-4-yl)-L-cysteinylglycine degradation | false | false | false | false | false | false | false | false | false | false | false | false | false | false | false | false | false |
| \|PWY-8303\| | reductive glycine pathway of autotrophic CO2 fixation | false | false | false | false | false | false | false | false | false | false | false | false | false | false | false | false | false |
| \|PWY-8304\| | methanogenesis from methoxylated aromatic compounds | false | false | false | false | false | false | false | false | false | false | false | false | false | false | false | false | false |
| \|PWY-8305\| | methoxylated aromatic compound degradation II | false | false | false | false | false | false | false | false | false | false | false | false | false | false | false | false | false |
| \|PWY-8307\| | L-carnitine degradation IV | false | false | false | false | false | false | false | false | false | false | false | true | false | false | false | false | false |
| \|PWY-8308\| | thiazole component of thiamine diphosphate biosynthesis IV | false | false | false | false | false | false | false | false | false | false | false | false | false | false | false | false | false |
| \|PWY-8314\| | inulin degradation | false | false | false | false | false | false | false | false | false | false | false | false | false | false | false | false | false |
| \|PWY-8317\| | cyclobis-(1→6)-α-nigerosyl biosynthesis | false | false | false | false | false | false | false | false | false | false | false | false | false | false | false | false | false |
| \|PWY-8318\| | L-fucose degradation III | false | false | false | false | false | false | false | false | false | false | false | false | false | false | false | false | false |
| \|PWY-8319\| | NiFe(CO)(CN)2 cofactor biosynthesis | false | false | false | false | false | false | false | false | false | false | false | false | false | false | false | false | false |
| \|PWY-8320\| | (3R)-N-[(2S)-1-hydroxy-6-[(3R)-3-isocyanobutanamido]hexan-2-yl]-3-isocyanobutanamide biosynthesis | false | false | false | false | false | false | false | false | false | false | false | false | false | false | false | false | false |
| \|PWY-8321\| | salicylate biosynthesis II | false | false | false | false | false | false | false | false | false | false | false | false | false | false | false | false | false |
| \|PWY-8322\| | protein N-glycosylation processing of proteins targeted for retention in cellular organelles (cis-Golgi, yeast) | false | false | false | false | false | false | false | false | false | false | false | false | false | false | false | false | false |
| \|PWY-8323\| | protein N-glycosylation processing of mannoproteins (cis-Golgi, yeast) | false | false | false | false | false | false | false | false | false | false | false | false | false | false | false | false | false |
| \|PWY-8324\| | D-tagatose degradation | false | false | false | false | false | false | false | false | false | false | false | false | false | false | false | false | false |
| \|PWY-8326\| | grayanate biosynthesis | false | false | false | false | false | false | false | false | false | false | false | false | false | false | false | false | false |
| \|PWY-8327\| | atranorin biosynthesis | false | false | false | false | false | false | false | false | false | false | false | false | false | false | false | false | false |
| \|PWY-8328\| | acetate and ATP formation from acetyl-CoA III | false | false | false | false | false | false | false | false | false | false | false | false | false | false | false | false | false |
| \|PWY-8329\| | L-arabinose degradation V | false | false | false | false | false | false | false | false | false | false | false | false | false | false | false | false | false |
| \|PWY-8330\| | D-xylose degradation VI | false | false | false | false | false | false | false | false | false | false | false | false | false | false | false | false | false |
| \|PWY-8331\| | D-arabinose degradation IV | false | false | false | false | false | false | false | false | false | false | false | false | false | false | false | false | false |
| \|PWY-8334\| | D-arabinose degradation V | false | false | false | false | false | false | false | false | false | false | false | false | false | false | false | false | false |
| \|PWY-8336\| | valinomycin biosynthesis | false | false | false | false | false | false | false | false | false | false | false | false | false | false | false | false | false |
| \|PWY-8337\| | γ-butyrobetaine degradation I | false | false | false | false | false | false | false | false | false | false | false | false | false | false | false | false | false |
| \|PWY-8339\| | γ-butyrobetaine degradation III | false | false | false | false | false | false | false | false | false | false | false | false | false | false | false | false | false |
| \|PWY-8340\| | isoorientin biosynthesis II | false | false | false | false | false | false | false | false | false | false | false | false | false | false | false | false | false |
| \|PWY-8341\| | N1-methyl-N3-aminocarboxypropyl-pseudouridine-modified rRNA biosynthesis | false | false | false | false | false | false | false | false | false | false | false | false | false | false | false | false | false |
| \|PWY-8342\| | coenzyme A biosynthesis III (archaea) | false | false | false | false | false | false | false | false | false | false | false | false | false | false | false | false | false |
| \|PWY-8343\| | leinamycin biosynthesis | false | false | false | false | false | false | false | false | false | false | false | false | false | false | false | false | false |
| \|PWY-8344\| | ansamitocin P biosynthesis | false | false | false | false | false | false | false | false | false | false | false | false | false | false | false | false | false |
| \|PWY-8345\| | malbrancheamide biosynthesis | false | false | false | false | false | false | false | false | false | false | false | false | false | false | false | false | false |
| \|PWY-8348\| | sulfoquinovose degradation IV | false | false | false | false | false | false | false | false | false | false | false | false | false | false | false | false | false |
| \|PWY-8349\| | sulfoquinovose degradation V | false | false | false | false | false | false | false | false | false | false | false | false | false | false | false | false | false |
| \|PWY-8350\| | sulfoquinovose degradation VI | false | false | false | false | false | false | false | false | false | false | false | false | false | false | false | false | false |
| \|PWY-8351\| | sulfoquinovosyl diacylglycerides and sulfoquinovosyl glycerol degradation | false | false | false | false | false | false | false | false | false | false | false | false | false | false | false | false | false |
| \|PWY-8352\| | NAD de novo biosynthesis III | false | false | false | false | false | false | false | false | false | false | false | false | false | false | false | false | false |
| \|PWY-8353\| | 3-thia-L-glutamate biosynthesis | false | false | false | false | false | false | false | false | false | false | false | false | false | false | false | false | false |
| \|PWY-8355\| | peptido-conjugates in tissue regeneration biosynthesis | false | false | false | false | false | false | false | false | false | false | false | false | false | false | false | false | false |
| \|PWY-8356\| | maresin biosynthesis | false | false | false | false | false | false | false | false | false | false | false | false | false | false | false | false | false |
| \|PWY-8357\| | protectin biosynthesis | false | false | false | false | false | false | false | false | false | false | false | false | false | false | false | false | false |
| \|PWY-8358\| | curacin A biosynthesis | false | false | false | false | false | false | false | false | false | false | false | false | false | false | false | false | false |
| \|PWY-8359\| | taurine biosynthesis III | false | false | false | false | false | false | false | false | false | false | false | false | false | false | false | false | false |
| \|PWY-8360\| | hectochlorin biosynthesis | false | false | false | false | false | false | false | false | false | false | false | false | false | false | false | false | false |
| \|PWY-8361\| | apratoxin A biosynthesis | false | false | false | false | false | false | false | false | false | false | false | false | false | false | false | false | false |
| \|PWY-8362\| | photorespiration II | false | false | false | false | false | false | false | false | false | false | false | false | false | false | false | false | false |
| \|PWY-8363\| | photorespiration III | false | false | false | false | false | false | false | false | false | false | false | false | false | false | false | false | false |
| \|PWY-8364\| | 7,8-dimethylmenaquinone biosynthesis | false | false | false | false | false | false | false | false | false | false | false | false | false | false | false | false | false |
| \|PWY-8365\| | C25,25 CDP-archaeol biosynthesis | false | false | false | false | false | false | false | false | false | false | false | false | false | false | false | false | false |
| \|PWY-8366\| | septacidin biosynthesis | false | false | false | false | false | false | false | false | false | false | false | false | false | false | false | false | false |
| \|PWY-8367\| | hygromycin B biosynthesis | false | false | false | false | false | false | false | false | false | false | false | false | false | false | false | false | false |
| \|PWY-8368\| | clorobiocin biosynthesis | false | false | false | false | false | false | false | false | false | false | false | false | false | false | false | false | false |
| \|PWY-8370\| | leporin B biosynthesis | false | false | false | false | false | false | false | false | false | false | false | false | false | false | false | false | false |
| \|PWY-8371\| | mannosylerythritol lipids biosynthesis | false | false | false | false | false | false | false | false | false | false | false | false | false | false | false | false | false |
| \|PWY-8372\| | erythritol biosynthesis I | false | false | false | false | false | false | false | false | false | false | false | false | false | false | false | false | false |
| \|PWY-8373\| | erythritol biosynthesis II | false | false | false | false | false | false | false | false | false | false | false | false | false | false | false | false | false |
| \|PWY-8374\| | antimycin biosynthesis | false | false | false | false | false | false | false | false | false | false | false | false | false | false | false | false | false |
| \|PWY-8375\| | benzene degradation IV (anaerobic) | false | false | false | false | false | false | false | false | false | false | false | false | false | false | false | false | false |
| \|PWY-8376\| | benzene degradation III (anaerobic) | false | false | false | false | false | false | false | false | false | false | false | false | false | false | false | false | false |
| \|PWY-8377\| | lactate fermentation to acetate, CO2 and hydrogen (Desulfovibrionales) | false | false | false | false | false | false | false | false | false | false | false | false | false | false | false | false | false |
| \|PWY-8378\| | Kdo-lipid A biosynthesis (Vibrio cholerae serogroup O1 El Tor) | false | false | false | false | false | false | false | false | false | false | false | false | false | false | false | false | false |
| \|PWY-8379\| | sibiromycin biosynthesis | false | false | false | false | false | false | false | false | false | false | false | false | false | false | false | false | false |
| \|PWY-8380\| | dTDP-sibirosamine biosynthesis | false | false | false | false | false | false | false | false | false | false | false | false | false | false | false | false | false |
| \|PWY-8381\| | sulfazecin biosynthesis | false | false | false | false | false | false | false | false | false | false | false | false | false | false | false | false | false |
| \|PWY-8382\| | diazepinomicin biosynthesis | false | false | false | false | false | false | false | false | false | false | false | false | false | false | false | false | false |
| \|PWY-8383\| | myxochelin A and B biosynthesis | false | false | false | false | false | false | false | false | false | false | false | false | false | false | false | false | false |
| \|PWY-8384\| | vitamin B6 degradation II | false | false | false | false | false | false | false | false | false | false | false | false | false | false | false | false | false |
| \|PWY-8386\| | benzene degradation II (aerobic) | false | false | false | false | false | false | false | false | false | false | false | false | false | false | false | false | false |
| \|PWY-8387\| | microcin C biosynthesis | false | false | false | false | false | false | false | false | false | false | false | false | false | false | false | false | false |
| \|PWY-8388\| | aspoquinolone biosynthesis | false | false | false | false | false | false | false | false | false | false | false | false | false | false | false | false | false |
| \|PWY-8389\| | penigequinolone biosynthesis | false | false | false | false | false | false | false | false | false | false | false | false | false | false | false | false | false |
| \|PWY-8390\| | D-glucuronate degradation III | false | false | false | false | false | false | false | false | false | false | false | false | false | false | false | false | false |
| \|PWY-8391\| | D-galacturonate degradation III | false | false | false | false | false | false | false | false | false | false | false | false | false | false | false | false | false |
| \|PWY-8392\| | D-arabinitol degradation II | false | false | false | false | false | false | false | false | false | false | false | false | false | false | false | false | false |
| \|PWY-8393\| | xylitol degradation II | false | false | false | false | false | false | false | false | false | false | false | false | false | false | false | false | false |
| \|PWY-8394\| | ribitol degradation II | false | false | false | false | false | false | false | false | false | false | false | false | false | false | false | false | false |
| \|PWY-8395\| | linoleate metabolites biosynthesis | false | false | false | false | false | false | false | false | false | false | false | false | false | false | false | false | false |
| \|PWY-8396\| | di-homo-γ-linolenate metabolites biosynthesis | false | false | false | false | false | false | false | false | false | false | false | false | false | false | false | false | false |
| \|PWY-8397\| | arachidonate metabolites biosynthesis | false | false | false | false | false | false | false | false | false | false | false | false | false | false | false | false | false |
| \|PWY-8398\| | α-linolenate metabolites biosynthesis | false | false | false | false | false | false | false | false | false | false | false | false | false | false | false | false | false |
| \|PWY-8399\| | icosapentaenoate metabolites biosynthesis | false | false | false | false | false | false | false | false | false | false | false | false | false | false | false | false | false |
| \|PWY-83\| | monolignol glucosides biosynthesis | false | false | false | false | false | false | false | false | false | false | false | false | false | false | false | false | false |
| \|PWY-8400\| | docosahexaenoate metabolites biosynthesis | false | false | false | false | false | false | false | false | false | false | false | false | false | false | false | false | false |
| \|PWY-8402\| | N-(1-deoxy-D-fructos-1-yl)-L-asparagine degradation | false | false | false | false | false | false | false | false | false | false | false | false | false | false | false | false | false |
| \|PWY-8407\| | fumagillin biosynthesis | false | false | false | false | false | false | false | false | false | false | false | false | false | false | false | false | false |
| \|PWY-8408\| | erinacine biosynthesis | false | false | false | false | false | false | false | false | false | false | false | false | false | false | false | false | false |
| \|PWY-8409\| | cyathin biosynthesis | false | false | false | false | false | false | false | false | false | false | false | false | false | false | false | false | false |
| \|PWY-8410\| | representative whole plasma arachidonate-derived icosanoids I | false | false | false | false | false | false | false | false | false | false | false | false | false | false | false | false | false |
| \|PWY-8411\| | representative whole plasma arachidonate-derived icosanoids II | false | false | false | false | false | false | false | false | false | false | false | false | false | false | false | false | false |
| \|PWY-8412\| | representative whole plasma DHA-derived icosanoids | false | false | false | false | false | false | false | false | false | false | false | false | false | false | false | false | false |
| \|PWY-8413\| | representative whole plasma EPA-derived icosanoids | false | false | false | false | false | false | false | false | false | false | false | false | false | false | false | false | false |
| \|PWY-8414\| | glycerol dibiphytanyl glycerol tetraether lipid biosynthesis | false | false | false | false | false | false | false | false | false | false | false | false | false | false | false | false | false |
| \|PWY-8415\| | macrocyclic archaeol phospholipid biosynthesis | false | false | false | false | false | false | false | false | false | false | false | false | false | false | false | false | false |
| \|PWY-8416\| | CDP-ribitol biosynthesis | false | false | false | false | false | false | false | false | false | false | false | false | false | false | false | false | false |
| \|PWY-8418\| | S-benzyl-L-cysteine degradation | false | false | false | false | false | false | false | false | false | false | false | false | false | false | false | false | false |
| \|PWY-8419\| | L-canavanine degradation II | false | false | false | false | false | false | false | false | false | false | false | false | false | false | false | false | false |
| \|PWY-8421\| | Helicobacter pylori 26695 lipopolysaccharide biosynthesis (final steps) | false | false | false | false | false | false | false | false | false | false | false | false | false | false | false | false | false |
| \|PWY-8422\| | Porphyromonas gingivalis lipopolysaccharide (O-LPS) biosynthesis (final steps) | false | false | false | false | false | false | false | false | false | false | false | false | false | false | false | false | false |
| \|PWY-8423\| | Salmonella typhimurium LT2 lipopolysaccharide biosynthesis (final steps) | false | false | false | false | false | false | false | false | false | false | false | false | false | false | false | false | false |
| \|PWY-8424\| | ascomycin biosynthesis | false | false | false | false | false | false | false | false | false | false | false | false | false | false | false | false | false |
| \|PWY-8425\| | (2S)-ethylmalonyl-CoA biosynthesis | false | false | false | false | false | false | false | false | false | false | false | false | false | false | false | false | false |
| \|PWY-8426\| | (2R)-methoxymalonyl-[acp] biosynthesis | false | false | false | false | false | false | false | false | false | false | false | false | false | false | false | false | false |
| \|PWY-8427\| | 2-allylmalonyl-CoA biosynthesis | false | false | false | false | false | false | false | false | false | false | false | false | false | false | false | false | false |
| \|PWY-8428\| | 2-O-(β-D-mannosyl-(1→2)-β-D-mannosyl)-bis(myo-inositol) 1,3'-phosphate biosynthesis | false | false | false | false | false | false | false | false | false | false | false | false | false | false | false | false | false |
| \|PWY-8429\| | 1-(sn-glycero-1-phospho)-1D-myo-inositol biosynthesis | false | false | false | false | false | false | false | false | false | false | false | false | false | false | false | false | false |
| \|PWY-842\| | starch degradation I | false | false | false | false | false | false | false | false | false | false | false | true | false | false | false | false | false |
| \|PWY-8430\| | 1-(sn-glycero-3-phospho)-1D-myo-inositol biosynthesis | false | false | false | false | false | false | false | false | false | false | false | false | true | true | true | true | true |
| \|PWY-8431\| | L-mimosine degradation | false | false | false | false | false | false | false | false | false | false | false | false | false | false | false | false | false |
| \|PWY-8432\| | 4-hydroxypyridine degradation | false | false | false | false | false | false | false | false | false | false | false | false | false | false | false | false | false |
| \|PWY-8434\| | cucurbitacin biosynthesis | false | false | false | false | false | false | false | false | false | false | false | false | false | false | false | false | false |
| \|PWY-8435\| | yeast mannan degradation | false | false | false | false | false | false | false | false | false | false | false | false | false | false | false | false | false |
| \|PWY-8436\| | mammalian high-mannose N-glycan degradation | false | false | false | false | false | false | false | false | false | false | false | false | false | false | false | false | false |
| \|PWY-8437\| | furaquinocin biosynthesis | false | false | false | false | false | false | false | false | false | false | false | false | false | false | false | false | false |
| \|PWY-8438\| | corallopyronin A biosynthesis | false | false | false | false | false | false | false | false | false | false | false | false | false | false | false | false | false |
| \|PWY-8439\| | poly-γ-D-glutamate biosynthesis | false | false | false | false | false | false | false | false | false | false | false | false | false | false | false | false | false |
| \|PWY-8440\| | nucleoside and nucleotide degradation (halobacteria) | false | false | false | false | false | false | false | false | false | false | false | false | false | false | false | false | false |
| \|PWY-8441\| | (4R)-4-[(E)-2-butenyl]-4-methyl-L-threonine biosynthesis | false | false | false | false | false | false | false | false | false | false | false | false | false | false | false | false | false |
| \|PWY-8442\| | hyperxanthone E biosynthesis | false | false | false | false | false | false | false | false | false | false | false | false | false | false | false | false | false |
| \|PWY-8443\| | cyclosporin A biosynthesis | false | false | false | false | false | false | false | false | false | false | false | false | false | false | false | false | false |
| \|PWY-8444\| | L-fucose degradation II | false | false | false | false | false | false | false | false | false | false | false | false | false | false | false | false | false |
| \|PWY-8445\| | puerarin degradation | false | false | false | false | false | false | false | false | false | false | false | false | false | false | false | false | false |
| \|PWY-8446\| | carminate degradation | false | false | false | false | false | false | false | false | false | false | false | false | false | false | false | false | false |
| \|PWY-8447\| | carminate biosynthesis | false | false | false | false | false | false | false | false | false | false | false | false | false | false | false | false | false |
| \|PWY-8449\| | mannopine and agropine biosynthesis | false | false | false | false | false | false | false | false | false | false | false | false | false | false | false | false | false |
| \|PWY-8450\| | pyruvate-derived opine biosynthesis I (bacteria) | false | false | false | false | false | false | false | false | false | false | false | false | false | false | false | false | false |
| \|PWY-8451\| | 2-oxoglutarate-derived opine biosynthesis | false | false | false | false | false | false | false | false | false | false | false | false | false | false | false | false | false |
| \|PWY-84\| | resveratrol biosynthesis | false | false | false | false | false | false | false | false | false | false | false | false | false | false | false | false | false |
| \|PWY-861\| | dhurrin biosynthesis | false | false | false | false | false | false | false | false | false | false | false | false | false | false | false | false | false |
| \|PWY-862\| | fructan degradation | false | false | false | false | false | false | false | false | false | false | false | false | true | true | true | false | true |
| \|PWY-881\| | trehalose biosynthesis II | false | false | false | false | false | false | false | false | false | false | false | false | false | false | false | false | false |
| \|PWY-882\| | L-ascorbate biosynthesis I (plants, L-galactose pathway) | false | false | false | false | false | false | false | false | false | false | false | false | false | false | false | false | false |
| \|PWYBJ2-1\| | bacillithiol biosynthesis | false | false | false | false | false | false | false | false | false | false | false | false | false | false | false | false | false |
| \|PWYBJ2-20\| | pulcherrimin biosynthesis | false | false | false | false | false | false | false | false | false | false | false | false | false | false | false | false | false |
| \|PWYBJ2-22\| | kanosamine biosynthesis II | false | false | false | false | false | false | false | false | false | false | false | false | false | false | false | false | false |
| \|PWYBJ2-25\| | S-(2-succinyl)-L-cysteine degradation | false | false | false | false | false | false | false | false | false | false | false | false | false | false | false | false | false |
| \|PWY-922\| | mevalonate pathway I (eukaryotes and bacteria) | false | false | false | false | false | false | false | false | false | false | false | false | false | false | false | false | false |
| \|PWY-981\| | salicylate biosynthesis III | false | false | false | false | false | false | false | false | false | false | false | false | false | false | false | false | false |
| \|PWYDQC-4\| | indole-3-acetate biosynthesis I | false | false | false | false | false | false | false | false | false | false | false | false | false | false | false | false | false |
| \|PWYG-321\| | mycolate biosynthesis | false | false | false | false | false | false | false | false | false | false | false | false | false | false | false | false | false |
| \|PWYI-14\| | lipid IVA biosynthesis (H. pylori) | false | false | false | false | false | false | false | false | false | false | false | false | false | false | false | false | false |
| \|PWYQT-4427\| | sulfoquinovosyl diacylglycerol biosynthesis | false | false | false | false | false | false | false | false | false | false | false | false | false | false | false | false | false |
| \|PWYQT-4429\| | CO2 fixation into oxaloacetate (anaplerotic) | false | false | false | false | false | false | false | false | false | false | false | false | false | false | false | false | false |
| \|PWYQT-4450\| | aliphatic glucosinolate biosynthesis, side chain elongation cycle | false | false | false | false | false | false | false | false | false | false | false | false | false | false | false | false | false |
| \|PWYQT-4471\| | glucosinolate biosynthesis from dihomomethionine | false | false | false | false | false | false | false | false | false | false | false | false | false | false | false | false | false |
| \|PWYQT-4472\| | glucosinolate biosynthesis from trihomomethionine | false | false | false | false | false | false | false | false | false | false | false | false | false | false | false | false | false |
| \|PWYQT-4473\| | glucosinolate biosynthesis from tetrahomomethionine | false | false | false | false | false | false | false | false | false | false | false | false | false | false | false | false | false |
| \|PWYQT-4474\| | glucosinolate biosynthesis from pentahomomethionine | false | false | false | false | false | false | false | false | false | false | false | false | false | false | false | false | false |
| \|PWYQT-4475\| | glucosinolate biosynthesis from hexahomomethionine | false | false | false | false | false | false | false | false | false | false | false | false | false | false | false | false | false |
| \|PWYQT-4476\| | indole glucosinolate activation (herbivore attack) | false | false | false | false | false | false | false | false | false | false | false | false | false | false | false | false | false |
| \|PWYQT-4477\| | indole glucosinolate activation (intact plant cell) | false | false | false | false | false | false | false | false | false | false | false | false | false | false | false | false | false |
| \|PYRIDNUCSAL-PWY\| | NAD salvage pathway I (PNC VI cycle) | false | false | false | false | false | false | false | false | false | false | false | false | false | false | false | false | false |
| \|PYRIDOXSYN-PWY\| | pyridoxal 5'-phosphate biosynthesis I | false | false | false | false | false | false | false | false | false | false | false | false | false | false | false | false | false |
| \|QUINATEDEG-PWY\| | quinate degradation I | false | false | false | false | false | false | false | false | false | false | false | false | false | false | false | false | false |
| \|REDCITCYC\| | TCA cycle VI (Helicobacter) | false | false | false | false | false | false | false | false | false | false | false | false | false | false | false | false | false |
| \|RHAMCAT-PWY\| | L-rhamnose degradation I | false | false | false | false | false | false | false | false | false | false | false | false | false | false | false | false | false |
| \|RIBITOLUTIL-PWY\| | ribitol degradation I | false | false | false | false | false | false | false | false | false | false | false | false | false | false | false | false | false |
| \|RIBOKIN-PWY\| | ribose phosphorylation | false | false | false | false | false | false | false | false | false | false | false | false | false | false | false | false | false |
| \|RIBOSYN2-PWY\| | flavin biosynthesis I (bacteria and plants) | false | false | false | false | false | false | false | false | false | false | false | false | false | false | false | false | false |
| \|RUMP-PWY\| | formaldehyde oxidation I | false | false | false | false | false | false | false | false | false | false | false | false | false | false | false | false | false |
| \|SALVADEHYPOX-PWY\| | adenosine nucleotides degradation II | false | false | false | false | false | false | false | false | false | false | false | false | false | false | false | false | false |
| \|SALVPURINE2-PWY\| | xanthine and xanthosine salvage | false | false | false | false | false | false | false | false | false | false | false | false | false | false | false | false | false |
| \|SERSYN-PWY\| | L-serine biosynthesis I | false | false | false | false | false | false | false | false | false | false | false | false | false | false | false | false | false |
| \|SHIKIMATEDEG-PWY\| | shikimate degradation I | false | false | false | false | false | false | false | false | false | false | false | false | false | false | false | false | false |
| \|SOPHOROSYLOXYDOCOSANOATE-DEG-PWY\| | sophorosyloxydocosanoate deacetylation | false | false | false | false | false | false | false | false | false | false | false | false | false | false | false | false | false |
| \|SOPHOROSYLOXYDOCOSANOATE-SYN-PWY\| | sophorolipid biosynthesis | false | false | false | false | false | false | false | false | false | false | false | false | false | false | false | false | false |
| \|SPHINGOLIPID-SYN-PWY\| | sphingolipid biosynthesis (yeast) | false | false | false | false | false | false | false | false | false | false | false | false | false | false | false | false | false |
| \|SUCROSEUTIL2-PWY\| | sucrose degradation VII (sucrose 3-dehydrogenase) | false | false | false | false | false | false | false | false | false | false | false | false | false | false | false | false | false |
| \|SUCUTIL-PWY\| | sucrose degradation I (sucrose phosphotransferase) | false | false | false | false | false | false | false | false | false | false | false | false | false | false | false | false | false |
| \|SULFMETII-PWY\| | assimilatory sulfate reduction II | false | false | false | false | false | false | false | false | false | false | false | false | false | false | false | false | false |
| \|SULFUROX-PWY\| | sulfur oxidation I (aerobic) | false | false | false | false | false | false | false | false | false | false | false | false | false | false | false | false | false |
| \|TEICHOICACID-PWY\| | poly(glycerol phosphate) wall teichoic acid biosynthesis | false | false | false | false | false | false | false | false | false | false | false | false | false | false | false | false | false |
| \|THIOREDOX-PWY\| | thioredoxin pathway | false | false | false | false | false | false | false | false | false | false | false | false | false | false | false | false | false |
| \|THIOSULFOX-PWY\| | thiosulfate oxidation I (to tetrathionate) | false | false | false | false | false | false | false | false | false | false | false | false | false | false | false | false | false |
| \|THRDLCTCAT-PWY\| | L-threonine degradation III (to methylglyoxal) | false | false | false | false | false | false | false | false | false | false | false | false | false | false | false | false | false |
| \|TOLSULFDEG-PWY\| | 4-toluenesulfonate degradation I | false | false | false | false | false | false | false | false | false | false | false | false | false | false | false | false | false |
| \|TOLUENE-DEG-2-OH-PWY\| | toluene degradation to 2-hydroxypentadienoate I (via o-cresol) | false | false | false | false | false | false | false | false | false | false | false | false | false | true | false | false | true |
| \|TOLUENE-DEG-3-OH-PWY\| | toluene degradation II (aerobic) (via 4-methylcatechol) | false | false | false | false | false | false | false | false | false | false | false | false | false | false | false | false | false |
| \|TOLUENE-DEG-4-OH-PWY\| | toluene degradation to 4-methylphenol | false | false | false | false | false | false | false | false | false | false | false | false | false | false | false | false | false |
| \|TOLUENE-DEG-CATECHOL-PWY\| | toluene degradation to benzoate | false | false | false | false | false | false | false | false | false | false | false | false | false | true | false | false | false |
| \|TOLUENE-DEG-DIOL-PWY\| | toluene degradation to 2-hydroxypentadienoate (via toluene-cis-diol) | false | false | false | false | false | false | false | false | false | false | false | false | false | false | false | false | false |
| \|TREDEGLOW-PWY\| | trehalose degradation I (low osmolarity) | false | false | false | false | false | false | false | false | false | false | false | false | false | false | false | false | false |
| \|TREHALOSESYN-PWY\| | trehalose biosynthesis III | false | false | false | false | false | false | false | false | false | false | false | false | false | false | false | false | false |
| \|TRESYN-PWY\| | trehalose biosynthesis I | false | false | false | false | false | false | false | false | false | false | false | false | true | true | true | true | true |
| \|TRIGLSYN-PWY\| | diacylglycerol and triacylglycerol biosynthesis | false | false | false | false | false | false | false | false | false | false | false | false | false | false | false | false | false |
| \|TRPCAT-PWY\| | L-tryptophan degradation I (via anthranilate) | false | false | false | false | false | false | false | false | false | false | false | false | false | false | false | false | false |
| \|TRPIAACAT-PWY\| | indole-3-acetate biosynthesis VI (bacteria) | false | false | false | false | false | false | false | false | false | false | false | false | false | false | false | false | false |
| \|TRPKYNCAT-PWY\| | L-tryptophan degradation IV (via indole-3-lactate) | false | false | false | false | false | false | false | false | false | false | false | false | false | false | false | false | false |
| \|TRPSYN-PWY\| | L-tryptophan biosynthesis | false | false | false | false | false | false | false | false | false | false | false | false | true | false | true | false | true |
| \|TRYPANOSYN-PWY\| | trypanothione biosynthesis | false | false | false | false | false | false | false | false | false | false | false | false | false | false | false | false | false |
| \|TYRFUMCAT-PWY\| | L-tyrosine degradation I | false | false | false | false | false | false | false | false | false | false | false | false | false | false | false | false | false |
| \|TYRSYN\| | L-tyrosine biosynthesis I | false | false | false | false | false | false | false | false | false | false | false | false | false | false | false | false | false |
| \|UDPNACETYLGALSYN-PWY\| | UDP-N-acetyl-D-glucosamine biosynthesis II | false | false | false | false | false | false | false | false | false | false | false | false | false | false | false | false | false |
| \|VALDEG-PWY\| | L-valine degradation I | false | false | false | false | false | false | false | false | false | false | false | false | true | true | true | true | false |
| \|XYLCAT-PWY\| | D-xylose degradation I | false | false | false | false | false | false | false | false | false | false | false | false | false | false | false | false | false |
| PWY1 | Oxidase test | false | false | false | false | false | false | false | false | false | false | false | false | false | false | false | false | false |
| PWY2 | Catalase test | false | false | false | false | false | false | false | false | false | false | false | false | false | false | false | false | false |
| PWY3 | leucyl aminopeptidase test | false | false | false | false | false | false | false | false | false | false | false | false | false | false | false | false | false |
| PWY4 | arginine deiminase test | false | false | false | false | false | false | false | false | false | false | false | false | true | false | true | true | false |
| 2FLHYD | 2-fucosyllactose degradation | false | false | false | false | false | false | false | false | false | false | false | false | true | true | true | true | true |
| LNT-DEGRADATION | Lacto-N-tetraose degradation | false | false | false | false | false | false | false | false | false | false | false | false | true | true | true | true | true |
| FUCDEGRA-NP | L-Fucose degradation (non-phosphorylating) | false | false | false | false | false | false | false | false | false | false | false | false | false | false | false | false | false |
| BIFIDOSHUNT2 | Bifidobacterium shunt II | false | false | false | false | false | false | false | false | false | false | false | false | true | true | true | true | true |
| MASS-SYNTHESIS | massetolide A biosynthesis | false | false | false | false | false | false | false | false | false | false | false | false | true | true | true | true | true |
| VISC-SYNTHESIS | viscosin biosynthesis | false | false | false | false | false | false | false | false | false | false | false | false | true | true | true | true | true |
| AGMA-TRANSP | Agmatine Transport | false | false | false | false | false | false | false | false | false | false | false | false | false | false | false | true | false |
| AGMA-PROD | Agmatine extracellular biosynthesis | false | false | false | false | false | false | false | false | false | false | false | false | true | true | true | true | true |
| FD-NADH-OXIDO | Ferredoxin:NAD+ oxidoreductase | false | false | false | false | false | false | false | false | false | false | false | false | true | true | true | true | true |
| BACTERIOFERRITIN-PWY | ferroxidase | false | false | false | false | false | false | false | false | false | false | false | false | true | true | true | true | true |
| SO4ASSIM-PWY2 | sulfate reduction I (assimilatory) | false | false | false | false | false | false | false | false | false | false | false | false | true | true | true | true | true |
| PWY-6151-GS | S-adenosyl-L-methionine cycle I | false | false | false | false | false | false | false | false | false | false | false | false | true | true | true | true | true |
| MQL-OXIDASE-PWY | menaquinol oxidase (cytochrom aa3-600) | false | false | false | false | false | false | false | false | false | false | false | false | true | true | true | true | true |
| MUC-GLYCAN_DEG | Mucus glycans degradation | false | false | false | false | false | false | false | false | false | false | false | false | true | true | true | true | true |
| AC-BUT-COATRANS | acetyl-CoA fermentation to butanoate clostridia | false | false | false | false | false | false | false | false | false | false | false | false | true | true | true | true | true |
| EXC-STARCH-N27 | Extracellular starch(27n) degradation | false | false | false | false | false | false | false | false | false | false | false | false | true | true | true | true | true |
| EXO-GALACTAN_DEG | Extracellular galactan(2n) degradation | false | false | false | false | false | false | false | false | false | false | false | false | true | true | true | true | true |
| FOLATE-SYN-GS1 | Folate biosynthesis from GTP and 4-aminobenzoate via dihydropterin-3-triphosphate | false | false | false | false | false | false | false | false | false | false | false | false | true | true | true | true | true |
| THIAZOLE-SYN-GS1 | thiazole biosynthesis I (facultative anaerobic bacteria) with tautomerase | false | false | false | false | false | false | false | false | false | false | false | false | true | true | true | true | true |
| F420-3-BIOSYN-ARCHAEA | F420 Biosynthesis until 3 glutamine residues | false | false | false | false | false | false | false | false | false | false | false | false | true | true | true | true | true |
| H4SPT-SYN | H4SPT-SYN | false | false | false | false | false | false | false | false | false | false | false | false | true | true | true | true | true |
| INULIN-DEGRADATION-EXO | Inulin degradation(11xFru,1xGlc) (extracellular) | false | false | false | false | false | false | false | false | false | false | false | false | true | true | true | true | true |
| INULIN-DEGRADATION | Inulin degradation(11xFru,1xGlc) (intracellular) | false | false | false | false | false | false | false | false | false | false | false | false | false | false | false | false | false |
| TYR-DEG-4HPAC | L-Tyrosine degradation to 4-hydroxyphenylacetate via tyramine | false | false | false | false | false | false | false | false | false | false | false | false | true | true | true | true | false |
| LWP-GS | Ljungdahl-Wood pathway (gapseq) | false | false | false | false | false | false | false | false | false | false | false | false | false | false | true | false | false |
| BAI-PWY | Bile acid 7alpha-dehydroxylation | false | false | false | false | false | false | false | false | false | false | false | false | true | true | true | true | true |
| QUERCETIN-ANAERO-DEG | Anaerobe quercetin degradation | false | false | false | false | false | false | false | false | false | false | false | false | false | false | false | true | false |
| PHLOROGLUCINOL-DEG-PWY | phloroglucinol degradation (anaerobic) | false | false | false | false | false | false | false | false | false | false | false | false | true | true | true | true | true |
| PWY-8086-GS | (S)-lactate fermentation to propanoate | false | false | false | false | false | false | false | false | false | false | false | false | true | false | false | true | true |
| ISOBILE-BIOSYN | iso-bile acids biosynthesis (NADH or NADPH dependent) | false | false | false | false | false | false | false | false | false | false | false | false | false | false | false | false | false |
| \|PWY-8017-2\| | L-tryptophan degradation XIII (reductive Stickland reaction) | false | false | false | false | false | false | false | false | false | false | false | false | true | true | true | true | true |
| \|PCEDEG-PWY2\| | tetrachloroethene degradation II | false | false | false | false | false | false | false | false | false | false | false | false | true | true | true | true | true |
| ARGSYN-SUCC-PWY | L-arginine biosynthesis via citrulline and succinylated intermediates | false | false | false | false | false | false | false | false | false | false | false | false | false | false | false | true | false |
| \|PWY-6753\| | S-methyl-5'-thioadenosine degradation III | false | false | false | true | false | false | false | false | false | false | false | false | true | true | true | true | true |
| \|PWY-1263\| | taurine degradation I | false | false | false | false | false | false | false | false | false | true | false | false | true | true | true | true | true |
| \|PWY-1781\| | β-alanine degradation II | false | false | false | false | false | false | false | false | false | true | false | false | true | true | true | true | true |
| \|PWY-3\| | putrescine degradation V | false | false | false | false | false | false | false | false | false | true | false | false | true | true | true | true | true |
| \|PWY-5514\| | UDP-N-acetyl-D-galactosamine biosynthesis II | false | false | false | false | false | false | false | false | false | true | false | false | true | true | true | true | true |
| \|PWY-7241\| | myo-inositol degradation II | false | false | false | false | false | false | false | false | false | true | false | false | false | false | false | true | false |
| \|PWY-7387\| | hypotaurine degradation | false | false | false | false | false | false | false | false | false | true | false | false | false | false | false | false | false |
| \|PWY-1881\| | formate oxidation to CO2 | false | false | false | false | false | false | false | false | true | false | false | false | true | true | true | true | true |
| \|PWY-5026\| | indole-3-acetate biosynthesis V (bacteria and fungi) | false | false | false | false | false | false | false | false | true | false | false | false | true | true | true | true | true |
| \|PWY-7309\| | acrylonitrile degradation II | false | false | false | false | false | false | false | false | true | false | false | false | false | false | false | false | false |
| \|P185-PWY\| | formaldehyde assimilation III (dihydroxyacetone cycle) | false | false | false | false | false | false | false | true | false | false | false | false | true | true | true | true | true |
| \|P21-PWY\| | pentose phosphate pathway (partial) | false | false | false | false | false | false | false | true | false | false | true | false | false | true | false | true | false |
| \|PWY-8178\| | pentose phosphate pathway (non-oxidative branch) II | false | false | false | false | false | false | false | true | false | false | true | false | false | false | false | false | false |
| MQL6-NADH-OXIDO | NADH:menaquinone6 oxidoreductase | false | false | false | false | false | false | false | true | false | false | false | false | false | false | false | true | false |
| \|P141-PWY\| | atrazine degradation I (aerobic) | false | false | false | false | false | false | false | true | false | true | false | false | true | true | true | true | true |
| \|DETOX1-PWY\| | superoxide radicals degradation | false | false | false | false | false | false | true | false | false | false | false | false | false | false | false | false | false |
| \|PWY-7220\| | adenosine deoxyribonucleotides de novo biosynthesis II | false | false | false | false | false | false | true | false | false | true | false | false | true | true | true | true | true |
| \|PWY-7222\| | guanosine deoxyribonucleotides de novo biosynthesis II | false | false | false | false | false | false | true | false | false | true | false | false | true | true | true | true | true |
| \|CENTFERM-PWY\| | pyruvate fermentation to butanoate | false | false | false | false | false | false | true | true | false | true | false | false | true | true | true | true | true |
| \|PWY0-1261\| | peptidoglycan recycling I | false | false | false | false | false | false | true | true | false | true | false | false | true | true | true | true | true |
| \|PWY-1042\| | glycolysis IV | false | false | false | false | false | false | true | true | false | true | false | false | false | false | false | true | false |
| \|PWY-5983\| | trehalose biosynthesis VI | false | false | false | false | false | false | true | true | false | true | false | false | true | true | true | true | true |
| \|PWY-5985\| | trehalose biosynthesis VII | false | false | false | false | false | false | true | true | false | true | false | false | false | false | false | true | false |
| \|PWY0-1312\| | acetate and ATP formation from acetyl-CoA I | false | false | false | false | false | false | true | true | true | true | false | false | false | false | false | true | false |
| \|PWY-5482\| | pyruvate fermentation to acetate II | false | false | false | false | false | false | true | true | true | true | false | false | true | true | true | true | true |
| \|PWY-8335\| | plasmalogen biosynthesis II (anaerobic) | false | false | false | false | false | false | true | true | true | true | false | false | true | true | true | true | true |
| \|ANAPHENOXI-PWY\| | L-phenylalanine degradation II (anaerobic) | false | false | false | false | true | false | false | false | false | false | false | false | true | true | true | true | true |
| \|PWY0-1280\| | ethylene glycol degradation | false | false | false | false | true | false | false | false | false | false | false | false | true | true | true | true | true |
| \|PWY-5188\| | tetrapyrrole biosynthesis I (from glutamate) | false | false | false | false | true | false | false | false | false | false | false | false | true | true | true | true | true |
| \|PWY-5372\| | carbon tetrachloride degradation II | false | false | false | false | true | false | false | false | false | false | false | false | false | false | false | false | false |
| \|PWY-6931\| | seleno-amino acid detoxification and volatilization I | false | false | false | false | true | false | false | false | false | false | false | false | true | true | true | true | true |
| \|PWY-7330\| | UDP-N-acetyl-β-L-fucosamine biosynthesis | false | false | false | false | true | false | false | false | false | false | false | false | true | false | true | true | true |
| \|PWY-7467\| | UDP-2-acetamido-4-amino-2,4,6-trideoxy-α-D-galactose biosynthesis | false | false | false | false | true | false | false | false | false | false | false | false | true | true | true | true | true |
| \|PWY-8001\| | felinine and 3-methyl-3-sulfanylbutan-1-ol biosynthesis | false | false | false | false | true | false | false | false | false | false | false | false | true | true | true | true | true |
| \|PWY-8159\| | dipyrromethane cofactor biosynthesis | false | false | false | false | true | false | false | false | false | false | false | false | true | true | true | true | true |
| \|SORBDEG-PWY\| | D-sorbitol degradation II | false | false | false | false | true | false | false | false | false | false | false | false | true | true | true | true | true |
| \|ASPARAGINE-BIOSYNTHESIS\| | L-asparagine biosynthesis I | false | false | false | true | true | false | false | false | false | false | false | false | true | true | true | true | true |
| \|P283-PWY\| | hydrogen oxidation I (aerobic) | false | false | false | true | true | false | false | false | false | false | false | false | false | false | true | true | false |
| \|PWY0-1300\| | 2-O-α-mannosyl-D-glycerate degradation | false | false | false | true | true | false | false | false | false | false | false | false | true | true | true | true | true |
| \|PWY-5690\| | TCA cycle II (plants and fungi) | false | false | false | true | true | false | false | false | false | false | false | false | true | true | true | true | true |
| \|PWY-6543\| | 4-aminobenzoate biosynthesis I | false | false | false | true | true | false | false | false | false | false | false | false | true | true | true | true | true |
| \|PWY-8060\| | 2-deoxy-D-ribose degradation I | false | false | false | true | true | false | false | false | false | false | false | true | true | true | true | true | true |
| \|PWY-8276\| | 4-aminobenzoate biosynthesis II | false | false | false | true | true | false | false | false | false | false | false | false | true | false | true | false | false |
| \|TCA\| | TCA cycle I (prokaryotic) | false | false | false | true | true | false | false | false | false | false | false | false | true | true | true | true | true |
| \|BETA-ALA-DEGRADATION-I-PWY\| | β-alanine degradation I | false | false | false | false | true | false | false | false | false | true | false | false | false | false | false | false | false |
| \|PWY-3722\| | glycine betaine biosynthesis II (Gram-positive bacteria) | false | false | false | false | true | false | false | false | false | true | false | false | false | false | false | true | false |
| \|PWY-5332\| | sulfur reduction I | false | false | false | true | true | false | false | false | false | true | true | false | false | true | true | true | true |
| \|PWY-5364\| | sulfur reduction II (via polysulfide) | false | false | false | true | true | false | false | false | false | true | true | true | true | true | true | true | true |
| \|PWY-8214\| | 3-sulfopropanediol degradation II | false | false | false | true | true | false | false | false | false | true | false | false | true | true | true | true | true |
| \|PWY-6321\| | homotaurine degradation | false | false | false | false | true | false | false | true | false | false | false | false | true | true | true | true | true |
| \|FORMASS-PWY\| | formaldehyde oxidation IV (thiol-independent) | false | false | false | false | true | false | true | true | false | true | false | false | true | true | true | true | true |
| \|PWY-7269\| | mitochondrial NADPH production (yeast) | false | false | false | false | true | false | true | true | false | true | false | false | true | true | true | true | true |
| \|PWY-7828\| | cyclobis-(1→6)-α-nigerosyl degradation | false | false | false | false | false | true | false | false | false | false | false | false | true | true | true | true | true |
| \|PWY-2723\| | trehalose degradation V | false | false | false | false | false | true | false | false | false | true | false | false | true | true | true | true | true |
| \|PWY-3982\| | uracil degradation I (reductive) | false | false | false | false | false | true | false | false | true | false | true | false | true | false | true | true | true |
| \|PWY-6430\| | thymine degradation | false | false | false | false | false | true | false | false | true | false | true | false | true | true | true | true | true |
| \|PWY-5109\| | propanoate fermentation to 2-methylbutanoate | false | false | false | false | false | true | false | true | false | true | false | false | true | false | false | false | false |
| \|ACETOACETATE-DEG-PWY\| | acetoacetate degradation (to acetyl CoA) | false | false | false | false | false | true | true | false | false | true | false | false | true | false | false | false | false |
| \|PWY-6348\| | phosphate acquisition | false | false | false | false | false | true | true | false | true | true | false | false | false | false | false | false | false |
| \|NADPHOS-DEPHOS-PWY\| | NAD phosphorylation and dephosphorylation | false | false | false | false | false | true | true | false | true | true | false | false | false | false | false | true | false |
| \|GLUAMCAT-PWY\| | N-acetylglucosamine degradation I | false | false | false | false | false | true | true | true | false | true | true | false | true | true | true | true | true |
| \|P163-PWY\| | L-lysine fermentation to acetate and butanoate | false | false | false | false | false | true | true | true | false | true | false | false | true | true | true | true | true |
| \|PWY-5177\| | glutaryl-CoA degradation | false | false | false | false | false | true | true | true | false | true | false | false | false | false | false | false | false |

| | | | | | | | | | | | | | | | | | | |
|---|---|---|---|---|---|---|---|---|---|---|---|---|---|---|---|---|---|---|
| \|PWY-8347\| | adipate biosynthesis | false | false | false | false | false | true | true | true | false | true | false | false | true | true | true | true | false |
| \|PWY-8354\| | adipate degradation | false | false | false | false | false | true | true | true | false | true | false | false | false | false | false | true | false |
| \|HISHP-PWY\| | L-histidine degradation VI | false | false | false | false | false | true | true | true | true | true | false | false | true | true | true | true | true |
| \|NADPHOS-DEPHOS-PWY-1\| | NAD phosphorylation and transhydrogenation | false | false | false | false | false | true | true | true | true | true | true | false | true | true | true | true | true |
| \|PWY-5030\| | L-histidine degradation III | false | false | false | false | false | true | true | true | true | true | false | false | true | true | true | true | true |
| \|TRYPDEG-PWY\| | L-tryptophan degradation II (via pyruvate) | false | false | false | false | false | true | true | true | true | true | true | false | true | true | true | true | true |
| \|PWY66-366\| | flavin salvage | false | false | false | true | false | true | true | true | true | true | true | false | false | true | false | false | false |
| \|PWY-7167\| | choline degradation III | false | false | false | true | true | true | true | true | true | true | false | false | true | true | true | true | true |
| \|ILEUSYN-PWY\| | L-isoleucine biosynthesis I (from threonine) | false | false | true | false | false | false | false | false | false | false | false | false | true | true | true | true | true |
| \|PWY-4861\| | UDP-α-D-galacturonate biosynthesis I (from UDP-D-glucuronate) | false | false | true | false | false | false | false | false | false | false | false | false | true | false | true | false | false |
| \|PWY-7346\| | UDP-α-D-glucuronate biosynthesis (from UDP-glucose) | false | false | true | false | false | false | false | false | false | false | false | false | true | true | true | true | true |
| \|PWY-5982\| | sulfoacetaldehyde degradation II | false | false | true | false | true | false | false | false | false | false | false | false | true | true | true | true | true |
| \|PWY-7117\| | C4 photosynthetic carbon assimilation cycle, PEPCK type | false | false | true | false | true | false | false | false | false | false | false | false | true | true | true | true | true |
| \|GLUTDEG-PWY\| | L-glutamate degradation II | false | false | true | true | true | false | false | false | false | false | false | false | false | false | false | true | false |
| \|PWY-6163\| | chorismate biosynthesis from 3-dehydroquinate | false | false | true | true | true | false | false | false | false | false | false | false | true | false | true | true | true |
| \|PWY-701\| | L-methionine degradation II | false | false | true | true | true | false | false | false | false | false | false | false | false | false | false | false | false |
| \|PWY-702\| | L-methionine biosynthesis II | false | false | true | true | true | false | false | false | false | false | false | false | true | true | true | true | true |
| \|PUTDEG-PWY\| | putrescine degradation I | false | false | true | false | true | false | false | true | false | true | false | false | true | true | true | true | true |
| \|PWY-7568\| | L-galactose degradation | false | false | true | false | true | false | true | true | false | true | false | false | true | true | true | true | true |
| \|PWY-6936\| | seleno-amino acid biosynthesis (plants) | false | false | true | true | true | false | true | true | false | true | true | false | true | true | true | true | true |
| \|CITRULLINE-DEG-PWY\| | L-citrulline degradation | false | false | true | false | false | true | false | true | true | true | false | false | true | true | true | true | true |
| \|ARGDEGRAD-PWY2\| | L-arginine degradation V (arginine deiminase pathway) | false | false | true | false | false | true | false | true | true | true | false | false | false | false | false | false | false |
| \|PWY-8448\| | D-alanine degradation | false | false | true | false | false | true | true | false | false | true | false | false | true | true | true | true | true |
| \|PWY-6828\| | linezolid resistance | false | false | true | true | false | true | true | true | true | true | false | false | true | true | true | true | true |
| \|TAURINEDEG-PWY\| | taurine degradation III | false | false | true | true | true | true | false | true | true | true | false | false | true | true | true | true | true |
| PWY-6559-2 | spermidine biosynthesis II (only one key enzyme) | false | false | true | true | true | true | true | true | true | true | true | false | true | true | true | true | true |
| \|PWY3O-4107\| | NAD salvage pathway V (PNC V cycle) | false | true | false | false | false | false | false | false | false | false | false | false | true | true | true | true | true |
| \|PWY4FS-2\| | phosphatidylcholine biosynthesis II | false | true | false | false | false | false | false | false | false | false | false | false | true | true | true | true | true |
| \|PWY-7331\| | UDP-N-acetyl-β-L-quinovosamine biosynthesis | false | true | false | false | false | false | false | false | false | false | false | false | true | true | true | true | true |
| \|PWY-8306\| | methoxylated aromatic compound degradation I | false | true | false | false | false | false | false | false | false | false | false | false | true | true | true | true | true |
| \|PWY66-389\| | phytol degradation | false | true | false | false | true | false | false | false | false | false | false | false | true | true | true | true | true |
| \|PWY-7090\| | UDP-2,3-diacetamido-2,3-dideoxy-α-D-mannuronate biosynthesis | false | true | false | false | true | false | false | false | false | false | true | false | true | true | true | true | true |
| \|PWY-8215\| | 3-sulfopropanediol degradation III | false | true | false | true | true | true | true | true | true | true | false | false | true | true | true | true | true |
| \|PWY-8266\| | arsenite methylation (bacterial) | false | true | true | true | false | false | false | false | false | false | false | false | true | true | true | true | true |
| \|PWY-6281\| | L-selenocysteine biosynthesis II (archaea and eukaryotes) | false | true | true | true | true | false | false | false | false | false | false | false | false | false | false | false | false |
